# Mark loss can strongly bias demographic rates in multi-state models: a case study with simulated and empirical datasets

**DOI:** 10.1101/2022.03.25.485763

**Authors:** Frédéric Touzalin, Eric J. Petit, Emmanuelle Cam, Claire Stagier, Emma C. Teeling, Sébastien J. Puechmaille

**Affiliations:** School of Biology and Environmental Science, Science Centre West, University College Dublin, Dublin, Ireland; Bretagne Vivante-SEPNB, Brest, France; DECOD (Ecosystem Dynamics and Sustainability), INRAE, Institut Agro, Ifremer, Rennes, France; Université de Bretagne occidentale, Brest, LEMAR, CNRS, IRD, Ifremer, F-29280 Plouzane, France; Zoological Institute and Museum, University of Greifswald, Greifswald, Germany; ISEM, Univ Montpellier, CNRS, IRD, Montpellier, France; Institut Universitaire de France, Paris, France

**Keywords:** Arnason-Schwarz model, Bayesian, bats, capture-mark-recapture, mark retention, *Myotis myotis*, multi-state, surgical glue

## Abstract

1. The development of methods for individual identification in wild species and the refinement of Capture-Mark-Recapture (CMR) models over the past few decades have greatly improved the assessment of population demographic rates to address ecological and conservation questions. In particular, multi-state models, which offer flexibility in analysing complex study systems, have gained popularity within the ecological community. In this study, we focus on the issue of mark loss and the associated recycling of remarked individuals, which requires further exploration given the increasing use of these models.
2. To fill this knowledge gap, we employed a wide range of simulation scenarios that reflect commonly encountered real case studies, drawing inspiration from the survival rates of 700 vertebrate species. Using a multi-state, Arnason-Schwartz (AS) modelling framework, we estimated the effects of mark loss and recycled individuals on parameter estimates. We assessed parameter bias by simulating a metapopulation system with varying capture and survival rates. Additionally, we demonstrated how mark loss can be easily estimated and accounted for using a 10-year empirical CMR dataset of bats. The bats were individually identified using Passive Integrated Transponder (PIT) tag technology as potentially lost marks and multi-locus genotypes as ’permanent marks’.
3. Our simulation results revealed that the occurrence of bias and the affected parameters were highly dependent on the study system, making it difficult to establish general rules to predict bias *a priori*. The model structure and the interdependency among parameters pose challenges in predicting the impact of bias on estimates.
4. Our findings underscore the importance of assessing the effect of mark loss when using AS models. Ignoring such violations of model assumptions can have significant implications for ecological inferences and conservation policies. In general, the use of permanent marks, such as genotypes, should always be preferred when modelling population dynamics. If that is not feasible, an alternative is to combine two independent types of temporary marks, such as PIT tags and bands.
5. Analysis of our empirical dataset on *Myotis myotis* bats revealed that tag loss is higher in juveniles than in adults during the first year after tagging. The use of surgical glue to close the injection hole reduces tag loss rate from 28% to 19% in juveniles, while it has no effect on the tag loss rate in adults (∼10%). The main bias observed in our metapopulation system appears in the survival rate, with up to a 20% underestimation if tag loss is not accounted for.

## 1 Introduction

Capture-mark-recapture (CMR) methods have become a standard approach for estimating demographic rates of wild species (Johnson *et al*., 1986; Williams *et al*., 2002). Accurately quantifying population dynamic parameters is crucial for assessing population status, understanding dynamics, and making effective management and conservation decisions. However, CMR models rely on certain assumptions of homogeneity (Johnson *et al*., 1986; Williams *et al*., 2002) that can introduce biases if violated. One common violation of CMR assumptions, first identified four decades ago (Nelson *et al*., 1980), is the loss of marks (see Supporting Information 1, Table S2). Mark loss has two implications: (1) when marks are shed, it leads to non-identifiability of individuals (detection heterogeneity), causing them to be considered dead or out of the study area, even when they are alive and present; (2) if these individuals are recaptured, they will go unrecognized and be counted as newly recruited individuals, referred to as “recycled” individuals (Malcolm-White *et al*., 2020). In open population models, estimates of abundance in the Jolly-Seber (JS) model (Jolly, 1965; Seber, 1965) or survival in the Cormac-Jolly-Seber (CJS) model (Cormack, 1964; Jolly, 1965; Seber, 1965) can be affected by mark loss (Arnason & Mills, 1981). Several statistical techniques have been developed to mitigate the confounding effect of mark loss in these models (Arnason & Mills, 1981; Cowen & Schwarz, 2006; Robson & Regier, 1966; Seber & Felton, 1981). However, the impact of mark loss on state transition in multi-state models, which have undergone significant development (Lebreton *et al*., 2009), has not been extensively studied.

In multi-state models, if survival is state dependent, it is the product of true survival and mark retention rate for individuals in a specific state (Lebreton *et al*., 1992; Lebreton *et al*., 2009). If the retention rate drops below one without being considered in the model, the estimation of survival in a particular state is underestimated and becomes confounded with the probability of presence of the mark. This is especially true if true survival is high, but it is unclear how state transitions are affected.

Let’s consider an example encounter history: “1011”, where “1” indicates the individual was captured and “0” indicates otherwise. If we denote *ϕ*_t_ as the survival rate between occasions t and t+1, and *p*_t_ as the capture probability at occasion t (with *q*_t_ = 1 - *p*_t_), this encounter history occurs with probability *ϕ*_1_*q*_2_*ϕ*_2_*p*_3_*ϕ*_3_*p*_4_. We can break down this probability product as follows: the individual survives between *t*_1_ and *t*_2_ (*ϕ*_1_), was not captured in *t*_2_ (*q*_2_), survives between *t*_2_ and *t*_3_ (*ϕ*_2_), was captured in *t*_3_ (*p*_3_), and finally survives between *t*_3_ and *t*_4_ (*ϕ*_3_) and was captured in *t*_4_ (*p*_3_). Now, let’s consider that the individual can transition between two states, and its history becomes “1022”, with the individual in state “1” at *t*_1_ and in state “2” at *t*_3_ and *t*_4_. At *t*_2_, when the individual was not detected, two possibilities arise: either it stayed in state 1 or made a transition to state 2. To account for these two possible histories, we introduce *ψ*^i,j^ as the transition probability from state *i* to state *j* (where *i* and *j* are in {1,2}), conditional on survival. The new encounter history is now the sum of two components o account for the two possible his histories “1122” or “1222”: *ϕ*_1_^(1)^*ψ*^*(1,1)*^*q*_2_^(1)^*ϕ*_2_^(2)^*ψ*^*(1,2)*^*p*_3_^(2)^*ϕ*_3_^(2)^*ψ*^(2,2)^*p*_4_^(2)^ + *ϕ*_1_^(1)^*ψ*^*(1,2)*^*q*_2_^(2)^*ϕ*_2_^(2)^*ψ*^*(2,2)*^*p*_3_^(2)^*ϕ*_3_^(2)^*ψ*^*(2,2)*^*p*_4_^(2)^, with the superscript denoting state-specific parameters in parentheses. However, if this individual loses its mark after its first capture (indicated when recaptured at *t*_3_ and not recognized), its encounter history becomes two different histories from two different individuals: one becomes “1000”, and the second becomes “0022”. In this case, survival and mark loss patterns differ. Not only is survival underestimated (at least for the “first” history), but the transition probabilities are also underestimated, since there is no longer a change of state (the second history starts directly at state 2).

Most studies using CMR techniques are affected by mark loss, which can vary based on several factors such as species (see Supporting Information 1, Table S2), mark type (Smout *et al*., 2011a), sex (Conn *et al*., 2004), mass (Schwarz *et al*., 2012), size (Acolas *et al*., 2007), mark location (Kaemingk *et al*., 2011), or physiological stage (Besnard *et al*., 2007). This mark loss has been previously shown to introduce negative bias in survival estimates and detection (Nichols *et al*., 1992; Nichols & Hines, 1993). To address situations where an individual’s “state” (e.g., location, behaviour, physiology, reproductive or social status) may affect its survival or detection probability, and where the individual can change “state” during its lifespan, multi-state models have been developed (reviewed in Lebreton *et al*., 2009). These models have gained popularity due to their flexibility in studying a wide range of ecological systems and biological questions. Biologists can easily use user-friendly software like Mark (White & Burnham, 1999), WinBUGS (Lunn *et al*., 2000), JAGS (Plummer, 2003), E-SURGE (Choquet, *et al*., 2009b), MultiBUGS (Goudie *et al*., 2020), NIMBLE (de Valpine *et al*., 2017), and STAN (STAN Development Team, 2022) to implement these models. They are used to examine ecological and evolutionary hypotheses related to variations in life history traits (state transitions) over an individual’s lifespan (Nichols & Kendall, 1995; see also Cam 2009 for a detailed discussion), density dependence effects (Schofield & Barker, 2008), co-evolution (Benkman *et al*., 2005), dispersal probability among subpopulations or habitats (Hestbeck *et al*., 1991; Spendelow *et al*., 1995), and disease prevalence in wild populations (Jennelle *et al*., 2007). However, there is limited literature on the impact of mark loss on the behaviour of multi-state models, and further exploration is needed in this area (Seber & Schofield, 2019).

To bridge this gap, we used simulation-based Arnason-Schwartz (AS) model approaches (Arnason, 1972, 1973; Schwarz *et al*., 1993) to investigate the impact of mark loss on estimates of model parameters within a Bayesian framework. With the increasing utilization of multi-state models, our aim is to evaluate the potential bias in the marginal posterior distributions of demographic parameter estimates using a metapopulation context, based on biologically realistic scenarios, and provide comprehensive guidelines for both fieldwork and data analyses. The AS model shares assumptions with the CJS model, specifically regarding mark loss, but additionally assumes that states are recorded without error. Similarly to the CJS model, we anticipated that the AS model might underestimate survival and transition probabilities in the event of tag loss and recycling (Nichols & Hines, 1993). Since transition probabilities are conditional on survival and detection of state in our AS model, we expected estimation errors to have varying effects on model parameters depending on the state transition rate.

To illustrate our approach with an empirical example, we used our decade-long mark recapture dataset of PIT-tagged and genotyped greater mouse-eared bat (*Myotis myotis*), a taxonomic group particularly vulnerable to PIT-tag loss (Freeland & Fry, 1995). We employed genotypes as individual permanent marks to estimate the bias between models that account for mark loss and recycling versus those that do not, and finally, we suggest recommendations for future studies.

## 2 Material and method

To quantify the potential bias induced by mark loss on parameter estimates in the AS framework, we defined various scenarios based on representative situations derived from a compilation data obtained from the Demographic Species Knowledge Index (Conde *et al*., 2019) and literature sources for fish and bat species (Fig. 1, Supporting Information 1, Table S1). These data were limited to published CMR studies or data from controlled conditions, specifically marked individuals with known outcomes, such as those observed in zoos. Among the six vertebrate classes (Actinopterygii, Chondrichthyes, Amphibia, Aves, Reptilia, and Mammalia) encompassing 700 species, the survival rates exhibit a wide range of values (Fig. 1). The relationship between adult and juvenile survival, available for 143 species (Supporting Information 1, Fig. S1), reveals that low adult survival is associated with low juvenile survival, while high adult survival can be accompanied by a diverse range of juvenile survival rates. To limit the number of scenarios to explore, we selected values towards the extremes. Consequently, we considered two hypothetical populations: one consisting of long-lived species with high survival rates in both age classes (e.g., large mammals), and the other consisting of short-lived species with low survival rates in both age classes (e.g., amphibians). For each population, we explored cases with high and low detection probabilities, and for each case, we tested three different mark loss rates, obtained from relevant literature sources (Supporting Information 1, Table S2). Initially, we present the procedure for generating our simulated scenarios, followed by the description of the two different models used to analyse these data: one model that does not account for mark loss, and another model that incorporates mark loss by utilizing a second permanent mark, allowing for comparison of estimations. Lastly, we outline the metrics used to evaluate the potential bias in parameter estimates when mark loss is not accounted for in the AS framework.

**Figure 1:**
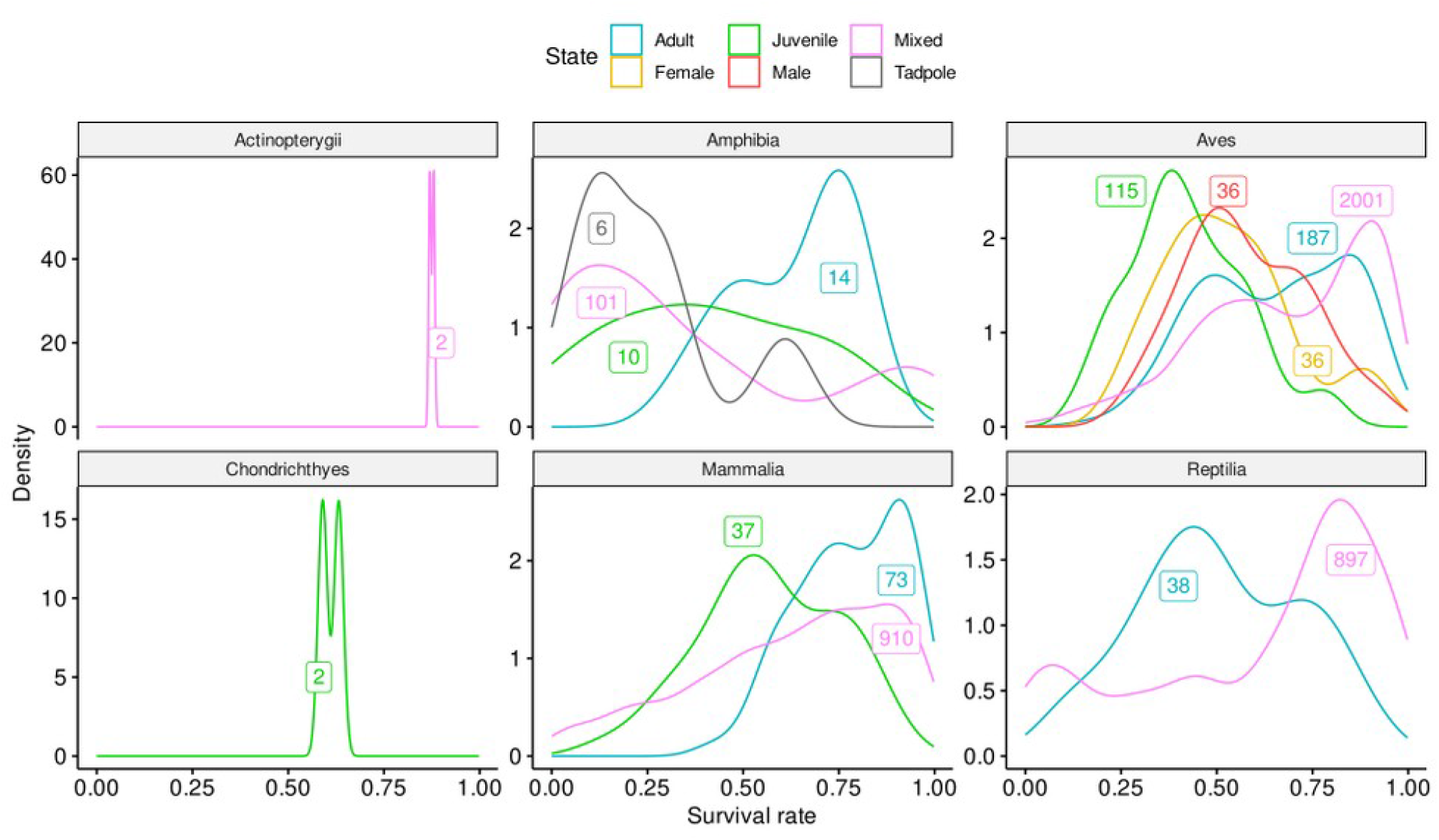
Density of probability of survival across age class and taxa for 700 species. Sample size are indicated by labels, with colour corresponding to the state of individual sampled.

### 2.1 Data generation

For each scenario, we conducted a simulation using data from a ten-year study period with multiple capture occasions. During these capture occasions, individuals could be in one of five possible states: “A”, “B”, “C”, “D”, or death itself. State “D” was considered an absorbing state, meaning that once an individual reached this state, it could not change to any other state. This can be used to model permanent emigration, for example. In the initial occasion, the individuals in states “A”, “B”, and “C” consisted of 40 juveniles (with a 1:1 sex ratio) and 60 adults (80% females, 20% males). No individuals were initially in state “D”. In each subsequent occasion, we marked 40 juveniles and 5 adults in each state (“A”, “B”, “C”), except for state “D”, where recapture was not possible and only observation was feasible. This conceptual system can be seen as having three breeding sites (“A”, “B”, “C”) where capture and resighting occurred annually, and a surrounding area (“D”) where only resighting was possible. For males only, we allowed a permanent transition to the “D” state. It is worth noting that sexual asymmetry in state change is common in various taxa, where one sex is more likely to disperse through permanent emigration (absorbing state). For each simulated scenario, all juveniles captured at a specific occasion were considered adults in the following occasion. All scenarios followed the same steps for data generation, as shown in Figure 2. We initiated the simulation by generating the survival of individual i at occasion t using a Bernoulli distribution:

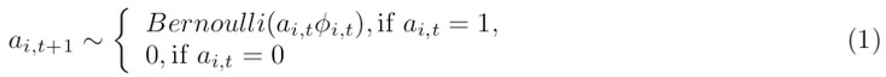

where *a_i,t_* = 1 if individual *i* is alive at time *t*, and 0 if not. *Φ_i,t_* represents the specific survival probability for each individual at a given state, time, and age. To introduce stochastic variations in annual survival for the two age classes, we drew survival probabilities from a logistic distribution. If an individual survived, we simulated the state transition using a Categorical distribution:

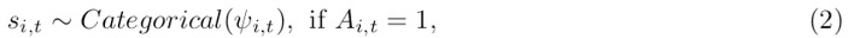

where *ψ*_i,t_ represents a state-, sex-, and age-specific transition probability. The study examined the transitions of females between states “A”, “B”, and “C” at a constant rate, regardless of their age. However, females were not allowed to transition to state “D”. Juvenile males, on the other hand, could only transition to state “D”, with the proportion depending on their initial state. Adult males, however, did not change their state throughout the study. The mark loss/retention process was simulated using a Bernoulli distribution:

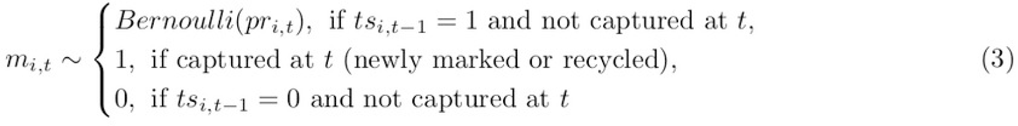

with *m_i,t_* indicating the presence of a mark on individual *i* at occasion *t* and *pr_i,t_* representing the probability of retention (see Supporting Information 2, Fig. S2), which is the complement of the mark loss probability *pl* (*pl_i,t_* = 1-*pr_i,t_*). The detection process was simulated using a Categorical distribution:

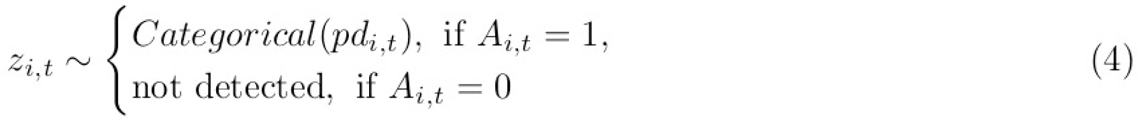

with *pd*_i,t_ representing the state-specific detection probability (Supporting Information 2, section 1.2). We examined two common methods for tracking individuals in our study: physical capture and resighting. Physical capture involves either recycling the individuals by re-marking them or checking their existing marks. Resighting, on the other hand, is a passive detection method that only includes individuals who have retained their marks. This approach is based on the observation that, in many studies, the probability of resighting is usually higher than the probability of recapture, making them susceptible to different estimation biases. We categorized the probability of detection into seven categories, with each category conditioned on the retention of the mark.

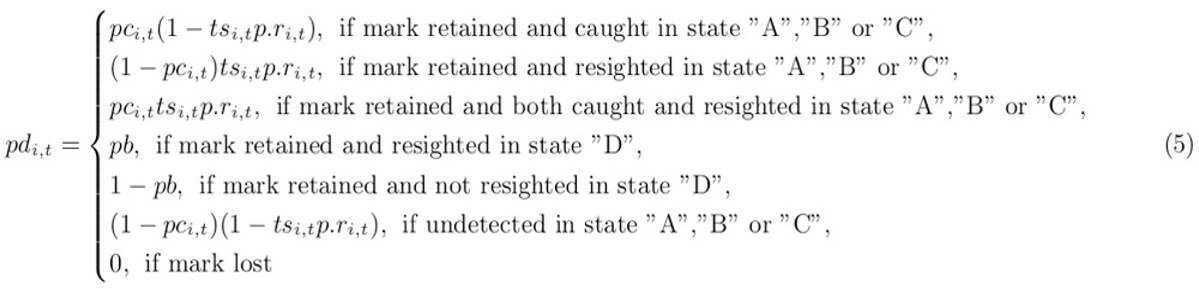

where *pc*_i,t_ was a state specific capture probability, *p.r*_i,t_ a state specific resighting probability and *pb* the detection probability in state “D”.

**Figure 2:**
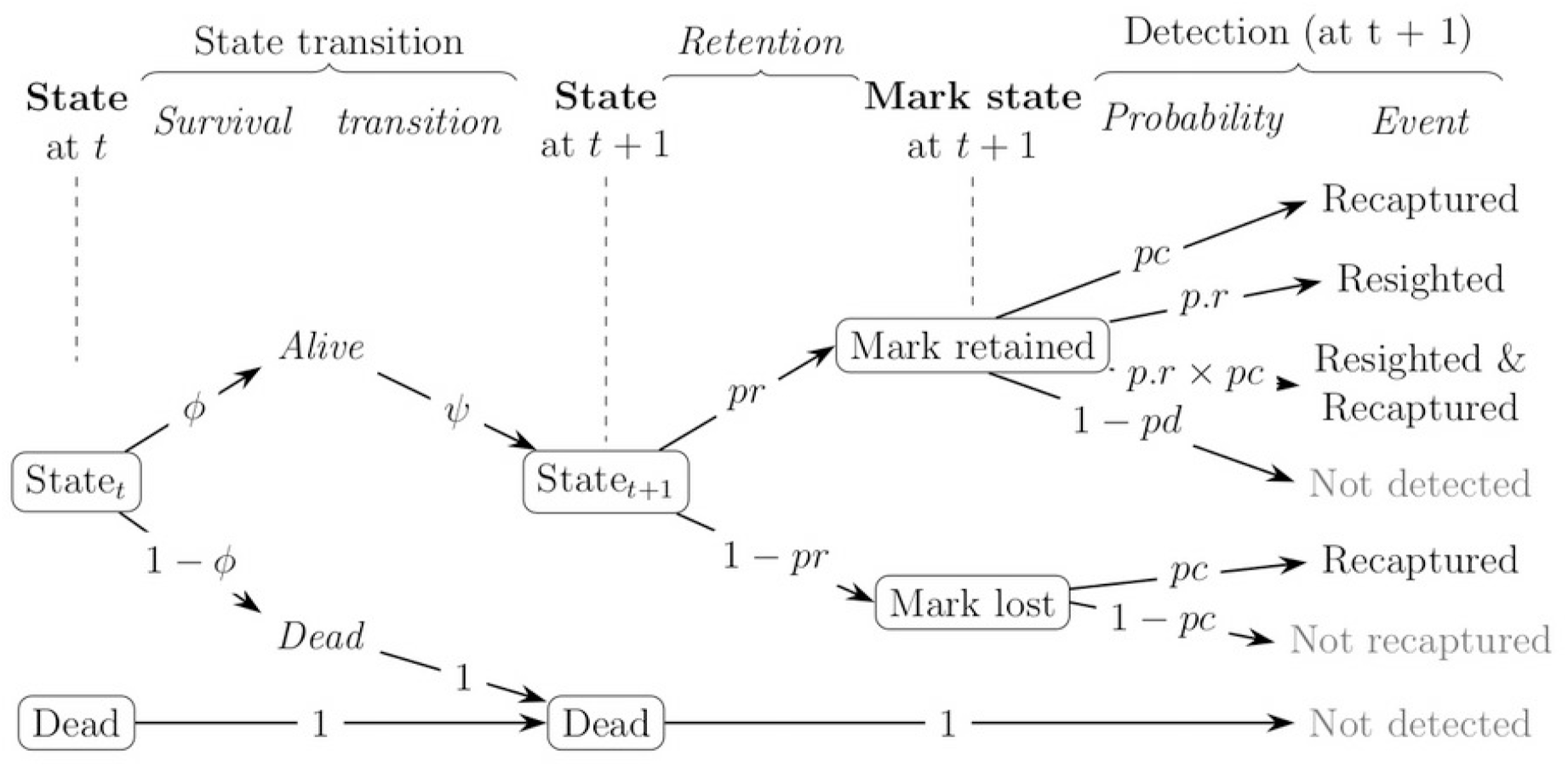
The possible fates of an individual between occasion t and t+1. We first consider the state transition process: if the individual dies between occasions, it can only remain dead and can no longer be detected, both with probability 1 (absorbing state). The individual can, however, survive between occasions with probability φ (depending on time and state at t) and can then change state with probability ψ (see Fig. 1). Second, the mark retention process: if it survives between occasions, then the individual can retain its mark with probability pr (depending on age and time since marking) or lose its mark with probability pl = 1-pr. Thirdly, the detection process: if this individual has lost his mark between occasions, he may possibly be recaptured with a probability pc (depending on time and state at t+1) and if this happens, he is marked again before being released. However, if the mark is retained, other events may occur: the individual may also be captured with probability pc, but it may also be resighted with probability p.r (depending on the state at t+1), or both with probability pc*p.r, or finally not be detected at all with probability 1-pd = (1-pc)*(1-p.r).

Since we can expect higher rates of recycling when recapture rates are high, and little recycling otherwise, we considered four scenarios (Table 1): (1) long-lived species with a high detection rate; (2) long-lived species with a low detection rate; (3) short-lived species with a low detection rate; (4) short-lived species with a high detection rate. The values of the simulated parameters can be found in Table 2 and Figure 3. Mark loss often depends on the time since marking, and in many species, it occurs soon after marking (Fabrizio *et al*., 1999; Fokidis *et al*., 2006; Kremers, 1988; Nichols & Hines, 1993). In our scenarios, we included this process and determined a range of mark loss rates from the literature (see Supporting Information 1, Table S2). We simulated three mark loss probabilities: low (*pl* = 0.05), medium (*pl* = 0.25), and high (*pl* = 0.4) during the first year after marking, and a constant rate of 0.05 thereafter. This generated a variety of cases of mark loss and recycled individuals (see Supporting Information 2, Fig. S2-3). This process enabled us to generate datasets that took mark loss into account in the presence of a second permanent mark. In order to generate datasets where mark loss was not considered (no second permanent mark), we created recycled individuals with life histories corresponding to the portion of their lives after mark loss, and replaced the original life history from mark loss with zeros. For example, a life history “1111” of an individual that lost its mark between occasions 2 and 3, and was re-marked at occasion 3, would become two new histories: (1) “1100”, representing the first part of the life before mark loss, and (2) “0011”, representing a second individual that was not recognized in the absence of a permanent mark, but considered newly recruited. Using this data generation process, we simulated 2 x 50 datasets for each of the 12 combinations of parameters (50 with and 50 without recycling), resulting in a total of 1200 simulated datasets (see Supporting Information 2, Fig. S1). The computational codes for a fully reproducible example dataset are provided in the Supporting Information 2.

**Figure 3:**
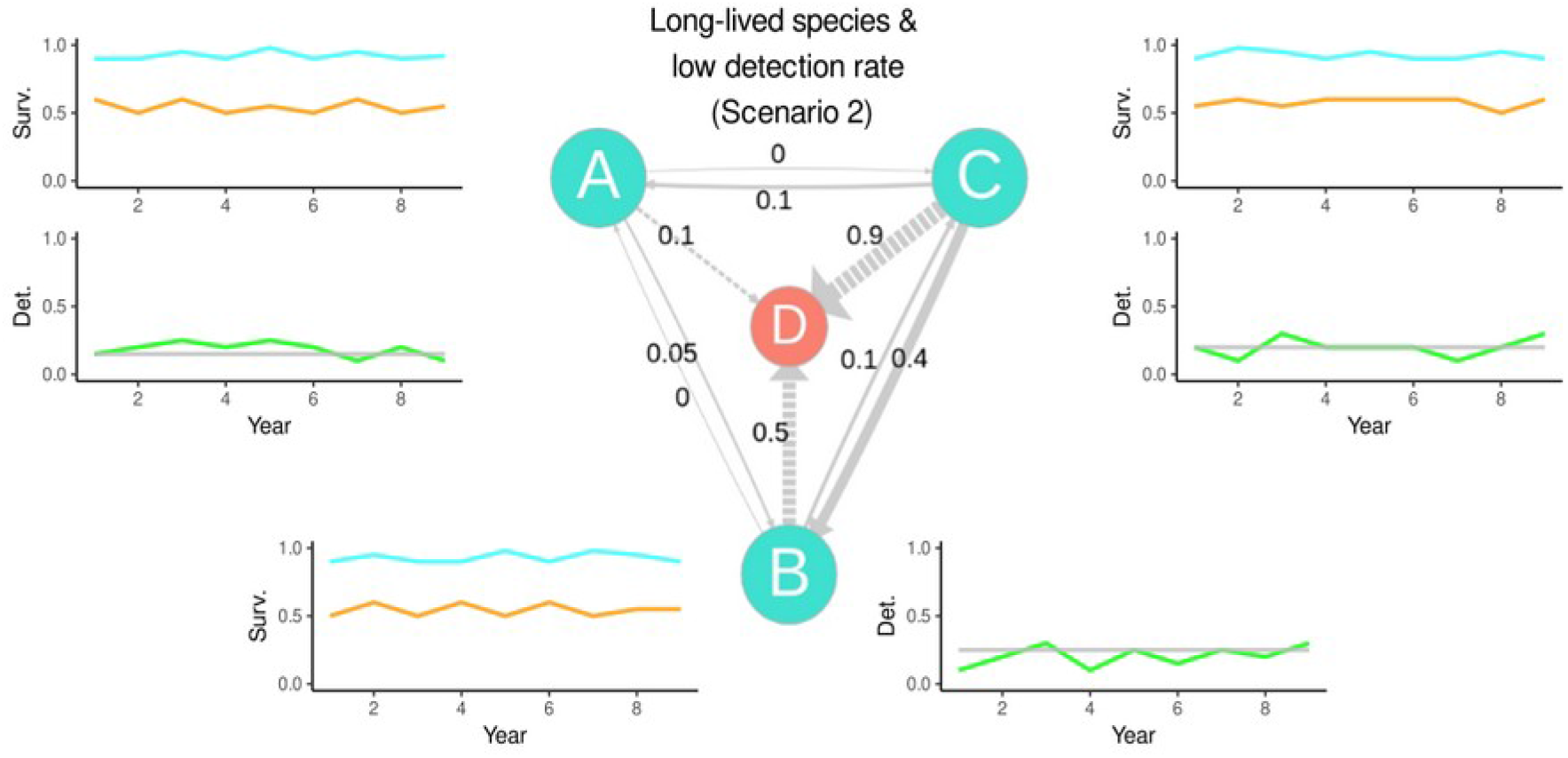
Schematic description of parameter values used to simulate data under scenario 2 (long-lived species with low detection). Central graph: solid arrows correspond to possible transitions of females and dashed arrows to those of males. The size of the arrows is proportional to the probability of transition indicated next to them, all were kept constant over time. Peripheral graph: simulated survival (Surv.) and detection (Det.) probabilities were displayed for states “A”, “B” and “C”. The light blue lines correspond to adult survival, the orange lines to juvenile survival and the green lines to the probability of capture, which are derived from Normal and Uniform distribution and therefore fluctuate over the years (see Table 1). The grey lines correspond to the probability of resighting, they differed between state but were set constant in time.

**Table 1:**
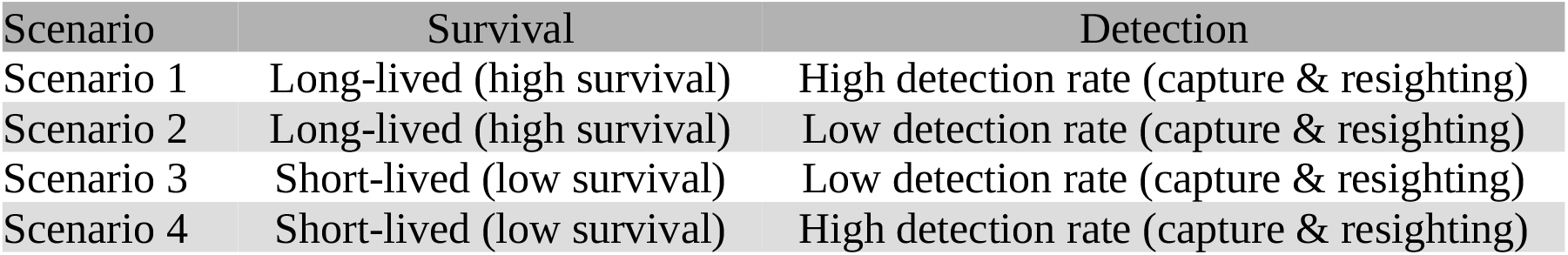
Summary of the characteristics of each simulated scenario.

| Scenario | Survival | Detection |
| --- | --- | --- |
| Scenario 1 | Long-lived (high survival) | High detection rate (capture & resighting) |
| Scenario 2 | Long-lived (high survival) | Low detection rate (capture & resighting) |
| Scenario 3 | Short-lived (low survival) | Low detection rate (capture & resighting) |
| Scenario 4 | Short-lived (low survival) | High detection rate (capture & resighting) |

**Table 2:** Parameter values used to simulate the 4 scenarios. For random values generated, the corresponding distribution is indicated with N (a, b) the normal distribution with mean a and variance b, and U(a,b) the uniform distribution with lower bound a and upper bound b. The square brackets show mean values on the probability scale. To simulate survival for short-lived species, we used the same distribution on as long-lived species but subtract generated values by 0.3 for adults and 0.2 for juveniles. In the same way, we obtained the low values of probability of capture and re-sighting by subtracting 0.5 from the high values. The probability of capture in state “D” is set to 0, as no capture is possible when individuals are in this state. For the transition values between states see Fig. 3.

| Parameter | Definition | Value |  |
| --- | --- | --- | --- |
|  |  | Long-lived species | Short-lived species |
| $\phi_{ad.}$ | Adult survival in state "A", "B", "C" | logit(N(2.5,0.3)) [0.92] | logit(N(2.5,0.3)-0.3) [0.62] |
| $\phi_{ad.}$ | Adult survival in state "D" | logit(N(1.5,0.3)) [0.81] | logit(N(1.5,0.3)-0.3) [0.51] |
| $\phi_{juv.}$ | Juvenile survival in state "A", "B", "C" | logit(N(0.2,0.3)) [0.55] | logit(N(0.2,0.3)-0.2) [0.35] |
| $\phi_{juv.}$ | Juvenile survival in state "D" | NA | NA |
|  |  | High | Low |
| $pc_A$ | Capture probability in state "A" | U(0.6,0.7) [0.65] | U(0.6,0.7)-0.5 [0.15] |
| $pc_B$ | Capture probability in state "B" | U(0.7,0.8) [0.75] | U(0.7,0.8)-0.5 [0.25] |
| $pc_C$ | Capture probability in state "C" | U(0.65,0.75) [0.7] | U(0.65,0.75)-0.5 [0.2] |
| $pc_D$ | Capture probability in state "D" | 0 | 0 |
| $p.r_A$ | Resighting probability in state "A" | 0.85 | 0.35 |
| $p.r_B$ | Resighting probability in state "B" | 0.95 | 0.45 |
| $p.r_C$ | Resighting probability in state "C" | 0.9 | 0.4 |
| $p.r_D$ | Resighting probability in state "D" | 0.7 | 0.2 |

### 2.2 Statistical models

As described, we simulated two different datasets for each combination, in which the assumption of permanent marking is violated. In the first dataset, individuals can still be identified even after the loss of their mark, thanks to a second permanent mark. In the second dataset, recycling occurred because there was no second permanent mark present (Supporting Information 2, Fig. S3). We developed two AS models to analyse these data. ModelA, used for the first dataset, included the estimation of mark loss. ModelW, used for the second dataset, ignored mark loss (Supporting Information 2, sections 1.7 and 2.2).

To estimate the state, time, and age-dependent survival *Φ* for both models, we used a Bernoulli distribution (see Eq. 1 above).

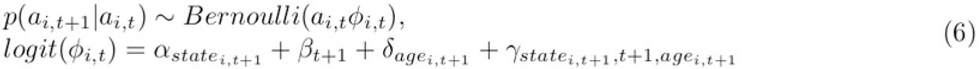

where *a_i,t_* is the life status of individual *i* at time *t* and coefficient α is a state effect, *β* a time effect, *δ* an age effect and *γ* a simultaneous effect of state, age and time. The transition and detection processes were estimated using the same distributions as described for data generation (see Eq. 2, 4, 5 above). Only in Model A, the fate of marks (whether they were retained or lost) for each individual was tracked. Various combinations of single, double, and permanent marks have been used to estimate mark loss (Laake *et al*., 2014). In Model A, we used a single mark loss approach, with the second mark being permanent. This allowed us to determine whether the non-permanent mark was lost or retained at each capture occasion, and resighting was conditional on mark retention (see Eq. 5). The mark retention process was directly incorporated into the model using a state space approach. While it is possible to model the presence-absence of marks as states with a transition matrix (McMahon & White, 2009), or as a hidden Markov process for unobserved individuals where mark retention status is unknown (Laake *et al*., 2014), we chose to model the mark retention process using a Bernoulli distribution with p(*m*_i,t_|*m*_i,t-1_) ∼ Bernoulli(*pr_i,t_*), if the individual was marked or retained its mark at *t-1* (*m_i,t-1_* = 1). We estimated three retention probabilities (*pr*) as a function of age and time since marking:

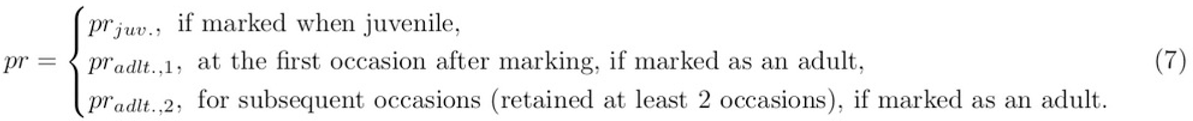

When an individual was lost and subsequently recaptured, a new mark was applied in modelA without altering its identity. In contrast, modelW does not incorporate mark retention, so if an individual lost its mark and was recaptured, it was regarded as newly recruited. The priors utilized for each parameter can be found in Supporting Information 2, section 1.6.

### 2.3 Application on a bats dataset

The simulations we conducted served two purposes: first, to test general hypotheses regarding the impact of mark loss on parameter estimates in multi-state models, and second, to validate a parametrization approach that can accurately estimate these parameters without bias. Based on this, we are able to accurately estimate the probability of mark loss in our own dataset and determine the importance of using a permanent mark in the long term (Juillet *et al*., 2011). Our empirical dataset consists of a 10-year study on the greater-mouse eared bat (*Myotis myotis*) in Brittany, France (2010-2019). We marked a total of 2,561 individuals in 5 maternity roosts: La Roche Bernard (47°31’N, 2°18’W), Férel (47°28’N, 2°20’W), Noyal-Muzillac (47°35’N, 2°27’W), Béganne (47°35’N, 2°14’W), and Limerzel (47°38’N, 2°21’W). The bats were individually tagged using Passive Integrated Transponders (PIT) tags, specifically the ID-100C (Troven®), which had a unique 10-digit code. These small tags (2.12×11mm, 0.1gr) allowed for identification using passive readers. All individuals captured in roosts without PIT-tags were consistently tagged, including both those who lost their tags and those who were never tagged before. Additionally, a second permanent marking method involved genotyping all newly tagged individuals. Genotypes were determined from DNA extracted from wing biopsies of all tagged individuals and all untagged males captured during swarming surveys (n=4,258 samples; see Supporting Information 3, Fig. S1). A total of 16 microsatellite markers optimized for *Myotis myotis* were used to establish individual genotypes (Foley *et al*., 2020). To minimize genotyping errors, we analysed two different samples per individual, whenever available. All samples were genotyped and scored twice by two different individuals (Frantz *et al*., 2003; Puechmaille & Petit, 2007). We also conducted genetic profile comparisons to identify errors. The error rate of genotypes was hypothesized to be sufficiently low to be negligible, and this source of uncertainty was not included in the models (but see Winiarski & McGarigal, 2016). In addition, we checked for lost tags on the floor of the maternity roosts during each winter. This allowed us to partially identify individuals that lost their tags (61.5% compared to genotype), with the remaining losses occurring outside of the roosts. Most tag losses occurred during the first year, as supported by the absence of records from passive reading detectors (Supporting Information 3, Fig. S2). Out of a total of 2,561 individuals, 252 (∼10%) were identified as having lost their tag at least once. Of these individuals, 94 were recaptured and retagged once, while three were retagged twice. For 13 individuals, retagging occurred during the last capture occasion, resulting in the recycling of 81 individuals out of the 94. To analyse these data, we employed a multisite model that accounted for movement between maternity roosts as transitions (similar to the AS model used for simulated data, Supporting Information 3). We divided the population into two age classes for survival analysis: juveniles (in their first year of life) and adults (one year and older). To examine departures from the AS model assumptions, we conducted goodness-of-fit tests using R2ucare (Gimenez *et al*., 2018b), an R package based on U-CARE (Choquet *et al*., 2009a). Minor overdispersion was observed in tests for transience (ĉ_3G.SR_=1.82) and memory (ĉ_WBWA_ = 1.96). Insufficient data in the individual contingency tables prevented the performance of other goodness-of-fit tests. Hence, we retained our two-age class structure for survival modelling. To account for unmeasured individual survival heterogeneity, we included a normally distributed random effect in the estimation of survival probability. This was necessary as other covariates were unable to capture the relevant variation. Individual heterogeneity plays a crucial role in population dynamics and evolution and is prevalent in wild populations (Gimenez *et al*., 2018a).

Emigration from the five subpopulations under study was assessed using capture and resighting data collected at swarming and wintering sites between capture occasions. We defined eight different detection states, which enabled us to separately estimate capture and resighting probabilities (see Supporting Information 3, Table S1). Based on empirical data suggesting potential movement of individuals between all subpopulations and outside areas, we did not restrict transitions between subpopulations, except for the movement of juveniles from outside the maternity roosts, which was not possible due to the absence of tagging outside the five roosts. Therefore, the probability for this type of movement was set to 0 (see Supporting Information 3, Fig. S3). This study also examined the impact of using surgical adhesive (Vetbond®) after PIT-tag injection to reduce tag loss compared to self-healing techniques (Lebl & Ruf, 2010; van Harten *et al*., 2020). In this model, tag retention probabilities were modelled similar to Eq. 7, taking into account time since marking (categorized as first year or subsequent years), individual age class (juvenile or adult), and the use of surgical adhesive (yes or no). Similar to the simulated datasets, two models were run on two datasets: one for estimating tag loss using genotyping (as a second permanent mark), and another model that ignored this information on a transformed dataset that included recycled individuals (following the same process as the simulated data).

### 2.4 Estimation procedures and assessments

Despite the possibility of using a frequentist approach for analysing simulated data (Lebreton *et al*., 2009), we opted for a Bayesian approach to ensure consistency with the analysis of empirical data, where these methods offer greater flexibility in accounting for individual heterogeneity in survival (Gimenez *et al*., 2018a). The simulated and empirical data were analysed using JAGS (Kruschke, 2014; Plummer, 2003) through the jagsUI package (Kellner, 2016) in R 3.6.0 (R Core Team, 2019). We employed four Monte Carlo Markov chains (MCMC) with 150,000 iterations each and discarded the initial 50,000 iterations (burn-in) when sampling from posterior distributions. We retained every twentieth iteration, resulting in 20,000 samples from the posterior distribution for each parameter. Chain convergence was assessed using the Gelman-Rubin statistic, denoted R-hat (Brooks & Gelman, 1998). Among the 1,200 simulations, some parameters exhibited R-hat values > 1.05, indicating convergence failure. Less than 0.4% of the estimated parameters for the model accounting for mark loss did not converge (Supporting Information 2, Table S1), particularly the coefficient *γ* (combined effect of state, time, and age on survival probability) and *γ.c* (combined effect of state and time on detection probability). The mean R-hat values for these parameters were less than 1.2 (Supporting Information 2, Fig. S4). For models that did not account for mark loss, 1.3% of the estimated parameters did not converge (Supporting Information 2, Table S2), especially in scenarios with low detection or low detection and survival rates (scenario 2 and 3, respectively), and mark loss probabilities set to 0.25 or 0.4. In most cases, the parameters that failed to converge exhibited a mean R-hat value less than 1.2, with only a few exceeding 1.5 (Supporting Information 2, Fig. S5). Convergence failures did not pertain to the probability of mark loss in any of the simulations. To avoid excessively long computing times, we did not increase the number of iterations to achieve complete convergence of the MCMC chains for these parameters in the affected simulations. Our results are based on 50 simulated datasets per scenario, and we assume that the lack of convergence for these few parameters has no substantial influence on our results. To assess bias in parameter estimates when mark loss or recycling is not accounted for, we calculated the Earth Mover Distance (EMD) using the EMD-L1 algorithm (Ling & Okada, 2007), which quantifies the difference between two distributions. This metric measures the minimum cost of transforming one distribution into another point by point. For each scenario, we also estimated a Region Of Practical Equivalence (ROPE, Kruschke, 2018) to assess the degree of difference between the distributions represented by the EMD metric. To define the ROPE for each scenario, we randomly constructed 1,000 pairs of models from the 50 simulations and calculated the associated 1,000 EMDs from the posterior distributions of the estimated parameters (for more details, see Supporting Information 2, Fig. S76). The resulting distributions of EMDs represented the expected variations in inferences obtained from simulations initiated with the same parameter values. The ROPE was then defined as the range between 0 and the upper value of the 80% highest posterior density interval (hdi) from the distribution of these EMDs. Finally, the proportion of EMDs for each simulated case that fell outside the ROPE was computed, providing a direct indication of bias; the higher this proportion, the greater the bias. Comparisons of EMDs between the models that accounted for tag loss and recycling, and those that did not, in relation to their respective ROPEs, depict cases in which not accounting for tag loss leads to estimates that substantially differ from estimates obtained when considering tag loss. We also evaluated parameter bias as the difference between the median of the posterior distribution and the simulated true value (median – truth). In the analysis of empirical data, we used the median of the posterior distribution of parameters from the model accounting for a secondary permanent mark as the true value. Finally, we calculated the precision of the parameter estimates in the simulated data using mean squared errors (MSE = bias^2^ + variance).

## 3 Results

### 3.1 Simulation results

The estimated numbers of lost marks and recycled individuals increased in scenarios with higher survival, detection, and mark loss rates (Supporting Information 2, Fig. S3). As expected, when the recapture rate was high (i.e. in scenarios 1 and 4), the proportion of recycled individuals relative to the number of lost marks was higher for an equivalent rate of mark loss. Regardless of the scenarios, there was no estimation bias on demographic parameters when the mark loss rate was set to 0.05 (Supporting Information 2, Fig. S28-S31). However, the number of parameters with biased estimates increased with the rate of mark loss, but the magnitude of bias varied across the simulated scenarios. Specifically, in scenario 1 (Fig. 4 & 5), there were substantial underestimations in some adult and juvenile survival rates. The probability of remaining in the same state was also underestimated among juveniles, leading to an overestimation of their probability of transitioning to another state (Fig. 4 & 6). The resighting probability was underestimated in all states except B (Fig. 4 & 6). The biases in state transitions were particularly high for transitions to absorbing states, such as juveniles transitioning to state “D” (Supporting Information 2, Fig. S57-S71). In simulations with the highest mark loss rate (0.4), scenarios 1 and 4 (which had high recapture probabilities) showed underestimates of adult survival only in the early years of the study (Supporting Information 2, Fig. S21-S22 and S27). The bias in juvenile survival was less pronounced, although moderate underestimations occurred for high mark loss rates, especially for state A and B in scenarios 1 and 4 (Supporting Information 2, Fig. S21-S22 and S27). Lack of precision in the estimate of juvenile survival was also observed when the model did not account for tag loss and the mark loss rate was high. The resighting probability showed substantial bias, with underestimates mainly in states A and C (Fig. 4 and 7.a), as well as a lack of precision for all scenarios and mark loss rates (Supporting Information 2, Fig. S28-S49). However, recapture, the second component of detection, showed little bias except during the second capture occasion for states A and C in scenario 1 when the mark loss rate was 0.25, and scenarios 1 and 4 when the mark loss rate was 0.4. There was also a decrease in the precision of the recapture parameter at high mark loss rates. When mark loss and recycling were ignored, a large percentage of the transition probability estimates were biased (Fig. 6b-e), with an overall underestimation of the probability to remain in the same state and, as a result, an overestimation of the probability of changing state. This decrease in precision mainly occurred at high mark loss rates (Supporting Information 2, Fig. S50-S71). Overestimations of transition probabilities were observed in juvenile males, except for transitions from state C to D, where the transition rate was set to 0.9. For females, when mark loss was set at a high level, the same pattern was observed in states A and B, from which state transitions were set at a low level. On the contrary, when the transition from state C to B was set at a high level (0.4), there was an underestimation of the transition probability and an overestimation of the probability to remain in state C, particularly in scenarios with low detection rates (scenario 2 and 3, Supporting Information 2, Fig. S67-S70).

**Figure 4:**
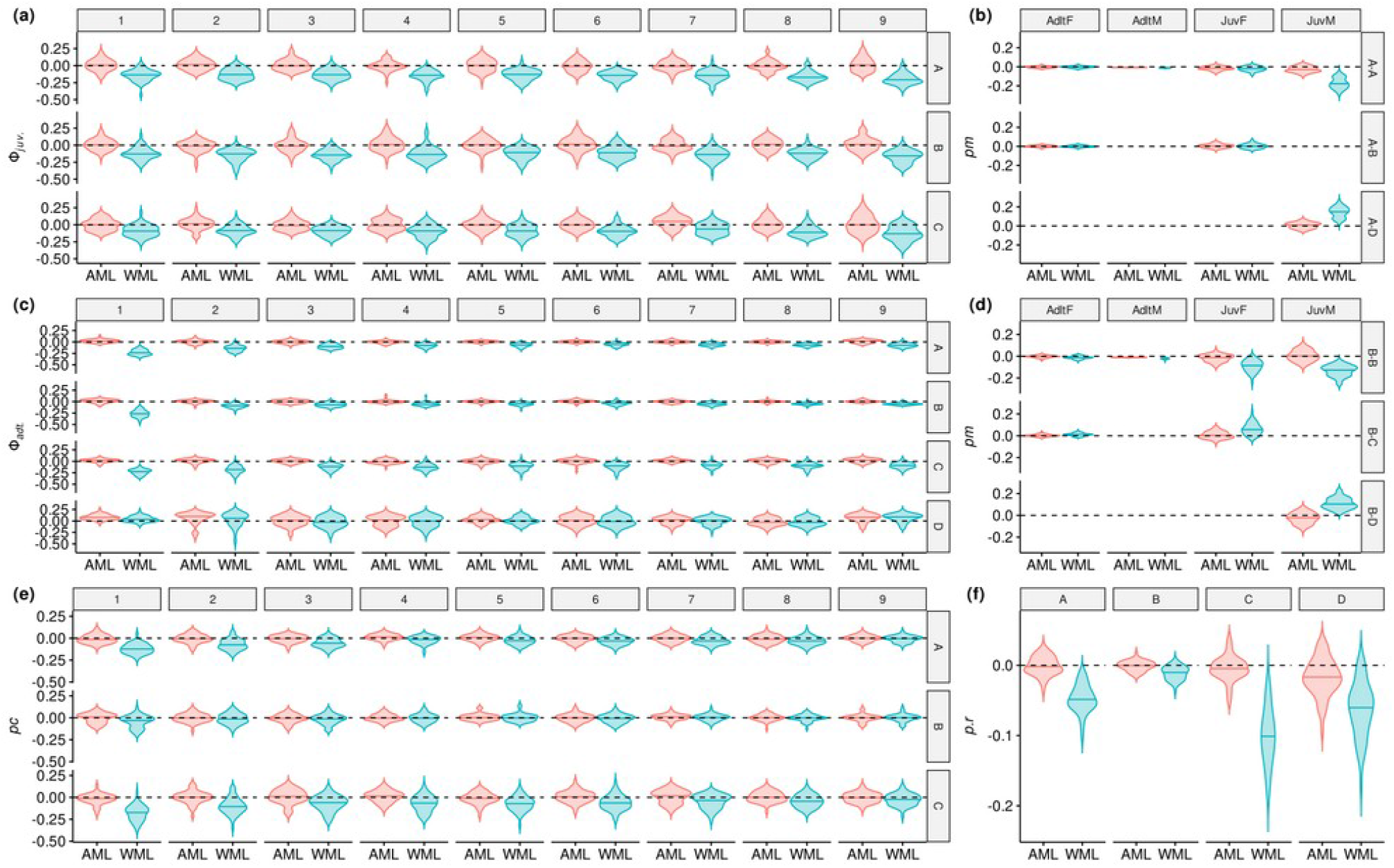
Comparison of bias for estimates of juvenile (a) and adult (c) survival, transition (b and d), capture (e) and resighting (f) probabilities between model accounting for mark loss (AML) or not (WML). All violin plots show the distribution of bias over 50 simulations from scenario 1 (long-lived species and high detection probabilities), with a simulated probability of mark loss of 0.4. The median of each simulated distribution is shown with a horizontal line. The numbers 1 to 9 are the recapture opportunities, the letters from A to D represent the different states, AdtF the adult females, AdtM the adult males, JuvF the juvenile females and JuvM the juvenile males.

**Figure 5:**
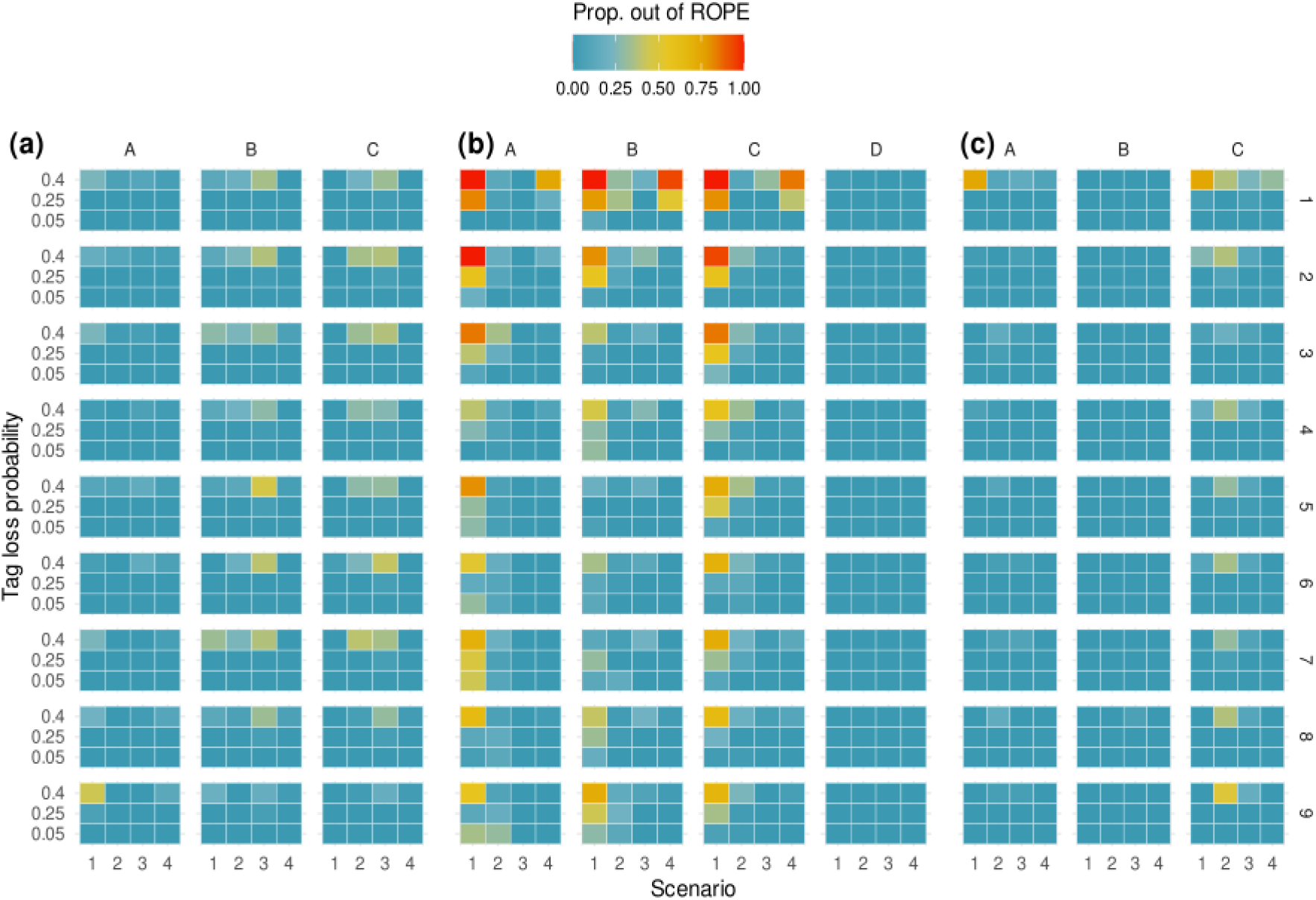
Tile-plots of the proportion of the distribution of the Earth Mover Distance (across 50 simulated datasets) out of the Region of Practical Equivalence (ROPE), between the model accounting for tag loss and recycling (ModelA) and the model ignoring them (ModelW). The ROPE corresponds to the interval including 80% hdi of the posterior density distribution of the “true value“ of a parameter which was estimated with ModelA. Each tile represents annual (right axis) juvenile survival (a), adult survival (b) and capture probability (c) for each scenario (y axis) and tag loss probabilities (x axis). The scenarios indicated at the bottom are: (1) long-lived species and high detection rate; (2) long-lived species and low detection rate; (3) short-lived species and low detection rate; (4) short-lived species and high detection rate. At the top of each panel, A, B, C and D correspond to the states.

**Figure 6:**
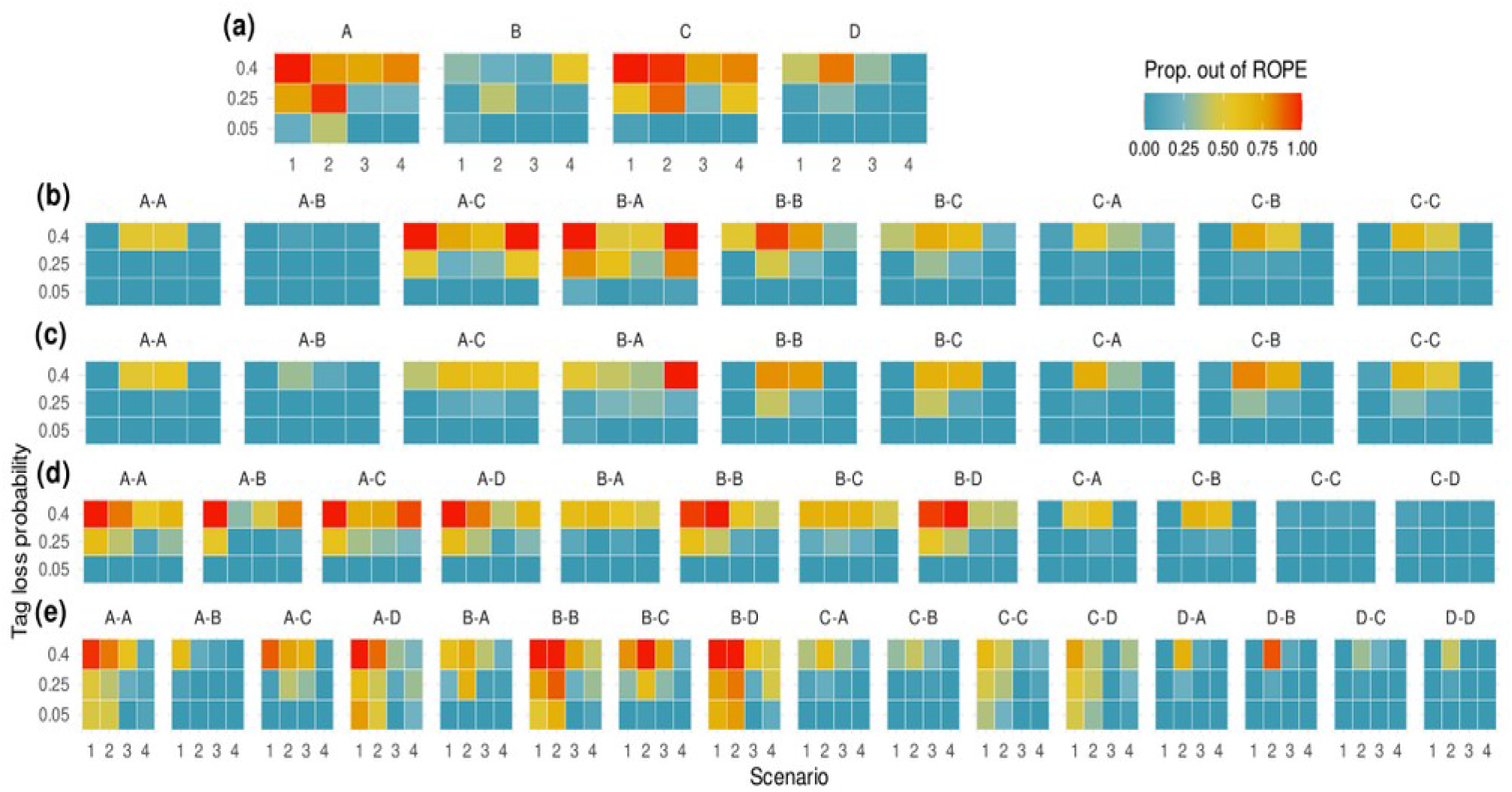
Tile-plots of the proportion of the distribution of the Earth Mover Distance (across 50 simulated datasets) out of the Region of Practical Equivalence (ROPE), between model accounting for tag loss and recycling (ModelA) and the model ignoring them (ModelW) for each simulated cases. The ROPE corresponds to the interval including 80% hdi of the posterior density distribution of the “true value“ of a parameter which was estimated with ModelA. Each tile represents resighting probability (a) and transition probabilities between subpopulations (direction, “from-to”, are indicated above each tile-plot, e.g. “A-B” correspond to state transition from A to B) of juvenile female (b), adult female (c), juvenile male (d) and adult male (e) for each scenario and tag loss probabilities. The scenario are indicated at the bottom: (1) long-lived species and high detection rate; (2) long-lived species and low detection rate; (3) short-lived species and low detection rate; (4) short-lived species and high detection rate. At the top of each panel, A, B, C and D correspond to the states.

**Figure 7:**
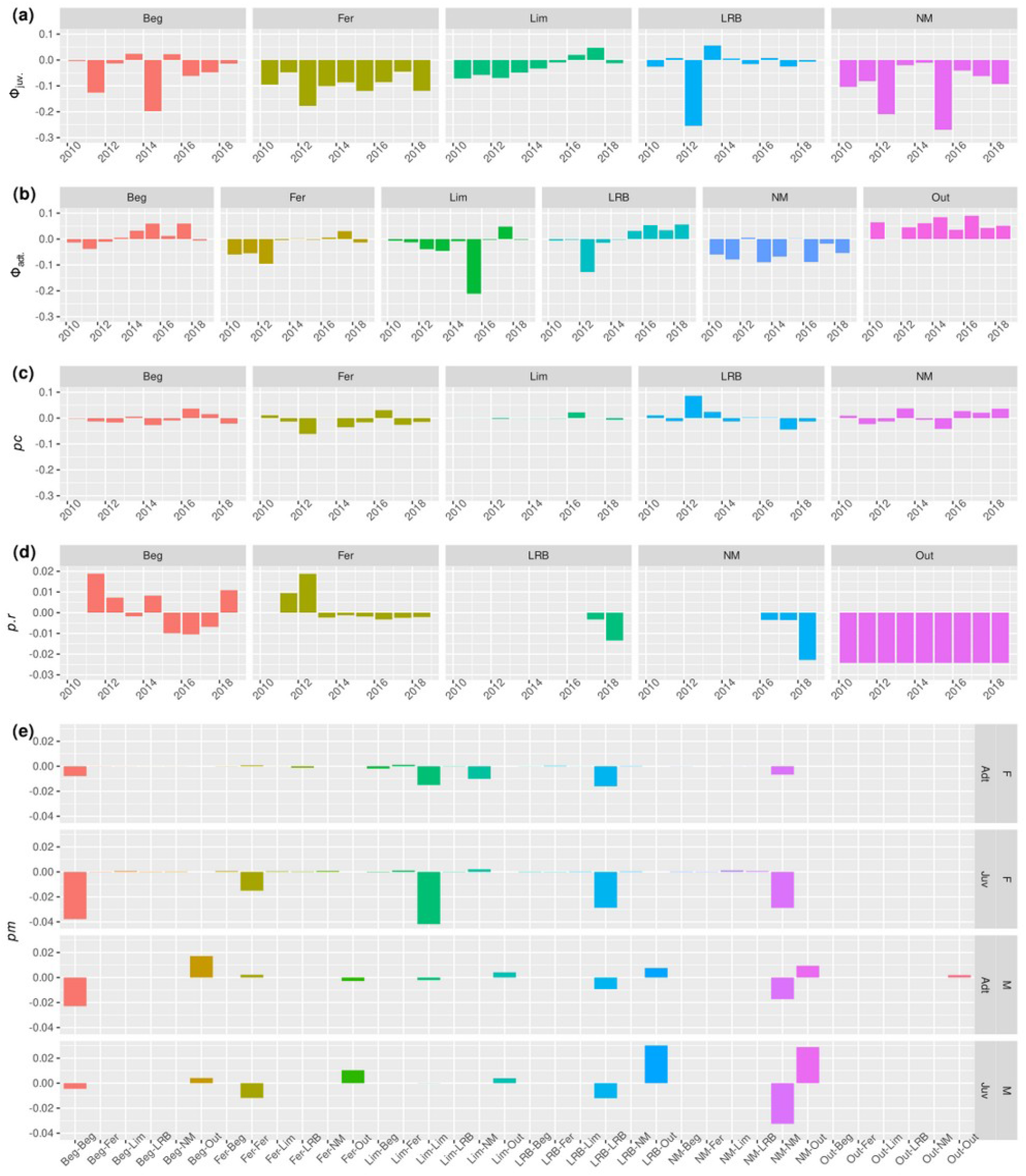
Differences in the medians of the posterior distributions of juvenile (a) and adult (b) survival, capture (c) resighting (d) and transition (e) probabilities between the model accounting for tag-loss and the model ignoring it, both estimated from the empirical data. Colonies are abbreviated: Beg = Beganne; Fer = Férel; LIM = Limerzel; LRB = La Roche Bernard; NM = Noyal-Muzillac. Movements between sites are indicated on x axis with direction “from-to”. Movements (e) are specified by age (Adt. = Adulte, Juv. = juvenile) and sex classes (M = male, F = Female), indicated on the right side of the plot (e).

### 3.2 Bat metapopulation

Most of the estimated parameters, such as survival, capture, resighting, and state transition probabilities, exhibited both negative and positive biases without a clear pattern (Fig. 7, Supporting Information 3, Fig. S4-S11). The largest biases were observed in survival estimates, with a median underestimation of survival of over 0.26 in juveniles (Fig. 7a) and 0.21 in adults (Fig. 7b). Emigration, defined as movement outside the studied maternity sites (“Out”), was overestimated by an average of 0.05 throughout the study (Fig. 7b). The probability of recapture varied from overestimation to underestimation by up to 0.1, depending on the occasion and roost (Fig. 7c). The estimated bias in the other parameters was minimal (Fig. 7d and 7e). Tag loss probability was significantly higher in juveniles, but the use of surgical glue substantially reduced this rate (Fig. 8). Specifically, the tag loss probability decreased by one third from 0.28 (90%hdi [0.23,0.33]) to 0.19 (90%hdi [0.16,0.22]) with the use of surgical glue. In adults, the use of surgical glue did not affect tag loss rate, with a 69% overlap in the probability distributions. The tag loss rate in adults was approximately 0.1, which is half the rate observed in juveniles when surgical glue was used. When considering the period following one year post-tagging, the probability of tag loss was higher when surgical glue was used (median 0.03, 90%hdi [0.02, 0.04]) compared to when it was not used (median 0.02, 90%hdi [0.01, 0.02]). However, this difference may be an artefact due to a lack of search for lost tags on the ground of the maternity roost in the first years of the study (Supplementary Information 3, part 3 and Fig. S12). Additional parameter estimates can be found in Supporting Information 3, part 2.6.

**Figure 8:**
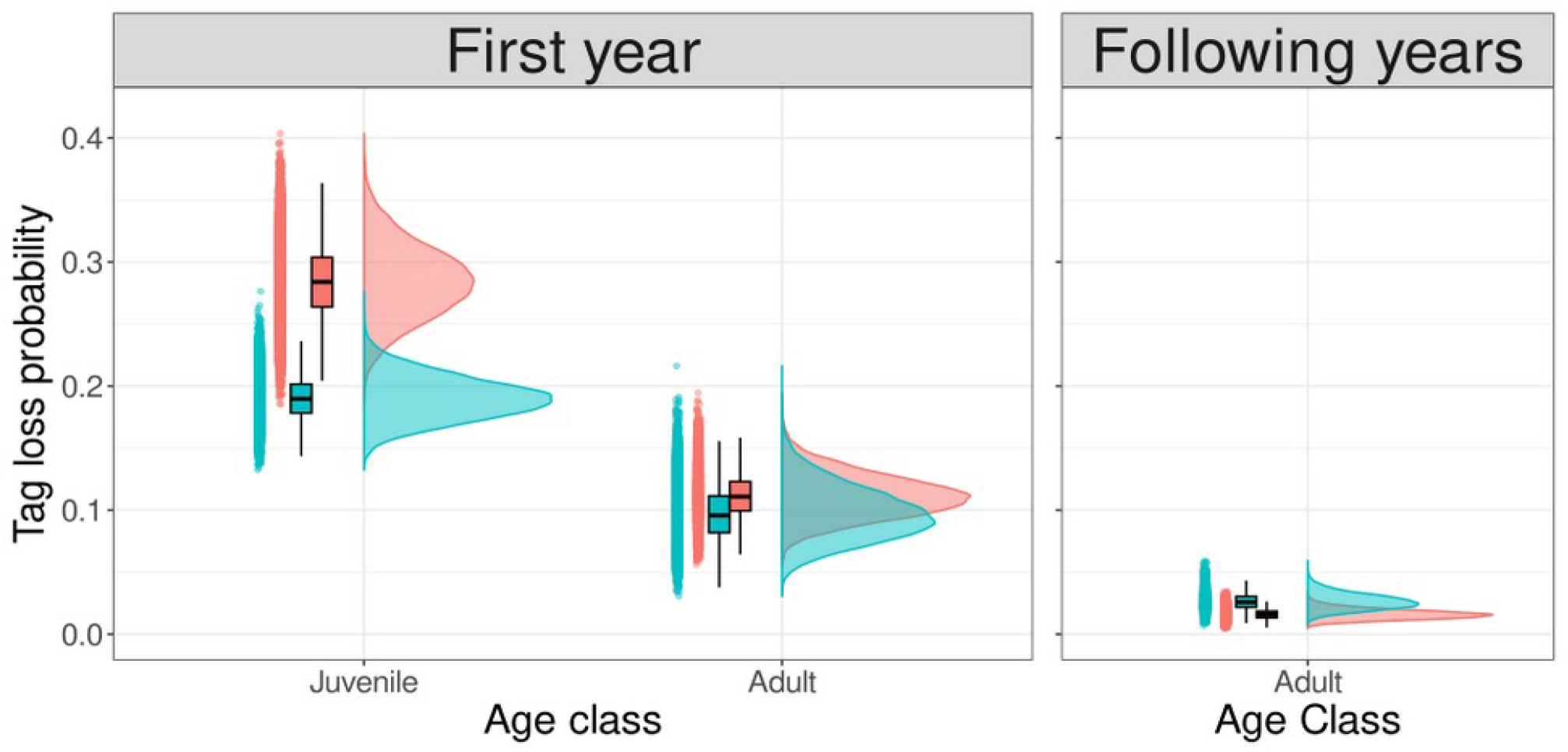
Posterior distribution of the tag loss probabilities according to age classes and time after marking in the Myotis myotis dataset. Left panel corresponds to tag shedding rate during the year following the tag injection and the right panel for the following years (constant in time). In blue, distribution if surgical adhesive was used after tag injection and in red, without surgical adhesive.

## 4 Discussion

The consideration of mark loss as a violation of CMR model assumptions has been the topic of numerous studies and model developments. Most of the existing research has focused on estimating survival, birth, or population size within the framework of Jolly-Seber models (Arnason & Mills, 1981; Malcolm-White *et al*., 2020; Schwarz *et al*., 2012; Smout *et al*., 2011a), recovery models (Kremers, 1988; Robson & Regier, 1966), CJS models (Laake *et al*., 2014; Nelson *et al*., 1980), and integrated population models (Riecke *et al*., 2019). However, there have been few proposed developments to account for mark loss specifically in AS models, such as those suggested by Besnard *et al*. (2007), Conn *et al*. (2004), and Johnson *et al*. (2016). These approaches have typically involved implicitly modelling mark status or using an adjustment factor, as seen in the work of Nishizawa *et al*. (2018). Additionally, we did not find any exploratory studies that have examined the impact of mark loss on parameter estimates. Therefore, in this study, we aim to fill this gap by exploring the effects of mark loss and recycled individuals on parameter estimates through simulations. We also examine the state of the mark (retained or lost) by modelling it as an independent Bernoulli process. By doing so, we can gain insights into how mark loss can affect the state transition of individuals when capture and survival probabilities vary over time, survival rates vary across age classes, and mark loss processes depend on the time since marking. Our results demonstrate that the violation of CMR model assumptions due to mark loss can significantly impact not only estimates of survival, but also capture, resighting, and state transition probabilities. Specifically, we found that survival is generally underestimated in cases where capture and detection rates are high. This underestimation becomes even more pronounced when survival rates are also high, thus moderating previous studies that suggested biases mainly occurred in species with high survival rates, catchability, and mark loss (Diefenbach & Alt, 1998). Our simulation results confirmed that the inaccuracy of model estimates is positively associated with the rate of mark loss. However, this inaccuracy can still occur even when the tag loss rate is low (e.g. 5%), and it can be independent of survival and capture rates. In datasets with a limited number of recycled individuals, which corresponds to low survival and capture rates, transition and resighting probabilities can be severely biased if mark loss is high. This implies that caution should be exercised when interpreting results from studies with low survival and capture rates if mark loss is suspected but not accounted for. Specifically, the probability of individuals remaining in the same state is underestimated when the transition from this state is low, but overestimated when the transition probabilities are high. The severity of these biases can also vary over time, with biases in survival and recapture decreasing over time as observed in our simulated datasets. This trend is partly influenced by the chosen mark loss pattern, highlighting the fact that even in studies conducted over short periods, parameters can still be substantially biased. It should be noted that if mark loss increases with time since marking (which was not investigated in this study), biases would be expected to increase accordingly.

The combination of simulation and empirical studies highlights how the complexity and interdependence of parameters can either exacerbate or mitigate estimation biases when mark loss modelling is absent. While the simulations revealed some general trends in the biases’ direction, the real-world example demonstrated the unexpected nature of bias patterns. It should also be noted that our simulation did not encompass the full range of parameter combinations that can be encountered in nature, despite aiming to cover demographic variations typically observed in vertebrates. This suggests that most study systems and monitoring methods have their own unique characteristics, and it is misleading to predict biases without simulating them. The uncertainty propagation in parameter estimates caused by mark loss remains challenging to predict and grows with system complexity. Hence, prior to planning a CMR study, we advocate for researchers and managers to conduct simulations to evaluate the conditions (i.e., parameter combinations) under which their study would yield reliable estimates of the parameters of interest (e.g., demographic, state transition). Preliminary studies with multiple marks could also be considered whenever possible (Smout *et al*., 2011a). This approach would allow for the optimization of CMR study design before its implementation, thereby minimizing biases from the outset.

Previous studies have developed AS models to estimate movement between sites, recruitment, dispersal, and temporary or permanent emigration (Lebreton *et al*., 2003, 2009; Schaub *et al*., 2004). Our simulation results indicate that state transition probabilities are sensitive to mark loss, even at low rates. For example, the probabilities of staying in the same state (philopatry, if transitions involve movement) or changing state (e.g., emigration) exhibited both underestimations and overestimations. These parameters are often essential in addressing ecological and demographic questions and used for management and conservation purposes (Cam *et al*., 2004; Horton *et al*., 2011). Although mark loss is frequently reported for various tag types and taxa, it is only marginally considered in studies focusing on estimating population dynamics parameters (Nelson *et al*., 1980; Ostrand *et al*., 2012; Smout *et al*., 2011b; Tavecchia *et al*., 2012). Most model developments to account for mark loss have focused on the JS model for abundance estimation, as mark loss and recycling in this model can introduce significant biases (Malcolm-White *et al*., 2020). Despite the growing use of AS models in ecology, demography, management, and conservation (Huntsman *et al*., 2020; Melnychuk *et al*., 2017), the issue of mark loss remains largely overlooked. Based on our study, we recommend the use of permanent or double temporary marks, ideally independent in terms of loss, or considering dependence in loss (Laake *et al*., 2014; McMahon & White, 2009; Schwarz *et al*., 2012). This is because any analysis of CMR data is potentially affected by this violation of model assumptions (Riecke *et al*., 2019).

Despite the fact that PIT tags are suitable for an increasing number of studies and allow data collection without physically re-capturing individuals, our case study emphasizes the importance of a second marking method to avoid potential bias in demographic rate estimations. Tag loss has long been known to occur in small mammal species, especially those that fly or glide (Freeland & Fry, 1995). In our study, we have confirmed that the use of surgical adhesive can reduce PIT tag shedding in the short term (Lebl & Ruf, 2010; van Harten *et al*., 2020).

However, our findings show that surgical adhesive alone is not enough to eliminate tag loss, and the use of additional data (e.g., evidence of tag loss) or a permanent mark (e.g., genotype) is necessary for the entire or a part of the studied population (Laake *et al*., 2014). Similar situations, where permanent marks should be considered, arise when marks deteriorate and become unreadable over time, such as with neck collars or ear tags (Conn *et al*., 2004; Diefenbach & Alt, 1998). In such cases, the accuracy of model parameter estimates is expected to decrease throughout the study duration, further supporting the use of permanent marks in capture-mark-recapture (CMR) studies.

Mark loss is typically not taken into account from ecological and management perspectives, except when researchers aim to understand factors influencing mark failures or improve their marking methods. Our results emphasize the need to assess the impact of mark loss whenever mark failure is suspected to avoid drawing misleading conclusions about the dynamics of the studied species. Based on our experience and the literature, PIT tags are prone to being shed regardless of the taxa being studied, often in the short term but sometimes even in the long term. A recent study on Gould’s wattled bats (*Chalinolobus gouldii*) over a relatively short period of 13-14 months reported low tag shedding (2.7%) and generalized this finding to all insectivorous bats (van Harten *et al*., 2020). However, our study shows that this generalization is partly incorrect and suggests that it is challenging to generalize such conclusions, as the pattern of mark loss is highly species-dependent, among other factors. Therefore, mark loss should be carefully considered in all CMR analyses and potentially in other studies using similar datasets. Explicit modelling of mark loss should be implemented when necessary to achieve more accurate estimations of population dynamics.

## Acknowledgements

This project was funded by an Irish Research Council Postdoctoral Fellowship Grant GOIPD/2018/256 awarded to F.T., a European Research Council Research Grant ERC-2012-StG311000 and an Irish Research Council Laureate Award awarded to E.C.T. The French field study was supported by the European Regional Development Fund EU000141 and a Contrat Nature grant awarded to Bretagne Vivante (BV). S.J.P. was supported by a Junior chair from the Institut Universitaire de France. We would like to thank the volunteers of BV for assistance in the fieldwork. Sampling was carried out in accordance with the French ethical and sampling guidelines issued in 3 successive permits delivered by the Préfet du Morbihan (Brittany) awarded to E.J.P., F.T. and S.J.P. for the period 2010–2020.

## Author contribution

F.T. designed the project with the other co-authors. E.C.T., F.T. acquired the funding. E.C.T., E.J.P., S.J.P. and E.C. supervised the project. E.C.T., F.T., E.J.P., and S.J.P supervised the fieldwork. E.C.T., E.J.P., F.T. and S.J.P. collected samples. S.J.P. & F.T. supervised the microsatelite genotyping. C.S. generated the genotypes. F.T. and E.C. developed the R scripts for simulating and analysing data with help of E.J.P. and S.J.P. F.T. led the writing of the manuscript and all authors contributed to manuscript revisions and gave final approval for publication.

### Data accessibility

R scripts for simulating the data, and analysing the data with JAGS, are accessible on Zenodo (https://zenodo.org/record/6453953).

## Notes

https://zenodo.org/record/6453953

### Competing Interest Statement

The authors have declared no competing interest.

### Summary of Updates

This version of the manuscript has been peer-reviewed and recommended by PCIEcology.

