## Supplementary Information 1 for "Mark loss can strongly bias demographic rates in multi-state models: a case study with simulated and empirical datasets"

### 1 Survival data in vertebrates

In order to cover a wide range of realistic case study of wild vertebrates in our simulation framework we gathered demographic data from published CMR studies and database. Most of the survival data for mammals, birds, reptiles, and amphibians come from the Demographic Species Knowledge Index (Conde *et al.* 2019) and were completed from literature in particular for fish and bat species (Lentini *et al.* 2015). Complementary data used are summarised in Table S1.

There is a wide range of values of survival rates, from 0.2 to over 0.97, across both age classes and taxa considered. For short-lived species, juveniles and adults have both low survival rate (Fig. S1). However, for longer-lived species, there is an greater dispersion in the data with increased survival in adults associated with both high or low survival in juveniles. To limit our simulation scenarios, we chose 2 cases where juvenile survival was low when adult survival was low, which we considered as short-lived species and the opposite as long-lived species.

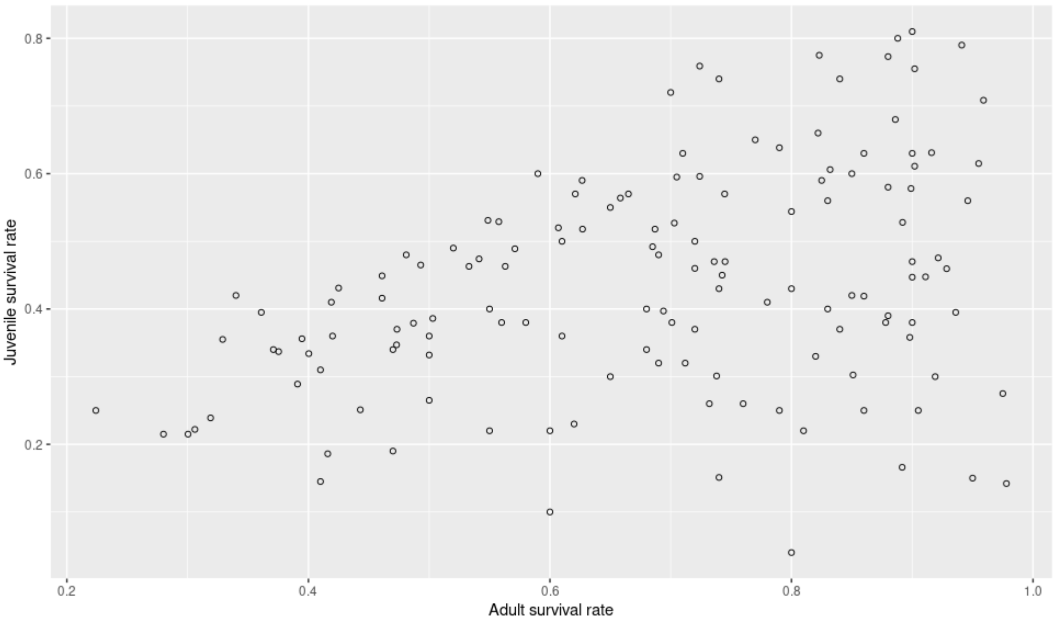


Figure S1: Relationship between adult and juvenile survival rates among 143 species.

| species | genus | family | order | class | phylum | kingdom | source | varname | varvalue |
|---|---|---|---|---|---|---|---|---|---|
| Odocoileus hemionus | Odocoileus | Cervidae | Artiodactyla | Mammalia | Chordata | Animalia | [1] | Adult_survival | 0.879 |
| Odocoileus hemionus | Odocoileus | Cervidae | Artiodactyla | Mammalia | Chordata | Animalia | [1] | Adult_survival | 0.823 |
| Rangifer tarandus | Rangifer | Cervidae | Artiodactyla | Mammalia | Chordata | Animalia | [1] | Adult_survival | 0.844 |
| Rangifer tarandus | Rangifer | Cervidae | Artiodactyla | Mammalia | Chordata | Animalia | [1] | Adult_survival | 0.94 |
| Tragelaphus strepsiceros | Tragelaphus | Cervidae | Artiodactyla | Mammalia | Chordata | Animalia | [1] | Adult_survival | 0.933 |
| Tragelaphus strepsiceros | Tragelaphus | Cervidae | Artiodactyla | Mammalia | Chordata | Animalia | [1] | Adult_survival | 0.889 |
| Ovis canadensis | Ovis | Cervidae | Artiodactyla | Mammalia | Chordata | Animalia | [1] | Adult_survival | 0.946 |
| Ovis canadensis | Ovis | Cervidae | Artiodactyla | Mammalia | Chordata | Animalia | [1] | Adult_survival | 0.911 |
| Capreolus capreolus | Capreolus | Cervidae | Artiodactyla | Mammalia | Chordata | Animalia | [1] | Adult_survival | 0.92 |
| Capreolus capreolus | capreolus | Cervidae | Artiodactyla | Mammalia | Chordata | Animalia | [1] | Adult_survival | 0.923 |
| Ovis aries | Ovis | Cervidae | Artiodactyla | Mammalia | Chordata | Animalia | [1] | Adult_survival | 0.902 |
| Alce alces | Alce | Cervidae | Artiodactyla | Mammalia | Chordata | Animalia | [1] | Adult_survival | 0.949 |
| Cervus elaphus | Cervus | Cervidae | Artiodactyla | Mammalia | Chordata | Animalia | [1] | Adult_survival | 0.955 |
| Oreamnos americanus | Oreamnos | Cervidae | Artiodactyla | Mammalia | Chordata | Animalia | [1] | Adult_survival | 0.916 |
| Equus ferus caballus | Equus | Cervidae | Artiodactyla | Mammalia | Chordata | Animalia | [1] | Adult_survival | 0.975 |
| Ovis dalli | Ovis | Cervidae | Artiodactyla | Mammalia | Chordata | Animalia | [1] | Adult_survival | 0.88 |
| Bos taurus | Bos | Cervidae | Artiodactyla | Mammalia | Chordata | Animalia | [1] | Adult_survival | 0.902 |
| Alce alces | Alce | Cervidae | Artiodactyla | Mammalia | Chordata | Animalia | [1] | Adult_survival | 0.976 |
| Alce alces | Alce | Cervidae | Artiodactyla | Mammalia | Chordata | Animalia | [1] | Adult_survival | 0.952 |
| Alce alces | Alce | Cervidae | Artiodactyla | Mammalia | Chordata | Animalia | [1] | Adult_survival | 0.976 |
| Capra sp | Capra | Cervidae | Artiodactyla | Mammalia | Chordata | Animalia | [1] | Adult_survival | 0.929 |
| Antilocapra americana | Antilocapra | Cervidae | Artiodactyla | Mammalia | Chordata | Animalia | [1] | Adult_survival | 0.978 |
| Odocoileus hemionus | Odocoileus | Cervidae | Artiodactyla | Mammalia | Chordata | Animalia | [1] | Juvenile_survival | 0.271 |
| Odocoileus hemionus | Odocoileus | Cervidae | Artiodactyla | Mammalia | Chordata | Animalia | [1] | Juvenile_survival | 0.334 |
| Rangifer tarandus | Rangifer | Cervidae | Artiodactyla | Mammalia | Chordata | Animalia | [1] | Juvenile_survival | 0.489 |
| Rangifer tarandus | Rangifer | Cervidae | Artiodactyla | Mammalia | Chordata | Animalia | [1] | Juvenile_survival | 0.567 |
| Tragelaphus strepsiceros | Tragelaphus | Cervidae | Artiodactyla | Mammalia | Chordata | Animalia | [1] | Juvenile_survival | 0.449 |
| Tragelaphus strepsiceros | Tragelaphus | Cervidae | Artiodactyla | Mammalia | Chordata | Animalia | [1] | Juvenile_survival | 0.446 |
| Ovis canadensis | Ovis | Cervidae | Artiodactyla | Mammalia | Chordata | Animalia | [1] | Juvenile_survival | 0.564 |
| Ovis canadensis | Ovis | Cervidae | Artiodactyla | Mammalia | Chordata | Animalia | [1] | Juvenile_survival | 0.355 |
| Capreolus capreolus | Capreolus | Cervidae | Artiodactyla | Mammalia | Chordata | Animalia | [1] | Juvenile_survival | 0.537 |
| Capreolus capreolus | capreolus | Cervidae | Artiodactyla | Mammalia | Chordata | Animalia | [1] | Juvenile_survival | 0.414 |
| Ovis aries | Ovis | Cervidae | Artiodactyla | Mammalia | Chordata | Animalia | [1] | Juvenile_survival | 0.611 |
| Alce alces | Alce | Cervidae | Artiodactyla | Mammalia | Chordata | Animalia | [1] | Juvenile_survival | 0.331 |
| Cervus elaphus | Cervus | Cervidae | Artiodactyla | Mammalia | Chordata | Animalia | [1] | Juvenile_survival | 0.615 |
| Oreamnos americanus | Oreamnos | Cervidae | Artiodactyla | Mammalia | Chordata | Animalia | [1] | Juvenile_survival | 0.631 |
| Equus ferus caballus | Equus | Cervidae | Artiodactyla | Mammalia | Chordata | Animalia | [1] | Juvenile_survival | 0.275 |
| Ovis dalli | Ovis | Cervidae | Artiodactyla | Mammalia | Chordata | Animalia | [1] | Juvenile_survival | 0.773 |
| Bos taurus | Bos | Cervidae | Artiodactyla | Mammalia | Chordata | Animalia | [1] | Juvenile_survival | 0.755 |
| Alce alces | Alce | Cervidae | Artiodactyla | Mammalia | Chordata | Animalia | [1] | Juvenile_survival | 0.813 |
| Alce alces | Alce | Cervidae | Artiodactyla | Mammalia | Chordata | Animalia | [1] | Juvenile_survival | 0.865 |
| Alce alces | Alce | Cervidae | Artiodactyla | Mammalia | Chordata | Animalia | [1] | Juvenile_survival | 0.825 |
| Capra sp | Capra | Cervidae | Artiodactyla | Mammalia | Chordata | Animalia | [1] | Juvenile_survival | 0.867 |
| Antilocapra americana | Antilocapra | Cervidae | Artiodactyla | Mammalia | Chordata | Animalia | [1] | Juvenile_survival | 0.142 |
| Common pipistrelle | Pipistrellus | Vespertilionidae | Chiroptera | Mammalia | Chordata | Animalia | [2] | Juvenile_survival | 0.527 |
| Common pipistrelle | Pipistrellus | Vespertilionidae | Chiroptera | Mammalia | Chordata | Animalia | [2] | Adult_survival | 0.799 |
| Nyctalus leisleri | Nyctalus | Vespertilionidae | Chiroptera | Mammalia | Chordata | Animalia | [3] | Adult_survival | 0.91 |

| species | genus | family | order | class | phylum | kingdom | source | varname | varvalue |
|---|---|---|---|---|---|---|---|---|---|
| Nyctalus leisleri | Nyctalus | Vespertilionidae | Chiroptera | Mammalia | Chordata | Animalia | [3] | Adult_survival | 0.613 |
| Halichoerus grypus | Nyctalus | Phocidae | Carnivora | Mammalia | Chordata | Animalia | [4] | Adult_survival | 0.89 |
| Halichoerus grypus | Nyctalus | Phocidae | Carnivora | Mammalia | Chordata | Animalia | [4] | Adult_survival | 0.97 |
| Carcharodon carcharias | Carcharodon | Lamnidae | Lamniformes | Chondrichthyesa | Chordata | Animalia | [5] | Juvenile_survival | 0.632 |
| Dipterus cf. intermedia | Dipterus | Rajidae | Rajiformes | Chondrichthyesa | Chordata | Animalia | [5] | Annual_survival | 0.632 |
| Rhincodon typus | Rhincodon | Rhincodontidae | Orectolobiformes | Chondrichthyesa | Chordata | Animalia | [6] | Juvenile_survival | 0.59 |
| Rhincodon typus | Rhincodon | Rhincodontidae | Orectolobiformes | Chondrichthyesa | Chordata | Animalia | [7] | Adult_survival | 0.825 |
| Fundulus heteroclitus | Fundulus | Fundulidae | Cyprinodontiformes | Actinopterygii | Chordata | Animalia | [8] | Annual_survival | 0.87 |
| Acipenser oxyrinchus oxyrinchus | Acipenser | Acipenseridae | Acipenseriformes | Actinopterygii | Chordata | Animalia | [9] | Annual_survival | 0.88 |
| Macroderma gigas | Macroderma | Megadermatidae | Chiroptera | Mammalia | Chordata | Animalia | [10] | juvenileSurvival.avg | 0.492 |
| Macroderma gigas | Macroderma | Megadermatidae | Chiroptera | Mammalia | Chordata | Animalia | [10] | adultSurvival.avg | 0.685 |
| Myotis myotis | Myotis | Vespertilionidae | Chiroptera | Mammalia | Chordata | Animalia | [11] | adultSurvival.avg | 0.719 |
| Myotis myotis | Myotis | Vespertilionidae | Chiroptera | Mammalia | Chordata | Animalia | [11] | adultSurvival.avg | 0.708 |
| Chalinolobus morio | Chalinolobus | Vespertilionidae | Chiroptera | Mammalia | Chordata | Animalia | [12] | adultSurvival.avg | 0.72 |
| Nyctophilus geoffroyi | Nyctophilus | Vespertilionidae | Chiroptera | Mammalia | Chordata | Animalia | [12] | adultSurvival.avg | 0.42 |
| Vespadelus vulturnus | Vespadelus | Vespertilionidae | Chiroptera | Mammalia | Chordata | Animalia | [12] | adultSurvival.avg | 0.59 |
| Eptesicus fuscus | Eptesicus | Vespertilionidae | Chiroptera | Mammalia | Chordata | Animalia | [13] | juvenileSurvival.avg | 0.77 |
| Myotis sodalis | Myotis | Vespertilionidae | Chiroptera | Mammalia | Chordata | Animalia | [14] | adultSurvival.avg | 0.624 |
| Corynorhinus townsendii | Corynorhinus | Vespertilionidae | Chiroptera | Mammalia | Chordata | Animalia | [15] | adultSurvival.avg | 0.6 |
| Corynorhinus townsendii | Corynorhinus | Vespertilionidae | Chiroptera | Mammalia | Chordata | Animalia | [15] | adultSurvival.avg | 0.58 |
| Corynorhinus townsendii | Corynorhinus | Vespertilionidae | Chiroptera | Mammalia | Chordata | Animalia | [15] | adultSurvival.avg | 0.65 |
| Corynorhinus townsendii | Corynorhinus | Vespertilionidae | Chiroptera | Mammalia | Chordata | Animalia | [15] | adultSurvival.avg | 0.54 |
| Corynorhinus townsendii | Corynorhinus | Vespertilionidae | Chiroptera | Mammalia | Chordata | Animalia | [15] | adultSurvival.avg | 0.67 |
| Corynorhinus townsendii | Corynorhinus | Vespertilionidae | Chiroptera | Mammalia | Chordata | Animalia | [15] | adultSurvival.avg | 0.67 |
| Eptesicus fuscus | Eptesicus | Vespertilionidae | Chiroptera | Mammalia | Chordata | Animalia | [16] | adultSurvival.avg | 0.68 |
| Eptesicus fuscus | Eptesicus | Vespertilionidae | Chiroptera | Mammalia | Chordata | Animalia | [16] | juvenileSurvival.avg | 0.52 |
| Eptesicus fuscus | Eptesicus | Vespertilionidae | Chiroptera | Mammalia | Chordata | Animalia | [16] | adultSurvival.avg | 0.81 |
| Eptesicus fuscus | Eptesicus | Vespertilionidae | Chiroptera | Mammalia | Chordata | Animalia | [16] | juvenileSurvival.avg | 0.69 |
| Eptesicus fuscus | Eptesicus | Vespertilionidae | Chiroptera | Mammalia | Chordata | Animalia | [16] | adultSurvival.avg | 0.87 |
| Eptesicus fuscus | Eptesicus | Vespertilionidae | Chiroptera | Mammalia | Chordata | Animalia | [16] | juvenileSurvival.avg | 0.5 |
| Eptesicus fuscus | Eptesicus | Vespertilionidae | Chiroptera | Mammalia | Chordata | Animalia | [16] | adultSurvival.avg | 0.86 |
| Eptesicus fuscus | Eptesicus | Vespertilionidae | Chiroptera | Mammalia | Chordata | Animalia | [16] | juvenileSurvival.avg | 0.76 |
| Eptesicus fuscus | Eptesicus | Vespertilionidae | Chiroptera | Mammalia | Chordata | Animalia | [16] | adultSurvival.avg | 0.73 |
| Eptesicus fuscus | Eptesicus | Vespertilionidae | Chiroptera | Mammalia | Chordata | Animalia | [16] | juvenileSurvival.avg | 0.59 |
| Myotis yumanensis | Myotis | Vespertilionidae | Chiroptera | Mammalia | Chordata | Animalia | [17] | adultSurvival.avg | 0.84 |
| Myotis yumanensis | Myotis | Vespertilionidae | Chiroptera | Mammalia | Chordata | Animalia | [17] | juvenileSurvival.avg | 0.74 |
| Pipistrellus pipistrellus | Pipistrellus | Vespertilionidae | Chiroptera | Mammalia | Chordata | Animalia | [18] | adultSurvival.avg | 0.56 |
| Pipistrellus pipistrellus | Pipistrellus | Vespertilionidae | Chiroptera | Mammalia | Chordata | Animalia | [18] | adultSurvival.avg | 0.75 |
| Eptesicus fuscus | Eptesicus | Vespertilionidae | Chiroptera | Mammalia | Chordata | Animalia | [19] | Survival | 0.465 |
| Myotis leibii | Myotis | Vespertilionidae | Chiroptera | Mammalia | Chordata | Animalia | [19] | Survival | 0.697 |
| Eptesicus fuscus | Eptesicus | Vespertilionidae | Chiroptera | Mammalia | Chordata | Animalia | [19] | Survival | 0.421 |
| Myotis leibii | Myotis | Vespertilionidae | Chiroptera | Mammalia | Chordata | Animalia | [19] | Survival | 0.757 |
| Myotis lucifugus | Myotis | Vespertilionidae | Chiroptera | Mammalia | Chordata | Animalia | [20] | Survival | 0.708 |
| Myotis lucifugus | Myotis | Vespertilionidae | Chiroptera | Mammalia | Chordata | Animalia | [20] | Survival | 0.816 |
| Chalinolobus tuberculatus | Chalinolobus | Vespertilionidae | Chiroptera | Mammalia | Chordata | Animalia | [21] | adultSurvival.avg | 0.75 |
| Myotis capaccinii | Myotis | Vespertilionidae | Chiroptera | Mammalia | Chordata | Animalia | [22] | adultSurvival.avg | 0.943 |
| Myotis capaccinii | Myotis | Vespertilionidae | Chiroptera | Mammalia | Chordata | Animalia | [22] | adultSurvival.avg | 0.903 |

| species | genus | family | order | class | phylum | kingdom | source | varname | varvalue |
|---|---|---|---|---|---|---|---|---|---|
| Eptesicus isabellinus | Eptesicus | Vespertilionidae | Chiroptera | Mammalia | Chordata | Animalia | [23] | adultSurvival.avg | 0.7 |
| Eptesicus isabellinus | Eptesicus | Vespertilionidae | Chiroptera | Mammalia | Chordata | Animalia | [23] | adultSurvival.avg | 0.81 |
| Eptesicus isabellinus | Eptesicus | Vespertilionidae | Chiroptera | Mammalia | Chordata | Animalia | [23] | adultSurvival.avg | 0.71 |
| Eptesicus isabellinus | Eptesicus | Vespertilionidae | Chiroptera | Mammalia | Chordata | Animalia | [23] | adultSurvival.avg | 0.61 |
| Eptesicus isabellinus | Eptesicus | Vespertilionidae | Chiroptera | Mammalia | Chordata | Animalia | [23] | adultSurvival.avg | 0.58 |
| Myotis bechsteinii | Myotis | Vespertilionidae | Chiroptera | Mammalia | Chordata | Animalia | [24] | adultSurvival.avg | 0.79 |
| Myotis daubentonii | Myotis | Vespertilionidae | Chiroptera | Mammalia | Chordata | Animalia | [24] | adultSurvival.avg | 0.79 |
| Nyctalus lasiopterus | Nyctalus | Vespertilionidae | Chiroptera | Mammalia | Chordata | Animalia | [24] | adultSurvival.avg | 0.74 |
| Nyctalus leisleri | Nyctalus | Vespertilionidae | Chiroptera | Mammalia | Chordata | Animalia | [24] | adultSurvival.avg | 0.74 |
| Plecotus auritus | Plecotus | Vespertilionidae | Chiroptera | Mammalia | Chordata | Animalia | [24] | adultSurvival.avg | 0.79 |
| Chalinolobus tuberculatus | Chalinolobus | Vespertilionidae | Chiroptera | Mammalia | Chordata | Animalia | [25] | adultSurvival.avg | 0.89 |
| Chalinolobus tuberculatus | Chalinolobus | Vespertilionidae | Chiroptera | Mammalia | Chordata | Animalia | [25] | adultSurvival.avg | 0.55 |
| Chalinolobus tuberculatus | Chalinolobus | Vespertilionidae | Chiroptera | Mammalia | Chordata | Animalia | [25] | adultSurvival.avg | 0.5 |
| Chalinolobus tuberculatus | Chalinolobus | Vespertilionidae | Chiroptera | Mammalia | Chordata | Animalia | [25] | adultSurvival.avg | 0.91 |
| Chalinolobus tuberculatus | Chalinolobus | Vespertilionidae | Chiroptera | Mammalia | Chordata | Animalia | [26] | adultSurvival.avg | 0.77 |
| Chalinolobus tuberculatus | Chalinolobus | Vespertilionidae | Chiroptera | Mammalia | Chordata | Animalia | [26] | adultSurvival.avg | 0.65 |
| Chalinolobus tuberculatus | Chalinolobus | Vespertilionidae | Chiroptera | Mammalia | Chordata | Animalia | [26] | juvenileSurvival.avg | 0.54 |
| Chalinolobus tuberculatus | Chalinolobus | Vespertilionidae | Chiroptera | Mammalia | Chordata | Animalia | [26] | adultSurvival.avg | 0.79 |
| Chalinolobus tuberculatus | Chalinolobus | Vespertilionidae | Chiroptera | Mammalia | Chordata | Animalia | [26] | juvenileSurvival.avg | 0.71 |
| Chalinolobus tuberculatus | Chalinolobus | Vespertilionidae | Chiroptera | Mammalia | Chordata | Animalia | [26] | adultSurvival.avg | 0.58 |
| Chalinolobus tuberculatus | Chalinolobus | Vespertilionidae | Chiroptera | Mammalia | Chordata | Animalia | [26] | juvenileSurvival.avg | 0.47 |
| Chalinolobus tuberculatus | Chalinolobus | Vespertilionidae | Chiroptera | Mammalia | Chordata | Animalia | [26] | adultSurvival.avg | 0.75 |
| Chalinolobus tuberculatus | Chalinolobus | Vespertilionidae | Chiroptera | Mammalia | Chordata | Animalia | [26] | juvenileSurvival.avg | 0.71 |
| Chalinolobus tuberculatus | Chalinolobus | Vespertilionidae | Chiroptera | Mammalia | Chordata | Animalia | [26] | adultSurvival.avg | 0.67 |
| Chalinolobus tuberculatus | Chalinolobus | Vespertilionidae | Chiroptera | Mammalia | Chordata | Animalia | [26] | juvenileSurvival.avg | 0.57 |
| Chalinolobus tuberculatus | Chalinolobus | Vespertilionidae | Chiroptera | Mammalia | Chordata | Animalia | [26] | adultSurvival.avg | 0.81 |
| Chalinolobus tuberculatus | Chalinolobus | Vespertilionidae | Chiroptera | Mammalia | Chordata | Animalia | [26] | juvenileSurvival.avg | 0.74 |
| Myotis nattereri | Myotis | Vespertilionidae | Chiroptera | Mammalia | Chordata | Animalia | [27] | Survival | 0.86 |
| Rhinolophus ferrumequinum | Rhinolophus | Rhinolophidae | Chiroptera | Mammalia | Chordata | Animalia | [28] | Survival | 0.49 |
| Rhinolophus ferrumequinum | Rhinolophus | Rhinolophidae | Chiroptera | Mammalia | Chordata | Animalia | [28] | Survival | 0.91 |
| Nyctalus leisleri | Nyctalus | Vespertilionidae | Chiroptera | Mammalia | Chordata | Animalia | [29] | juvenileSurvival.avg | 0.45 |
| Nyctalus leisleri | Nyctalus | Vespertilionidae | Chiroptera | Mammalia | Chordata | Animalia | [29] | adultSurvival.avg | 0.76 |
| Nyctalus leisleri | Nyctalus | Vespertilionidae | Chiroptera | Mammalia | Chordata | Animalia | [29] | adultSurvival.avg | 0.69 |

Table S1: Complementary data to Conde et al. (2019) used to explore cross taxa survival rates (varvalue) in vertebrates, according to age class (varname).

### 2 Tag loss data in vertebrates

| Species | Scientific Name | Class | Mark type | Sample size | Loss rate | Duration | Location | Conditions | Control | Source |
|---|---|---|---|---|---|---|---|---|---|---|
| Brown trout | Salmo trutta | Actinopterygii | PIT tag | 145 | 0.2-0.3 | 4 weeks | Peritoneal cavity | Laboratory | None | [30] |
| Bluegills | Lepomis macrochirus | Actinopterygii | PIT tag | 20 | 0-0.4 | 42 days | Dorsal musculature | Laboratory | None | [31] |
| Bluegills | Lepomis macrochirus | Actinopterygii | PIT tag | 20 | 0.1-0.2 | 42 days | Peritoneal cavity | Laboratory | None | [31] |
| Bluegills | Lepomis macrochirus | Actinopterygii | PIT tag | 20 | 0.4-1 | 42 days | Isthmus | Laboratory | None | [31] |
| Yellow perch | Perca flavescens | Actinopterygii | PIT tag | 20 | 0-0.5 | 42 days | Dorsal musculature | Laboratory | None | [31] |
| Yellow perch | Perca flavescens | Actinopterygii | PIT tag | 20 | 0-0.2 | 42 days | Peritoneal cavity | Laboratory | None | [31] |
| Yellow perch | Perca flavescens | Actinopterygii | PIT tag | 20 | 0.4-1 | 42 days | Isthmus | Laboratory | None | [31] |
| Muskellunge | Esox masquinongy | Actinopterygii | PIT tag | 300 | 0-0.005 | 48 hours | Peritoneal cavity | Laboratory | None | [32] |
| Muskellunge | Esox masquinongy | Actinopterygii | PIT tag | 300 | 0-0.011 | 48 hours | Dorsal musculature | Laboratory | None | [32] |
| Steelhead trout | Oncorhynchus mykiss | Actinopterygii | Acoustic tags | 80 | 0.2 | 143 days | Peritoneal cavity | Laboratory | None | [33] |
| Lake trout | Salvelinus namaycush | Actinopterygii | Anchor tag | 660 | 0.06-0.5 | 19 years | Dorsal musculature | Wild | Double tag | [34] |
| Hellbenders | Cryptobranchus alleganiensis | Amphibia | PIT tag | 78 | 0 | 2 years | Tail musculature | Wild | Genotype | [35] |
| Common guillemots | Uria aalge | Aves | Metal ring | 5594 | 0.24 | 20 years | Tarsus | Wild | None | [36] |
| Greater snow goose | Chen caerulescens atlantica | Aves | Neck collar | 7799 | 0.05 | 12 years | Neck | Wild | Metal ring | [37] |
| Lesser snow goose | Chen caerulescens caerulescens | Aves | Neck collar | 3078 | 0.45-0.65 | 6 years | Neck | Wild | Metal ring | [38] |
| Red kites | Milvus milvus | Aves | Radiotransmitter | 142 | 0.11 | 11 years | Backpack harness | Wild | Plastic wing tag | [39] |
| Abalone | Haliotis kamtschatkana | Gastropoda | PIT tag | 62 | 0.1-0.9 | 6-15 days | Shell,/foot muscle | Laboratory | None | [40] |
| Grasshoppers | Prionotropis hystrix rhodanica | Insecta | Radio-tracking | 442 | 0.06-0.09 | 36 days | Pronotum (Glued) | Wild | None | [41] |
| New Zealand fur seal | Arctocephalus forsteri | Mammalia | Plastic tags | 1255 | 0-0.6 | 3 years | Fore flipper | Wild | Double tag | [42] |
| Elephant seal | Mirounga leonina | Mammalia | Plastic tags | 8567 | 0.2 | 7 years | Hind flipper | Wild | Hot-iron branding | [43] |
| Elephant seals | Mirounga leonina | Mammalia | Plastic tags | 14000 | 0-0.4 | 17 years | Hind flipper | Wild | Hot-iron branding | [44] |
| Grey seals | Halichoerus grypus | Mammalia | Plastic tags | 725 | 0.04-0.27 | 28 years | Flipper | Wild | Double tag/brand | [45] |
| Black bears | Ursus americanus | Mammalia | Plastic tags | 298 | 0.094 | 12 years | Ear | Wild | Double tag/tattoo | [46] |
| Elephant seal | Mirounga leonina | Mammalia | Plastic tags | 8568 | 0-0.364 | 8 years | Hind flipper | Wild | Double tag/brand | [47] |
| Silvery marmoset | Callithrix argentata | Mammalia | PIT tag | 4 | 0.75 | 8 months | ? | Zoo | None | [48] |
| White-handed gibbon | Hylobates lar | Mammalia | PIT tag | 2 | 0 | 10 months | ? | Zoo | None | [48] |
| Naked mole rat | Heterocephalus glaber | Mammalia | PIT tag | 2 | 0 | 8 months | ? | Zoo | None | [48] |
| Himalayan tahr | Hemitragus jemlahicus | Mammalia | PIT tag | 9 | 0 | 10 months | ? | Zoo | None | [48] |
| Corn snake | Elaphe gutatta | Reptilia | PIT tag | 15 | 0.53 | 9 weeks | neck region | Laboratory | None | [49] |

Table S2: Non exhaustive list of tag loss rate from bibliography. Only studies using CMR methods or studies in control conditions (Laboratory and Zoo) are reported. The mark type, the sample size of the individuals marked, the duration of the study and the location of the mark were reported. If a method is used as permanent mark, the type of 'control' mark is indicated.

#### Sources in Table S1 & S2:

[10] Hoyle, S. D., Pople, A. R., & Toop, G. J. (2001). Mark–recapture may reveal more about ecology than about population trends: Demography of a threatened ghost bat (Macroderma gigas) population. Austral Ecology, 26(1), 80–92.
https://doi.org/10.1111/ j.1442-9993.2001.01092.pp.x

[11] Amengual, B., Bourhy, H., Lopez-Roig, M., & Serra-Cobo, J. (2007). Temporal Dynamics of European Bat Lyssavirus Type 1 and Survival of Myotis myotis Bats in Natural Colonies. Plos One, 6(2).
https://doi.org/10.1371/journal.pone.0000566

[12] Baker, G. B., Lumsden, L. F., Dettmann, E. B., Schedvin, N. K., Schulz, M., Watkins, D., & Jansen, L. (2001). The effect of forearm bands on insectivorous bats (Microchiroptera) in Australia. Wildlife Research, 28(3), 229–237.
https://doi.org/10.1071/ wr99068

[13] Beer, J. R. (1955). Survival and Movements of Banded Big Brown Bats. Journal of Mammalogy, 36(2), 242–248.
https://doi.org/10.2307/1375883

[14] Boyles, J. G., Walters, B. L., Whitaker, J. O., & Cope, J. B. (2007). A reanalysis of apparent survival rates of Indiana myotis (Myotis sodalis). Acta Chiropterologica, 9(1), 127–132.
https://doi.org/10.3161/1733-5329(2007)9[127:AROASR]2.0.CO;2

[15] Ellison, L. E. (2010). A Retrospective Survival Analysis of Townsend's Big-Eared Bat (Corynorhinus townsendii) from Washington State. Northwestern Naturalist, 91(2), 172–182.
https://doi.org/10.1898/NWN09-10.1

[16] Ellison, L. E., O'Shea, T. J., Neubaum, D. J., Neubaum, M. A., Pearce, R. D., & Bowen, R. A. (2007). A comparison of conventional capture versus PIT reader techniques for estimating survival and capture probabilities of big brown bats (Eptesicus fuscus). Acta Chiropterologica, 9(1), 149–160.
https://doi.org/10.3161/1733-5329(2007)9 [149:ACOCCV]2.0.CO;2

[17] Frick, W. F., Rainey, W. E., & Pierson, E. D. (2007). Potential Effects of Environmental Contamination on Yuma Myotis Demography and Population Growth. Ecological Applications, 17(4), 1213–1222.
https://doi.org/10.1890/06-1021

[18] Gerell, R., & Lundberg, K. (1990). Sexual Differences in Survival Rates of Adult Pipistrelle Bats (Pipistrellus pipistrellus) in South Sweden. Oecologia, 83(3), 401–404.

[19] Hitchcock, H. B., Keen, R., & Kurta, A. (1984). Survival Rates of Myotis leibii and Eptesicus fuscus in Southeastern Ontario. Journal of Mammalogy, 65(1), 126–130.
https://doi.org/10.2307/1381210

[20] Humphrey, S. R., & Cope, J. B. (1976). Population ecology of the little brown bat, Myotis lucifugus, in Indiana and north-central Kentucky. American Society of Mammalogists. http://archive.org/details/populationecolog00hump

[21] O'Donnell, C. F. J. (2002). Timing of breeding, productivity and survival of long-tailed bats Chalinolobus tuberculatus (Chiroptera: Vespertilionidae) in cold-temperate rainforest in New Zealand. Journal of Zoology, 257(3), 311–323.
https://doi.org/10.1017/ S0952836902000912

[22] Papadatou, E., Butlin, R. K., Pradel, R., & Altringham, J. D. (2009). Sex-specific roost movements and population dynamics of the vulnerable long-fingered bat, Myotis capaccinii. Biological Conservation, 142(2), 280–289.

[23] Papadatou, Elena, Pradel, R., Schaub, M., Dolch, D., Geiger, H., Ibanez, C., Kerth, G., Popa-Lisseanu, A., Schorcht, W., Teubner, J., & Gimenez, O. (2012). Comparing survival among species with imperfect detection using multilevel analysis of markure-capture data: A case study on bats. Ecography, 35(2), 153–161.
https://doi.org/10.1111/ j.1600-0587.2011.07084.x

[24] Papadatou, Eleni, Ibáñez, C., Pradel, R., Juste, J., & Gimenez, O. (2011). Assessing survival in a multi-population system: A case study on bat populations. Oecologia,

165(4), 925–933.
https://doi.org/10.1007/s00442-010-1771-5

[25] Pryde, M. A, O'Donnell, C. F. J., & Barker, R. J. (2005). Factors influencing survival and long-term population viability of New Zealand long-tailed bats (Chalinolobus tuberculatus): Implications for conservation. Biological Conservation, 126(2), 175–185.

[26] Pryde, Moira A., Lettink, M., & O'Donnell, C. F. J. (2006). Survivorship in two populations of long-tailed bats (Chalinolobus tuberculatus) in New Zealand. New Zealand Journal of Zoology, 33(2), 85–95.
https://doi.org/10.1080/03014223.2006.9518433

[27] Rivers, N. M., Butlin, R. K., & Altringham, J. D. (2006). Autumn swarming behaviour of Natterer's bats in the UK: Population size, catchment area and dispersal. Biological Conservation, 127(2), 215–226.
https://doi.org/10.1016/j.biocon.2005.08.010

[28] Schaub, M., Gimenez, O., Sierro, A., & Arlettaz, R. (2007). Use of integrated modeling to enhance estimates of population dynamics obtained from limited data. Conservation Biology, 21(4), 945–955.

[29] Schorcht, W., Bontadina, F., & Schaub, M. (2009). Variation of adult survival drives population dynamics in a migrating forest bat. Journal of Animal Ecology, 78(6), 1182–1190.

[30] Acolas, M. L., Roussel, J. M., Lebel, J. M., and Bagliniere, J. L. (2007). Laboratory experiment on survival, growth and tag retention following PIT injection into the body cavity of juvenile brown trout (Salmo trutta). Fisheries Research, 86(2–3), 280–284.
https://doi.org/10.1016/j.fishres.2007.05.011

[31] Kaemingk, M. A., Weber, M. J., McKenna, P. R., and Brown, M. L. (2011). Effect of Passive Integrated Transponder Tag Implantation Site on Tag Retention, Growth, and Survival of Two Sizes of Juvenile Bluegills and Yellow Perch. North American Journal of Fisheries Management, 31(4), 726–732.
https://doi.org/10.1080/02755947.2011.611863

[32] Wagner, C. P., Jennings, M. J., Kampa, J. M., and Wahl, D. H. (2007). Survival, growth, and tag retention in age-0 Muskellunge implanted with passive integrated transponders. North American Journal of Fisheries Management, 27(3), 873–877.
https://doi.org/10.1577/M06-196.1

[33] Sandstrom, P. T., Ammann, A. J., Michel, C., Singer, G., Chapman, E. D., Lindley, S., … Klimley, A. P. (2013). Growth, survival, and tag retention of steelhead trout (Oncorhynchus mykiss) and its application to survival estimates. Environmental Biology of Fishes, 96(2–3), 145–164.
https://doi.org/10.1007/s10641-012-0051-0

[34] Fabrizio, M. C., Nichols, J. D., Hines, J., Swanson, B. L., and Schram, S. T. (1999). Modeling data from double-tagging experiments to estimate heterogeneous rates of tag shedding in lake trout (Salvelinus namaycush).
https://doi.org/10.1139/F99-069

[35]Unger, S. D., Burgmeier, N. G., and Williams, R. N. (2012). Genetic markers reveal high PIT tag retention rates in giant salamanders (Cryptobranchus alleganiensis). Amphibia-Reptilia, 33(2), 313–317.
https://doi.org/10.1163/156853812X641712

[36] Reynolds, T. J., King, R., Harwood, J., Frederiksen, M., Harris, M. P., and Wanless, S. (2009). Integrated data analysis in the presence of emigration and mark loss. Journal of Agricultural, Biological, and Environmental Statistics, 14(4), 411–431.
https://doi.org/10.1198/jabes.2009.080089

[37] Juillet, C., Choquet, R., Gauthier, G., and Pradel, R. (2011). A Capture–Recapture

model with double-marking, live and dead encounters, and heterogeneity of reporting due to auxiliary mark loss. Journal of Agricultural, Biological, and Environmental Statistics, 16(1), 88–104.
https://doi.org/10.1007/s13253-010-0035-5

[38] Conn, P. B., Kendall, W. L., and Samuel, M. D. (2004). A General Model for the Analysis of Mark-Resight, Mark-Recapture, and Band-Recovery Data under Tag Loss. Biometrics, 60(4), 900–909.
https://doi.org/10.1111/j.0006-341X.2004.00245.x

[39] Tavecchia, G., Adrover, J., Navarro, A. M., and Pradel, R. (2012). Modelling mortality causes in longitudinal data in the presence of tag loss: application to raptor poisoning and electrocution. Journal of Applied Ecology, 49(1), 297–305.
https://doi.org/10.1111/j.1365-2664.2011.02074.x

[40] Hale, J. R., Bouma, J. V., Vadopalas, B., and Friedman, C. S. (2012). Evaluation of Passive Integrated Transponders for Abalone: Tag Placement, Retention, and Effect on Survival. Journal of Shellfish Research, 31(3), 789–794.
https://doi.org/10.2983/035.031.0324

[41] Besnard, A., Piry, S., Berthier, K., Lebreton, J.-D., and Streiff, R. (2007). Modeling survival and mark loss in molting animals: Recapture, dead recoveries, and exuvia recoveries. Ecology, 88(2), 289–295.
https://doi.org/10.1890/05-0811

[42] Bradshaw, C. J. A., Bark6er, R. J., and Davis, L. S. (2000). Modeling tag loss in New Zealand fur seal pups. Journal of Agricultural Biological and Environmental Statistics, 5(4), 475–485.
https://doi.org/10.2307/1400661

[43] Malcolm-White, E., McMahon, C. R., and Cowen, L. L. E. (2020). Complete tag loss in capture–recapture studies affects abundance estimates: An elephant seal case study. Ecology and Evolution, 10(5), 2377–2384.
https://doi.org/10.1002/ece3.6052

[44] Schwarz, L. K., Hindell, M. A., McMahon, C. R., and Costa, D. P. (2012). The implications of assuming independent tag loss in southern elephant seals. Ecosphere, 3(9), art81.
https://doi.org/10.1890/ES12-00132.1

[45] Smout, S., King, R., and Pomeroy, P. (2011). Integrating heterogeneity of detection and mark loss to estimate survival and transience in UK grey seal colonies. Journal of Applied Ecology, 48(2), 364–372.
https://doi.org/10.1111/j.1365-2664.2010.01913.x

[46] Laake, J. L., Johnson, D. S., Diefenbach, D. R., and Ternent, M. A. (2014). Hidden Markov Model for Dependent Mark Loss and Survival Estimation. Journal of Agricultural Biological and Environmental Statistics, 19(4), 524–540.
https://doi.org/10.1007/s13253-014-0190-1

[47] McMahon, C. R., and White, G. C. (2009). Tag loss probabilities are not independent: Assessing and quantifying the assumption of independent tag transition probabilities from direc9t observations. Journal of Experimental Marine Biology and Ecology, 372(1–2), 36–42.
https://doi.org/10.1016/j.jembe.2009.02.006

[48] Elbin, S., and Burger, J. (1994). Implantable Microchips for Individual Identification in Wild and Captive Populations. Wildlife Society Bulletin, 22(4), 677–683.

[49] Roark, A. W., and Dorcas, M. E. (2000). Regional Body Temperature Variation in Corn Snakes Measured Using Temperature-Sensitive Passive Integrated Transponders. Journal of Herpetology, 34(3), 481–485.

https://doi.org/10.2307/1565378
