## Supplementary Information 2 for "Mark loss can strongly bias demographic rates in multi-state models: a case study with simulated and empirical datasets"

4 **ROPE estimation for EMDs** **95**

### 1 Data simulation without recycled individuals and modelling

#### 1.1 Experimental design

To designed our simulations, we considered a set of parameter values inspired from our bibliographic review (Supporting Information 1), a range of values that might be encountered in practice and able to produce realistic and informative capture–recapture cases. First we defined 4 scenarios that set survival and detection probabilities (Fig. S 1). In scenario 1, we considered species with high survival rate in adults and juveniles (long-lived) and high detection. For detection probability, we distinguished in the simulation framework below: (1) resighting, a frequently used passive detection method that does not necessitate to capture individuals after first catch (e.g. the observation of colour rings by distance or automatic reading of PIT-tags through fixed antennas); (2) physical captures and recaptures. In scenario 1, both were set high. In scenario 2, survival rate stay the same (long-lived species) but we considered the case when detection is low (both resigthing and capture probabilities). In scenario 3, both survival and and detection were set low, which mimicked cases when the study species is short-lived and re-encounter is rare. Finally in scenario 4, we considered the case when detection is high but survival rate is low, corresponding to studies with short-lived species but designed for high re-encounter probabilities.

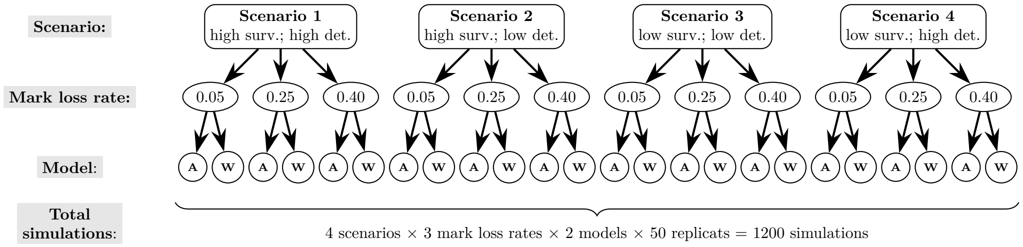

Figure S1: **Graphical representation of the simulation design.** Four biologically relevant scenarios were defined with different level of survival (surv.) and detection (det.) probabilities. Three different mark loss rates were also defined for all scenarios (0.05, 0.25, 0.4). For each of the 12 resulting cases we estimated parameters from 2 different models: one accounting for mark loss and recycling (A) and another one without (W). 50 simulated datasets were then ran for each case, leading to a total of 1200 runs.

For each scenario we set three different values of mark loss rate (Fig. S 1). In the literature (Supporting Information 1) the estimated mark loss rates varied widely between studies and finally stand in the range [0,1]. Since it was not realistic to simulate all possible cases, we decided to restrict our simulations to a range of common values between low mark loss rate (5%), medium (25%) and that we considered as high (40%), higher rates are reported but the value of such a study must be questioned in light of the study's objectives. We restrict our simulation to cases when mark loss is time dependent and can be higher for the first time interval between marking and the following occasion, than subsequently (Fig. 2). Other situations exist, with for example marks that wear out over time and become illegible, in this case the loss of mark increases with the time since marking (Conn et al., 2004; Diefenbach & Alt, 1998).

For each of these 4×3=12 cases we fitted 2 models, one accounting for mark loss and recycling of individuals and another without (Fig. S1). As we used stochastic values for the simulation of survival and capture probabilities, we fit 50 simulations per model in order to account for variability in data collection. Finally, 1200 simulations were ran (Fig. S1).

#### 1.2 Structure of the simulations

In our simulations, females were only allowed to change state among states A, B or C, but not to transit to state D, which resulted in the following state transition matrix:

$$
\begin{array}{c c}
 & \begin{array}{ccccc} \text{state A} & \text{state B} & \text{state C} & \text{state D} & \text{dead} \end{array} \\
\begin{array}{c} \text{state A} \\ \text{state B} \\ \text{state C} \\ \text{state D} \\ \text{dead} \end{array} &
\begin{bmatrix}
\phi_{A,t}(1-\psi_{AB}-\psi_{AC}) & \phi_{A,t}\psi_{AB} & \phi_{A,t}\psi_{AC} & 0 & 1-\phi_{A,t} \\
\phi_{B,t}\psi_{BA} & \phi_{B,t}(1-\psi_{BA}-\psi_{BC}) & \phi_{B,t}\psi_{BC} & 0 & 1-\phi_{B,t} \\
\phi_{C,t}\psi_{CA} & \phi_{C,t}\psi_{CB} & \phi_{C,t}(1-\psi_{CA}-\psi_{CB}) & 0 & 1-\phi_{C,t} \\
0 & 0 & 0 & 0 & 0 \\
0 & 0 & 0 & 0 & 1
\end{bmatrix}
\end{array}
$$

Juvenile males could only stay in their original state or change permanently to state D when juveniles, which resulted in the state transition matrix:

$$
\begin{array}{c|ccccc}
 & \text{state A} & \text{state B} & \text{state C} & \text{state D} & \text{dead} \\
\hline
\text{state A} & \phi_{A,t}(1-\psi_{AD}) & 0 & 0 & \phi_{A,t}\psi_{AD} & 1-\phi_{A,t} \\
\text{state B} & 0 & \phi_{B,t}(1-\psi_{BD}) & 0 & \phi_{B,t}\psi_{BD} & 1-\phi_{B,t} \\
\text{state C} & 0 & 0 & \phi_{C,t}(1-\psi_{CD}) & \phi_{C,t}\psi_{AD} & 1-\phi_{C,t} \\
\text{state D} & 0 & 0 & 0 & 0 & 0 \\
\text{dead} & 0 & 0 & 0 & 0 & 1
\end{array}
$$

The state transition probabilities $\psi$ depended on transition direction (from-to) but were constant in time (see Fig.3 in main text). Adult males could only remain in the same state throughout their lives.

The mark loss process was dependent on time since marking and age of the individuals at marking, with 2 discrete classes for time since marking: (1) the first time between marking occasion and the next occasion; (2) the time since the occasion following marking to the individual death or mark shedding. For (1), we simulated 3 different values inspired from literature (5%, 25% and 40%), but we set (2) to 5% in all simulated scenarios (Fig. S2).

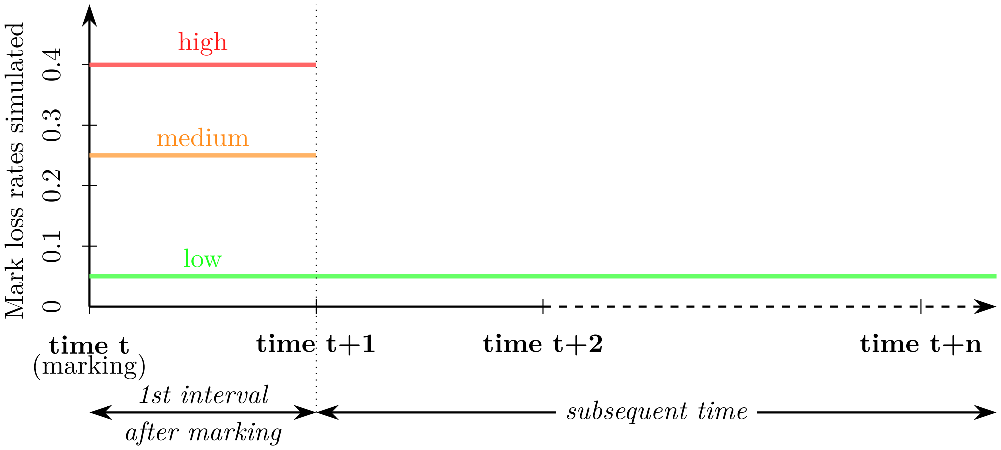

Figure S2: **Mark loss design.** We simulated different mark loss rates during the first time interval (between marking and the next capture occasion) and set it low (5%) after.

The observation matrix link the true states to the observed states:

$$
\begin{array}{c|ccccc}
 & \text{seen in A} & \text{seen in B} & \text{seen in C} & \text{seen in D} & \text{Not seen} \\
\hline
\text{state A} & pd_A & 0 & 0 & 0 & 1-pd_A \\
\text{state B} & 0 & pd_B & 0 & 0 & 1-pd_B \\
\text{state C} & 0 & 0 & pd_C & 0 & 1-pd_C \\
\text{state D} & 0 & 0 & 0 & pb & 1-pb \\
\text{dead} & 0 & 0 & 0 & 0 & 1
\end{array}
$$

with $pd$ the capture probability depending on state ("A","B", or "C"), and $pb$ the resighting probability in D. The observation process for state "A","B", and "C" was in fact more complexe, includind $pc$, the capture probability, and $p.r$, the resighting probability, both state-dependant (see main text).

#### 1.3 Data simulation functions

We simulated data using function inspired by R codes from Kéry and Schaub (2012) book (chapter 7 and 9). We defined two functions to simulate matrix for survival and transition (A), capture-history (CH), an indicator matrix for accasion of marking (TM) and mark retention-history (TR). The first function (simul.msj) simulated data for individual caught when they were juveniles (in their first year), while the second (simul.msa) simulated data for the individuals caught as adults:

```
simul.msj <- function(sex, Pr, Pc, Pout, PSI, PSI_out, mean.phij, mean.phia, pl,
                      marked, n.occasions, state, pz=NULL){
  A <- matrix(NA, ncol = n.occasions, nrow = sum(marked)) # real state
  CH <- matrix(NA, ncol = n.occasions, nrow = sum(marked)) # detection history
  TR <- matrix(NA, ncol = n.occasions, nrow = sum(marked)) # mark retention history
  TM <- matrix(NA, ncol = n.occasions, nrow = sum(marked)) # marking occasions
  PSI1 <- PSI2 <- numeric()
  # Define a vector with the occasion of marking
  mark.occ <- rep(1:length(marked), marked[1:length(marked)])
  # Fill the matrix
  for (i in 1:sum(marked)){
    A[i, mark.occ[i]] <- state[i] # Write the initial state at the release occasion
    TR[i, mark.occ[i]] <- 0 # Write an 0 at first marking occasion
    TM[i, mark.occ[i]] <- 1 # Write an 1 at first marking occasion
    CH[i, mark.occ[i]] <- 1 # Write an 1 at the release occasion
    for (t in (mark.occ[i]+1):n.occasions){
      if(t==(mark.occ[i]+1) & sex[i]==2){
        # Bernoulli trial: does juvenile males survive occasion and change state?
        PSI1 <- rbinom(1,1,mean.phij[A[i,(t-1)],t-1])*PSI_out
        statej <- which(rmultinom(1, 1, c(PSI1,1-sum(PSI1)))==1)
        if (statej==3) break     # If dead, move to the next individual
        A[i,t] <- ifelse(state==1,4,state[i]) # if state =1 emigrate (state= 4),
                                                # if state = 2 stay in subpopulation of birth
        # Bernoulli trial: is mark retain?
        TR[i,t] <- rbinom(1, 1, 1-pl[1])
        # Bernoulli trial: is individual recapture?
        if(A[i,t]==4) pz <- c(0,0,0,Pout,1-Pout) else
          pz <- c(Pc[A[i,t],t-1]*(1-Pr[A[i,t]]),
                  TR[i,t]*Pr[A[i,t]]*(1-Pc[A[i,t],t-1]),
                  Pc[A[i,t],t-1]*Pr[A[i,t]],
                  0,
                  (1-Pc[A[i,t],t-1])*(1-TR[i,t]*Pr[A[i,t]]))
        CH[i,t] <- which(rmultinom(1, 1, pz)==1)
        # if the individual is caught without mark he is marked again
        if(CH[i,t] %in% c(1,3) & TR[i,t]==0) TM[i,t] <- 1 else TM[i,t] <- 0
      }
      if(t>(mark.occ[i]+1) & sex[i]==2){
        # Bernoulli trial: does adult males survive occasion and change state?
        PSI2 <- rbinom(1,1,mean.phia[A[i,(t-1)],t-1])
        state <- which(rmultinom(1, 1, c(PSI2,1-PSI2))==1)
        if (state==2) break # If dead, move to the next individual
        A[i,t] <- A[i,(t-1)]
        # Bernoulli trial: is mark retain?
        tr <- ifelse(TM[i,t-1]==1, rbinom(1, 1, 1-pl[1]),
                     rbinom(1, 1, 1-pl[2]))
        if(TM[i,t-1]==0 & TR[i,t-1]==0) TR[i,t] <- 0 else TR[i,t] <- tr
```

```
        # Bernoulli trial: is individual recapture?
        if(A[i,t]==4) pz <- c(0,0,0,Pout,1-Pout) else
          pz <- c(Pc[A[i,t],t-1]*(1-Pr[A[i,t]]),
                  TR[i,t]*Pr[A[i,t]]*(1-Pc[A[i,t],t-1]),
                  Pc[A[i,t],t-1]*Pr[A[i,t]],
                  0,
                  (1-Pc[A[i,t],t-1])*(1-TR[i,t]*Pr[A[i,t]]))
        CH[i,t] <- which(rmultinom(1, 1, pz)==1)
        # if the individual is caught without mark he is marked again
        if(CH[i,t] %in% c(1,3) & TR[i,t]==0) TM[i,t] <- 1 else TM[i,t] <- 0
      }
      if(t==(mark.occ[i]+1) & sex[i]==1) {
        # Bernoulli trial: does juvenile females survive occasion and change state?
        PSI1 <- rbinom(1,1,mean.phij[A[i,(t-1)],t-1])*PSI[A[i,(t-1)],]
        state <- which(rmultinom(1, 1, c(PSI1,1-sum(PSI1)))==1)
        if (state==4) break     # If dead, move to the next individual
        A[i,t] <- state
        # Bernoulli trial: is mark retain?
        TR[i,t] <- rbinom(1, 1, 1-Pl[1])
        # Bernoulli trial: is individual recapture?
        pz <- c(Pc[A[i,t],t-1]*(1-Pr[A[i,t]]),
                TR[i,t]*Pr[A[i,t]]*(1-Pc[A[i,t],t-1]),
                Pc[A[i,t],t-1]*Pr[A[i,t]],
                0,
                (1-Pc[A[i,t],t-1])*(1-TR[i,t]*Pr[A[i,t]]))
        CH[i,t] <- which(rmultinom(1, 1, pz)==1)
        # if the individual is caught without mark he is marked again
        if(CH[i,t] %in% c(1,3) & TR[i,t]==0) TM[i,t] <- 1 else TM[i,t] <- 0
      }
      if(t>(mark.occ[i]+1) & sex[i]==1) {
        # Bernoulli trial: does adult females survive occasion and change state?
        PSI2 <- rbinom(1,1,mean.phia[A[i,(t-1)],t-1])*PSI[A[i,(t-1)],]
        state <- which(rmultinom(1, 1, c(PSI2,1-sum(PSI2)))==1)
        if (state==4) break     # If dead, move to the next individual
        A[i,t] <- state
        # Bernoulli trial: is mark retain?
        tr <- ifelse(TM[i,t-1]==1, rbinom(1, 1, 1-Pl[1]),
                     rbinom(1, 1, 1-Pl[2]))
        if(TM[i,t-1]==0 & TR[i,t-1]==0) TR[i,t] <- 0 else TR[i,t] <- tr
        # Bernoulli trial: is individual recapture?
        pz <- c(Pc[A[i,t],t-1]*(1-Pr[A[i,t]]),
                TR[i,t]*Pr[A[i,t]]*(1-Pc[A[i,t],t-1]),
                Pc[A[i,t],t-1]*Pr[A[i,t]],
                0,
                (1-Pc[A[i,t],t-1])*(1-TR[i,t]*Pr[A[i,t]]))
        CH[i,t] <- which(rmultinom(1, 1, pz)==1)
        # if the individual is caught without mark he is marked again
        if(CH[i,t] %in% c(1,3) & TR[i,t]==0) TM[i,t] <- 1 else TM[i,t] <- 0
      }
    }
  }
  return(list(A = A, CH = CH, TR = TR, TM = TM))
}
```

```
simul.msa <- function(sex, Pr, Pc, Pout, PSI, mean.phia, Pl, marked, n.occasions,
                      state, pz=NULL){
  A <- matrix(NA, ncol = n.occasions, nrow = sum(marked)) # real state
  CH <- matrix(NA, ncol = n.occasions, nrow = sum(marked)) # detection history
  TR <- matrix(NA, ncol = n.occasions, nrow = sum(marked)) # mark retention history
  TM <- matrix(NA, ncol = n.occasions, nrow = sum(marked)) # marking occasions
  PSI2 <- numeric()
  # Define a vector with the occasion of marking
  mark.occ <- rep(1:length(marked), marked[1:length(marked)])
  # Fill the matrix
  for (i in 1:sum(marked)){
    A[i, mark.occ[i]] <- state[i] # Write initial state at the release occasion
    CH[i, mark.occ[i]] <- 1 # Write an 1 at the release occasion
    TR[i, mark.occ[i]] <- 0 # Write an 0 at first marking occasion
    TM[i, mark.occ[i]] <- 1 # Write an 1 at first marking occasion
    for (t in (mark.occ[i]+1):n.occasions){
      # binomial trial: does adult male  survive occasion?
      surv <- rbinom(1,1,mean.phia[A[i,(t-1)],t-1])
      if(surv==0) break # If dead, move to next individual
      A[i,t] <- ifelse(sex[i]==1, which(rmultinom(1, 1, PSI[A[i,(t-1)],])==1), A[i,(t-1)])
      # Bernoulli trial: is mark retain?
      tr <- ifelse(TM[i,t-1]==1, rbinom(1, 1, 1-Pl[1]),
                   rbinom(1, 1, 1-Pl[2]))
      if(TM[i,t-1]==0 & TR[i,t-1]==0) TR[i,t] <- 0 else TR[i,t] <- tr
      # Bernoulli trial: is individual recapture?
      pz <- c(Pc[A[i,t],t-1]*(1-Pr[A[i,t]]),
              TR[i,t]*Pr[A[i,t]]*(1-Pc[A[i,t],t-1]),
              Pc[A[i,t],t-1]*Pr[A[i,t]],
              0,
              (1-Pc[A[i,t],t-1])*(1-TR[i,t]*Pr[A[i,t]]))
      CH[i,t] <- which(rmultinom(1, 1, pz)==1)
      # if the individual is caught without mark he is marked again
      if(CH[i,t] %in% c(1,3) & TR[i,t]==0) TM[i,t] <- 1 else TM[i,t] <- 0
    }
  }
  return(list(A = A, CH = CH, TR = TR, TM = TM))
}
```

#### 1.4 Specifying real parameter values for simulation

Specific dataset were defined for each subpopulation (A, B and C), as survival and detection probability for males in D. We use set.seed function (random number generator) with different values for each of the 50 replicated datasets we simulated for each combination of parameters, to ensure that generated values always differ between datasets. Here we present an example for simulating the long-lived species with high detection scenario, with a mark loss rate of 0.4.

```
### Data ####
n.occasions <- 10  # Number of capture occasions
pout <- 0.7 # detection probability in D
### Subpopulation A ####
marked.j_A <- rep(40, n.occasions-1) # Annual number of newly marked juveniles
```

```
marked.a_A <- c(60,rep(5, n.occasions-2)) # Annual number of newly marked adults
sex.J_A <- rep(rep(1:2,length.out=40), n.occasions-1) # sex of juveniles
sex.A_A <- c(rep(1,50),rep(2,10),rep(c(2,rep(1,4)), n.occasions-2)) # sex of adults
set.seed(1)
mean.phij_A <- plogis(rnorm(9, 0.2, 0.3)) # Juvenile annual survival
set.seed(1)
mean.phia_A <- plogis(rnorm(9, 2.5, 0.3))  # Adult annual survival
set.seed(1)
pcA <- runif(9, 0.6, 0.7)  # Recapture
prA <- 0.85 #  Resighting
psiAout <- 0.1 # transition from state A to D for male juveniles
PSI_outA <- c(psiAout,1-psiAout) # Define cell transition probability of juvenile males
psiAB <- 0.05 # transition A -> B
psiAC <- 0 # transition A -> C
PSI_A <- c(1-psiAB-psiAC, psiAB, psiAC) # Define cell transition probability for female

### Subpopulation B ####
marked.j_B <- rep(40, n.occasions-1) # Annual number of newly marked juveniles
marked.a_B <- c(60,rep(5, n.occasions-2))  # Annual number of newly marked adults
sex.J_B <- rep(rep(1:2,length.out=40), n.occasions-1) # sex of juvenals
sex.A_B <- c(rep(1,50),rep(2,10),rep(c(2,rep(1,4)), n.occasions-2)) # sex of adults
set.seed(12)
mean.phij_B <- plogis(rnorm(9, 0.2, 0.3)) # Juvenile annual survival
set.seed(12)
mean.phia_B <- plogis(rnorm(9, 2.5, 0.3))  # Adult annual survival
set.seed(12)
pcB <- runif(9, 0.7, 0.8) # Recapture
prB <- 0.95  #  Resighting
psiBout <- 0.5 # transition B -> out for male juveniles
PSI_outB <- c(psiBout,1-psiBout) # Define cell transition probability for juvenile males
psiBA <- 0 # transition B -> A
psiBC <- 0.1 # transition B -> C
PSI_B <- c(psiBA,1-psiBA-psiBC,psiBC) # Define cell transition probability for female

### Subpopulation C ####
marked.j_C <- rep(40, n.occasions-1) # Annual number of newly marked juveniles
marked.a_C <- c(60,rep(5, n.occasions-2))  # Annual number of newly marked adults
sex.J_C <- rep(rep(1:2,length.out=40), n.occasions-1) # sex of juvenals
sex.A_C <- c(rep(1,50),rep(2,10),rep(c(2,rep(1,4)), n.occasions-2)) # sex of adults
set.seed(123)
mean.phij_C <- plogis(rnorm(9, 0.2, 0.3)) # Juvenile annual survival
set.seed(123)
mean.phia_C <- plogis(rnorm(9, 2.5, 0.3))  # Adult annual survival
set.seed(123)
pcC <- runif(9, 0.65, 0.75) # Recapture
prC <- 0.9 #  Resighting
psiCout <- 0.9 # transition C -> out for male juveniles
PSI_outC <- c(psiCout,1-psiCout) # Define cell transition probability for juvenile males
psiCA <- 0.1 # transition C -> A
psiCB <- 0.4 # transition C -> B
PSI_C <- c(psiCA, psiCB, 1-psiCA-psiCB) # Define cell transition probability for female
```

```
# define probability for mark loss : 0.4 1st year and 0.05 the following years
Pl <- c(0.4,0.05)

# survival for males in state D
set.seed(1)
mean.phi_out <- plogis(rnorm(9, 1.5, 0.3))
```

##### 1.5 Simulating data

```
###  Subpopulation A
set.seed(12)
sm <- simul.msj(sex = sex.J_A,
                Pr=c(prA,prB,prC,0),
                Pc=matrix(c(pcA,pcB,pcC,rep(0,(n.occasions-1))),byrow=T,nrow=4),
                Pout=pout,
                PSI=matrix(c(PSI_A,PSI_B,PSI_C),nrow=3,byrow = T),
                PSI_out=PSI_outA,
                mean.phij = matrix(c(mean.phij_A,mean.phij_B,mean.phij_C),byrow=T,nrow=3),
                mean.phia = matrix(c(mean.phia_A,mean.phia_B,mean.phia_C,mean.phi_out),
                                   byrow=T,nrow=4),
                Pl = Pl,
                marked = marked.j_A,
                n.occasions=n.occasions,
                state = rep(1,sum(marked.j_A)))

A_A.J <- sm$A
TM_A.J <- sm$TM
TR_A.J <- sm$TR
z_A.J <- sm$CH

set.seed(12)
sm <- simul.msa(sex = sex.A_A,
                Pr=c(prA,prB,prC),
                Pc=matrix(c(pcA,pcB,pcC,rep(0,(n.occasions-1))),byrow=T,nrow=4),
                Pout=pout,
                PSI=matrix(c(PSI_A,PSI_B,PSI_C),nrow=3,byrow = T),
                mean.phia = matrix(c(mean.phia_A,mean.phia_B,mean.phia_C),byrow=T,nrow=3),
                Pl = Pl,
                marked = marked.a_A,
                n.occasions=n.occasions,
                state = rep(1,sum(marked.a_A)))

A_A.A <- sm$A
TR_A.A <- sm$TR
TM_A.A <- sm$TM
z_A.A <- sm$CH

A_A <- rbind(A_A.J,A_A.A)
TM_A <- rbind(TM_A.J, TM_A.A)
TR_A <- rbind(TR_A.J, TR_A.A)
z_A <- rbind(z_A.J,z_A.A)
```

```
### Subpopulation B
set.seed(12)
sm <- simul.msj(sex = sex.J_B,
                Pr=c(prA,prB,prC,0),
                Pc=matrix(c(pcA,pcB,pcC,rep(0,(n.occasions-1))),byrow=T,nrow=4),
                Pout=pout,
                PSI=matrix(c(PSI_A,PSI_B,PSI_C),nrow=3,byrow = T),
                PSI_out=PSI_outB,
                mean.phij = matrix(c(mean.phij_A,mean.phij_B,mean.phij_C),byrow=T,nrow=3),
                mean.phia = matrix(c(mean.phia_A,mean.phia_B,mean.phia_C,mean.phi_out),
                                   byrow=T,nrow=4),
                Pl = Pl,
                marked = marked.j_B,
                n.occasions = n.occasions,
                state = rep(2,sum(marked.j_B)))

A_B.J <- sm$A
TR_B.J <- sm$TR
TM_B.J <- sm$TM
z_B.J <- sm$CH

set.seed(12)
sm <- simul.msa(sex = sex.A_B,
                Pr=c(prA,prB,prC),
                Pc=matrix(c(pcA,pcB,pcC,rep(0,(n.occasions-1))),byrow=T,nrow=4),
                Pout=pout,
                PSI=matrix(c(PSI_A,PSI_B,PSI_C),nrow=3,byrow = T),
                mean.phia = matrix(c(mean.phia_A,mean.phia_B,mean.phia_C),byrow=T,nrow=3),
                Pl = Pl,
                marked = marked.a_B,
                n.occasions = n.occasions,
                state = rep(2,sum(marked.a_B)))

A_B.A <- sm$A
TR_B.A <- sm$TR
TM_B.A <- sm$TM
z_B.A <- sm$CH

A_B <- rbind(A_B.J,A_B.A)
TM_B <- rbind(TM_B.J, TM_B.A)
TR_B <- rbind(TR_B.J, TR_B.A)
z_B <- rbind(z_B.J,z_B.A)

### Subpopulation C
set.seed(12)
sm <- simul.msj(sex = sex.J_C,
                Pr=c(prA,prB,prC,0),
                Pc=matrix(c(pcA,pcB,pcC,rep(0,(n.occasions-1))),byrow=T,nrow=4),
                Pout=pout,
                PSI=matrix(c(PSI_A,PSI_B,PSI_C),nrow=3,byrow = T),
                PSI_out=PSI_outC,
                mean.phij = matrix(c(mean.phij_A,mean.phij_B,mean.phij_C),byrow=T,nrow=3),
```

```
                mean.phia = matrix(c(mean.phia_A,mean.phia_B,mean.phia_C,mean.phi_out),
                                   byrow=T,nrow=4),
                Pl = Pl,
                marked = marked.j_C,
                n.occasions=n.occasions,
                state = rep(3,sum(marked.j_C)))

A_C.J <- sm$A
TR_C.J <- sm$TR
TM_C.J <- sm$TM
z_C.J <- sm$CH

set.seed(12)
sm <- simul.msa(sex = sex.A_C,
                Pr=c(prA,prB,prC),
                Pc=matrix(c(pcA,pcB,pcC,rep(0,(n.occasions-1))),byrow=T,nrow=4),
                Pout=pout,
                PSI=matrix(c(PSI_A,PSI_B,PSI_C),nrow=3,byrow = T),
                mean.phia = matrix(c(mean.phia_A,mean.phia_B,mean.phia_C),byrow=T,nrow=3),
                Pl = Pl,
                marked = marked.a_C,
                n.occasions=n.occasions,
                state = rep(3,sum(marked.a_C)))

A_C.A <- sm$A
TR_C.A <- sm$TR
TM_C.A <- sm$TM
z_C.A <- sm$CH

A_C <- rbind(A_C.J,A_C.A)
TM_C <- rbind(TM_C.J, TM_C.A)
TR_C <- rbind(TR_C.J, TR_C.A)
z_C <- rbind(z_C.J,z_C.A)

### Bundle data ####
sex <- c(sex.J_A,sex.A_A,sex.J_B,sex.A_B,sex.J_C,sex.A_C)
state <- rbind(A_A, A_B, A_C)
a <- rbind(A_A, A_B, A_C)
a[a>1] <- 1
z <- rbind(z_A,z_B,z_C)
z[is.na(z)] <- 5
f <- apply(z, 1, function(x) min(which(x<5)))
l <- rep(n.occasions,nrow(z))
a[z==5] <- NA
for(i in 1:nrow(a)){
  a[i,(f[i]:max(which(a[i,]==1)))] <- 1
}
state[z==5] <- NA

# age index
age <- a
for(i in 1:nrow(a)){
```

```
  age[i,((f[i]+1):n.occasions)] <- 2
}
idage <- c(rep(1,sum(marked.j_A)),rep(2,sum(marked.a_A)),rep(1,sum(marked.j_B)),
           rep(2,sum(marked.a_B)),rep(1,sum(marked.j_C)),rep(2,sum(marked.a_C)))
for(i in 1:nrow(a)){
  if(idage[i] == 2) age[i,f[i]] <- 2
}

TM <- rbind(TM_A, TM_B, TM_C)
TM[is.na(TM)] <- 0

TR <- rbind(TR_A, TR_B, TR_C)
TR[z==5] <- NA
```

##### 1.6 Prior distributions for the AS model

The state transition process had two consecutive steps, first survival and second transition. The state of an individual $i$ at the first capture can be "A", "B" or "C" with survival probability equal 1, as he is alive for certain. The probability to change or stay in the same state for the subsequent occasion was a product of survival and transition. Survival from time $t-1$ to time $t$ was modelled as a Bernoulli process:

$$\begin{cases} p(A_{i,t}/A_{i,t-1}) \sim Bernoulli(A_{i,t-1}\phi_{i,t-1}), \\ logit(\phi_{i,t}) = \alpha_{state_{i,t}} + \beta_t + \delta_{age_{i,t}} + \gamma_{state_{i,t},t,age_{i,t}} \end{cases}$$

$$\begin{aligned} &\alpha \sim Normal(0, 0.01)[-10, 10] \\ &\beta \sim Student(0, 0.16, 3) \\ &\delta \sim Student(0, 0.16, 3) \\ &\gamma \sim Student(0, 0.16, 3) \end{aligned}$$

with $\phi_{i,t}$ the probability to survive from time $t-1$ to time $t$ conditioned on been alive at $t-1$ ($A_{i,t-1} = 1$).

The probability of changing state from $t-1$ to $t$ was modelled as a categorical process depending on age and sex classes. We used an uninformative Dirichlet prior for state transition probability:

$$\psi \sim Dirichlet(1, 1, 1, 1)$$

State transitions matrix were the same as whose used for data generation described in section 1.2.

The third process was mark retention, which conditioned resighting probability (see observation matrix, section 1.2), as individual without mark (not yet marked or after mark loss) could not be resighted. The mark retention process was a first order Markovian process, where mark loss probability (whose complement is retention probability) varied according to age and the previous state of the mark. For juveniles we have only one probability $pr_{juv.}$, as they were mark at their first catch. But we distinguished if adults were caught and marked during the previous occasion, we defined then their retention probability $pr_{adlt.,1}$, and if they were marked before the previous occasion, their retention probability was defined $pr_{adlt.,2}$. We used weakly informative priors:

$$\begin{cases} pr_{juv.} \sim Beta(1, 1), \\ pr_{adlt.,1} \sim Beta(1, 1), \\ pr_{adlt.,2} \sim Beta(1, 1), \end{cases}$$

The observation process and the observation matrix were the same as those used for data generation described in section 1.2. Two different observation/detection methods have been distinguished, resighting:

$$p.r \sim Beta(1, 1)$$

and recapture:

$$\begin{cases} logit(pc) = \alpha.c_{state_{i,t}} + \beta.c_t + \gamma.c_{state_{i,t},t} \\ \alpha.c \sim Normal(0, 0.01)[-10, 10] \\ \beta.c \sim Student(0, 0.16, 3) \\ \gamma.c \sim Student(0, 0.16, 3) \end{cases}$$

#### 1.7 Model code

Model accounting for mark loss (no recycling), assuming individuals were recognized even in case of mark loss.

```
sink("ModelA.jags")
cat("
model{

  for(i in 1:nind){
    for(j in (f[i]+1):l[i]){
      ap[i,j] <- a[i,j] + 1 # indicator variable (1=dead, 2=alive)
      # This specifies the distribution [a|sv]
      a[i,j] ~ dbern(sv[i,j-1])
      sv[i,j-1] <- a[i,j-1]*phi[i,j-1]
      # State transition probability
      state[i,j] ~ dcat(psi[state[i,j-1],sex[i],age[i,j-1],1:nstate])
      # Detection process
      z[i,j] ~ dcat(pdet[ap[i,j],i,j-1,1:5])
      # indicator variable (1=marked, 2=not marked)
      tm[i,j-1] <- 2-TM[i,j-1]
      # Mark retention process: 1 = retained, 0 = lost
      TR[i,j] ~ dbern(pret[age[i,j-1],tm[i,j-1],i,j-1])
    }
  }

  for(i in 1:nind){
    # specify presence or absence of a mark in the individual
    m[i,f[i]] <- 1 # indicator variable (1 at first marking)
    for(j in (f[i]+1):l[i]){
      m[i,j] <- TR[i,j] + TM[i,j]  # indicator variable (1=mark present, 0=mark absent)
     }
  }

  for(i in 1:nind){
    for(j in (f[i]+1):l[i]){
      # detection probabilities
      pdet[1,i,j-1,1] <- 0 # dead individual can not be caugth
      pdet[1,i,j-1,2] <- 0 # dead individual can not be resighted
      pdet[1,i,j-1,3] <- 0 # dead individual can not be caugth and resighted
      pdet[1,i,j-1,4] <- 0 # dead individual can not be detected in D
      pdet[1,i,j-1,5] <- 1 # dead individual is always undetected

      pdet[2,i,j-1,1] <- ifelse(state[i,j]==4,0,1)*p.c[state[i,j],j-1]*(1 - m[i,j]*
      p.r[state[i,j]]) # only caugth
      pdet[2,i,j-1,2] <- ifelse(state[i,j]==4,0,1)*m[i,j]*p.r[state[i,j]]*(1 -
```

```
    p.c[state[i,j],j-1]) # only resighted
    pdet[2,i,j-1,3] <- ifelse(state[i,j]==4,0,1)*m[i,j]*p.r[state[i,j]]*
    p.c[state[i,j],j-1] # caught and resighted
    pdet[2,i,j-1,4] <- ifelse(state[i,j]==4, pb, 0) # detection in D
    pdet[2,i,j-1,5] <- ifelse(state[i,j]==4, 1-pb, (1 - p.c[state[i,j],j-1])*(1 -
    m[i,j]*p.r[state[i,j]])) # undetected

    # retention probability (see below for description)
    pret[1,1,i,j-1] <- pr[1]
    pret[1,2,i,j-1] <- pr[2]
    pret[2,1,i,j-1] <- pr[3]
    pret[2,2,i,j-1] <- TR[i,j-1]*pr[4]
    }

  for(j in f[i]:(l[i]-1)){
    # survival probability
    logit(phi[i,j]) <- alpha[state[i,j]]  + beta[j]  + delta[age[i,j]] +
    gamma[state[i,j],j,age[i,j]]
  }
}

for(c in 1:(nstate-1)){
  for(j in 1:(nocc-1)){
    logit(p.c[c,j]) <- alpha.c[c] + beta.c[j] + gamma.c[c,j]
    # mean detection probability in Subpopulation A,B or C
    pd[c,j] <- p.c[c,j] + p.r[c] - p.c[c,j]*p.r[c]
  }
}

# Priors and constraints
for(j in 1:(nocc-1)){
  p.c[4,j] <- 0
  p.d[4,j] <- 0
}

pr[1] ~ dbeta(1,1) # retention for juv. after marking
pr[2] <- 0
pr[3] ~ dbeta(1,1) # retention for adlt. after marking
pr[4] ~ dbeta(1,1) # retention for adlt. subsequent time

for(c in 1:nstate){
  alpha[c] ~ dnorm(0,0.01)T(-10,10)
}

for(c in 1:(nstate-1)){
  alpha.c[c] ~ dnorm(0,0.01)T(-10,10)
}

beta[1] <- 0
beta.c[1] <- 0
delta[1] <- 0
delta[2] ~ dt(0,0.16,3)
```

```
for(j in 2:(nocc-1)){
  beta[j] ~ dt(0,0.16,3)
  beta.c[j] ~ dt(0,0.16,3)
}

for(c in 1:nstate){
  for(j in 1:(nocc-1)){
    for(a in 1:2){
      gamma[c,j,a] ~ dt(0,0.16,3)
    }
  }
}

for(c in 1:(nstate-1)){
  gamma.c[c,1] <- 0
}

for(j in 2:(nocc-1)){
  gamma.c[1,j] <- 0
  for(c in 2:3){
    gamma.c[c,j] ~ dt(0,0.16,3)
  }
}

pb ~ dbeta(1,1)

for(c in 1:(nstate-1)){
  p.r[c] ~ dbeta(1,1)
}

p.r[4] <- 0

# state transition probability
for(c in 1:3){
  for(a in 1:2){
    psi[c,1,a,1:3] ~ ddirch(alpha.psi1[])
    psi[c,1,a,4] <- 0
    psi[4,1,a,c] <- 0
  }
}

psi[4,1,1,4] <- 0
psi[4,1,2,4] <- 0

for(c in 1:3){
  psi[c,2,1,1:4] ~ ddirch(alpha.psi2[])
  psi[4,2,1,c] <- 0
}

psi[4,2,1,4] <- 0

for(c in 1:nstate){
  psi[c,2,2,1:nstate] ~ ddirch(alpha.psi2[])
```

```
  }

  for(c in 1:3){
    alpha.psi1[c] <- 1
  }

  for(c in 1:nstate){
    alpha.psi2[c] <- 1
  }

}
",fill = TRUE)
sink()
```

Multistate CJS model initialisation, parameter saving and fiting:

```
data <- list(a = a, state = state, z = z, f = f, l = l,  nocc = n.occasions, nstate=4,
             nind = length(f), age = age, sex = sex, TR = TR, TM = TM)

# inits
a.ini <- a
a.ini[is.na(a)] <- 1
a.ini[!is.na(a)] <- NA
for(i in 1:nrow(a.ini)){
  if(f[i]>1)   a.ini[i,1:f[i]] <- NA
}

state.ini <- state
for(i in 1:nrow(state.ini)){
  if(any(na.omit(state[i, (f[i]+1):l[i]])==4)) state.ini[i, (f[i]+1):l[i]] <- 4 else
    state.ini[i, (f[i]+1):l[i]] <- state.ini[i,f[i]]
}
state.ini[!is.na(state)] <- NA

tr.ini <- TR+TM
for(i in 1:nrow(tr.ini)){
  for(j in 1:ncol(tr.ini)){
    if(j<(f[i]+1) | j==10) next
    if(is.na(tr.ini[i,j]) & any(!is.na(tr.ini[i,(j+1):10]))) tr.ini[i,j] <-
        tr.ini[i,(j+min(which(!is.na(tr.ini[i,(j+1):10]))))]
  }
}
tr.ini[!is.na(TR)] <- NA

inits <- function(){list(a = a.ini, state = state.ini, TR = tr.ini, alpha = rnorm(4,0,1),
                         alpha.c = rnorm(3,0,1), beta = c(NA,rnorm((n.occasions-2),0,1)),
                         beta.c = c(NA,rnorm((n.occasions-2),0,1)),
                         delta = c(NA,rnorm(1,0,1)), gamma = array(rnorm(4*(n.occasions-
                         1)*2,0,1),dim=c(4,(n.occasions-1),2)), gamma.c = matrix(c(
                         rep(NA,(n.occasions-1)), NA,rnorm((n.occasions-2),0,1),NA,
                         rnorm((n.occasions-2),0,1)),nrow=3,byrow=T),
                         p.r = c(runif(3,0,1),NA), pb = runif(1,0,1),
                         pr = c(runif(1,0,1),NA,runif(2,0,1)))}
```

```r
param <- c("alpha", "alpha.c",  "beta", "beta.c", "gamma", "gamma.c", "delta",
           "p.r", "pb", "pd", "psi", "pr")

# MCMC settings
ni <- 150000
nt <- 20
nb <- 50000
nc <- 4

# Call JAGS from R
library(jagsUI)
tlm <- jags(data, inits, param, "ModelA.jags", n.chains = nc, n.adapt=100,
            n.iter = ni, n.burnin = nb, n.thin = nt, parallel=T, store.data=T)
```

### 2 Data simulation with recycled individuals and modelling

#### 2.1 Creating dataset from previous simulations

Here are the codes to modify previous simulated dataset for creating the corresponding dataset with recycled individuals, i.e in the situation when their is no permanent mark and recognition of individual who have lost their mark is not possible.

```r
# New dataset generating from the previous dataset
# row index of individuals marked >1 time
idwt <- which(apply(TM,1,function(x) length(which(x==1))>1))

znew <- z
# remove history after second marking (state 5, undetected)
for(i in idwt){
  znew[i,(which(TM[i,]==1)[2]):10] <- 5
}
# second marking
idwt2 <- which(apply(TM,1,function(x) length(which(x==1))==2))
# remove individuals marked at last occasion
if(length(which(TM[idwt2,10]==1))==0) idwt2 <- idwt2 else idwt2 <- idwt2[-which(
  TM[idwt2,10]==1)]
znew2 <- z[idwt2,]
# remove history before second marking
for(i in 1:nrow(znew2)){
  znew2[i,1:(which(TM[idwt2[i],]==1)[2]-1)] <- 5
}
# third marking
idwt3 <- which(apply(TM,1,function(x) length(which(x==1))==3))
# remove individuals marked at last occasion
if(length(which(TM[idwt3,10]==1))==0) idwt3 <- idwt3 else idwt3 <-
  idwt3[-which(TM[idwt3,10]==1)]
znew3 <- z[idwt3,]
# remove history before third marking
for(i in 1:nrow(znew3)){
```

```
  if(nrow(znew3)!=0) znew3[i,1:(which(TM[idwt3[i],]==1)[3]-1)] <- 5
}
# fourth marking
idwt4 <- which(apply(TM,1,function(x) length(which(x==1))==4))
# remove individuals marked at last occasion
if(length(which(TM[idwt4,10]==1))==0) idwt4 <- idwt4 else idwt4 <-
  idwt4[-which(TM[idwt4,10]==1)]
znew4 <- z[idwt4,]
# remove history before fourth marking
for(i in 1:nrow(znew4)){
  if(nrow(znew4)!=0) znew4[i,1:(which(TM[idwt4[i],]==1)[4]-1)] <- 5
}
# fifth marking
idwt5 <- which(apply(TM,1,function(x) length(which(x==1))==5))
# remove individuals marked at last occasion
if(length(which(TM[idwt5,10]==1))==0) idwt5 <- idwt5 else idwt5 <-
  idwt5[-which(TM[idwt5,10]==1)]
znew5 <- z[idwt5,]
# remove history before fifth marking
for(i in 1:nrow(znew5)){
  if(nrow(znew5)!=0) znew5[i,1:(which(TM[idwt5[i],]==1)[5]-1)] <- 5
}
# sixth marking
idwt6 <- which(apply(TM,1,function(x) length(which(x==1))==6))
# remove individuals marked at last occasion
if(length(which(TM[idwt6,10]==1))==0) idwt6 <- idwt6 else idwt6 <-
  idwt6[-which(TM[idwt6,10]==1)]
znew6 <- z[idwt6,]
# remove history before sixth marking
for(i in 1:nrow(znew6)){
  if(nrow(znew6)!=0) znew6[i,1:(which(TM[idwt6[i],]==1)[6]-1)] <- 5
}
# seventh marking
idwt7 <- which(apply(TM,1,function(x) length(which(x==1))==7))
# remove individuals marked at last occasion
if(length(which(TM[idwt7,10]==1))==0) idwt7 <- idwt7 else idwt7 <-
  idwt7[-which(TM[idwt7,10]==1)]
znew7 <- matrix(z[idwt7,], nrow = length(idwt7))
# remove history before seventh marking
for(i in 1:nrow(znew7)){
  if(nrow(znew7)!=0) znew7[i,1:(which(TM[idwt7[i],]==1)[7]-1)] <- 5
}

znew <- rbind(znew,znew2,znew3,znew4,znew5,znew6,znew7)

anew <- a
for(i in idwt){
  anew[i,(which(TM[i,]==1)[2]):10] <- NA
}
anew2 <- a[idwt2,]
# remove history before second marking
for(i in 1:nrow(anew2)){
```

```
  if(nrow(anew2)!=0) anew2[i,1:(which(TM[idwt2[i],]==1)[2]-1)] <- NA
}
anew3 <- a[idwt3,]
# remove history before third marking
for(i in 1:nrow(anew3)){
  if(nrow(anew3)!=0) anew3[i,1:(which(TM[idwt3[i],]==1)[3]-1)] <- NA
}
anew4 <- a[idwt4,]
# remove history before fourth marking
for(i in 1:nrow(anew4)){
  if(nrow(anew4)!=0) anew4[i,1:(which(TM[idwt4[i],]==1)[4]-1)] <- NA
}
anew5 <- a[idwt5,]
# remove history before fifth marking
for(i in 1:nrow(anew5)){
  if(nrow(anew5)!=0) anew5[i,1:(which(TM[idwt5[i],]==1)[5]-1)] <- NA
}
anew6 <- matrix(a[idwt6,], nrow = length(idwt6))
# remove history before sixth marking
for(i in 1:nrow(anew6)){
  if(nrow(anew6)!=0) anew6[i,1:(which(TM[idwt6[i],]==1)[6]-1)] <- NA
}
anew7 <- matrix(a[idwt7,], nrow = length(idwt7))
# remove history before seventh marking
for(i in 1:nrow(anew7)){
  if(nrow(anew7)!=0) anew7[i,1:(which(TM[idwt7[i],]==1)[7]-1)] <- NA
}

anew <- rbind(anew,anew2,anew3,anew4,anew5,anew6,anew7)

agenew <- age
agenew2 <- age[idwt2,]
# remove history before second marking
for(i in 1:nrow(agenew2)){
  if(nrow(agenew2)!=0) agenew2[i,1:(which(TM[idwt2[i],]==1)[2]-1)] <- NA
}
agenew3 <- age[idwt3,]
# remove history before third marking
for(i in 1:nrow(agenew3)){
  if(nrow(agenew3)!=0) agenew3[i,1:(which(TM[idwt3[i],]==1)[3]-1)] <- NA
}
agenew4 <- age[idwt4,]
# remove history before fourth marking
for(i in 1:nrow(agenew4)){
  if(nrow(agenew4)!=0) agenew4[i,1:(which(TM[idwt4[i],]==1)[4]-1)] <- NA
}
agenew5 <- age[idwt5,]
# remove history before fifth marking
for(i in 1:nrow(agenew5)){
  if(nrow(agenew5)!=0) agenew5[i,1:(which(TM[idwt5[i],]==1)[5]-1)] <- NA
}
agenew6 <- matrix(age[idwt6,], nrow = length(idwt6))
```

```
# remove history before sixth marking
for(i in 1:nrow(agenew6)){
  if(nrow(agenew6)!=0) agenew6[i,1:(which(TM[idwt6[i],]==1)[6]-1)] <- NA
}
agenew7 <- matrix(age[idwt7,], nrow = length(idwt7))
# remove history before seventh marking
for(i in 1:nrow(agenew7)){
  if(nrow(agenew7)!=0) agenew7[i,1:(which(TM[idwt7[i],]==1)[7]-1)] <- NA
}

agenew <- rbind(agenew,agenew2,agenew3,agenew4,agenew5,agenew6,agenew7)

statenew <- state
for(i in idwt){
  statenew[i,(which(TM[i,]==1)[2]):10] <- NA
}
statenew2 <- state[idwt2,]
# remove history before second marking
for(i in 1:nrow(statenew2)){
  if(nrow(statenew2)!=0) statenew2[i,1:(which(TM[idwt2[i],]==1)[2]-1)] <- NA
}
statenew3 <- state[idwt3,]
# remove history before third marking
for(i in 1:nrow(statenew3)){
  if(nrow(statenew3)!=0) statenew3[i,1:(which(TM[idwt3[i],]==1)[3]-1)] <- NA
}
statenew4 <- state[idwt4,]
# remove history before fourth marking
for(i in 1:nrow(statenew4)){
  if(nrow(statenew4)!=0) statenew4[i,1:(which(TM[idwt4[i],]==1)[4]-1)] <- NA
}
statenew5 <- state[idwt5,]
# remove history before fifth marking
for(i in 1:nrow(statenew5)){
  if(nrow(statenew5)!=0) statenew5[i,1:(which(TM[idwt5[i],]==1)[4]-1)] <- NA
}
statenew6 <- matrix(state[idwt6,], nrow = length(idwt6))
# remove history before sixth marking
for(i in 1:nrow(statenew6)){
  if(nrow(statenew6)!=0) statenew6[i,1:(which(TM[idwt6[i],]==1)[6]-1)] <- NA
}
statenew7 <- matrix(state[idwt7,], nrow = length(idwt7))
# remove history before seventh marking
for(i in 1:nrow(statenew7)){
  if(nrow(statenew7)!=0) statenew7[i,1:(which(TM[idwt7[i],]==1)[7]-1)] <- NA
}

statenew <- rbind(statenew,statenew2,statenew3,statenew4,statenew5,statenew6,statenew7)

fnew <- apply(anew,1,function(x) min(which(x==1)))
lnew <- rep(10,length(fnew))
sexnew <- c(sex,sex[idwt2],sex[idwt3],sex[idwt4],sex[idwt5],sex[idwt6],sex[idwt7])
```

#### 2.2 Model code

Model for estimating parameters when mark loss is not account for (data with recycling), assuming individuals are not recognized if recaptured. This is the same model as the previous one but mark fate tracking ($m$) and mark loss estimation were removed.

```
sink("ModelW.jags")
cat("
model{

  for(i in 1:nind){
    for(j in (f[i]+1):l[i]){
      ap[i,j] <- a[i,j] + 1 # indicator variable (1=dead, 2=alive)
      # This specifies the distribution [a|sv]
      a[i,j] ~ dbern(sv[i,j-1])
      sv[i,j-1] <- a[i,j-1]*phi[i,j-1]
      # State transition process
      state[i,j] ~ dcat(psi[state[i,j-1],sex[i],age[i,j-1],1:nstate])
      # Detection process
      z[i,j] ~ dcat(pdet[ap[i,j],i,j-1,1:5])
    }
  }

  for(i in 1:nind){
    for(j in (f[i]+1):l[i]){
      # detection probabilities
      pdet[1,i,j-1,1] <- 0 # dead individual can not be caugth
      pdet[1,i,j-1,2] <- 0 # dead individual can not be resighted
      pdet[1,i,j-1,3] <- 0 # dead individual can not be caugth and resighted
      pdet[1,i,j-1,4] <- 0 # dead individual can not be detected in D
      pdet[1,i,j-1,5] <- 1 # dead individual is always undetected

      pdet[2,i,j-1,1] <- ifelse(state[i,j]==4,0,1)*p.c[state[i,j],j-1]*(1 -
      p.r[state[i,j]]) # only caught
      pdet[2,i,j-1,2] <- ifelse(state[i,j]==4,0,1)*p.r[state[i,j]]*(1 -
      p.c[state[i,j],j-1]) # only resigted
      pdet[2,i,j-1,3] <- ifelse(state[i,j]==4,0,1)*p.r[state[i,j]]*
      p.c[state[i,j],j-1] # caugth and resighted
      pdet[2,i,j-1,4] <- ifelse(state[i,j]==4, pb, 0)  # detection in D
      pdet[2,i,j-1,5] <- ifelse(state[i,j]==4, 1-pb, (1 - p.c[state[i,j],j-1])*(1 -
      p.r[state[i,j]])) # undetected
      }

    for(j in f[i]:(l[i]-1)){
      # survival probability )
      logit(phi[i,j]) <- alpha[state[i,j]]  + beta[j]  + delta[age[i,j]] +
      gamma[state[i,j],j,age[i,j]]
    }
  }

  for(c in 1:(state[-1))){
    for(j in 1:(nocc-1)){
      logit(p.c[c,j]) <- alpha.c[c] + beta.c[j] + gamma.c[c,j]
      # mean detection probability in Subpopulation A,B,C
```

```
    pd[c,j] <- p.c[c,j] + p.r[c] - p.c[c,j]*p.r[c]
  }
}

# Priors and constraints
for(j in 1:(nocc-1)){
  p.c[4,j] <- 0
  p.d[4,j] <- 0
}

for(c in 1:state[){
  alpha[c] ~ dnorm(0,0.01)T(-10,10)
}

for(c in 1:(state[-1)){
  alpha.c[c] ~ dnorm(0,0.01)T(-10,10)
}

beta[1] <- 0
beta.c[1] <- 0
delta[1] <- 0
delta[2] ~ dt(0,0.16,3)

for(j in 2:(nocc-1)){
  beta[j] ~ dt(0,0.16,3)
  beta.c[j] ~ dt(0,0.16,3)
}

for(c in 1:state[){
  for(j in 1:(nocc-1)){
    for(a in 1:2){
      gamma[c,j,a] ~ dt(0,0.16,3)
    }
  }
}

for(c in 1:(state[-1)){
  gamma.c[c,1] <- 0
}

for(j in 2:(nocc-1)){
  gamma.c[1,j] <- 0
  for(c in 2:3){
    gamma.c[c,j] ~ dt(0,0.16,3)
  }
}

pb ~ dbeta(1,1)

for(c in 1:(state[-1)){
  p.r[c] ~ dbeta(1,1)
}
```

```
  p.r[4] <- 0

  # movement probability
  for(c in 1:3){
    for(a in 1:2){
      psi[c,1,a,1:3] ~ ddirch(alpha.psi1[])
      psi[c,1,a,4] <- 0
      psi[4,1,a,c] <- 0
    }
  }

  psi[4,1,1,4] <- 0
  psi[4,1,2,4] <- 0

  for(c in 1:3){
    psi[c,2,1,1:4] ~ ddirch(alpha.psi2[])
    psi[4,2,1,c] <- 0
  }

  psi[4,2,1,4] <- 0

  for(c in 1:state[]){
    psi[c,2,2,1:state[] ~ ddirch(alpha.psi2[])
  }

  for(c in 1:3){
    alpha.psi1[c] <- 1
  }

  for(c in 1:state[]){
    alpha.psi2[c] <- 1
  }

}
",fill = TRUE)
sink()
```

Multistate CJS model initialisation, parameter saving and fiting:

```
data <- list(a = anew, state = statenew, z = znew, f = fnew, l = lnew,  nocc = n.occasions,
             state[=4, nind = length(fnew), age = agenew, sex = sexnew)

# inits
a.ini <- anew
a.ini[is.na(anew)] <- 1
a.ini[!is.na(anew)] <- NA
for(i in 1:nrow(a.ini)){
  if(fnew[i]>1)   a.ini[i,1:fnew[i]] <- NA
}

state.ini <- statenew
for(i in 1:nrow(state.ini)){
  if(any(na.omit(statenew[i, (fnew[i]+1):lnew[i]])==4)) state.ini[i,
```

```
                                                          (fnew[i]+1):lnew[i]] <- 4 else
    state.ini[i, (fnew[i]+1):lnew[i]] <- state.ini[i,fnew[i]]
}
state.ini[!is.na(statenew)] <- NA

inits <- function(){list(a = a.ini, state = state.ini, alpha = rnorm(4,0,1),
                         alpha.c = rnorm(3,0,1),
                         beta = c(NA,rnorm((n.occasions-2),0,1)), beta.c = c(NA,
                         rnorm((n.occasions-2),0,1)),
                         delta = c(NA,rnorm(1,0,1)), gamma = array(rnorm(4*
                         (n.occasions-1)*2,0,1),dim=c(4,(n.occasions-1),2)),
                         gamma.c = matrix(c(rep(NA,(n.occasions-1)),NA,
                         rnorm((n.occasions-2),0,1),NA,rnorm((n.occasions-2),0,1)),
                         nrow=3,byrow=T),
                         p.r = c(runif(3,0,1),NA), pb = runif(1,0,1),
                         pret = matrix(c(runif(2,0,1),NA,runif(1,0,1)),nrow=2))}

param <- c("alpha", "alpha.c", "beta", "beta.c", "gamma", "gamma.c", "delta",
           "p.r", "pb", "pd", "psi")

# MCMC settings
ni <- 150000
nt <- 20
nb <- 50000
nc <- 4

# Call JAGS from R
library(jagsUI)
tlm <- jags(data, inits, param, "ModelW.jags", n.chains = nc, n.adapt=100,
            n.iter = ni, n.burnin = nb, n.thin = nt, parallel=T, store.data=T)
```

### 3 Results

#### 3.1 Mark loss (TL) and recycling (TR)

The number of individuals recycled can vary significantly depending on many parameters, such as mark loss, survival and recapture rates. Below is the total number of recycling that occurred in our 12 simulated study cases during the 10 occasions defined as the duration of the study. The proportion of recycling in scenario where recapture was high (1, 4) is closer to the number of mark loss compare to those were recapture set low (2,3). The total number of recycling even is higher for scenarios where survival was set high(1,2).

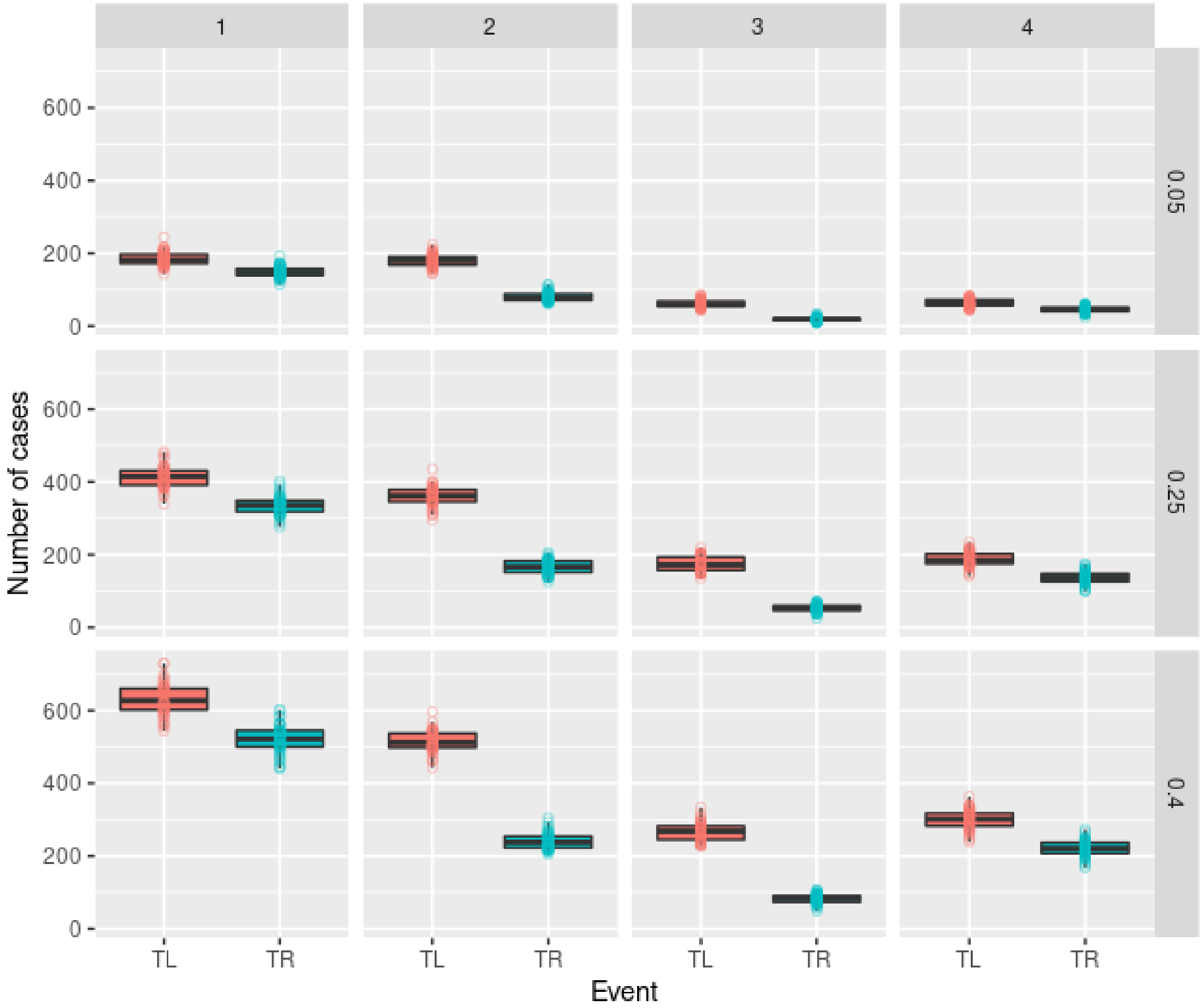

Figure S3: Boxplot of the number of mark loss (TL) and recycling (TR) events for each simulated scenario (1, 2, 3, 4) × mark loss rates (0.05, 0.25, 0.4) across the 50 simulated datasets.

#### 3.2 Parameter convergence

##### 3.2.1 Model that accounted for tag loss

| Scenario | TL prob. | Parameter | Failure | Total | Proportion |
|---|---|---|---|---|---|
| 1 | 0.05 | $\gamma$ | 25 | 3600 | 0.007 |
| 2 | 0.05 | $\gamma$ | 54 | 3600 | 0.015 |
| 3 | 0.05 | $\gamma$ | 70 | 3600 | 0.019 |
| 3 | 0.05 | $\gamma.c$ | 1 | 800 | 0.001 |
| 4 | 0.05 | $\gamma$ | 13 | 3600 | 0.004 |
| 4 | 0.05 | $\gamma.c$ | 2 | 800 | 0.003 |
| 1 | 0.25 | $\gamma$ | 25 | 3600 | 0.007 |
| 2 | 0.25 | $\beta$ | 3 | 400 | 0.008 |
| 2 | 0.25 | $\gamma$ | 75 | 3600 | 0.021 |
| 3 | 0.25 | $\gamma$ | 3 | 3600 | 0.001 |
| 4 | 0.25 | $\gamma$ | 14 | 3600 | 0.004 |
| 4 | 0.25 | $\gamma.c$ | 1 | 800 | 0.001 |
| 1 | 0.4 | $\gamma$ | 25 | 3600 | 0.007 |
| 2 | 0.4 | $\beta$ | 1 | 400 | 0.003 |
| 2 | 0.4 | $\gamma$ | 85 | 3600 | 0.024 |
| 2 | 0.4 | $pb$ | 1 | 50 | 0.020 |
| 3 | 0.4 | $\gamma$ | 46 | 3600 | 0.013 |
| 3 | 0.4 | $\gamma.c$ | 1 | 800 | 0.001 |
| 4 | 0.4 | $\gamma$ | 23 | 3600 | 0.006 |

Table S1: **Parameters that did not converge ($\hat{R} > 1.05$)**.Data are displayed for each, scenario, each tag loss probability (TL prob.) simulated and each parameter. Failure indicate the number of times a parameter failed to converge for all 50 simulations. Total is the total number of values estimated for a particular parameter for all 50 simulations. Proportion is the proportion of failure among all value estimated for a particular parameter for all 50 simulations. $\gamma$ correspond to the combine effect of the state, time and age on survival probability; $\beta$ is the effect of time on survival probability; $p_b$ is the probability of detection in state "D"; $\gamma.c$ is the combine effect of state and time on capture probability. See details of the parameter specifications in section 1.6.

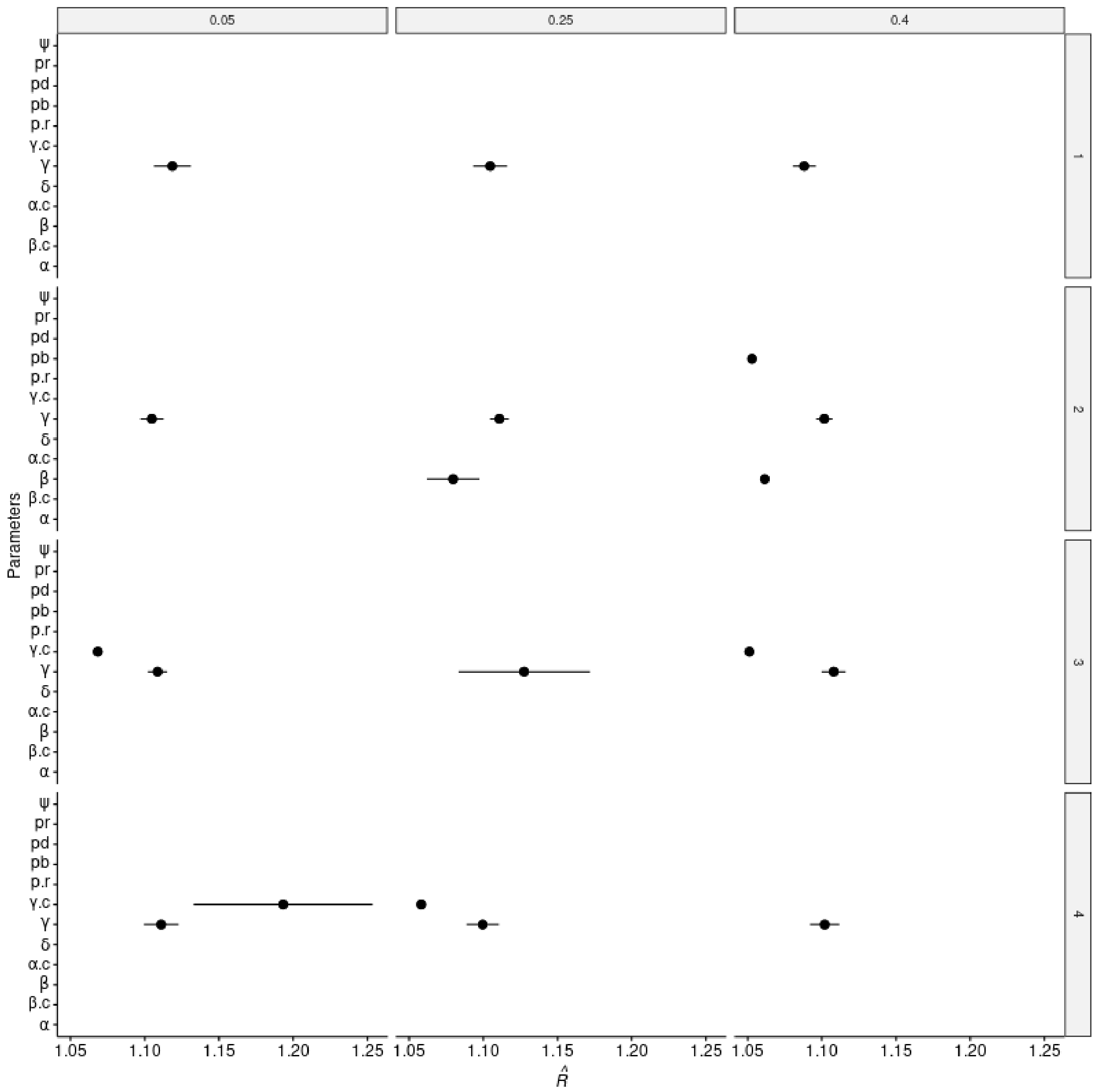

Figure S4: $\hat{R} > 1.05$. Mean and standard error of $\hat{R}$ values that increase 1.05, showing convergence failure of the concerned parameters. Scenarios are indicated on the right side of the plot and tag loss rate simulated on the top. See details of parameter in section 1.6.

##### 3.2.2 Model that did not account for tag loss

| Scenario | TL prob. | Parameter | Failure | Total | Proportion |
|---|---|---|---|---|---|
| 1 | 0.05 | $\gamma$ | 22 | 3600 | 0.006 |
| 2 | 0.05 | $\beta$ | 2 | 400 | 0.005 |
| 2 | 0.05 | $\gamma$ | 76 | 3600 | 0.021 |
| 3 | 0.05 | $\gamma$ | 66 | 3600 | 0.018 |
| 4 | 0.05 | $\gamma$ | 12 | 3600 | 0.003 |
| 1 | 0.25 | $\gamma$ | 19 | 3600 | 0.005 |

| Scenario | TL prob. | Parameter | Failure | Total | Proportion |
|---|---|---|---|---|---|
| 2 | 0.25 | $\alpha$ | 6 | 200 | 0.030 |
| 2 | 0.25 | $\alpha.c$ | 3 | 150 | 0.020 |
| 2 | 0.25 | $\beta$ | 10 | 400 | 0.025 |
| 2 | 0.25 | $\gamma$ | 126 | 3600 | 0.035 |
| 2 | 0.25 | $\gamma.c$ | 4 | 800 | 0.005 |
| 2 | 0.25 | $\delta$ | 2 | 50 | 0.040 |
| 2 | 0.25 | $p.r$ | 5 | 150 | 0.033 |
| 2 | 0.25 | $pb$ | 2 | 50 | 0.040 |
| 2 | 0.25 | $pd$ | 44 | 1350 | 0.033 |
| 2 | 0.25 | $\psi$ | 64 | 2300 | 0.028 |
| 3 | 0.25 | $\alpha$ | 3 | 200 | 0.015 |
| 3 | 0.25 | $\alpha.c$ | 1 | 150 | 0.007 |
| 3 | 0.25 | $\beta$ | 1 | 400 | 0.003 |
| 3 | 0.25 | $\gamma$ | 60 | 3600 | 0.017 |
| 3 | 0.25 | $\gamma.c$ | 1 | 800 | 0.001 |
| 3 | 0.25 | $\delta$ | 1 | 50 | 0.020 |
| 3 | 0.25 | $p.r$ | 2 | 150 | 0.013 |
| 3 | 0.25 | $pd$ | 14 | 1350 | 0.010 |
| 3 | 0.25 | $\psi$ | 23 | 2300 | 0.010 |
| 4 | 0.25 | $\gamma$ | 14 | 3600 | 0.004 |
| 1 | 0.4 | $\gamma$ | 22 | 3600 | 0.006 |
| 2 | 0.4 | $\alpha$ | 21 | 200 | 0.105 |
| 2 | 0.4 | $\alpha.c$ | 8 | 150 | 0.053 |
| 2 | 0.4 | $\beta$ | 22 | 400 | 0.055 |
| 2 | 0.4 | $\beta$.c | 4 | 400 | 0.010 |
| 2 | 0.4 | $\gamma$ | 157 | 3600 | 0.044 |
| 2 | 0.4 | $\gamma.c$ | 14 | 800 | 0.018 |
| 2 | 0.4 | $\delta$ | 3 | 50 | 0.060 |
| 2 | 0.4 | $p.r$ | 18 | 150 | 0.120 |
| 2 | 0.4 | $pb$ | 11 | 50 | 0.220 |
| 2 | 0.4 | $pd$ | 134 | 1350 | 0.099 |
| 2 | 0.4 | $\psi$ | 177 | 2300 | 0.077 |
| 3 | 0.4 | $\alpha$ | 8 | 200 | 0.040 |
| 3 | 0.4 | $\alpha.c$ | 5 | 150 | 0.033 |
| 3 | 0.4 | $\beta$ | 7 | 400 | 0.018 |
| 3 | 0.4 | $\gamma$ | 105 | 3600 | 0.029 |
| 3 | 0.4 | $\gamma.c$ | 5 | 800 | 0.006 |
| 3 | 0.4 | $\delta$ | 1 | 50 | 0.020 |
| 3 | 0.4 | $p.r$ | 14 | 150 | 0.093 |
| 3 | 0.4 | $pb$ | 3 | 50 | 0.060 |
| 3 | 0.4 | $pd$ | 96 | 1350 | 0.071 |
| 3 | 0.4 | $\psi$ | 115 | 2300 | 0.050 |
| 4 | 0.4 | $\gamma$ | 16 | 3600 | 0.004 |

Table S2: **Parameters that did not converge ($\hat{R} > 1.05$)**.Data are displayed for each, scenario, each tag loss probability (TL prob.) simulated and each parameter. Failure indicate the number of times a parameter failed to converge for all 50 simulations. Total is the total number of values estimated for a particular parameter for all 50 simulations. Proportion is the proportion of failure among all value estimated for a particular parameter for all 50 simulations. $\alpha$ is the effect of state on survival probability; $\beta$ is the effect of time on survival probability; $\delta$ is the effect of age on survival probability; $\gamma$ correspond to the combine effect of the state, time and age on survival probability; $\alpha.c$ is the effect of state on detection probability; $\beta.c$ is the effect of time on detection probability; $\gamma.c$ correspond to the combine effect of the state and time on detection probability; $p.r$ is the probability of resighting; $p_b$ is the probability of detection in state "D"; $p_d$ is the probability of detection; $\psi$ is the state transition probability. See details of the parameter specifications in section 1.6.

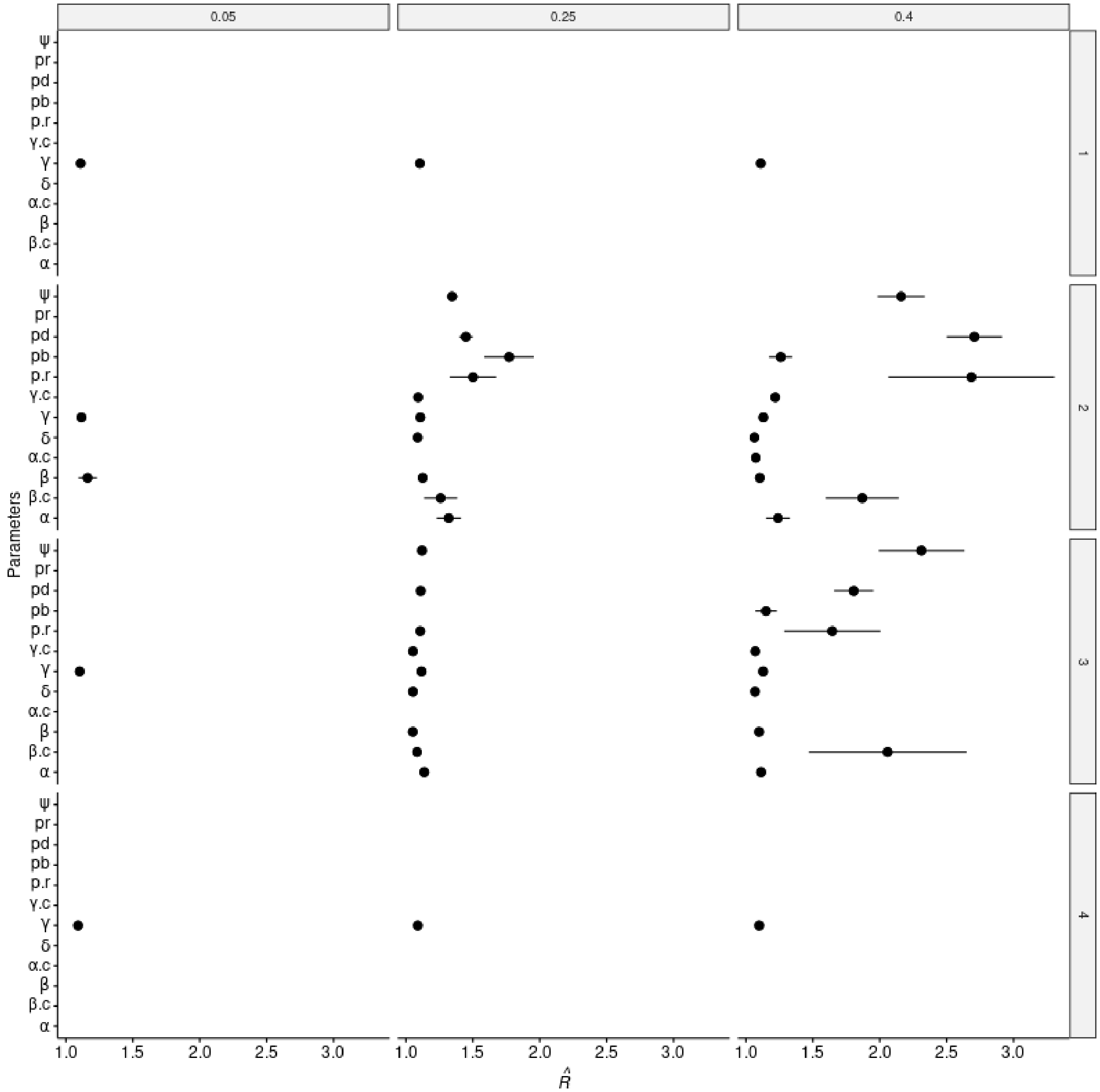

Figure S5: $\hat{R} > 1.05$. Mean and standard error of $\hat{R}$ values that increase 1.05, showing convergence failure of the concerned parameters. Scenarios are indicated on the right side of the plot and tag loss rate simulated on the top. See details of parameter in section 1.6.

#### 3.3 Comparison of survival probability estimates

Below, we compared the bias (median - truth) and the precision (mean squared errors MSE = bias$^2$+ variance) of the survival probability between the model without recycling and with recycling. We also use the Earth Mover Distance (EMD) to compare the distribution of the medians of these parameters. The density distribution of these medians were also displayed. In our simulation framework, only juvenile males could enter the "D" state during their first year of life and became adult as soon as they reached their second year of life if they survived. Survival in state "D" can then only be estimated for adult males.

##### 3.3.1 Simulations with mark loss rate of 0.05

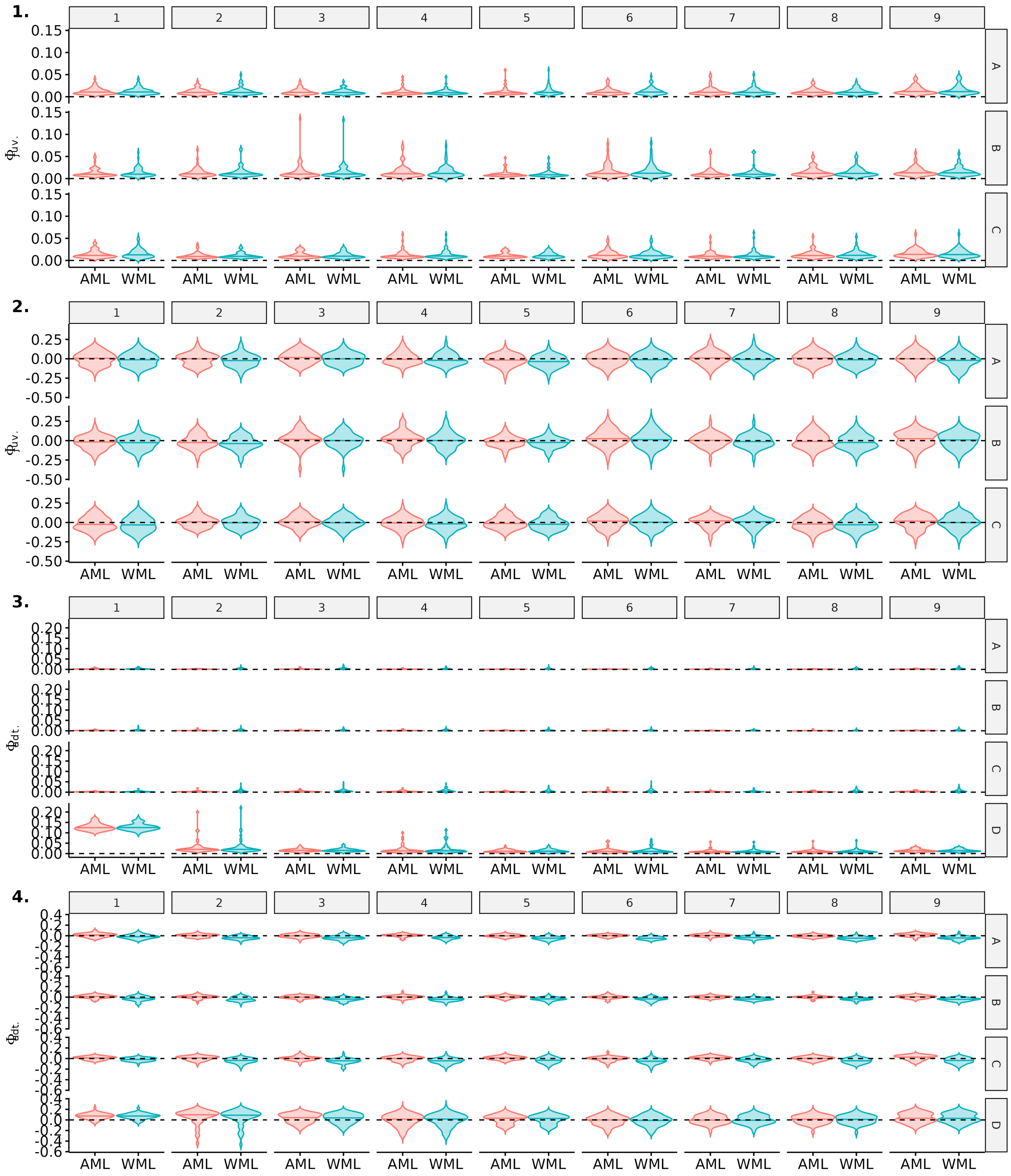

Figure S6: Comparison of precision and bias for estimates of juvenile and adult survival probabilities between model accounting for mark loss (AML) or not (WML), over the 9 recapture occasions. Here, scenario 1 (long-lived species with high detection) is shown for a simulated mark loss probability of 0.05. Violin plots show the distributions of mean precision (1,3) and bias (2,4) over 50 simulations. The median of each distribution is shown with an horizontal line.

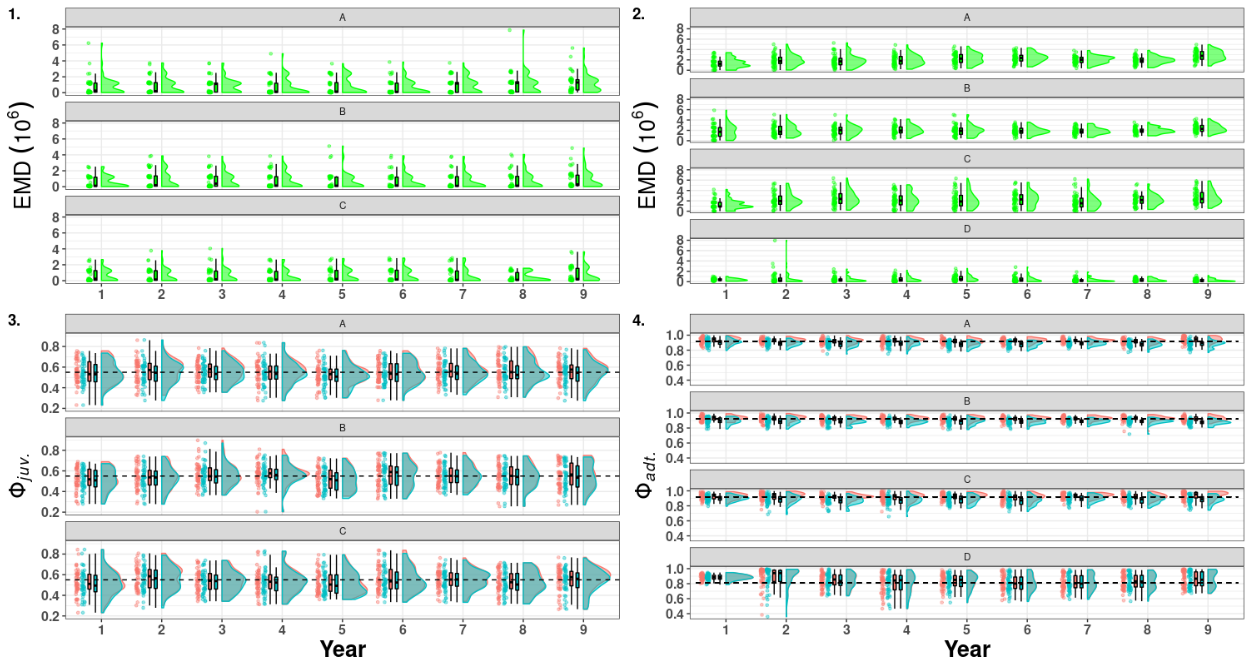

Figure S7: Raincloud plots of the Earth Mover Distance (green) between the medians of the posterior distribution of survival estimated for the model accounting for mark loss and recycling (red) and not accounting for mark loss (blue). On the left juvenile survival ($\phi_{juv.}$, 1 & 3), on the right adult survival ($\phi_{adt.}$, 2 & 4) in scenario 1 (long-lived species with high detection) with a simulated mark loss probability of 0.05. A, B, C, D denotes the name of the states and the dashed line correspond to the mean of the true values.

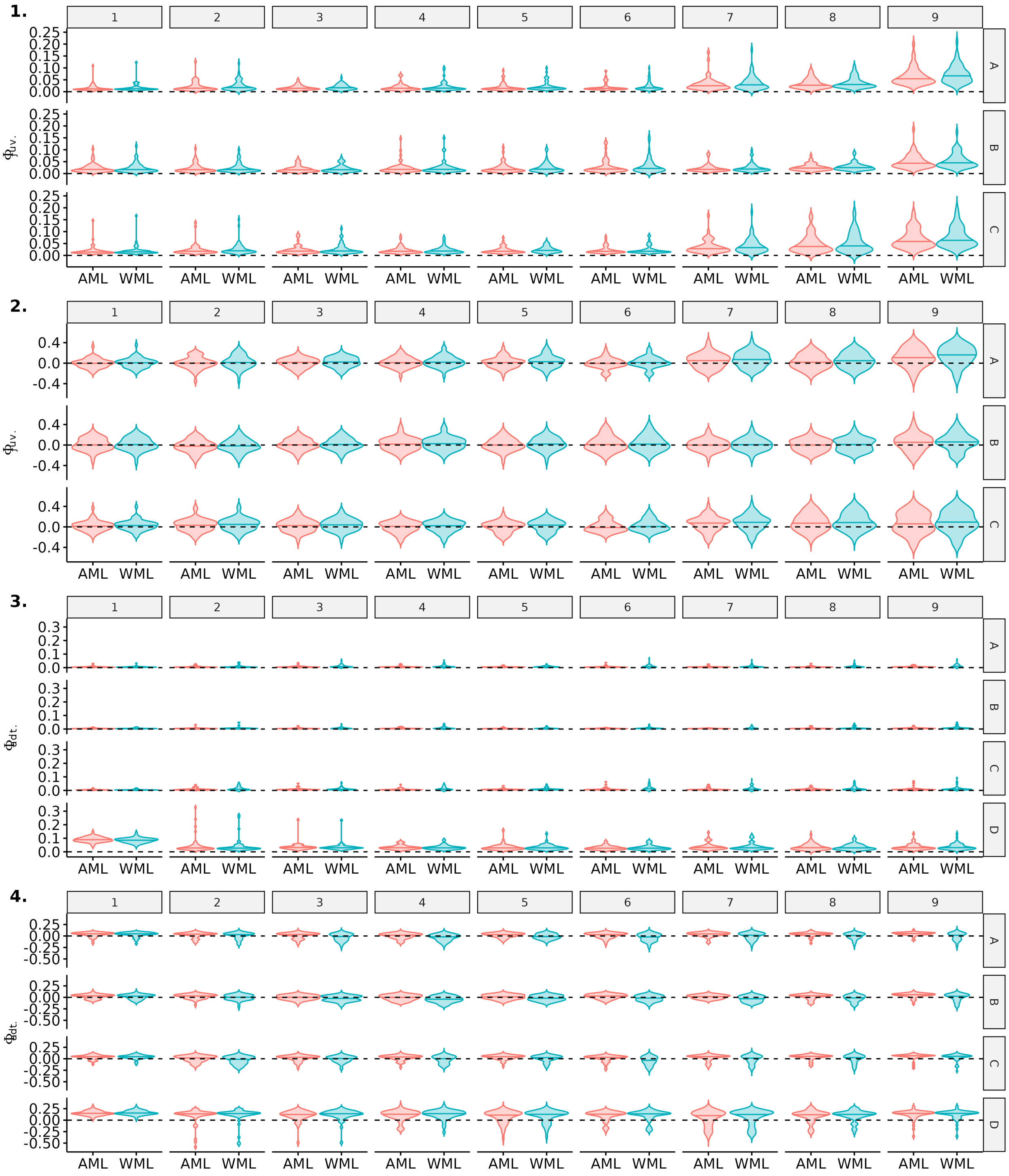

Figure S8: Comparison of precision and bias for estimates of juvenile and adult survival probabilities between model accounting for mark loss (AML) or not (WML), over the 9 recapture occasions. Here, scenario 2 (long-lived species with low detection) is shown for a simulated mark loss probability of 0.05. Violin plots show the distributions of mean precision (1,3) and bias (2,4) over 50 simulations. The median of each distribution is shown with an horizontal line.

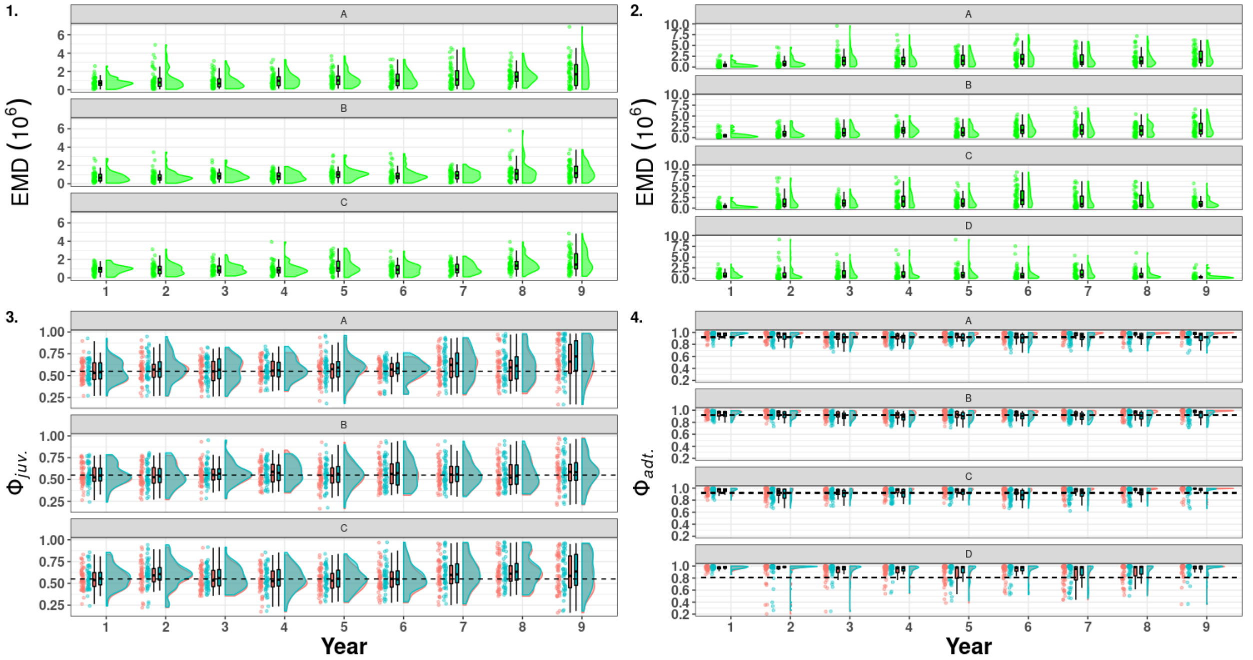

Figure S9: Raincloud plots of the Earth Mover Distance (green) between the medians of the posterior distribution of survival estimated for the model accounting for mark loss and recycling (red) and not accounting for mark loss (blue). On the left juvenile survival ($\phi_{juv.}$, 1 & 3), on the right adult survival ($\phi_{adt.}$, 2 & 4) in scenario 2 (long-lived species with low detection) with a simulated mark loss probability of 0.05. A, B, C, D denotes the name of the states and the dashed line correspond to the mean of the true values.

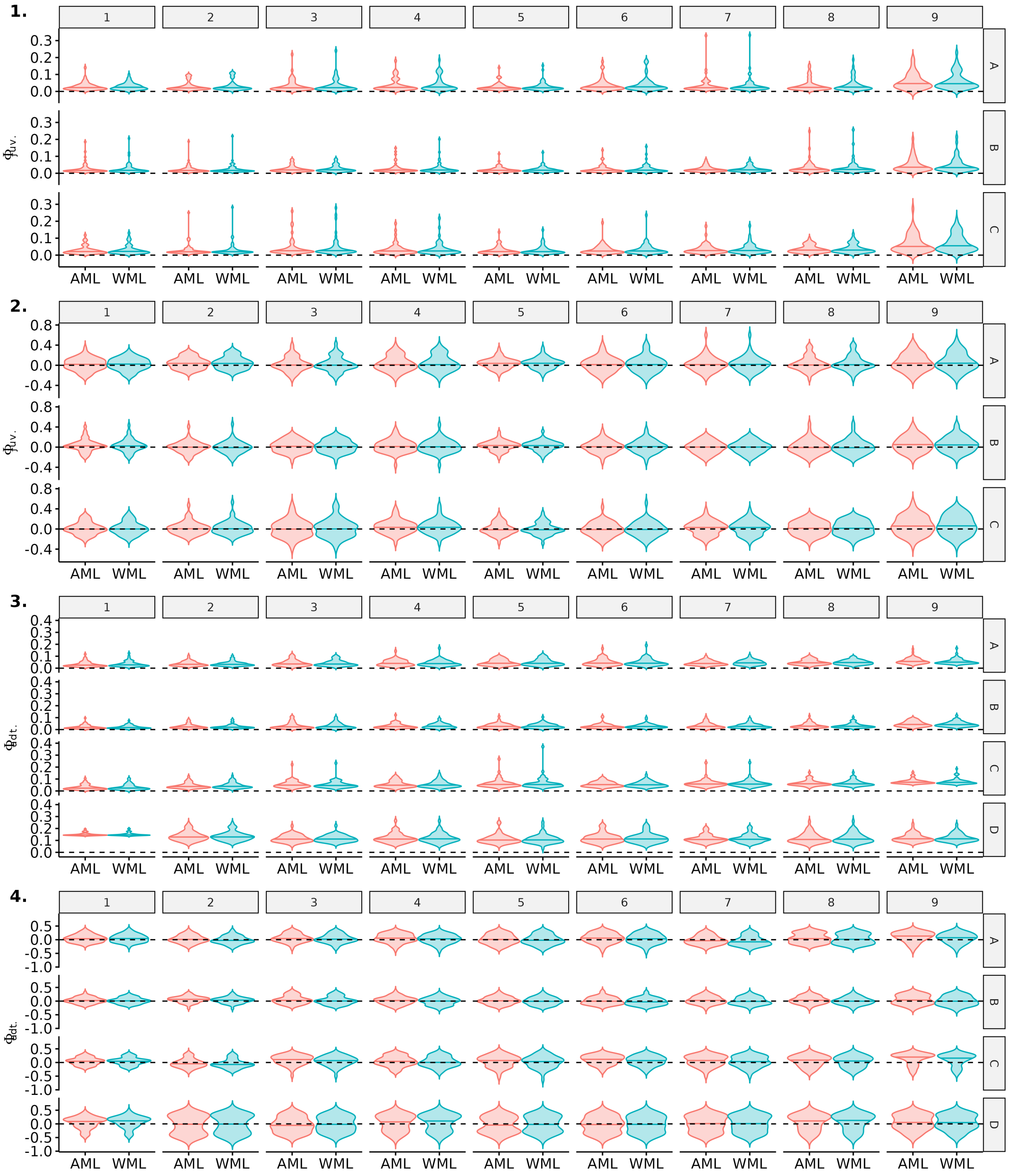

Figure S10: Comparison of precision and bias for estimates of juvenile and adult survival probabilities between model accounting for mark loss (AML) or not (WML), over the 9 recapture occasions. Here, scenario 3 (short-lived species with low detection) is shown for a simulated mark loss probability of 0.05. Violin plots show the distributions of mean precision (1,3) and bias (2,4) over 50 simulations. The median of each distribution is shown with an horizontal line.

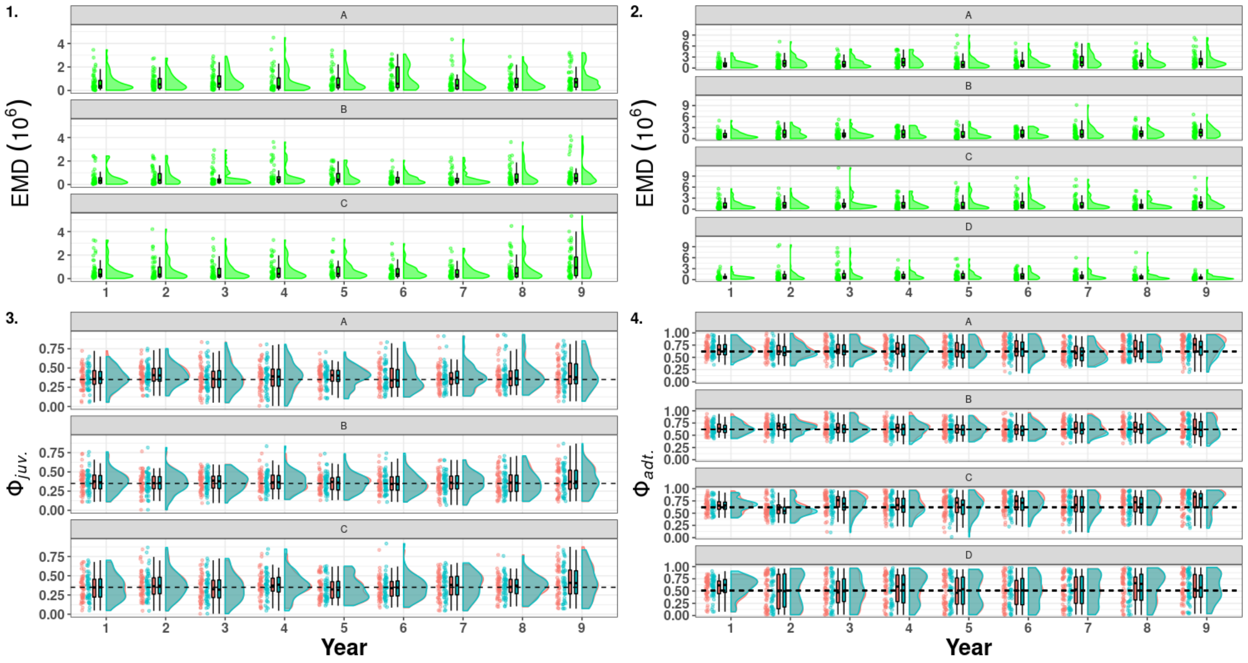

Figure S11: Raincloud plots of the Earth Mover Distance (green) between the medians of the posterior distribution of survival estimated for the model accounting for mark loss and recycling (red) and not accounting for mark loss (blue). On the left juvenile survival ($\phi_{juv.}$, 1 & 3), on the right adult survival ($\phi_{adt.}$, 2 & 4) in scenario 3 (short-lived species with low detection) with a simulated mark loss probability of 0.05. A, B, C, D denotes the name of the states and the dashed line correspond to the mean of the true values.

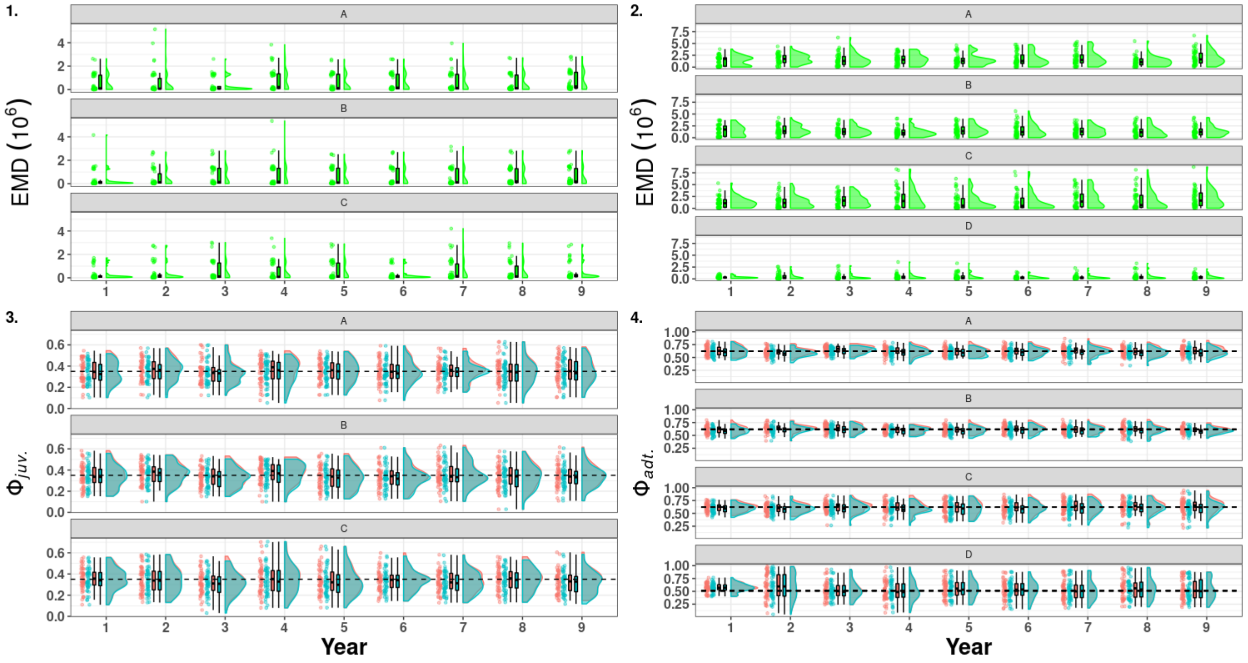

Figure S12: Raincloud plots of the Earth Mover Distance (green) between the medians of the posterior distribution of survival estimated for the model accounting for mark loss and recycling (red) and not accounting for mark loss (blue). On the left juvenile survival ($\phi_{juv.}$, 1 & 3), on the right adult survival ($\phi_{adt.}$, 2 & 4) in scenario 4 (short-lived species with high detection) with a simulated mark loss probability of 0.05. A, B, C, D denotes the name of the states and the dashed line correspond to the mean of the true values.

##### 3.3.2 Simulations with mark loss rate of 0.25

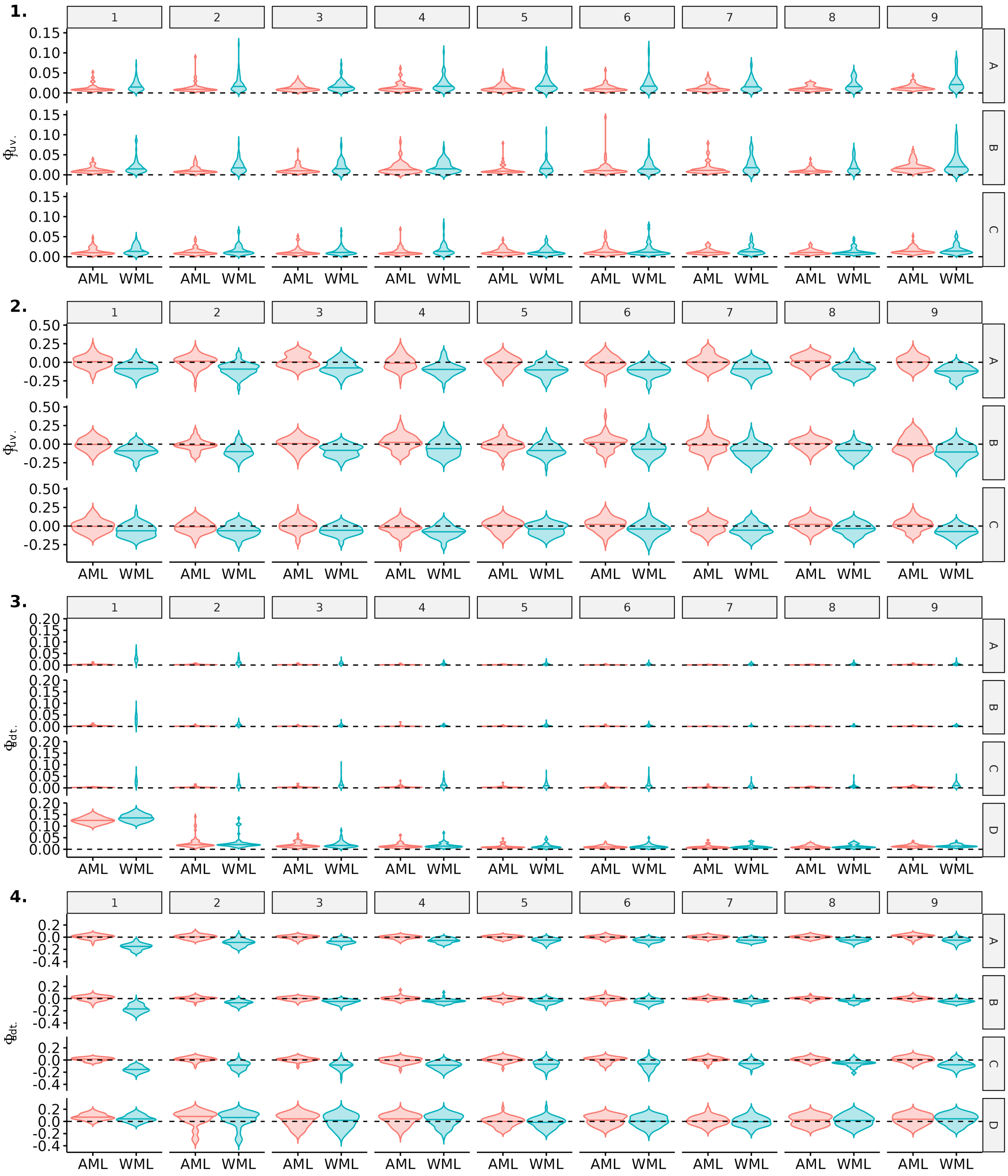

Figure S13: Comparison of precision and bias for estimates of juvenile and adult survival probabilities between model accounting for mark loss (AML) or not (WML), over the 9 recapture occasions. Here, scenario 1 (long-lived species with high detection) is shown for a simulated mark loss probability of 0.25. Violin plots show the distributions of mean precision (1,3) and bias (2,4) over 50 simulations. The median of each distribution is shown with an horizontal line.

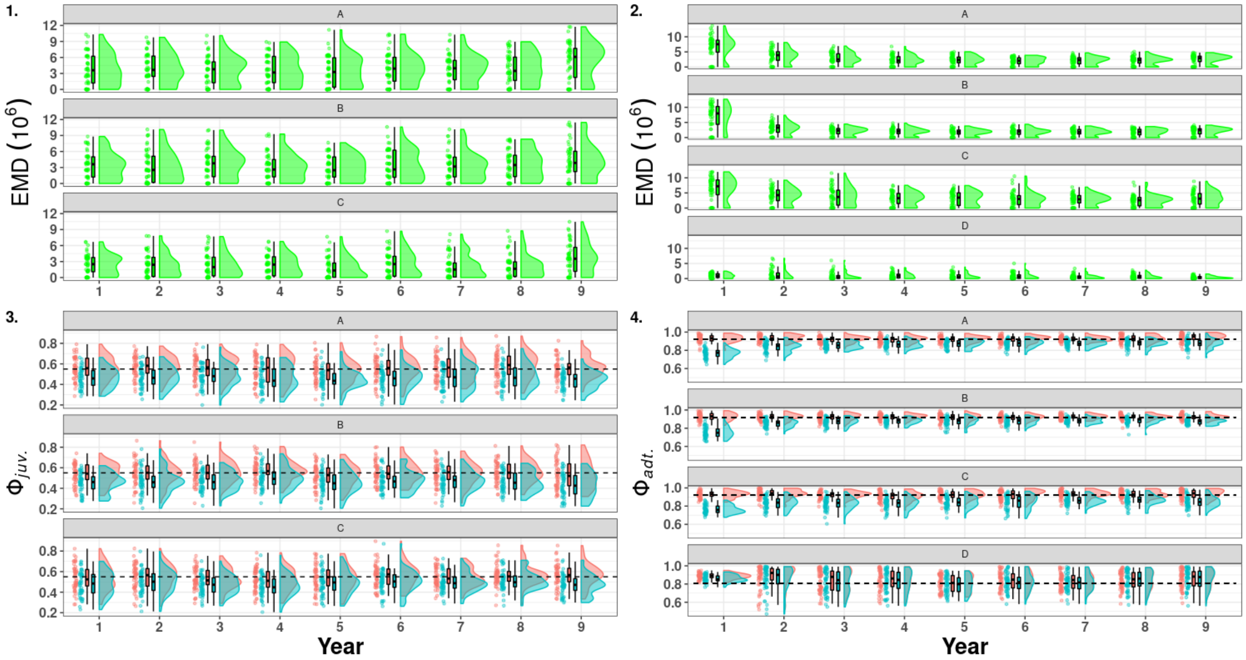

Figure S14: Raincloud plots of the Earth Mover Distance (green) between the medians of the posterior distribution of survival estimated for the model accounting for mark loss and recycling (red) and not accounting for mark loss (blue). On the left juvenile survival ($\phi_{juv.}$, 1 & 3), on the right adult survival ($\phi_{adt.}$, 2 & 4) in scenario 1 (long-lived species with detection) with a simulated mark loss probability of 0.25. A, B, C, D denotes the name of the states and the dashed line correspond to the mean of the true values.

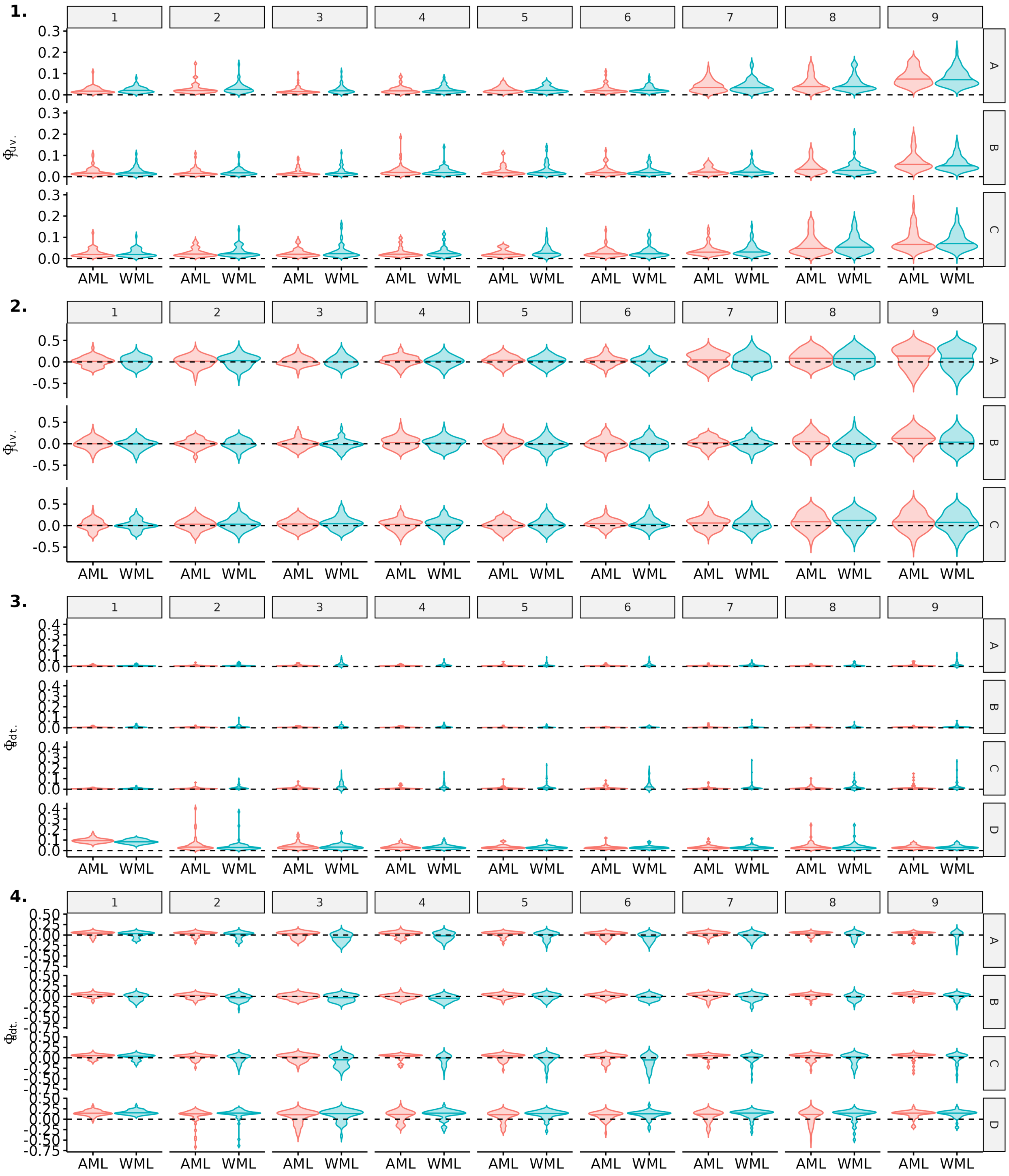

Figure S15: Comparison of precision and bias for estimates of juvenile and adult survival probabilities between model accounting for mark loss (AML) or not (WML), over the 9 recapture occasions. Here, scenario 2 (long-lived species with low detection) is shown for a simulated mark loss probability of 0.25. Violin plots show the distributions of mean precision (1,3) and bias (2,4) over 50 simulations. The median of each distribution is shown with an horizontal line.

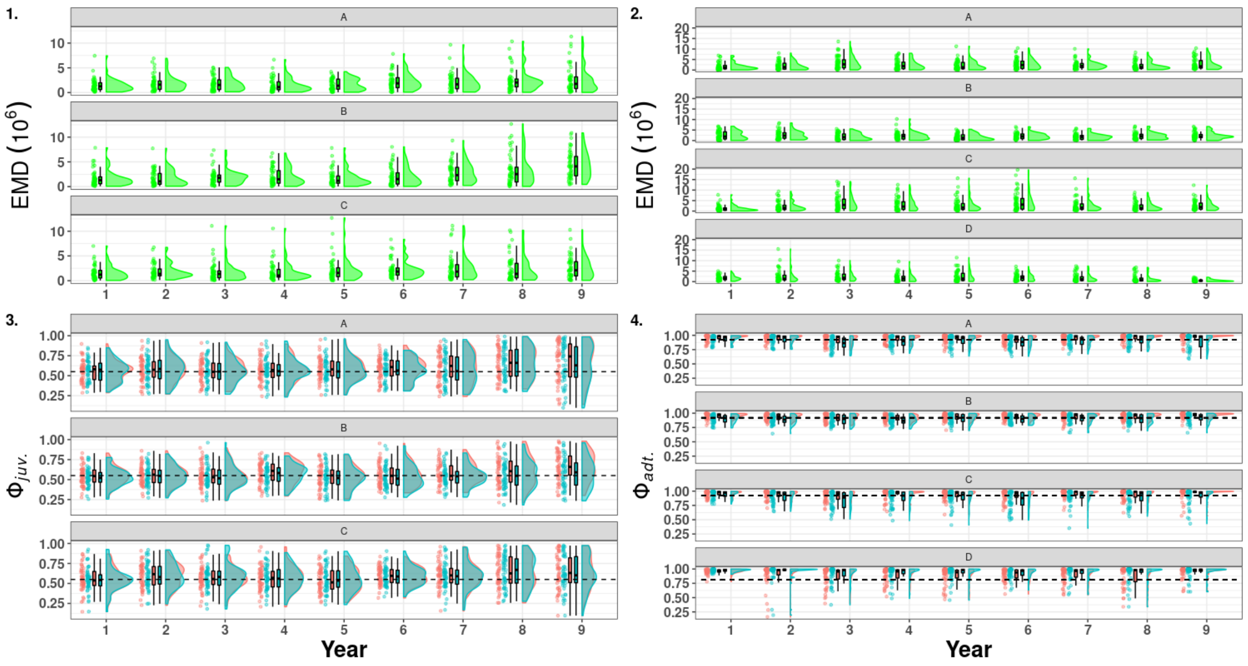

Figure S16: Raincloud plots of the Earth Mover Distance (green) between the medians of the posterior distribution of survival estimated for the model accounting for mark loss and recycling (red) and not accounting for mark loss (blue). On the left juvenile survival ($\phi_{juv.}$, 1 & 3), on the right adult survival ($\phi_{adt.}$, 2 & 4) in scenario 2 (long-lived species with low detection) with a simulated mark loss probability of 0.25. A, B, C, D denotes the name of the states and the dashed line correspond to the mean of the true values.

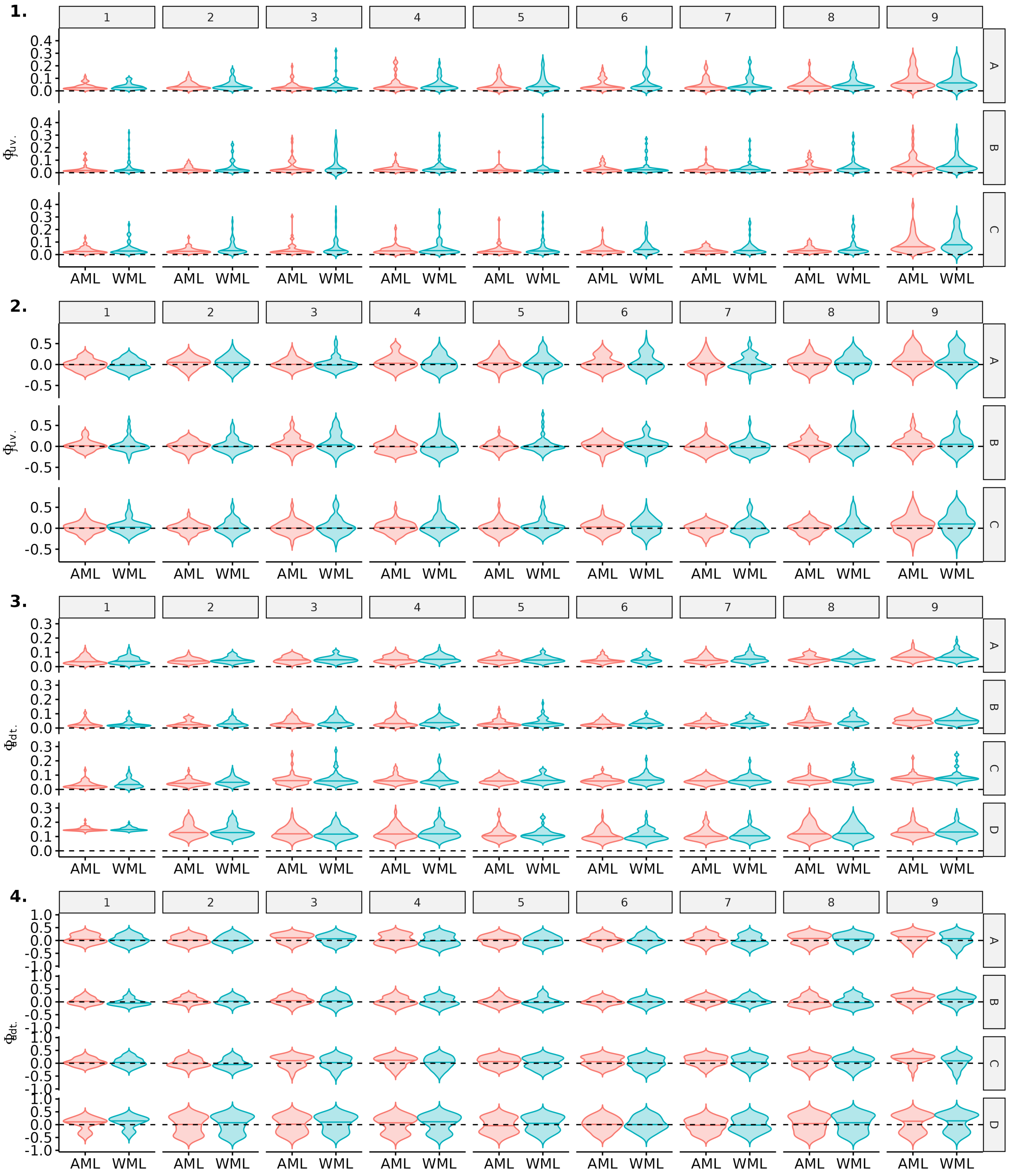

Figure S17: Comparison of precision and bias for estimates of juvenile and adult survival probabilities between model accounting for mark loss (AML) or not (WML), over the 9 recapture occasions. Here, scenario 3 (short-lived species with low detection) is shown for a simulated mark loss probability of 0.25. Violin plots show the distributions of mean precision (1,3) and bias (2,4) over 50 simulations. The median of each distribution is shown with an horizontal line.

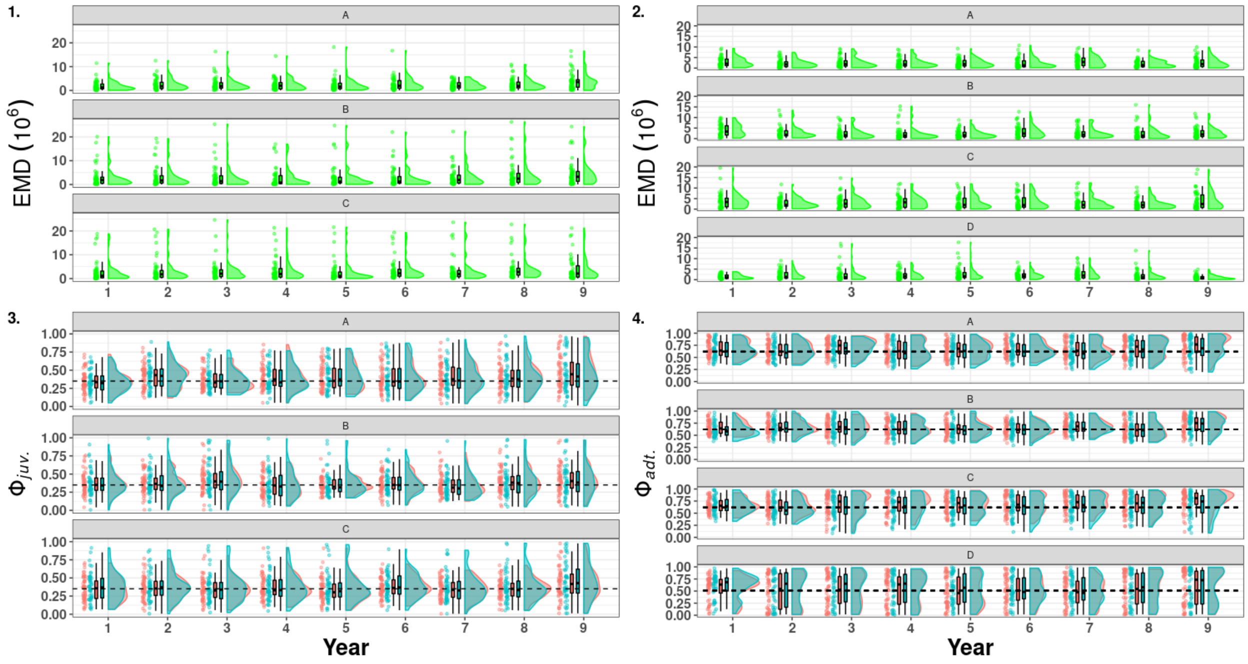

Figure S18: Raincloud plots of the Earth Mover Distance (green) between the medians of the posterior distribution of survival estimated for the model accounting for mark loss and recycling (red) and not accounting for mark loss (blue). On the left juvenile survival ($\phi_{juv.}$, 1 & 3), on the right adult survival ($\phi_{adt.}$, 2 & 4) in scenario 3 (short-lived species with low detection) with a simulated mark loss probability of 0.25. A, B, C, D denotes the name of the states and the dashed line correspond to the mean of the true values.

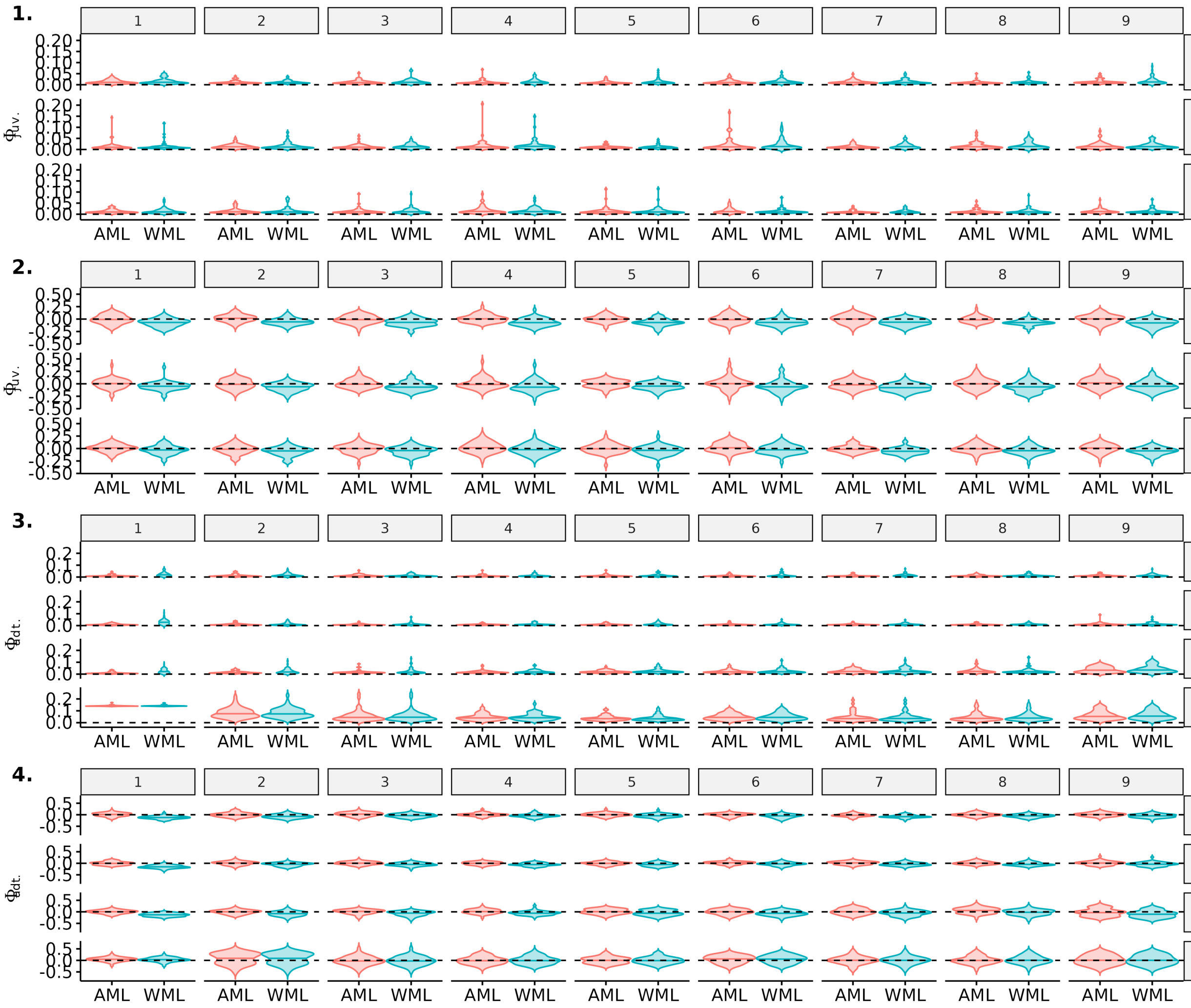

Figure S19: Comparison of precision and bias for estimates of juvenile and adult survival probabilities between model accounting for mark loss (AML) or not (WML), over the 9 recapture occasions. Here, scenario 4 (short-lived species with high detection) is shown for a simulated mark loss probability of 0.25. Violin plots show the distributions of mean precision (1,3) and bias (2,4) over 50 simulations. The median of each distribution is shown with an horizontal line.

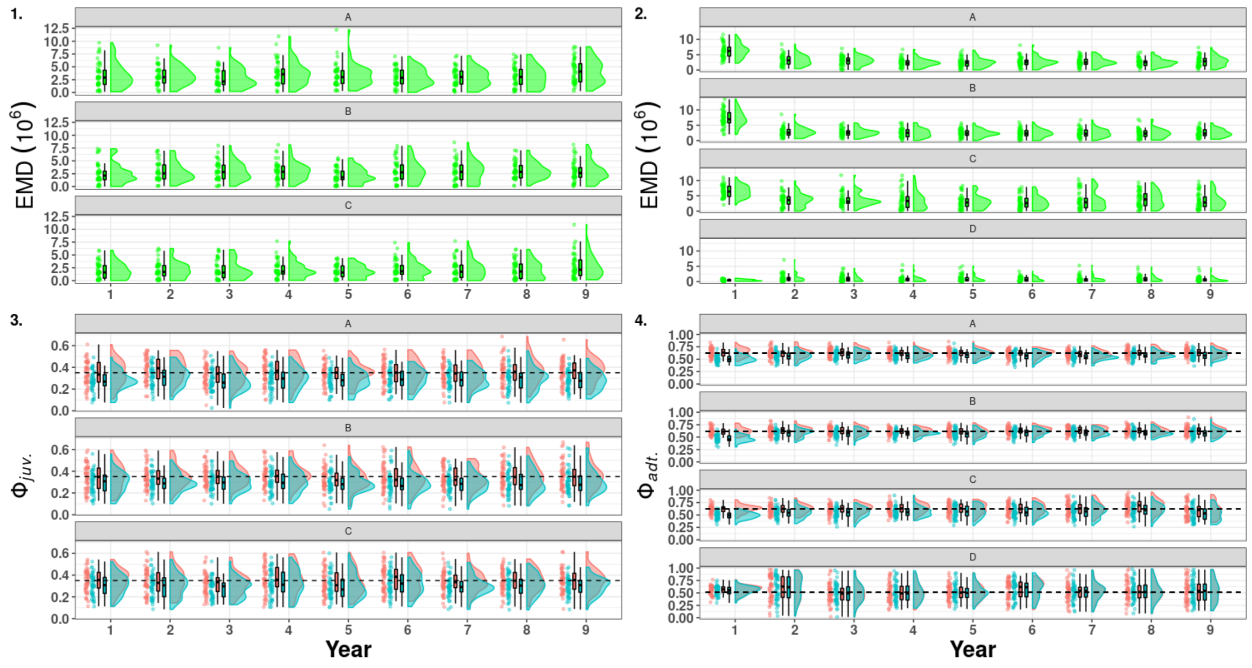

Figure S20: Raincloud plots of the Earth Mover Distance (green) between the medians of the posterior distribution of survival estimated for the model accounting for mark loss and recycling (red) and not accounting for mark loss (blue). On the left juvenile survival ($\phi_{juv.}$, 1 & 3), on the right adult survival ($\phi_{adt.}$, 2 & 4) in scenario 4 (short-lived species with high detection) with a simulated mark loss probability of 0.25. A, B, C, D denotes the name of the states and the dashed line correspond to the mean of the true values.

##### 3.3.3 Simulations with mark loss rate of 0.4

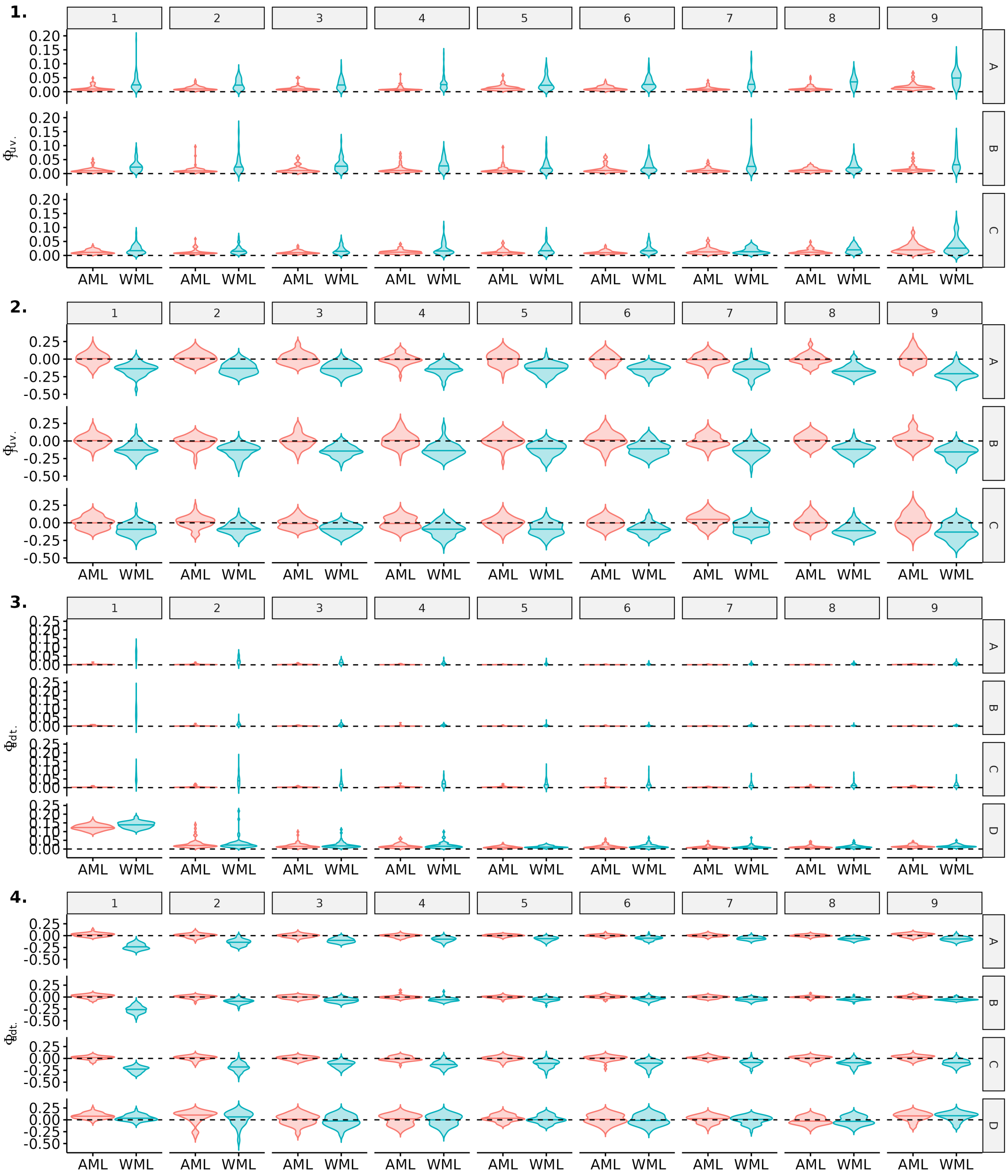

Figure S21: Comparison of precision and bias for estimates of juvenile and adult survival probabilities between model accounting for mark loss (AML) or not (WML), over the 9 recapture occasions. Here, scenario 1 (long-lived species with high detection) is shown for a simulated mark loss probability of 0.4. Violin plots show the distributions of mean precision (1,3) and bias (2,4) over 50 simulations. The median of each distribution is shown with an horizontal line.

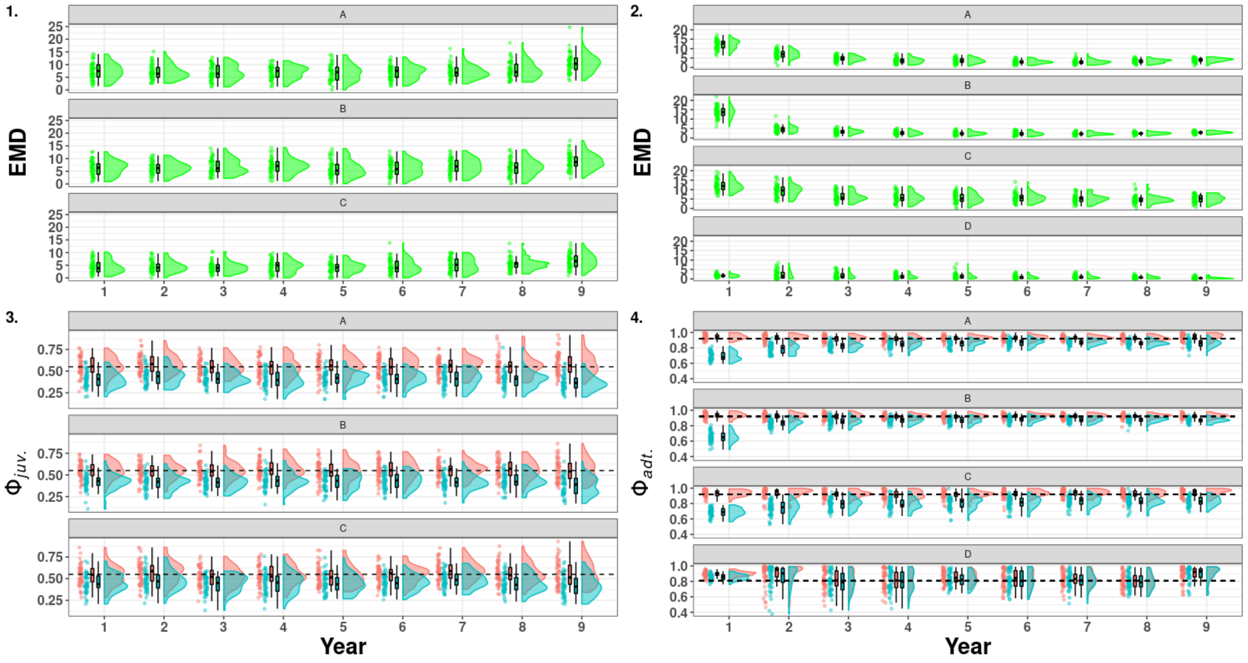

Figure S22: Raincloud plots of the Earth Mover Distance (green) between the medians of the posterior distribution of survival estimated for the model accounting for mark loss and recycling (red) and not accounting for mark loss (blue). On the left juvenile survival ($\phi_{juv.}$, 1 & 3), on the right adult survival ($\phi_{adt.}$, 2 & 4) in scenario 1 (long-lived species with detection) with a simulated mark loss probability of 0.4. A, B, C, D denotes the name of the states and the dashed line correspond to the mean of the true values.

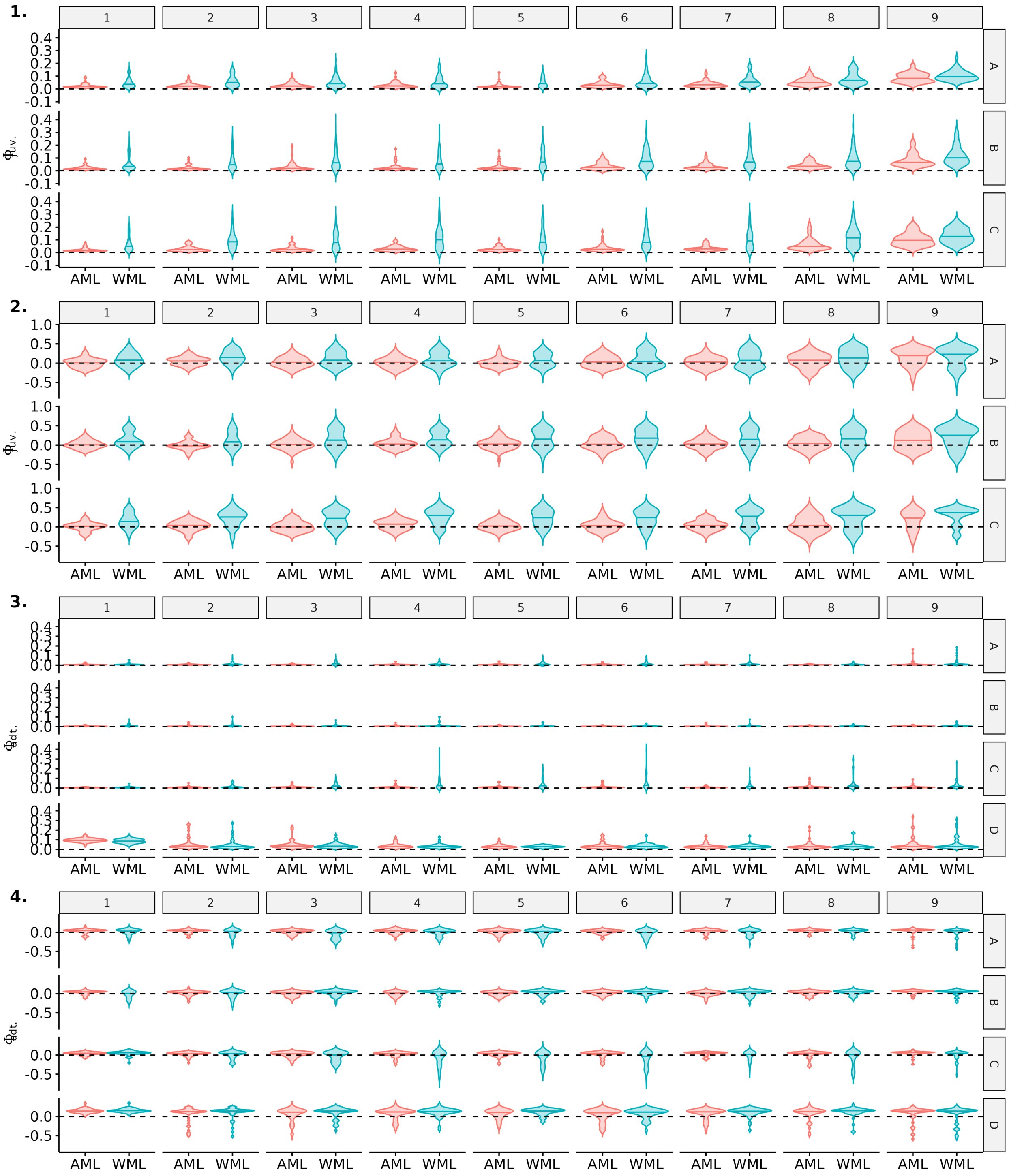

Figure S23: Comparison of precision and bias for estimates of juvenile and adult survival probabilities between model accounting for mark loss (AML) or not (WML), over the 9 recapture occasions. Here, scenario 2 (long-lived species with low detection) is shown for a simulated mark loss probability of 0.4. Violin plots show the distributions of mean precision (1,3) and bias (2,4) over 50 simulations. The median of each distribution is shown with an horizontal line.

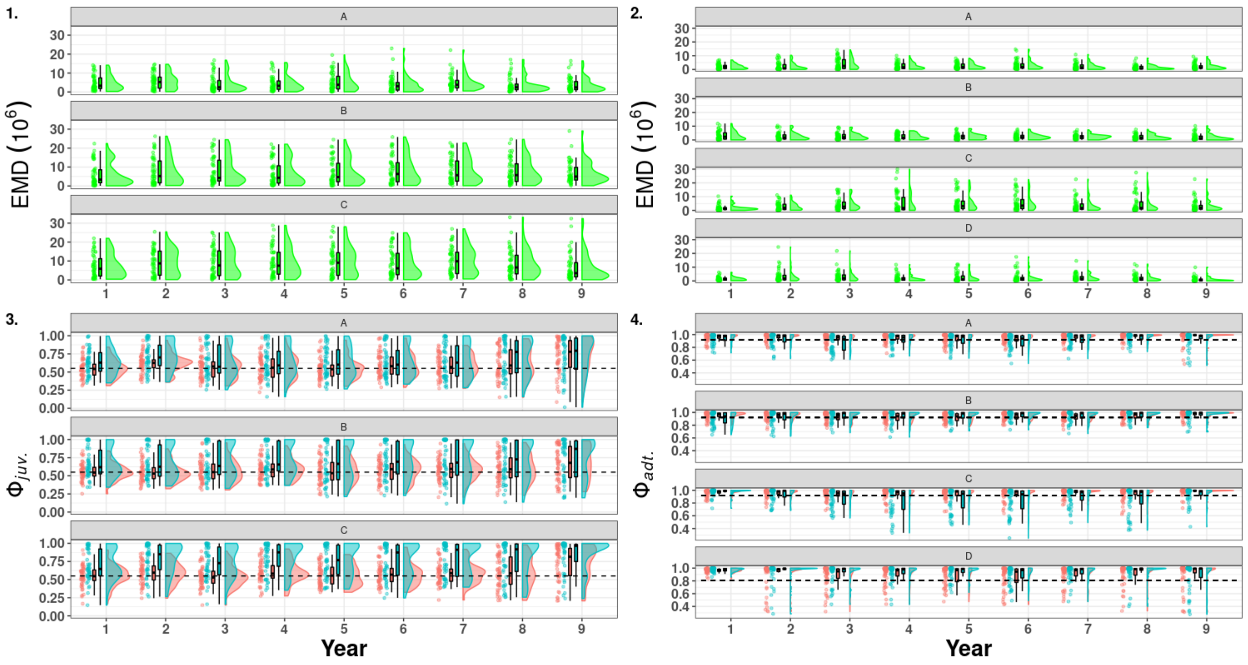

Figure S24: Raincloud plots of the Earth Mover Distance (green) between the medians of the posterior distribution of survival estimated for the model accounting for mark loss and recycling (red) and not accounting for mark loss (blue). On the left juvenile survival ($\phi_{juv.}$, 1 & 3), on the right adult survival ($\phi_{adt.}$, 2 & 4) in scenario 2 (long-lived species with low detection) with a simulated mark loss probability of 0.4. A, B, C, D denotes the name of the states and the dashed line correspond to the mean of the true values.

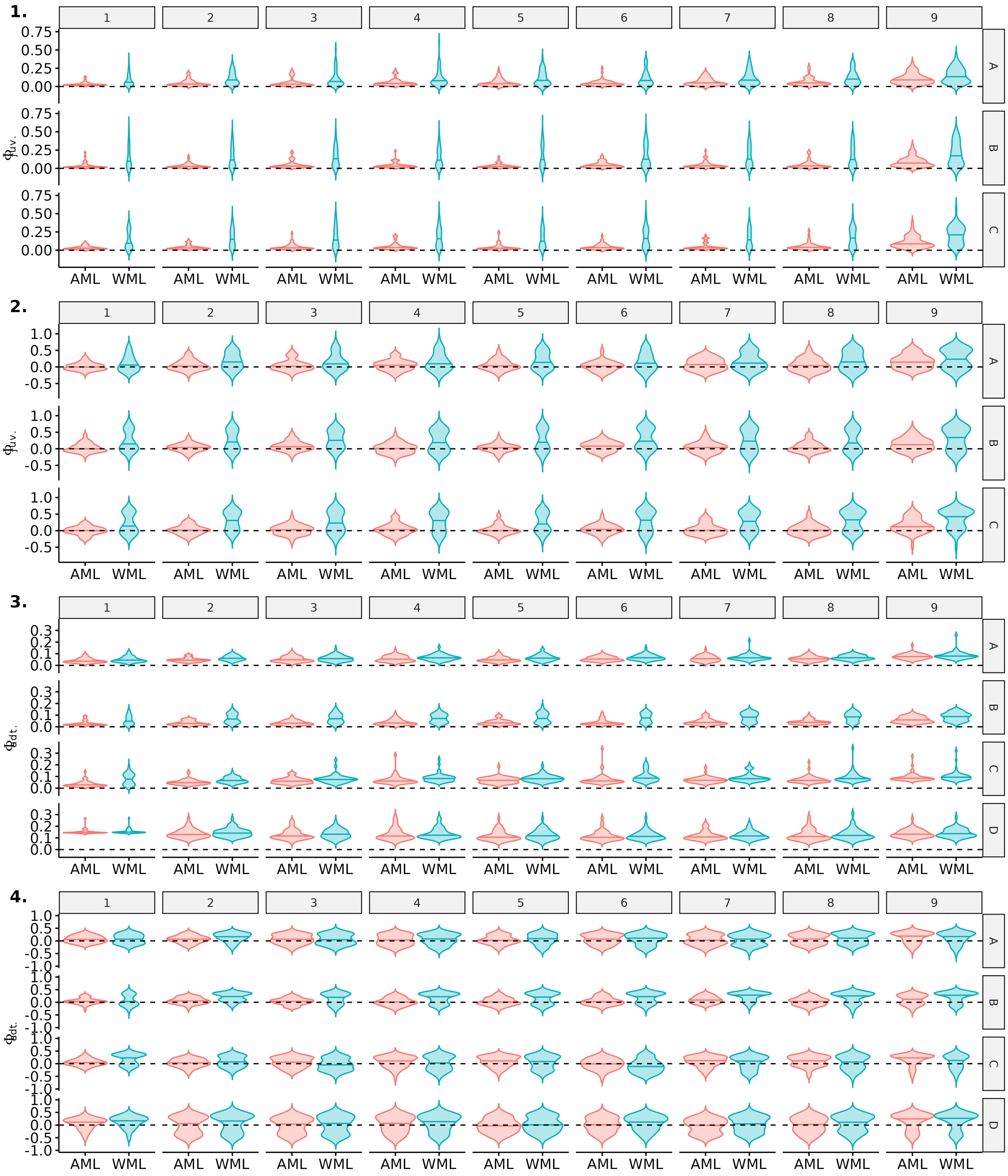

Figure S25: Comparison of precision and bias for estimates of juvenile and adult survival probabilities between model accounting for mark loss (AML) or not (WML), over the 9 recapture occasions. Here, scenario 3 (short-lived species with low detection) is shown for a simulated mark loss probability of 0.4. Violin plots show the distributions of mean precision (1,3) and bias (2,4) over 50 simulations. The median of each distribution is shown with an horizontal line.

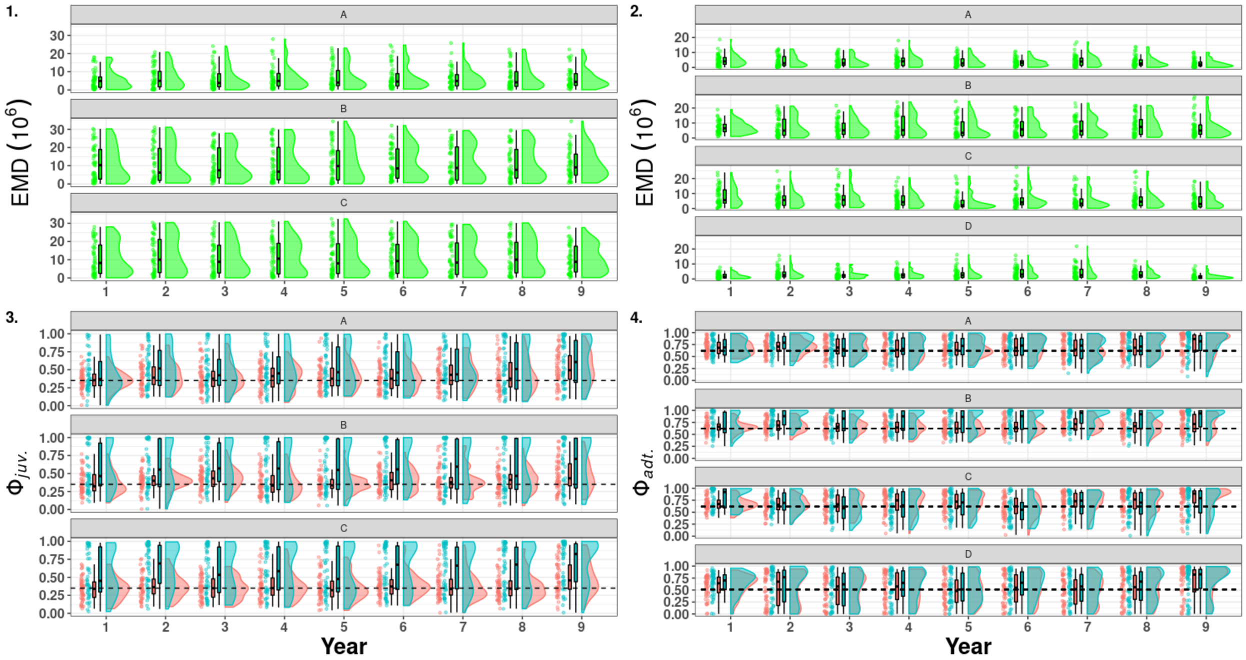

Figure S26: Raincloud plots of the Earth Mover Distance (green) between the medians of the posterior distribution of survival estimated for the model accounting for mark loss and recycling (red) and not accounting for mark loss (blue). On the left juvenile survival ($\phi_{juv.}$, 1 & 3), on the right adult survival ($\phi_{adt.}$, 2 & 4) in scenario 3 (short-lived species with low detection with a simulated mark loss probability of 0.4). A, B, C, D denotes the name of the states and the dashed line correspond to the mean of the true values.

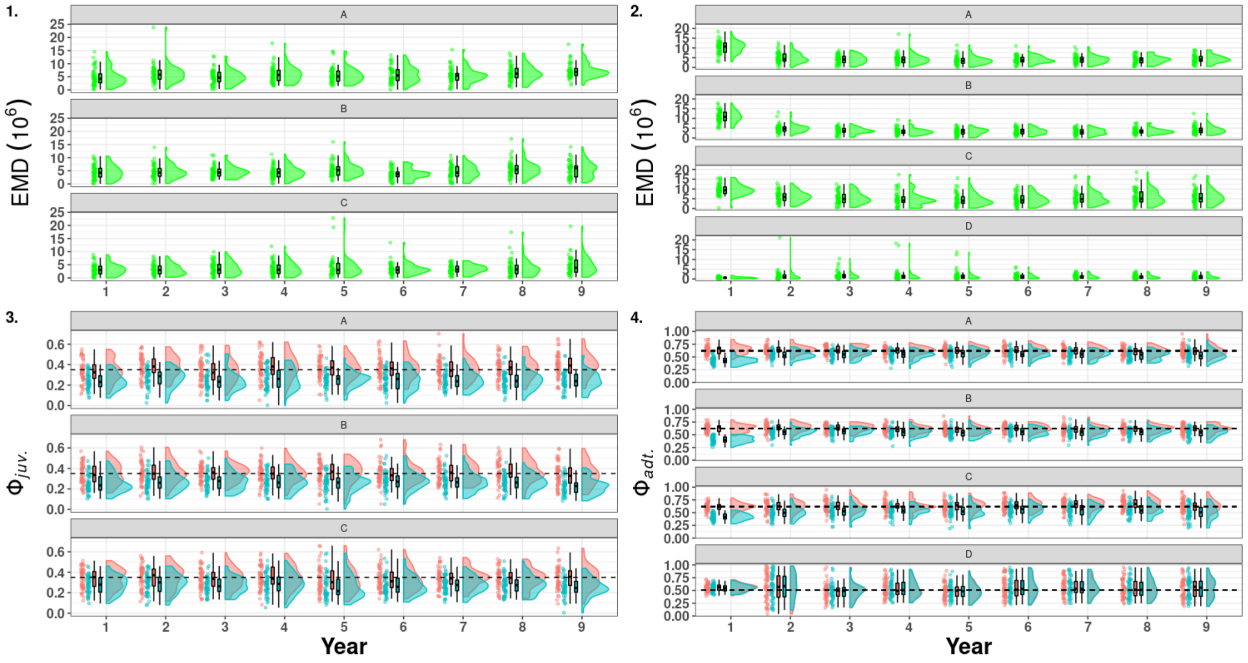

Figure S27: Raincloud plots of the Earth Mover Distance (green) between the medians of the posterior distribution of survival estimated for the model accounting for mark loss and recycling (red) and not accounting for mark loss (blue). On the left juvenile survival ($\phi_{juv.}$, 1 & 3), on the right adult survival ($\phi_{adt.}$, 2 & 4) in scenario 4 (short-lived species with high detection) with a simulated mark loss probability of 0.4. A, B, C, D denotes the name of the states and the dashed line correspond to the mean of the true values.

#### 3.4 Capture and resighting probability estimates

Below, we compared the bias (median - truth) and the precision (mean squared errors MSE = bias$^2$+ variance) of the detection probability (capture and resighting) between the model without recycling and with recycling. We also use the Earth Mover Distance (EMD) to compare the distribution of the medians of these parameters. The density distribution of these medians were also displayed. In our simulations, the individuals could only be captured when in states A, B or C but they could be resighted in every states including D.

##### 3.4.1 Simulations with mark loss rate of 0.05

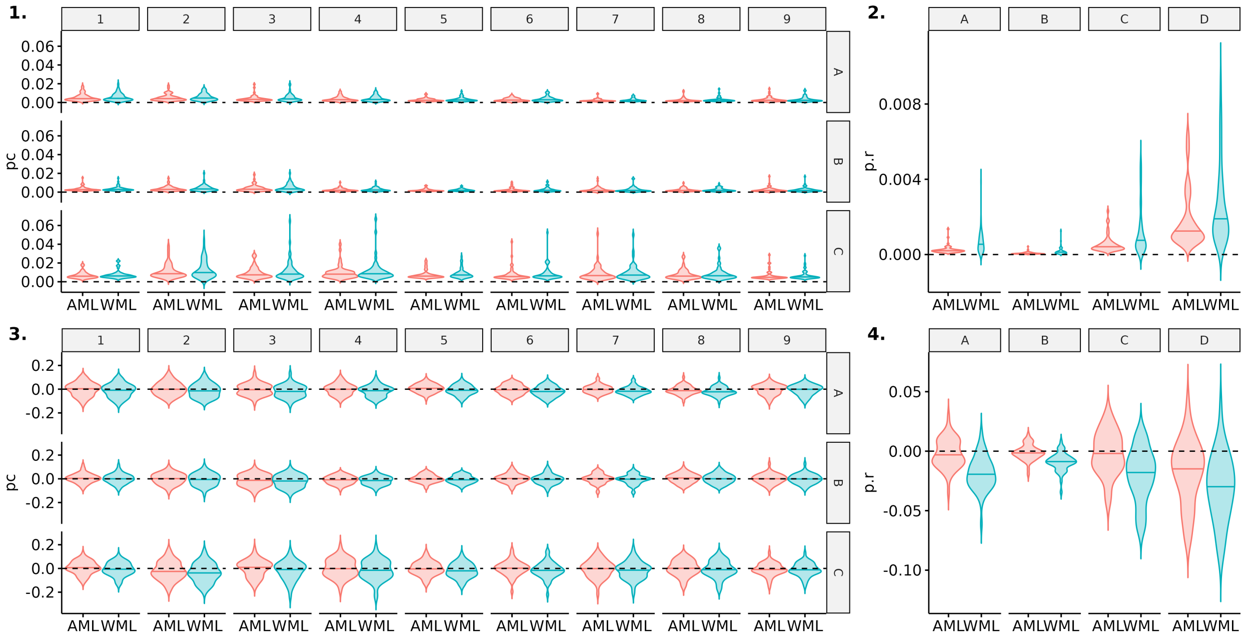

Figure S28: Comparison of precision and bias for estimates of capture ($pc$) and resigthing probabilities ($p.r$) between model accounting for mark loss (AML) or not (WML). Here, scenario 1 (long-lived species with high detection) is shown for a simulated mark loss probability of 0.05. Violin plots show the distributions of mean precision (1,2) and bias (3,4) over 50 simulations. The median of each distribution is shown with an horizontal line.

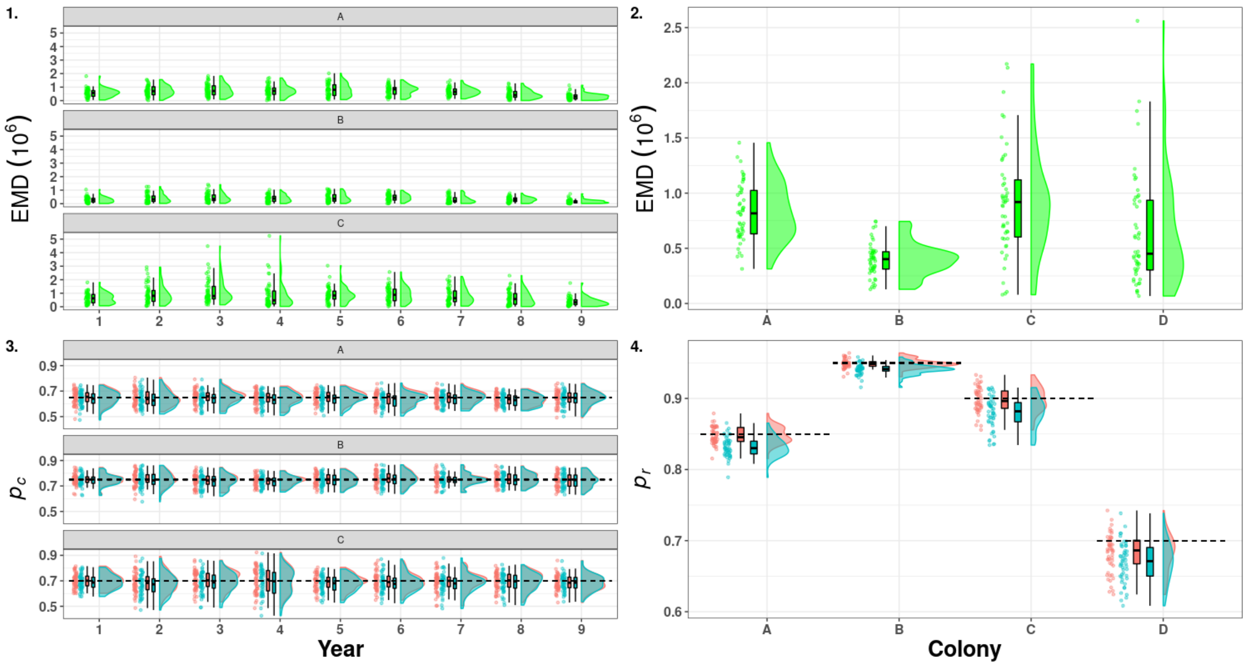

Figure S29: Raincloud plots of the Earth Mover Distance (green) between the medians of the posterior distribution of detection estimated for the model accounting for mark loss and recycling (red) and not accounting for mark loss (blue). On the left capture probability ($p_c$, 1 & 3), on the right resighting probability ($p_r$, 2 & 4) in scenario 1 (long-lived species with high detection). A, B, C, D denotes the name of states and the dashed line correspond to the mean of the true values.

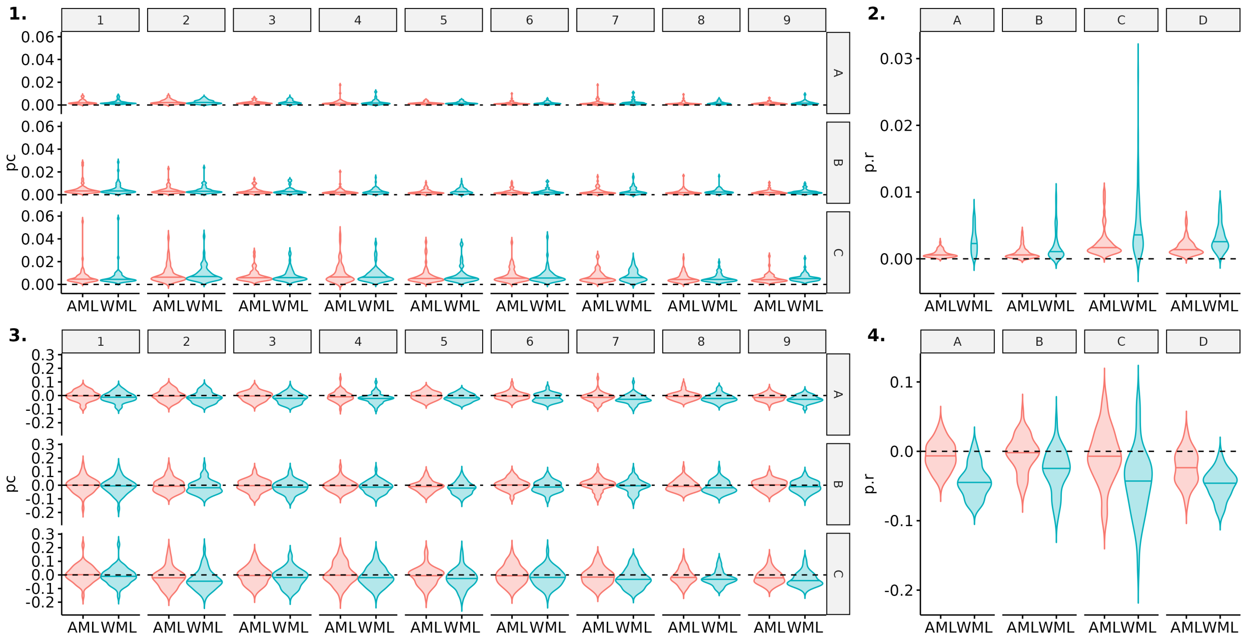

Figure S30: Comparison of precision and bias for estimates of capture ($pc$) and resigthing probabilities ($p.r$) between model accounting for mark loss (AML) or not (WML). Here, scenario 2 (long-lived species with low detection) is shown for a simulated mark loss probability of 0.05. Violin plots show the distributions of mean precision (1,2) and bias (3,4) over 50 simulations. The median of each distribution is shown with an horizontal line.

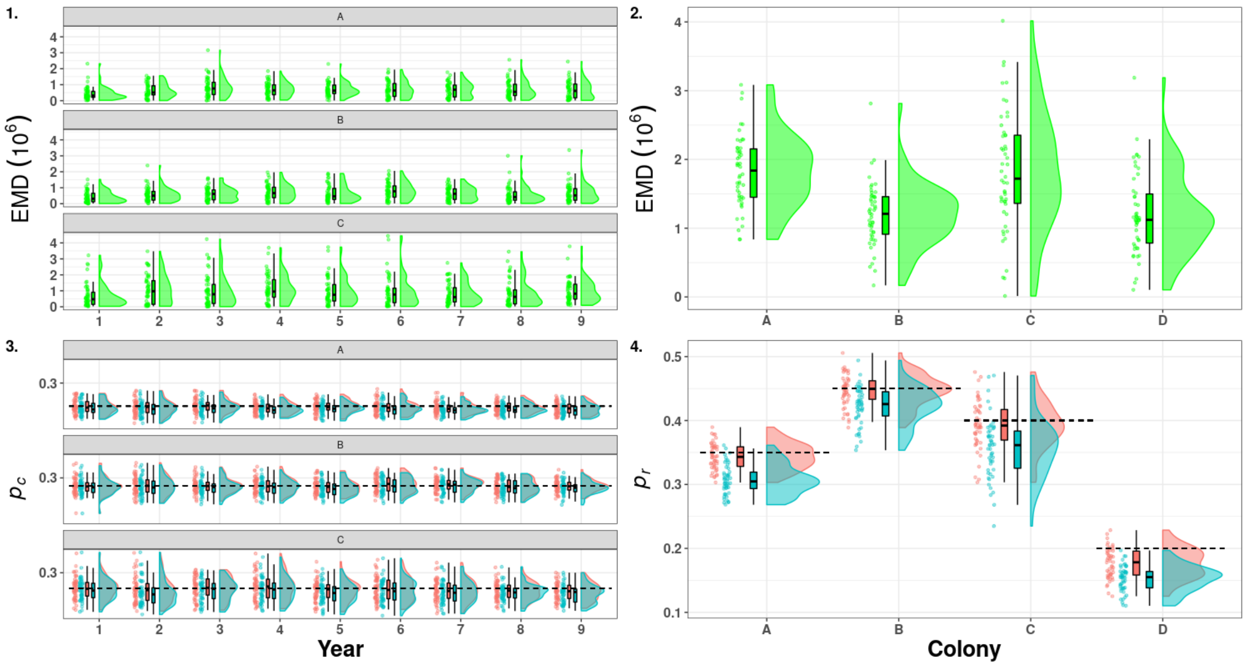

Figure S31: Raincloud plots of the Earth Mover Distance (green) between the medians of the posterior distribution of detection estimated for the model accounting for mark loss and recycling (red) and not accounting for mark loss (blue). On the left capture probability ($p_c$, 1 & 3), on the right resighting probability ($p_r$, 2 & 4) in scenario 2 (long-lived species with low detection). A, B, C, D denotes the name of states and the dashed line correspond to the mean of the true values.

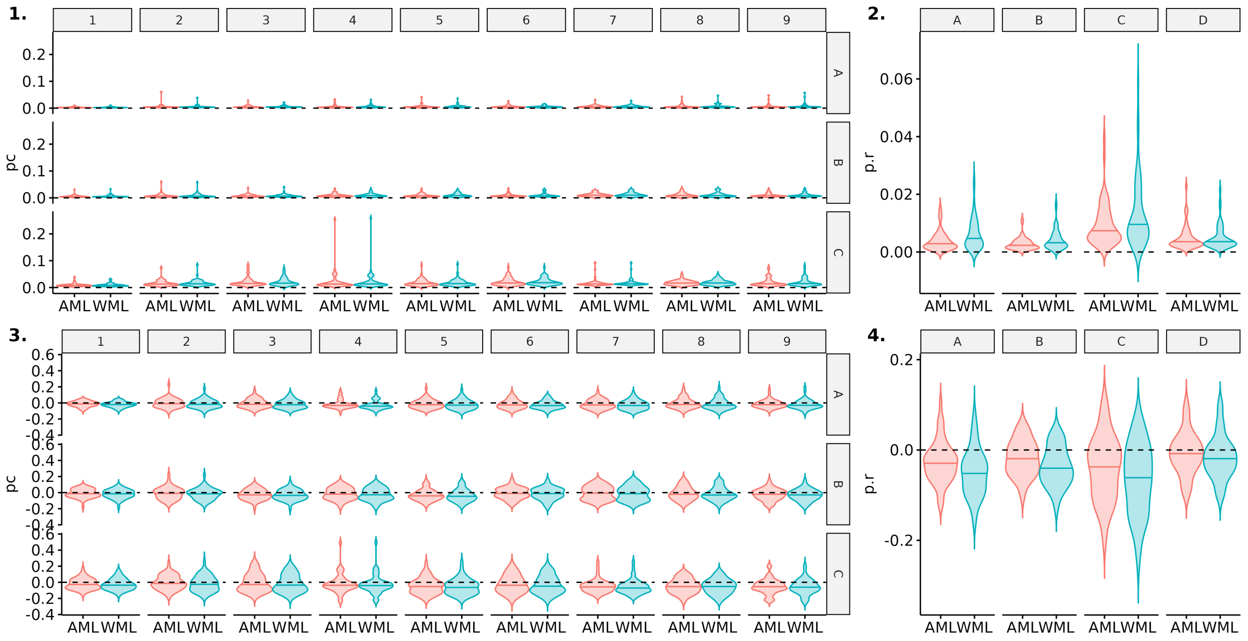

Figure S32: Comparison of precision and bias for estimates of capture ($pc$) and resigthing probabilities ($p.r$) between model accounting for mark loss (AML) or not (WML). Here, scenario 3 (short-lived species with low detection) is shown for a simulated mark loss probability of 0.05. Violin plots show the distributions of mean precision (1,2) and bias (3,4) over 50 simulations. The median of each distribution is shown with an horizontal line.

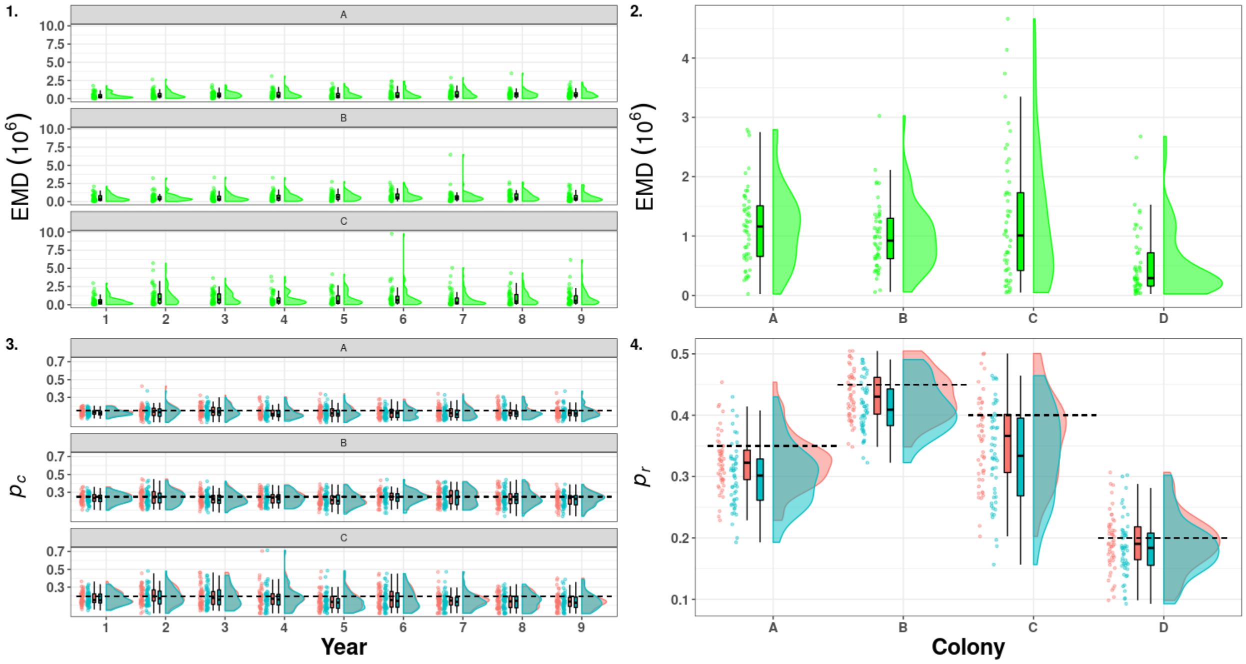

Figure S33: Raincloud plots of the Earth Mover Distance (green) between the medians of the posterior distribution of detection estimated for the model accounting for mark loss and recycling (red) and not accounting for mark loss (blue). On the left capture probability ($p_c$, 1 & 3), on the right resighting probability ($p_r$, 2 & 4) in scenario 3 (short-lived species with low detection). A, B, C, D denotes the name of states and the dashed line correspond to the mean of the true values.

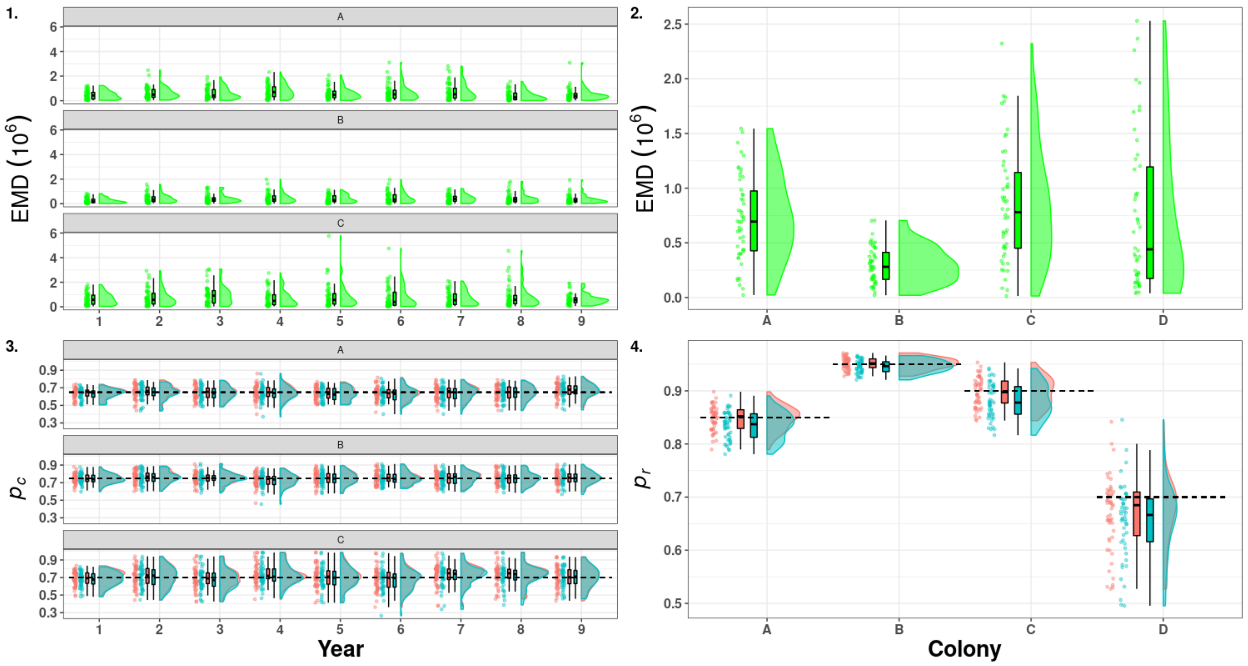

Figure S34: Raincloud plots of the Earth Mover Distance (green) between the medians of the posterior distribution of detection estimated for the model accounting for mark loss and recycling (red) and not accounting for mark loss (blue). On the left capture probability ($p_c$, 1 & 3), on the right resighting probability ($p_r$, 2 & 4) in scenario 4 (short-lived species with high detection). A, B, C, D denotes the name of states and the dashed line correspond to the mean of the true values.

##### 3.4.2 Simulations with mark loss rate of 0.25

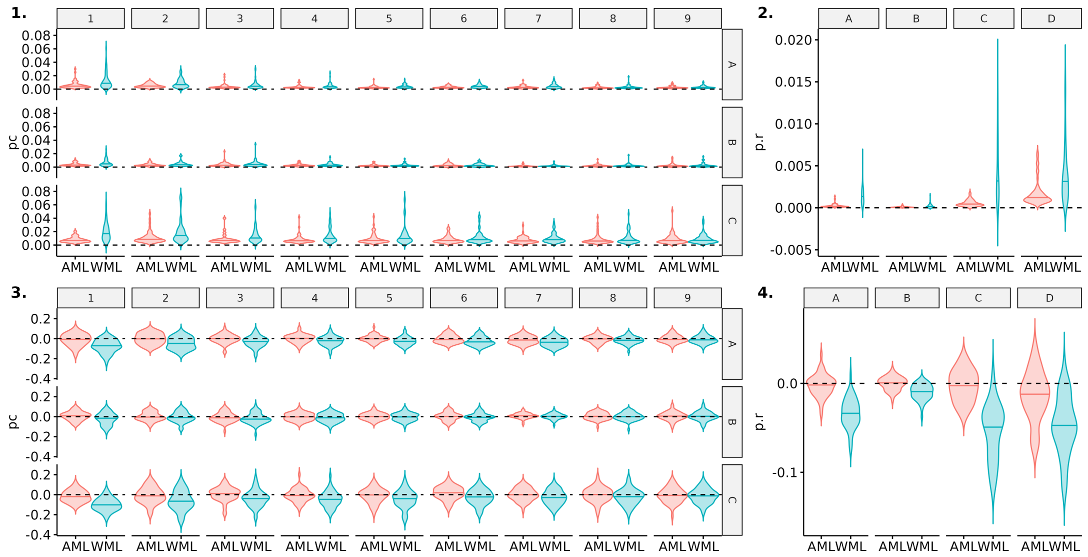

Figure S35: Comparison of precision and bias for estimates of capture ($pc$) and resigthing probabilities ($p.r$) between model accounting for mark loss (AML) or not (WML). Here, scenario 1 (long-lived species with high detection) is shown for a simulated mark loss probability of 0.25. Violin plots show the distributions of mean precision (1,2) and bias (3,4) over 50 simulations. The median of each distribution is shown with an horizontal line.

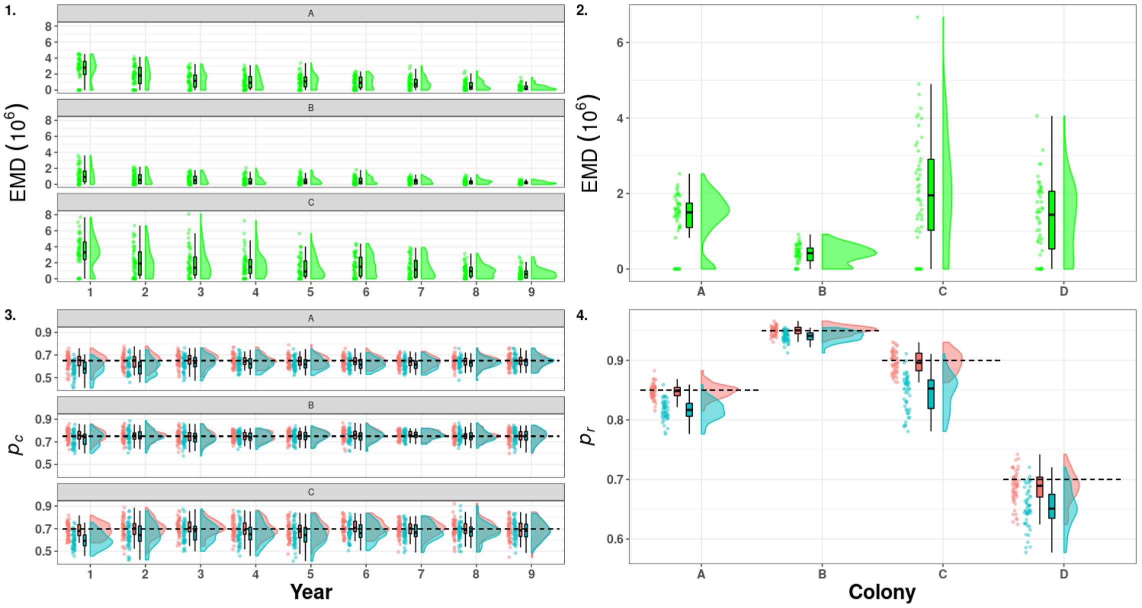

Figure S36: Raincloud plots of the Earth Mover Distance (green) between the medians of the posterior distribution of detection estimated for the model accounting for mark loss and recycling (red) and not accounting for mark loss (blue). On the left capture probability ($p_c$, 1 & 3), on the right resighting probability ($p_r$, 2 & 4) in scenario 1 (long-lived species with high detection). A, B, C, D denotes the name of states and the dashed line correspond to the mean of the true values.

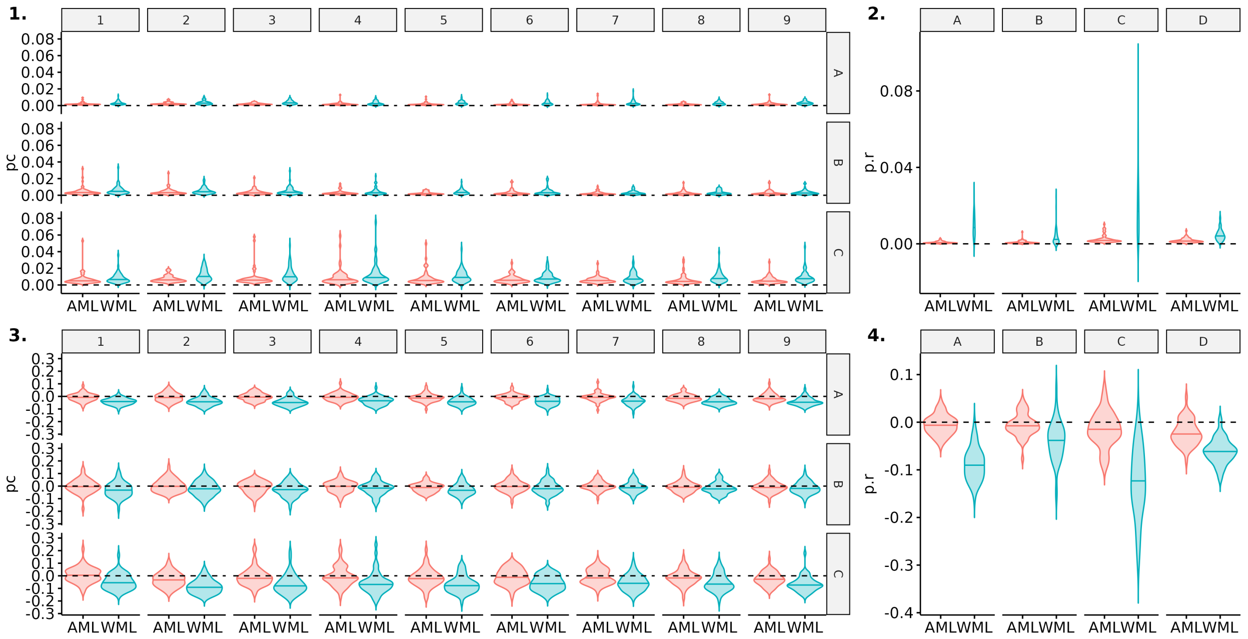

Figure S37: Comparison of precision and bias for estimates of capture ($pc$) and resigthing probabilities ($p.r$) between model accounting for mark loss (AML) or not (WML). Here, scenario 2 (long-lived species with low detection) is shown for a simulated mark loss probability of 0.25. Violin plots show the distributions of mean precision (1,2) and bias (3,4) over 50 simulations. The median of each distribution is shown with an horizontal line.

.

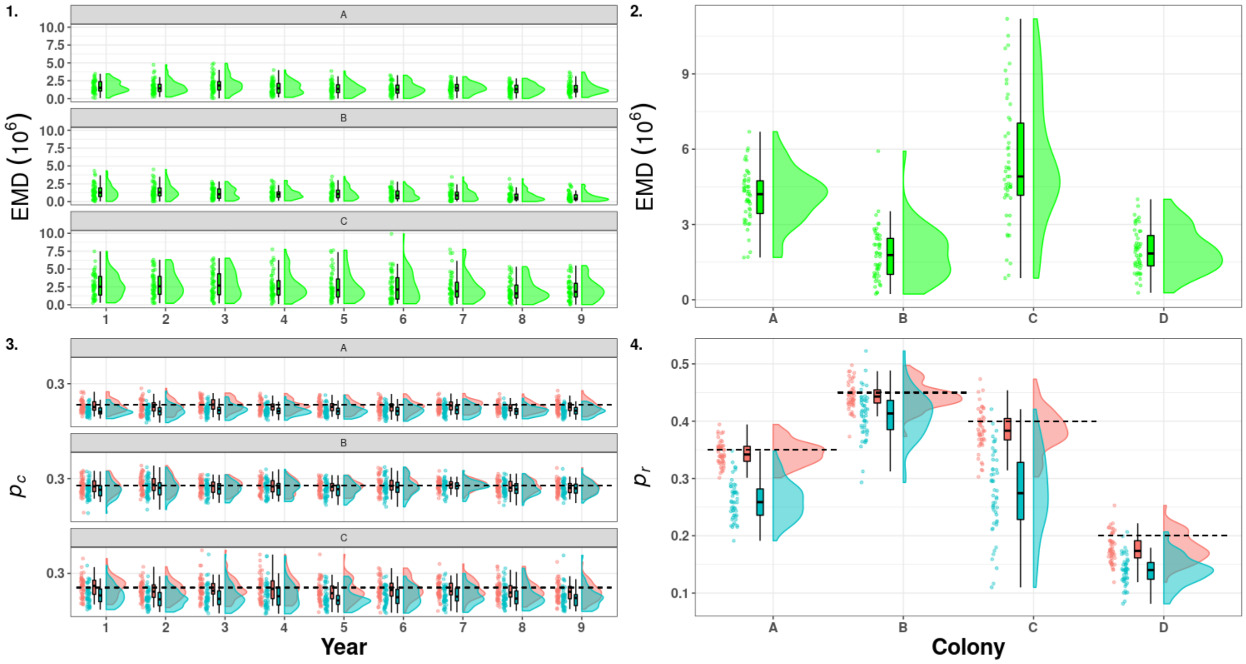

Figure S38: Raincloud plots of the Earth Mover Distance (green) between the medians of the posterior distribution of detection estimated for the model accounting for mark loss and recycling (red) and not accounting for mark loss (blue). On the left capture probability ($p_c$, 1 & 3), on the right resighting probability ($p_r$, 2 & 4) in scenario 2 (long-lived species with low detection). A, B, C, D denotes the name of states and the dashed line correspond to the mean of the true values.

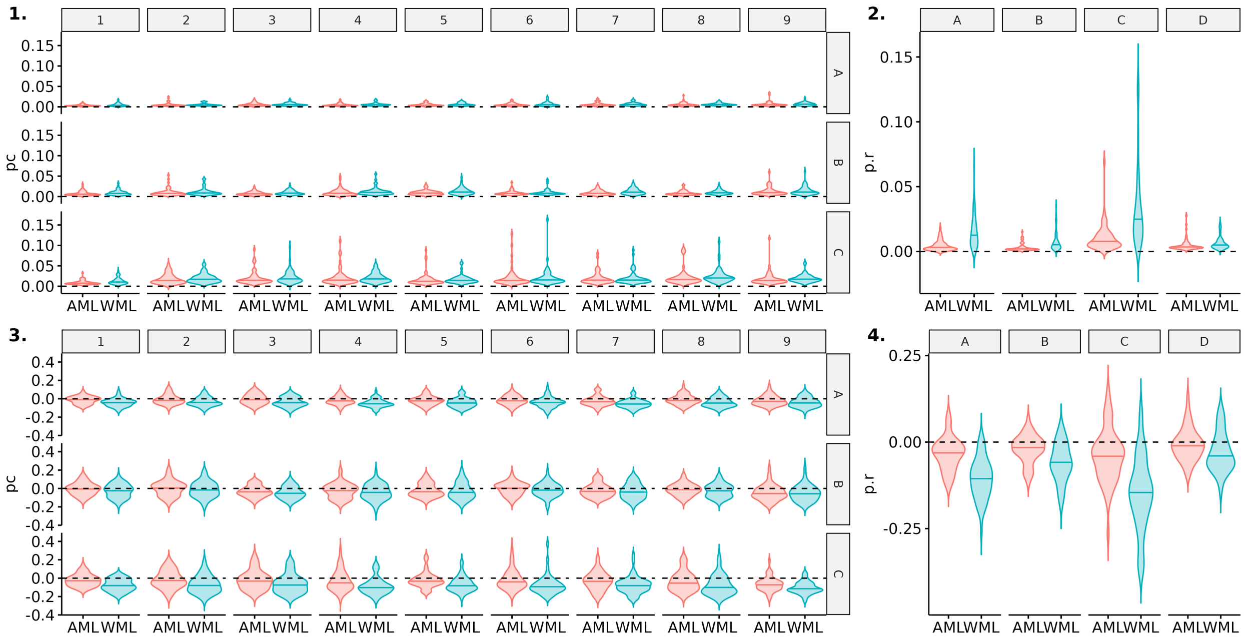

Figure S39: Comparison of precision and bias for estimates of capture ($pc$) and resigthing probabilities ($p.r$) between model accounting for mark loss (AML) or not (WML). Here, scenario 3 (short-lived species with low detection) is shown for a simulated mark loss probability of 0.25. Violin plots show the distributions of mean precision (1,2) and bias (3,4) over 50 simulations. The median of each distribution is shown with an horizontal line.

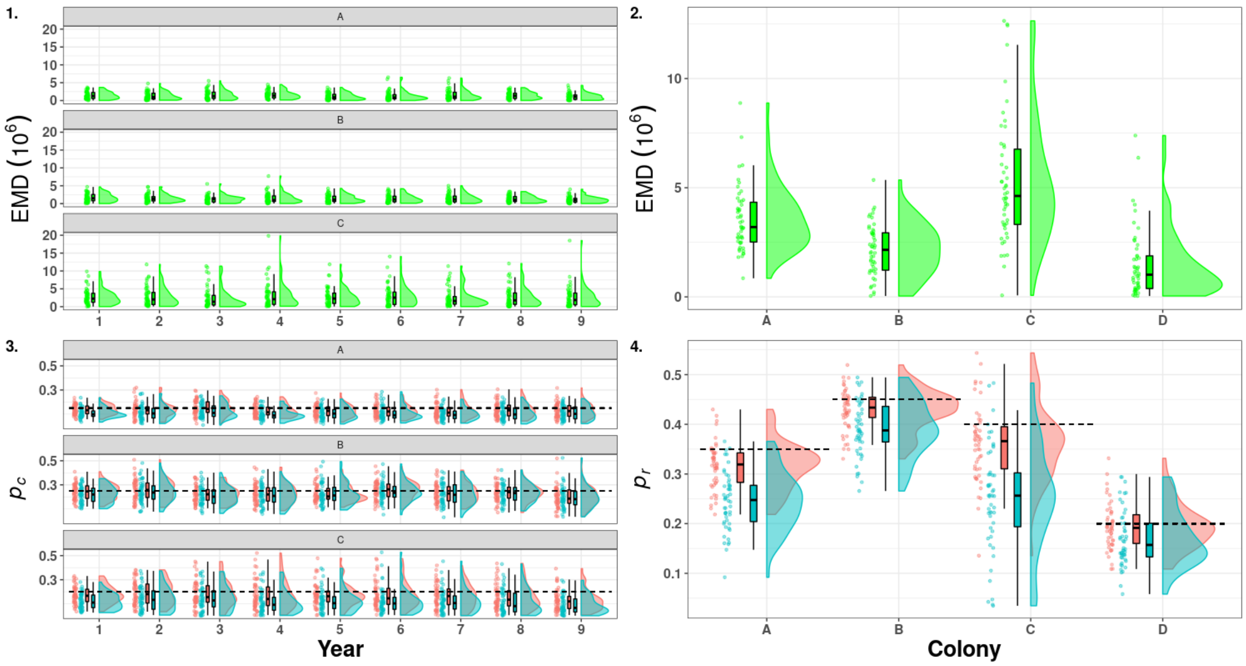

Figure S40: Raincloud plots of the Earth Mover Distance (green) between the medians of the posterior distribution of detection estimated for the model accounting for mark loss and recycling (red) and not accounting for mark loss (blue). On the left capture probability ($p_c$, 1 & 3), on the right resighting probability ($p_r$, 2 & 4) in scenario 3 (short-lived species with low detection). A, B, C, D denotes the name of states and the dashed line correspond to the mean of the true values.

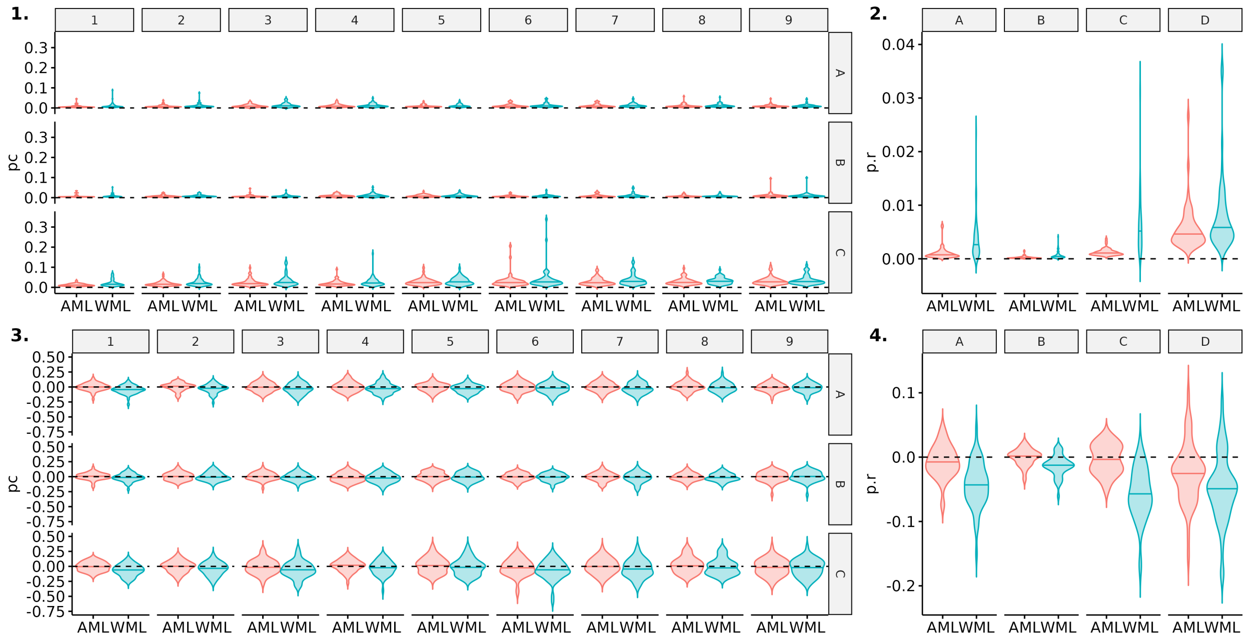

Figure S41: Comparison of precision and bias for estimates of capture ($pc$) and resigthing probabilities ($p.r$) between model accounting for mark loss (AML) or not (WML). Here, scenario 4 (short-lived species with high detection) is shown for a simulated mark loss probability of 0.25. Violin plots show the distributions of mean precision (1,2) and bias (3,4) over 50 simulations. The median of each distribution is shown with an horizontal line.

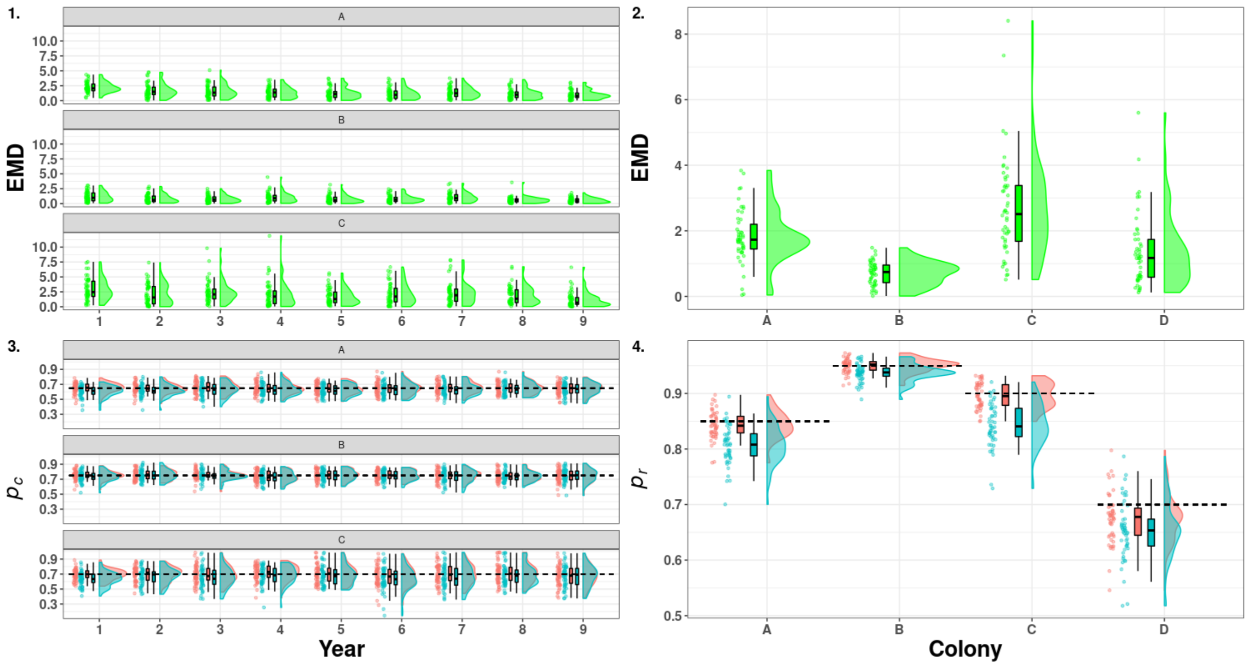

Figure S42: Raincloud plots of the Earth Mover Distance (green) between the medians of the posterior distribution of detection estimated for the model accounting for mark loss and recycling (red) and not accounting for mark loss (blue). On the left capture probability ($p_c$, 1 & 3), on the right resighting probability ($p_r$, 2 & 4) in scenario 2 (short-lived species with high detection). A, B, C, D denotes the name of states and the dashed line correspond to the mean of the true values.

##### 3.4.3 Simulations with mark loss rate of 0.4

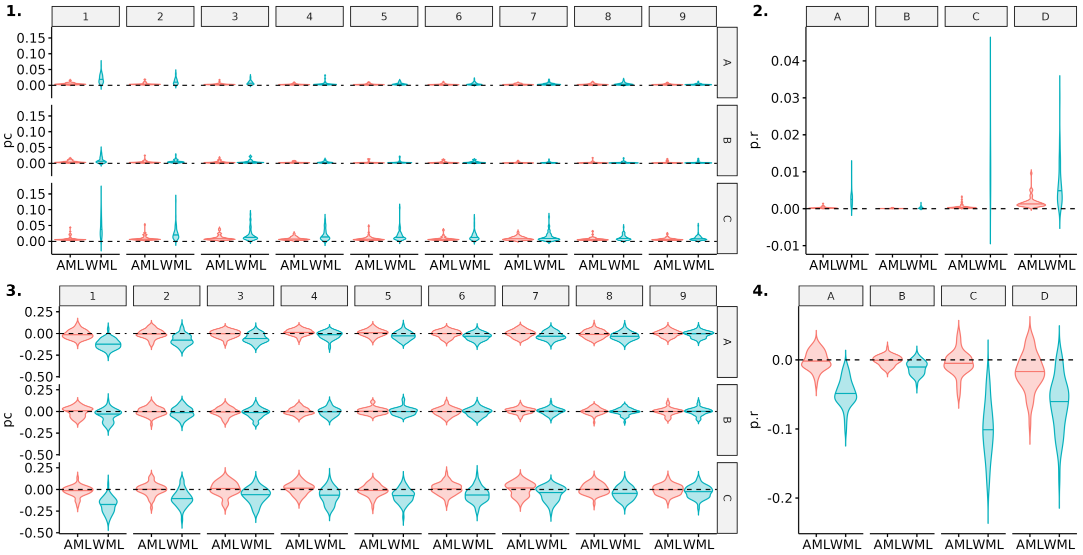

Figure S43: Comparison of precision and bias for estimates of capture ($pc$) and resigthing probabilities ($p.r$) between model accounting for mark loss (AML) or not (WML). Here, scenario 1 (long-lived species with high detection) is shown for a simulated mark loss probability of 0.4. Violin plots show the distributions of mean precision (1,2) and bias (3,4) over 50 simulations. The median of each distribution is shown with an horizontal line.

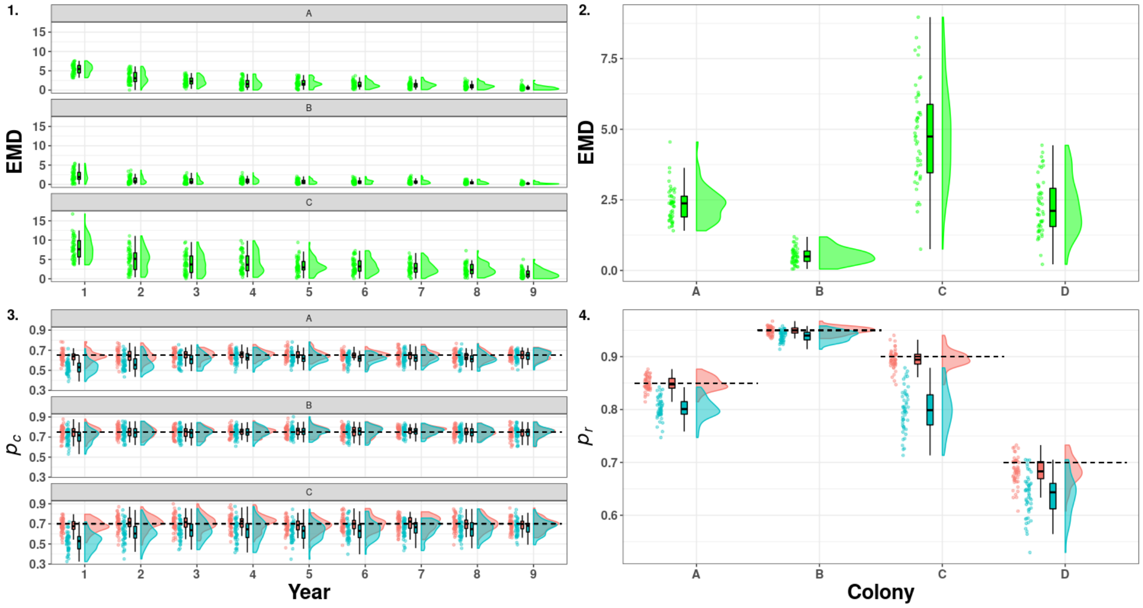

Figure S44: Raincloud plots of the Earth Mover Distance (green) between the medians of the posterior distribution of detection estimated for the model accounting for mark loss and recycling (red) and not accounting for mark loss (blue). On the left capture probability ($p_c$, 1 & 3), on the right resighting probability ($p_r$, 2 & 4) in scenario 1 (long-lived species with high detection). A, B, C, D denotes the name of states and the dashed line correspond to the mean of the true values.

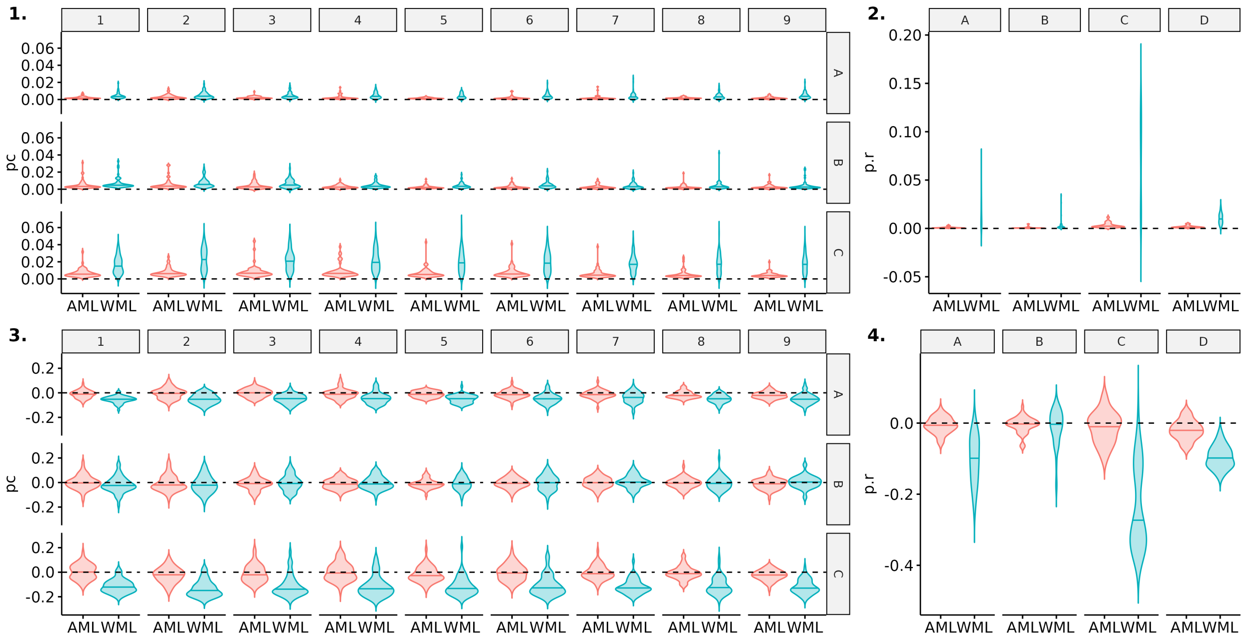

Figure S45: Comparison of precision and bias for estimates of capture (*pc*) and resigthing probabilities (*p.r*) between model accounting for mark loss (AML) or not (WML). Here, scenario 2 (long-lived species with low detection) is shown for a simulated mark loss probability of 0.4. Violin plots show the distributions of mean precision (1,2) and bias (3,4) over 50 simulations. The median of each distribution is shown with an horizontal line.

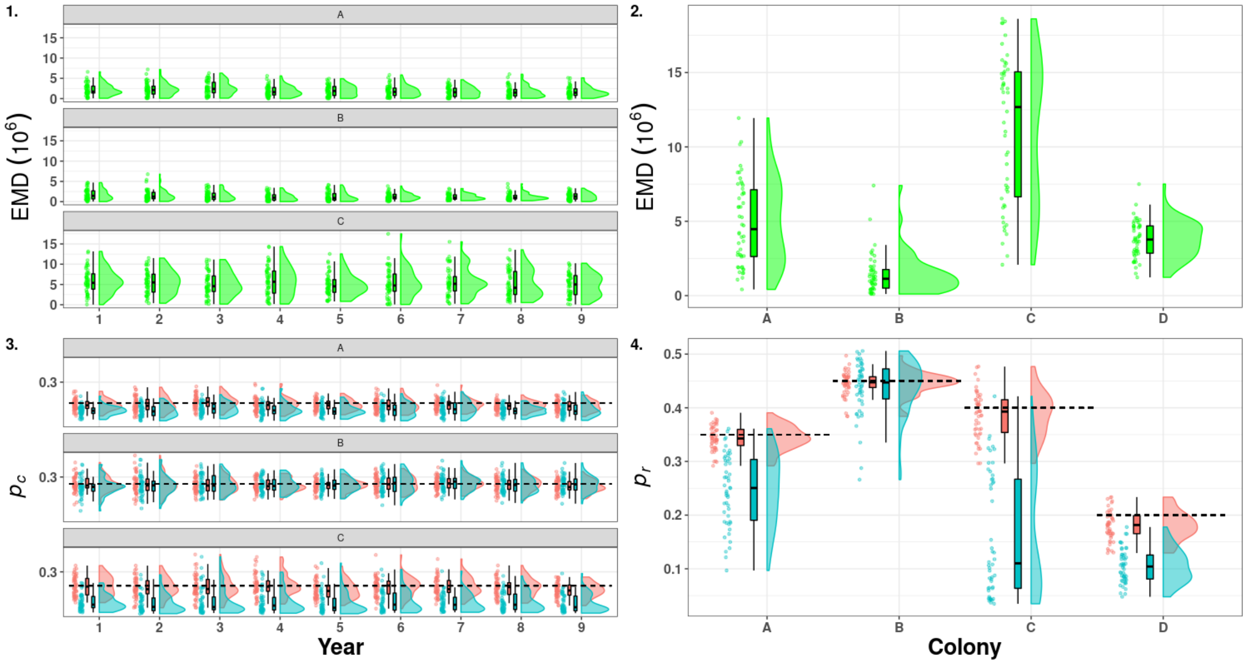

Figure S46: Raincloud plots of the Earth Mover Distance (green) between the medians of the posterior distribution of detection estimated for the model accounting for mark loss and recycling (red) and not accounting for mark loss (blue). On the left capture probability ($p_c$, 1 & 3), on the right resighting probability ($p_r$, 2 & 4) in scenario 2 (long-lived species with low detection). A, B, C, D denotes the name of states and the dashed line correspond to the mean of the true values.

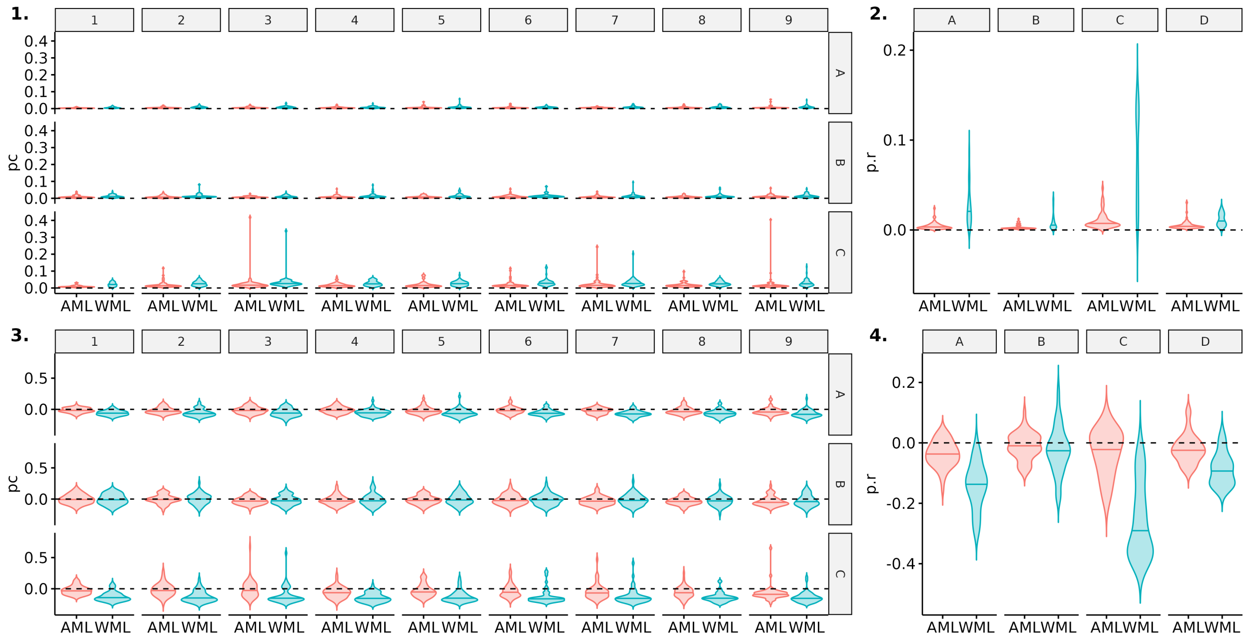

Figure S47: Comparison of precision and bias for estimates of capture ($pc$) and resigthing probabilities ($p.r$) between model accounting for mark loss (AML) or not (WML). Here, scenario 3 (short-lived species with low detection) is shown for a simulated mark loss probability of 0.4. Violin plots show the distributions of mean precision (1,2) and bias (3,4) over 50 simulations. The median of each distribution is shown with an horizontal line.

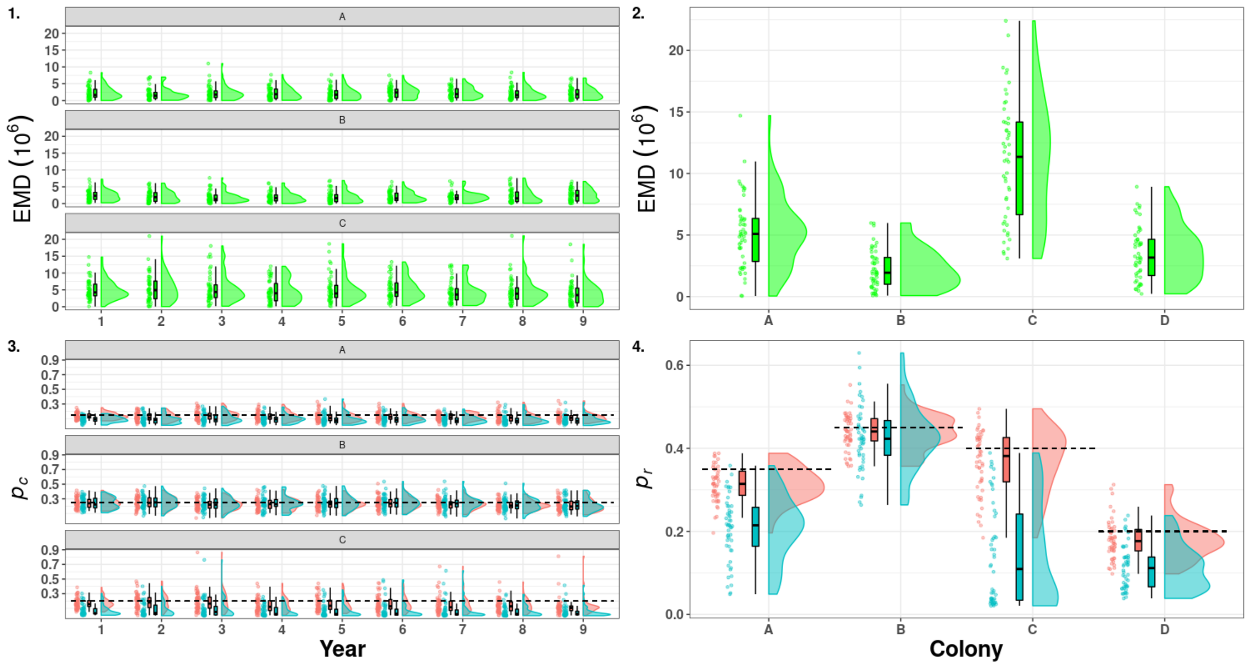

Figure S48: Raincloud plots of the Earth Mover Distance (green) between the medians of the posterior distribution of detection estimated for the model accounting for mark loss and recycling (red) and not accounting for mark loss (blue). On the left capture probability ($p_c$, 1 & 3), on the right resighting probability ($p_r$, 2 & 4) in scenario 3 (short-lived species with low detection). A, B, C, D denotes the name of states and the dashed line correspond to the mean of the true values.

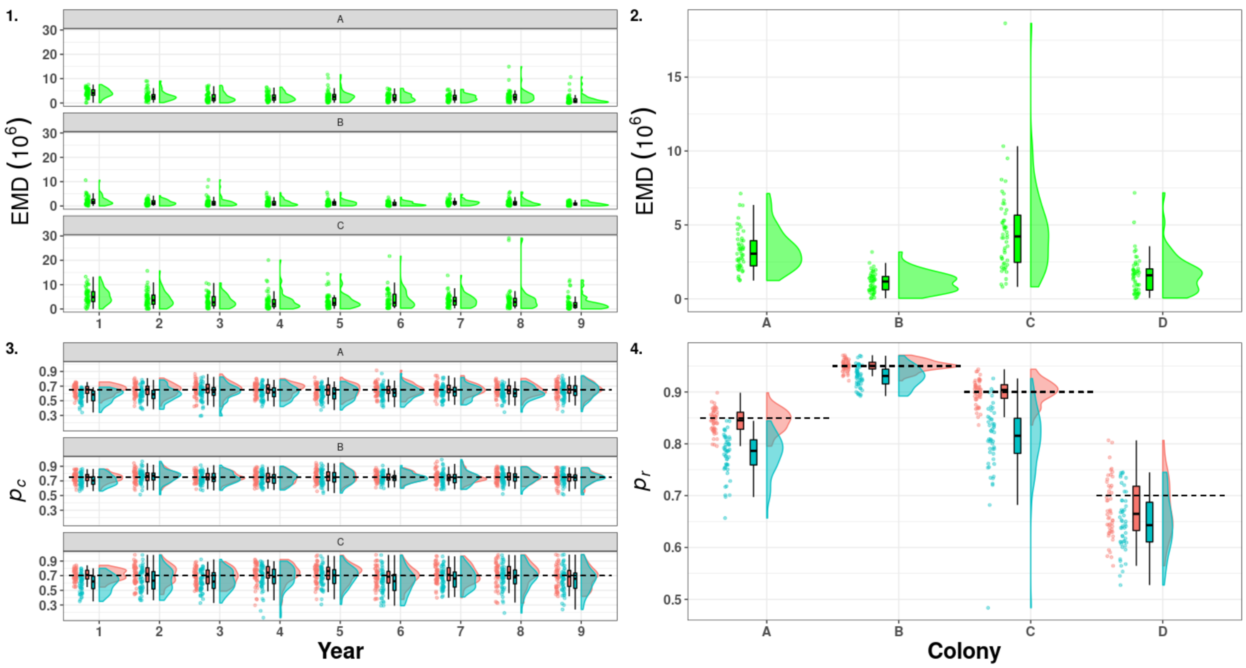

Figure S49: Raincloud plots of the Earth Mover Distance (green) between the medians of the posterior distribution of detection estimated for the model accounting for mark loss and recycling (red) and not accounting for mark loss (blue). On the left capture probability ($p_c$, 1 & 3), on the right resighting probability ($p_r$, 2 & 4) in scenario 4 (short-lived species with high detection). A, B, C, D denotes the name of states and the dashed line correspond to the mean of the true values.

#### 3.5 State transition probability estimates

Below, we compared the bias (median - truth) and the precision (mean squared errors MSE = bias$^2$+ variance) of the state transition probabilities between the model without recycling (red distribution) and with recycling (blue distribution). We also use the Earth Mover Distance (EMD) to compare the distribution of the medians of these parameters (green distribution). The density distribution of these medians were also displayed. In our simulation study, we constrained females to change only between state A, B, C, and male from initial state (either A, B, C) to D when they were juveniles, with fixed rates during the entire study period. Males no longer change state (A, B, C or D) when they reach one year of age.

##### 3.5.1 Simulations with mark loss rate of 0.05

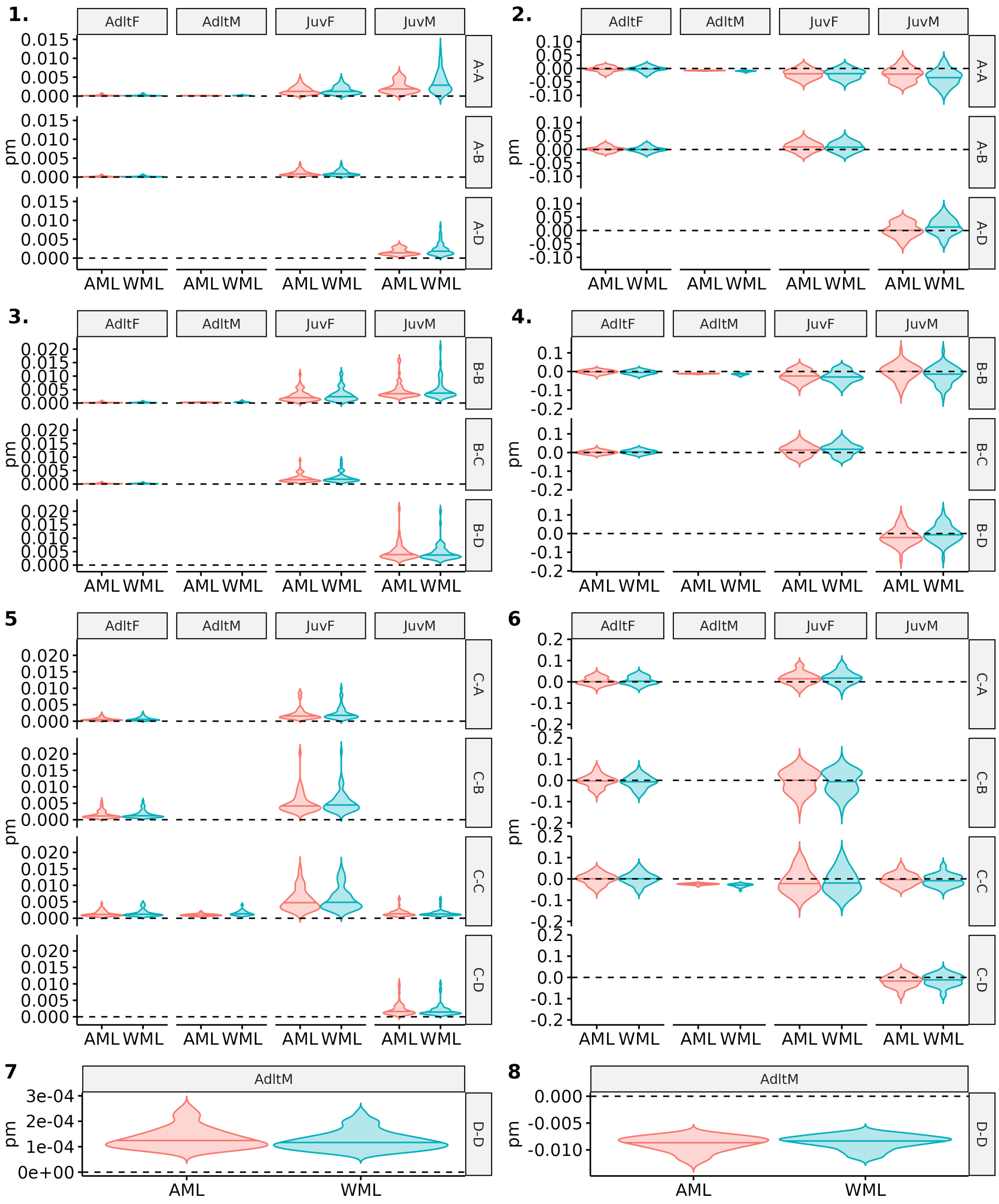

Figure S50: Comparison of precision and bias for estimates of state transition probabilities between model accounting for mark loss (AML) or not (WML). Scenario 1, long-lived and high detection, with a simulated probability of mark loss of 0.05 is shown. Age (Adlt=adult, Juv=juvenile) and sex (F=female, M=male) are specified at the top of each plot and transition "from-to" on the right side, with A, B, C, D the name of the simulated states. Violin plots show the distributions of mean precision (1,3,5,7) and bias (2,4,6,8) over 50 simulations. The median of each distribution is shown with an horizontal line.

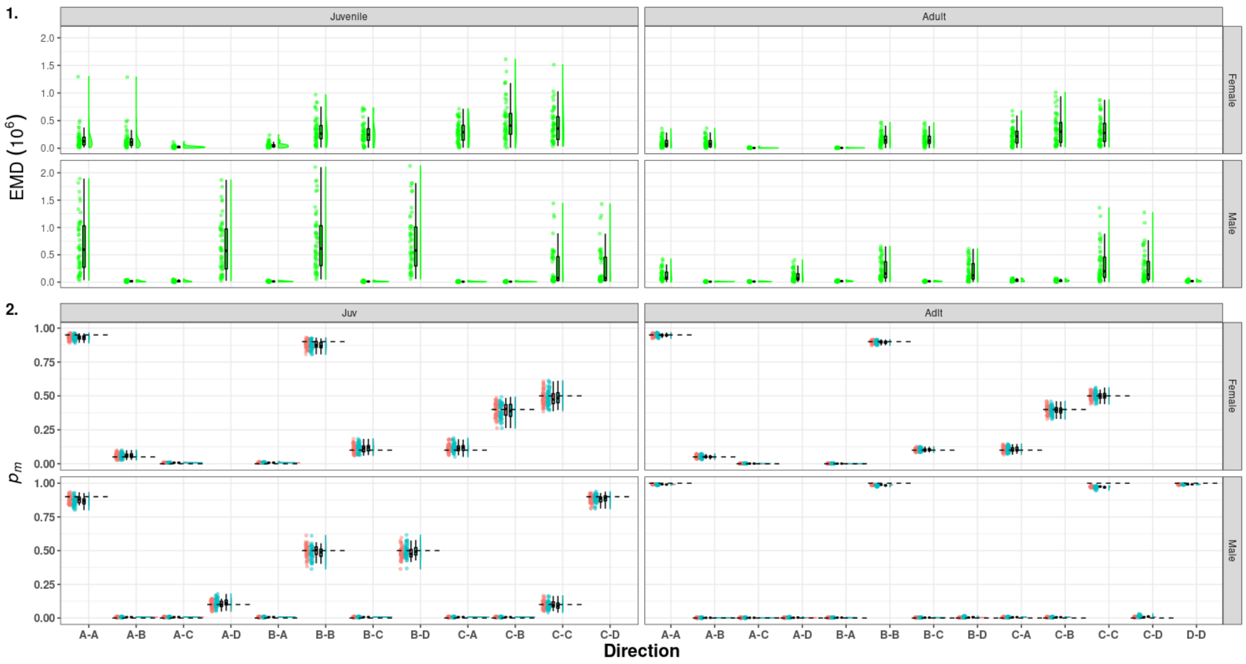

Figure S51: Raincloud plots of the Earth Mover Distance (green) between the medians of the posterior distribution of detection estimated for the model accounting for mark loss and recycling (red) and not accounting for mark loss (blue). On the left capture probability ($p_c$, 1 & 3), on the right resighting probability ($p_r$, 2 & 4) in scenario 1 (long-lived species with high detection). A, B, C, D denotes the name of states and the dashed line correspond to the mean of the true values.

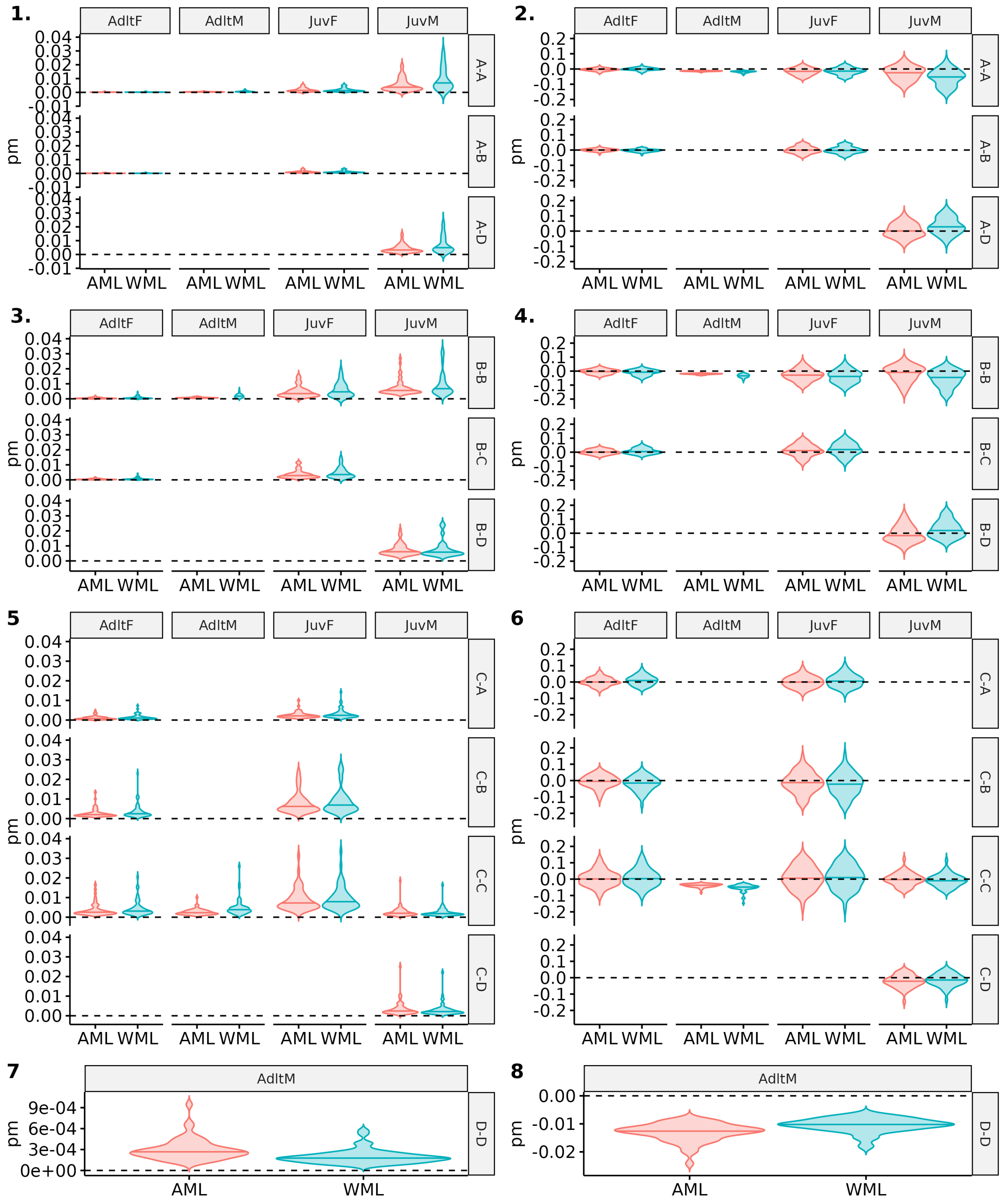

Figure S52: Comparison of precision and bias for estimates of state transition probabilities between model accounting for mark loss (AML) or not (WML). Scenario 2, long-lived and low detection, with a simulated probability of mark loss of 0.05 is shown. Age (Adlt=adult, Juv=juvenile) and sex (F=female, M=male) are specified at the top of each plot and transition "from-to" on the right side, with A, B, C, D the name of the simulated states. Violin plots show the distributions of mean precision (1,3,5,7) and bias (2,4,6,8) over 50 simulations. The median of each distribution is shown with an horizontal line.

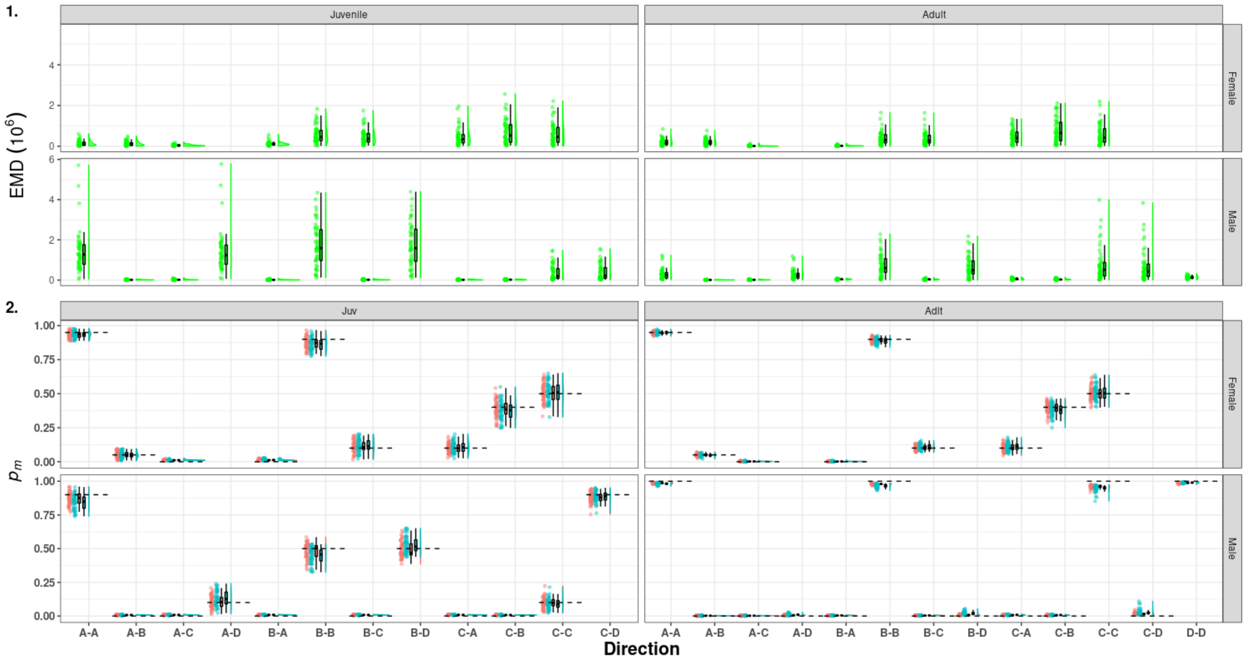

Figure S53: Raincloud plots of the Earth Mover Distance (green) between the medians of the posterior distribution of detection estimated for the model accounting for mark loss and recycling (red) and not accounting for mark loss (blue). On the left capture probability ($p_c$, 1 & 3), on the right resighting probability ($p_r$, 2 & 4) in scenario 2 (short-lived species with high detection). A, B, C, D denotes the name of states and the dashed line correspond to the mean of the true values.

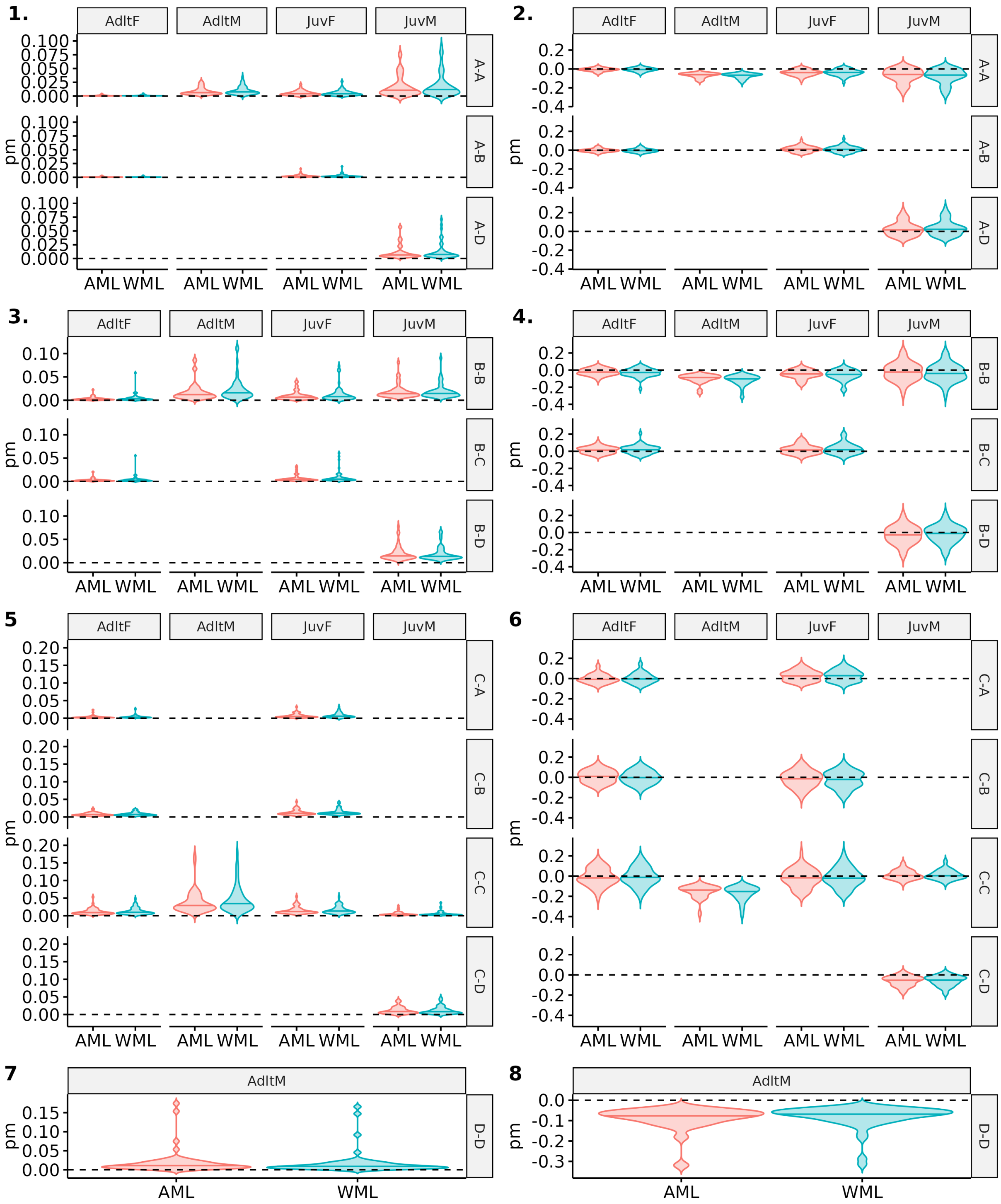

Figure S54: Comparison of precision and bias for estimates of state transition probabilities between model accounting for mark loss (AML) or not (WML). Scenario 3, short-lived and low detection, with a simulated probability of mark loss of 0.05 is shown. Age (Adlt=adult, Juv=juvenile) and sex (F=female, M=male) are specified at the top of each plot and transition "from-to" on the right side, with A, B, C, D the name of the simulated states. Violin plots show the distributions of mean precision (1,3,5,7) and bias (2,4,6,8) over 50 simulations. The median of each distribution is shown with an horizontal line.

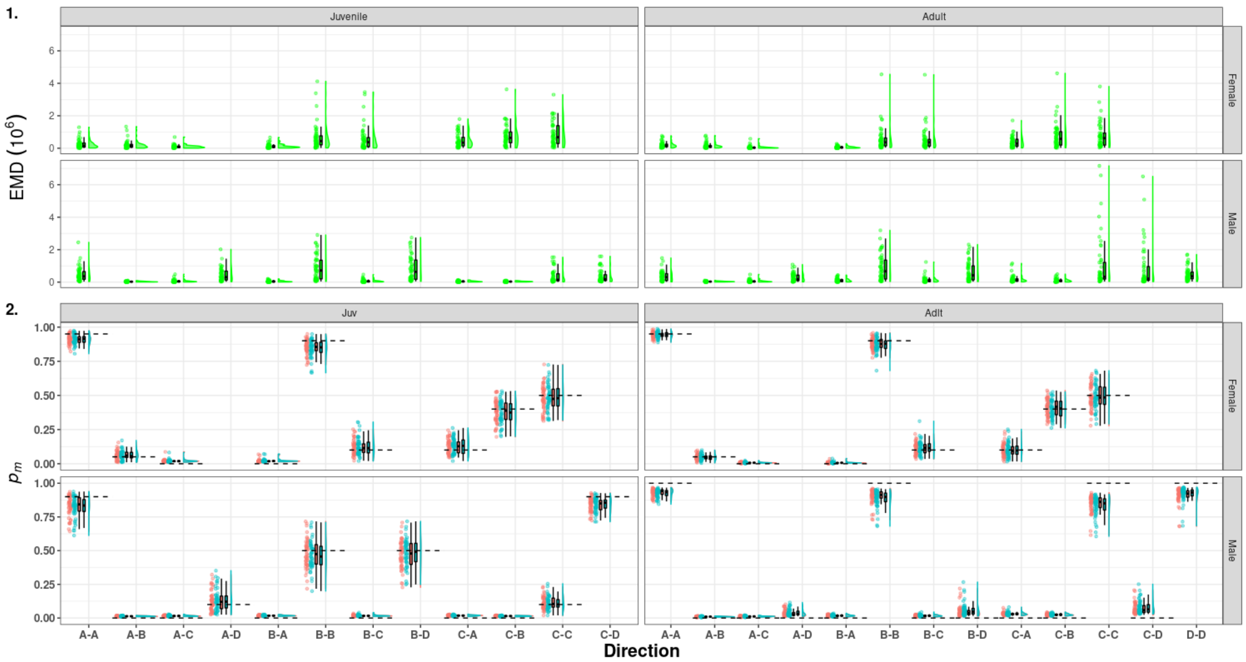

Figure S55: Raincloud plots of the Earth Mover Distance (green) between the medians of the posterior distribution of detection estimated for the model accounting for mark loss and recycling (red) and not accounting for mark loss (blue). On the left capture probability ($p_c$, 1 & 3), on the right resighting probability ($p_r$, 2 & 4) in scenario 3 (short-lived species with low detection). A, B, C, D denotes the name of states and the dashed line correspond to the mean of the true values.

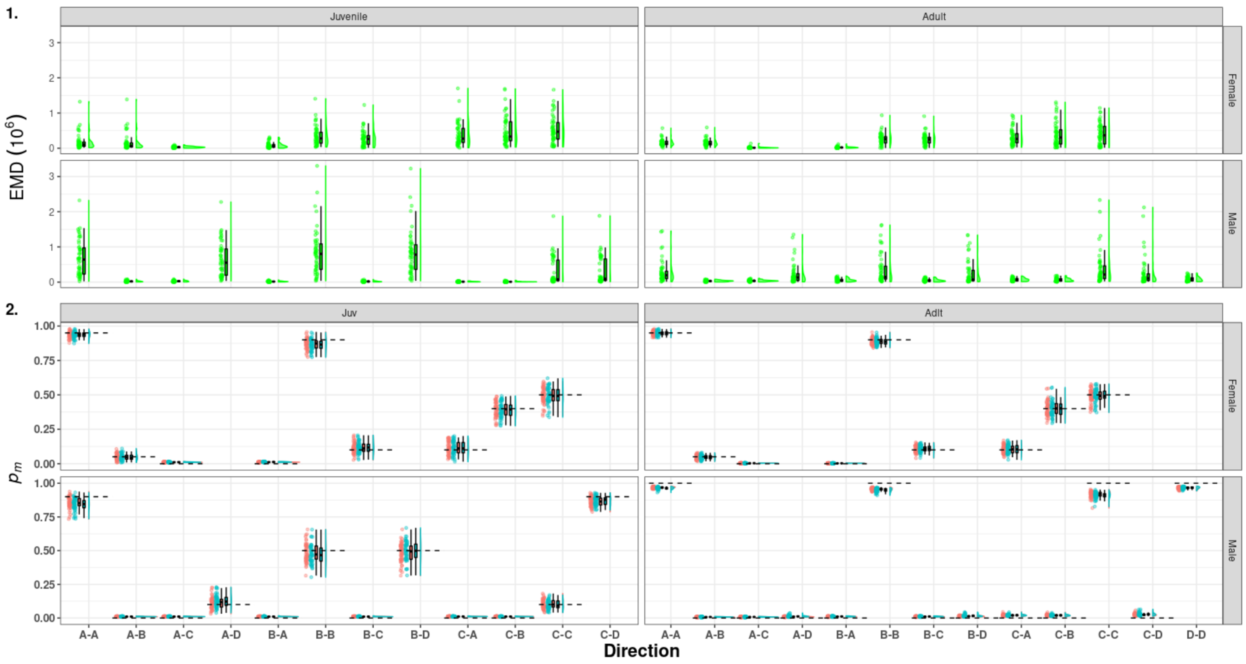

Figure S56: Raincloud plots of the Earth Mover Distance (green) between the medians of the posterior distribution of detection estimated for the model accounting for mark loss and recycling (red) and not accounting for mark loss (blue). On the left capture probability ($p_c$, 1 & 3), on the right resighting probability ($p_r$, 2 & 4) in scenario 4 (long-lived species with low detection). A, B, C, D denotes the name of states and the dashed line correspond to the mean of the true values.

##### 3.5.2 Simulations with mark loss rate of 0.25

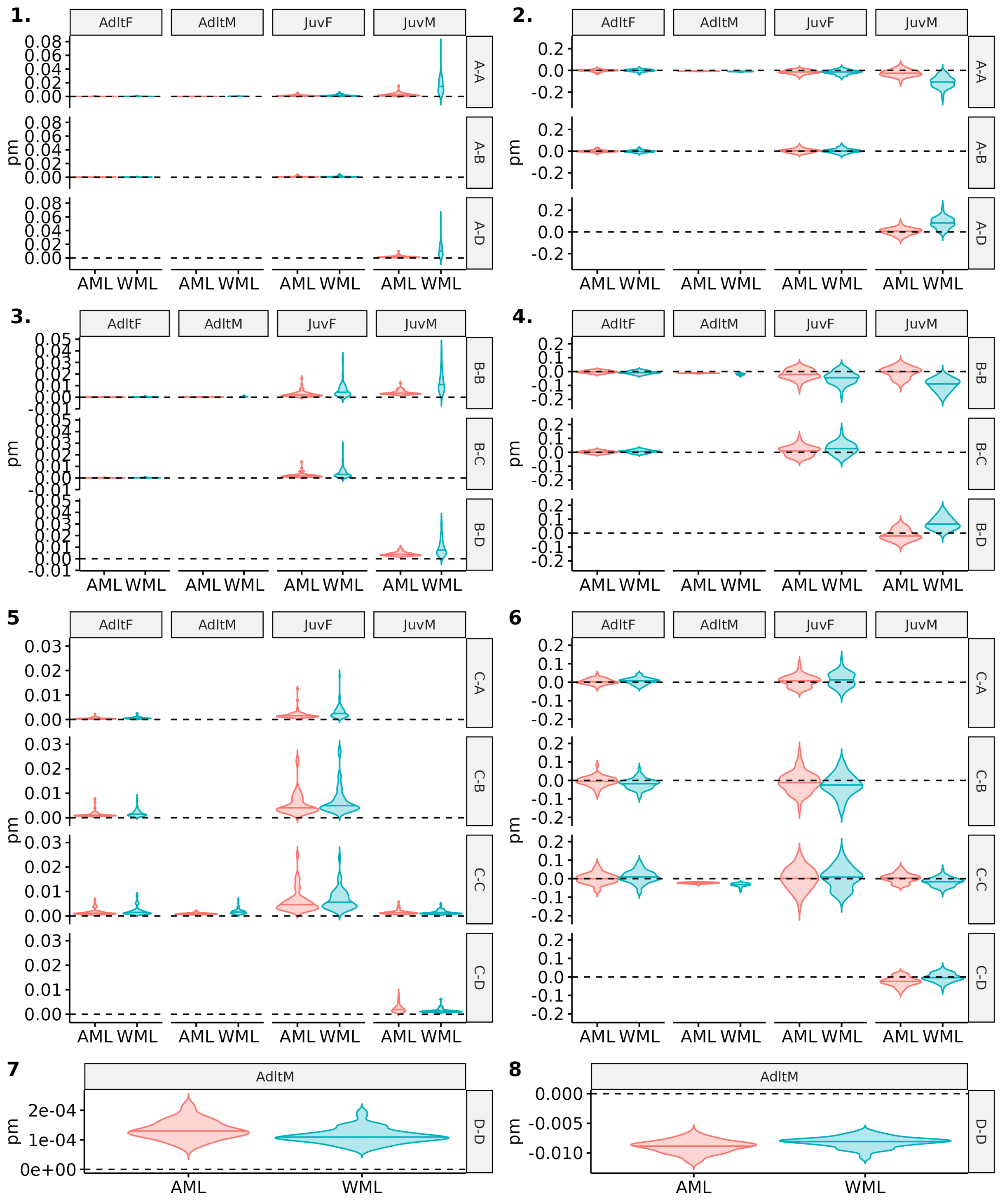

Figure S57: Comparison of precision and bias for estimates of state transition probabilities between model accounting for mark loss (AML) or not (WML). Scenario 1, long-lived and high detection, with a simulated probability of mark loss of 0.25 is shown. Age (Adlt=adult, Juv=juvenile) and sex (F=female, M=male) are specified at the top of each plot and transition "from-to" on the right side, with A, B, C, D the name of the simulated states. Violin plots show the distributions of mean precision (1,3,5,7) and bias (2,4,6,8) over 50 simulations. The median of each distribution is shown with an horizontal line.

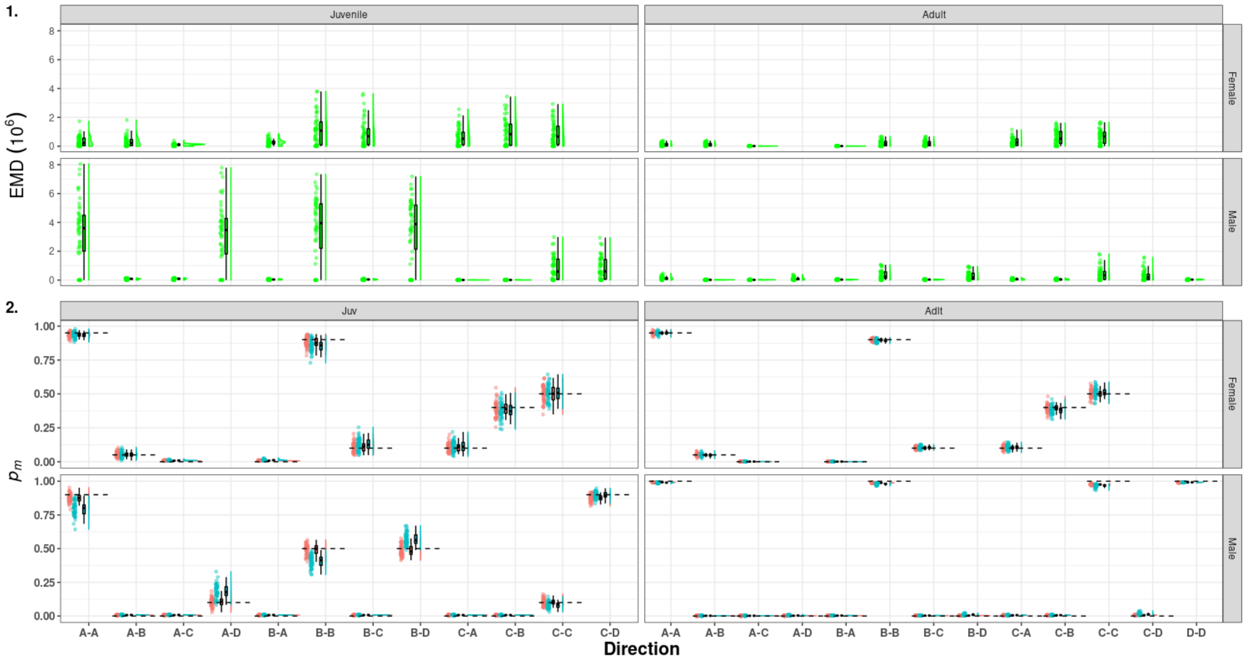

Figure S58: (1) Raincloud plots of the Earth Mover Distance (green) between the medians of the posterior distribution of detection estimated for the model accounting for mark loss and recycling (red) and not accounting for mark loss (blue). On the left capture probability ($p_c$, 1 & 3), on the right resighting probability ($p_r$, 2 & 4) in scenario 1 (long-lived species with high detection). A, B, C, D denotes the name of states and the dashed line correspond to the mean of the true values.

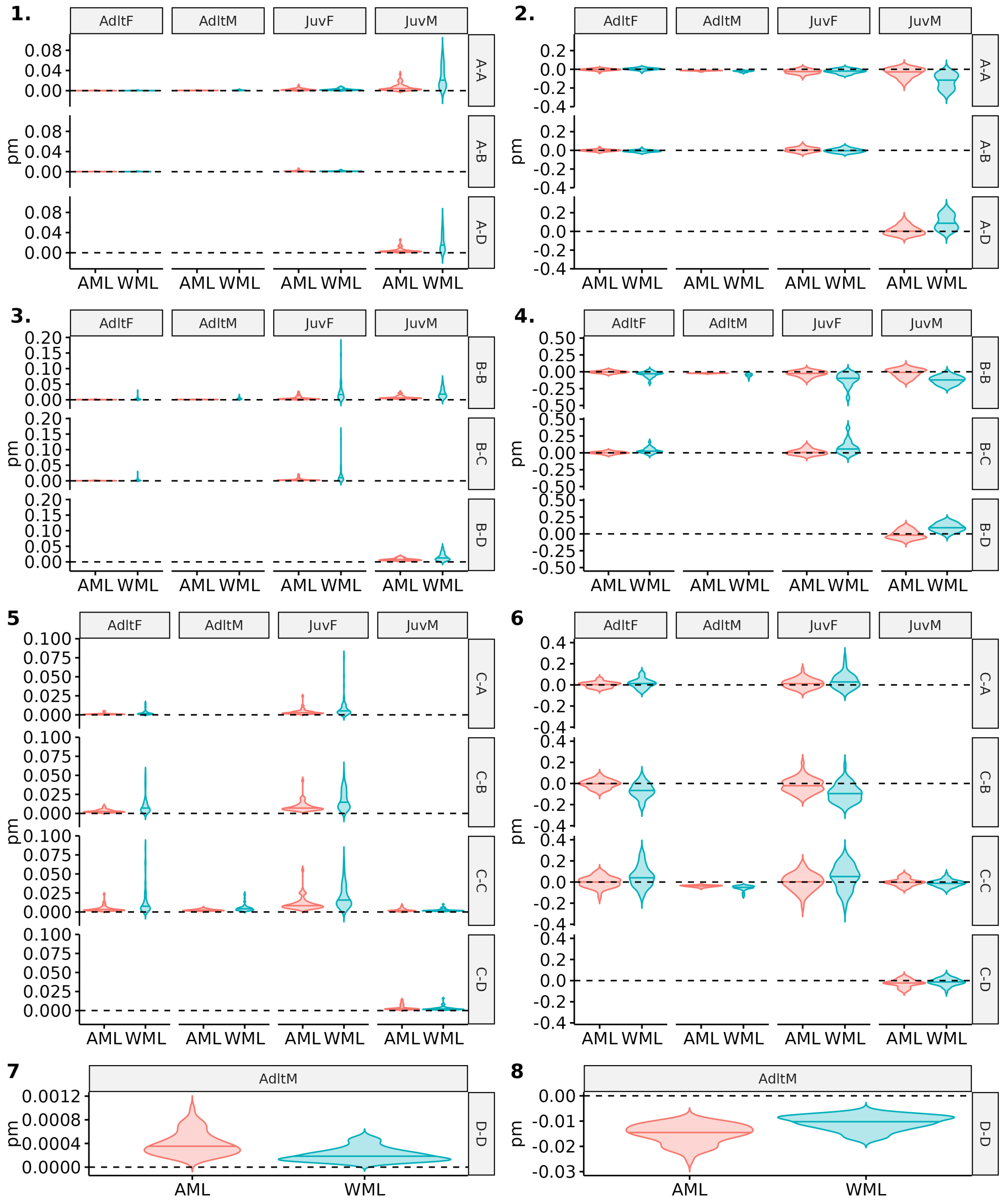

Figure S59: Comparison of precision and bias for estimates of state transition probabilities between model accounting for mark loss (AML) or not (WML). Scenario 2, long-lived and low detection, with a simulated probability of mark loss of 0.25 is shown. Age (Adlt=adult, Juv=juvenile) and sex (F=female, M=male) are specified at the top of each plot and transition "from-to" on the right side, with A, B, C, D the name of the simulated states. Violin plots show the distributions of mean precision (1,3,5,7) and bias (2,4,6,8) over 50 simulations. The median of each distribution is shown with an horizontal line.

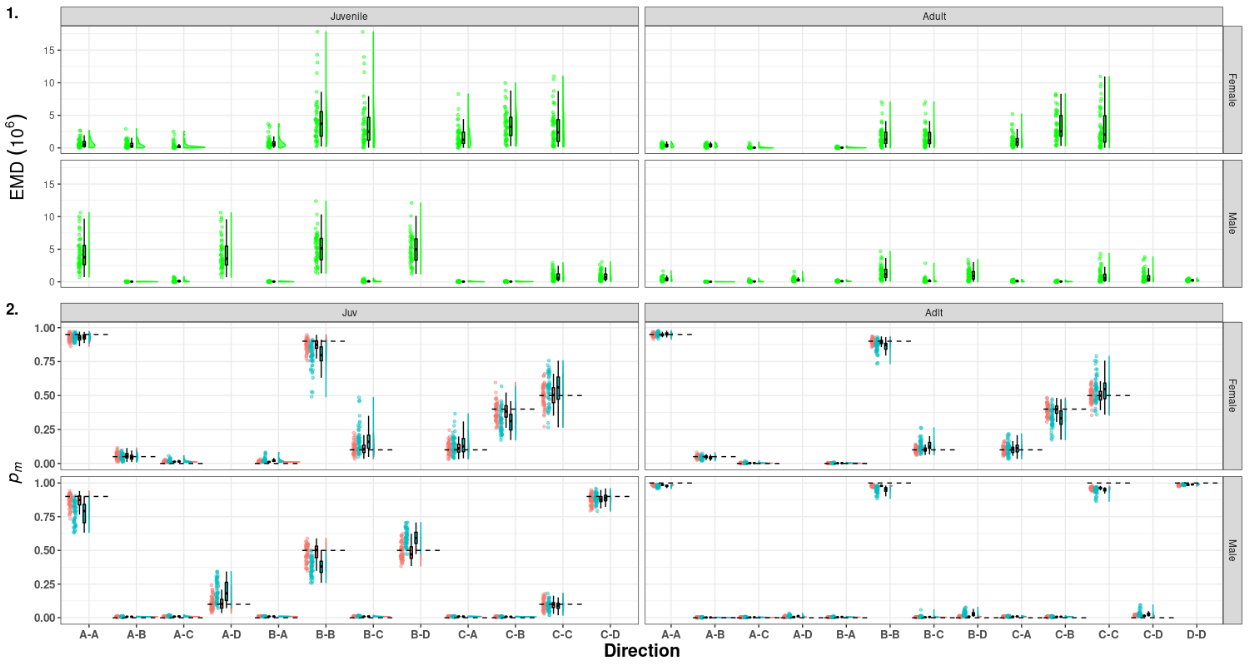

Figure S60: Raincloud plots of the Earth Mover Distance (green) between the medians of the posterior distribution of detection estimated for the model accounting for mark loss and recycling (red) and not accounting for mark loss (blue). On the left capture probability ($p_c$, 1 & 3), on the right resighting probability ($p_r$, 2 & 4) in scenario 2 (short-lived species with high detection). A, B, C, D denotes the name of states and the dashed line correspond to the mean of the true values.

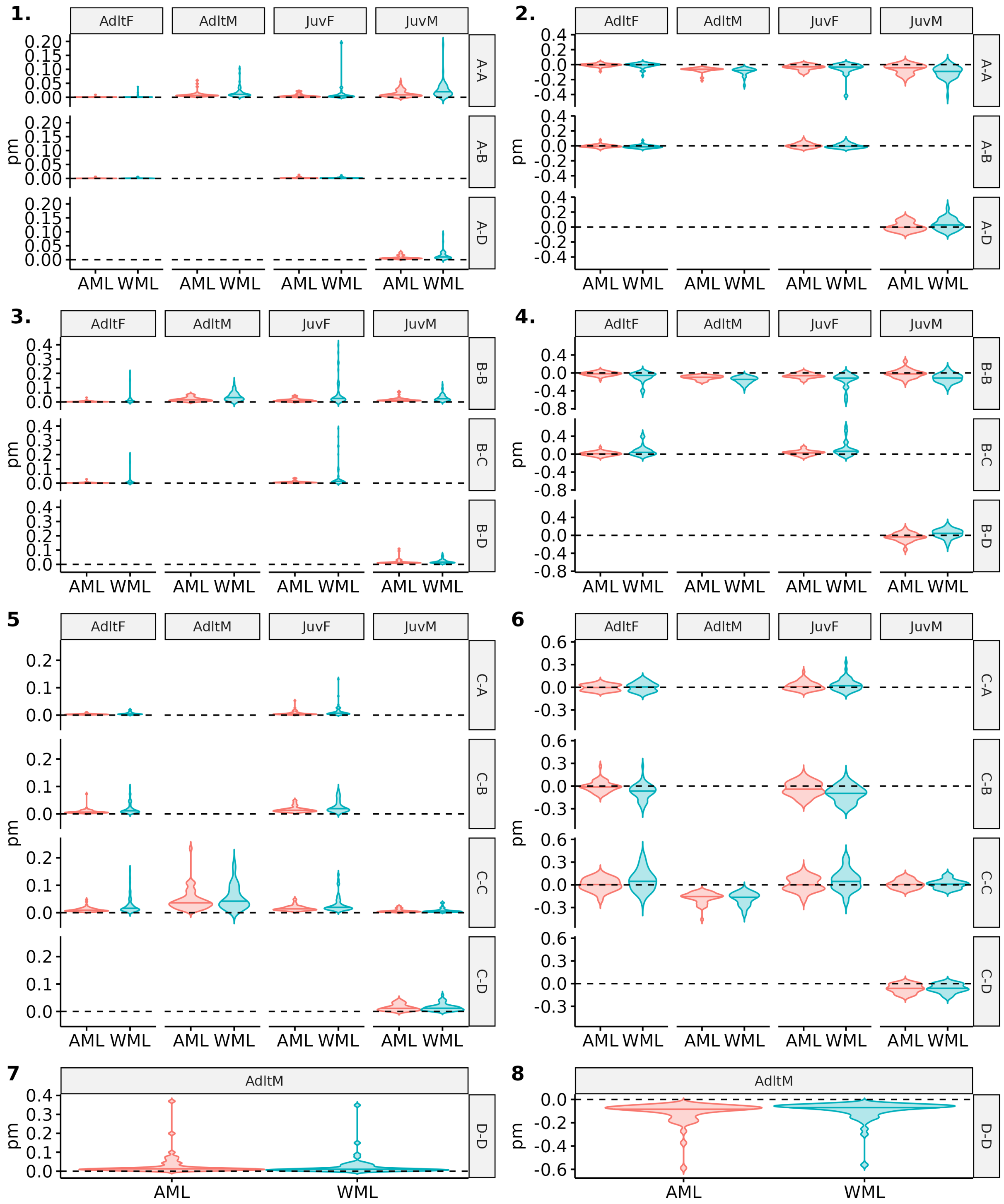

Figure S61: Comparison of precision and bias for estimates of state transition probabilities between model accounting for mark loss (AML) or not (WML). Scenario 3, short-lived and low detection, with a simulated probability of mark loss of 0.25 is shown. Age (Adlt=adult, Juv=juvenile) and sex (F=female, M=male) are specified at the top of each plot and transition "from-to" on the right side, with A, B, C, D the name of the simulated states. Violin plots show the distributions of mean precision (1,3,5,7) and bias (2,4,6,8) over 50 simulations. The median of each distribution is shown with an horizontal line.

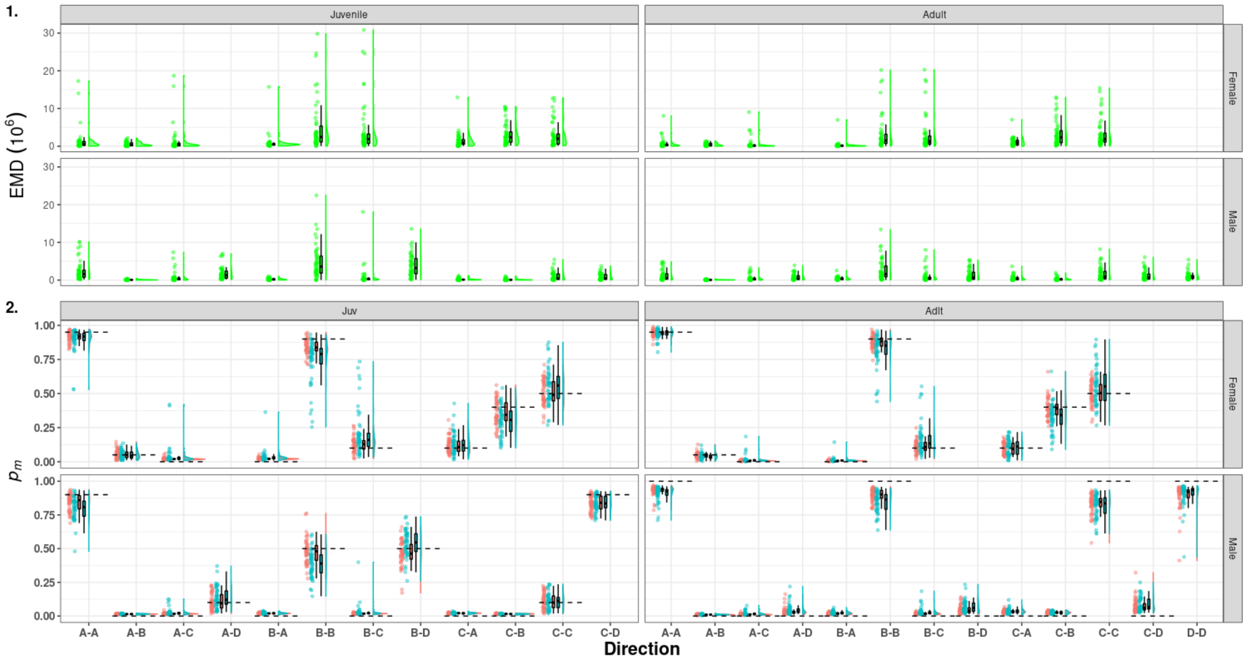

Figure S62: (1) Raincloud plots of the Earth Mover Distance (green) between the medians of the posterior distribution of detection estimated for the model accounting for mark loss and recycling (red) and not accounting for mark loss (blue). On the left capture probability ($p_c$, 1 & 3), on the right resighting probability ($p_r$, 2 & 4) in scenario 3 (short-lived species with low detection). A, B, C, D denotes the name of states and the dashed line correspond to the mean of the true values.

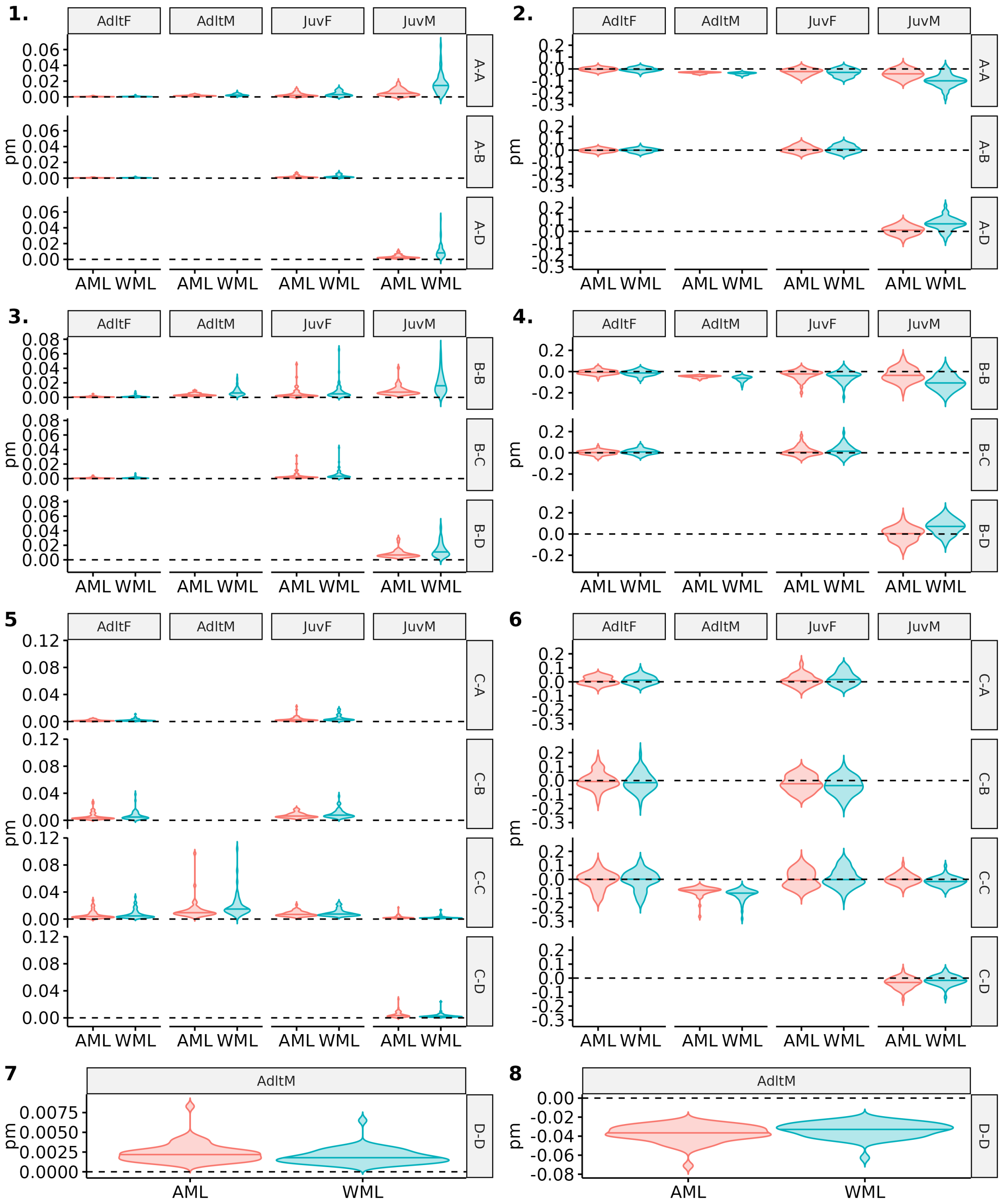

Figure S63: Comparison of precision and bias for estimates of state transition probabilities between model accounting for mark loss (AML) or not (WML). Scenario 4, short-lived and high detection, with a simulated probability of mark loss of 0.25 is shown. Age (Adlt=adult, Juv=juvenile) and sex (F=female, M=male) are specified at the top of each plot and transition "from-to" on the right side, with A, B, C, D the name of the simulated states. Violin plots show the distributions of mean precision (1,3,5,7) and bias (2,4,6,8) over 50 simulations. The median of each distribution is shown with an horizontal line.

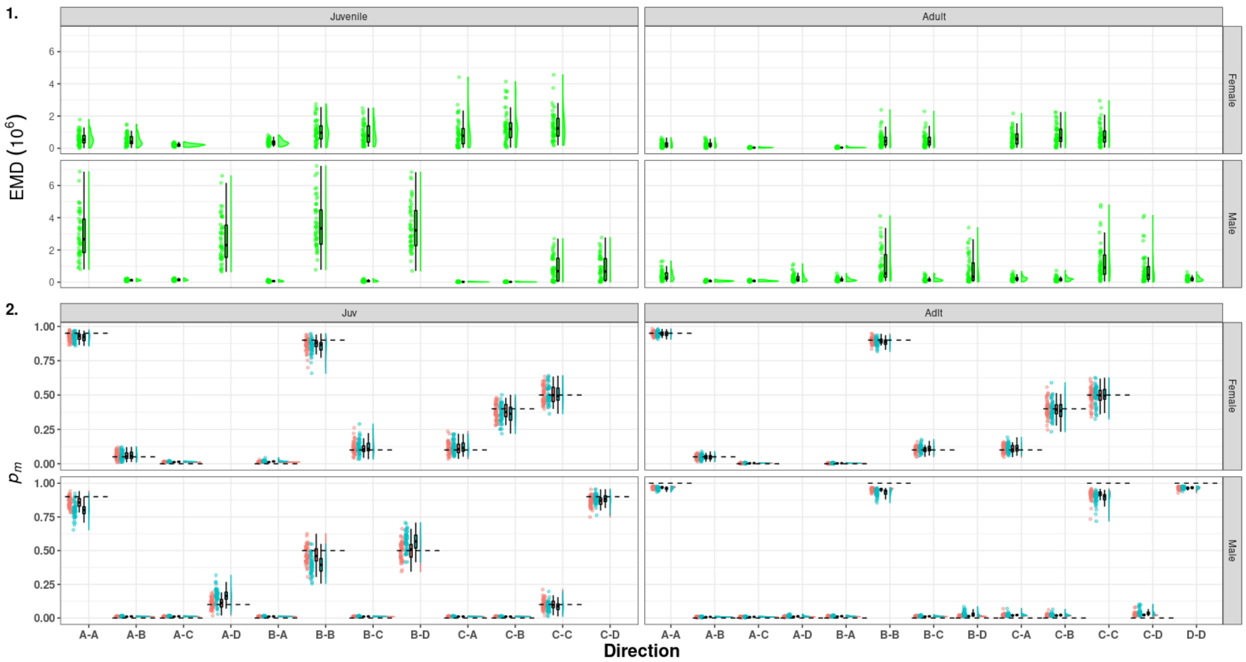

Figure S64: Raincloud plots of the Earth Mover Distance (green) between the medians of the posterior distribution of detection estimated for the model accounting for mark loss and recycling (red) and not accounting for mark loss (blue). On the left capture probability ($p_c$, 1 & 3), on the right resighting probability ($p_r$, 2 & 4) in scenario 4 (long-lived species with low detection). A, B, C, D denotes the name of states and the dashed line correspond to the mean of the true values.

##### 3.5.3 Simulations with mark loss rate of 0.4

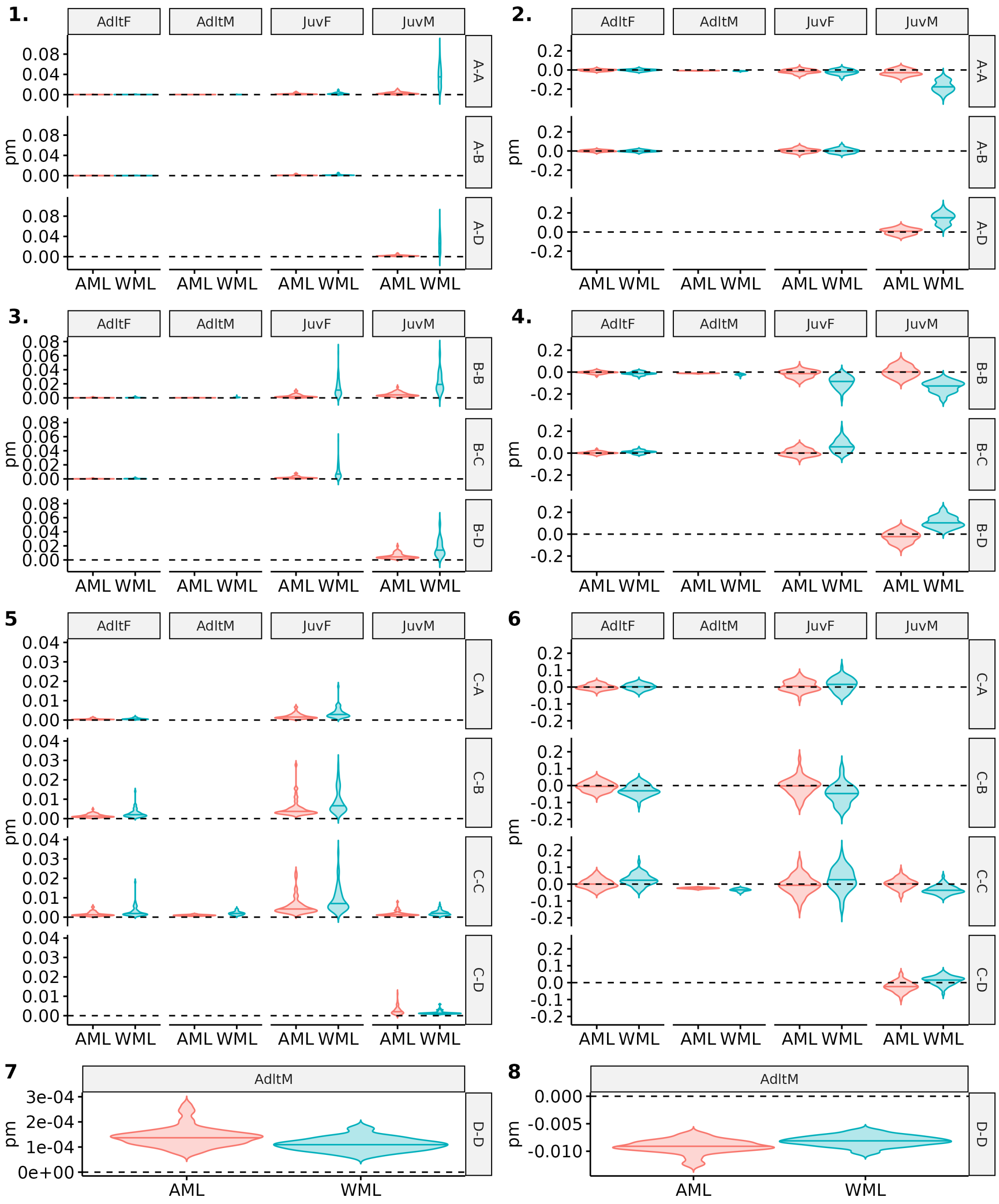

Figure S65: Comparison of precision and bias for estimates of state transition probabilities between model accounting for mark loss (AML) or not (WML). Scenario 1, long-lived and high detection, with a simulated probability of mark loss of 0.4 is shown. Age (Adlt=adult, Juv=juvenile) and sex (F=female, M=male) are specified at the top of each plot and transition "from-to" on the right side, with A, B, C, D the name of the simulated states. Violin plots show the distributions of mean precision (1,3,5,7) and bias (2,4,6,8) over 50 simulations. The median of each distribution is shown with an horizontal line.

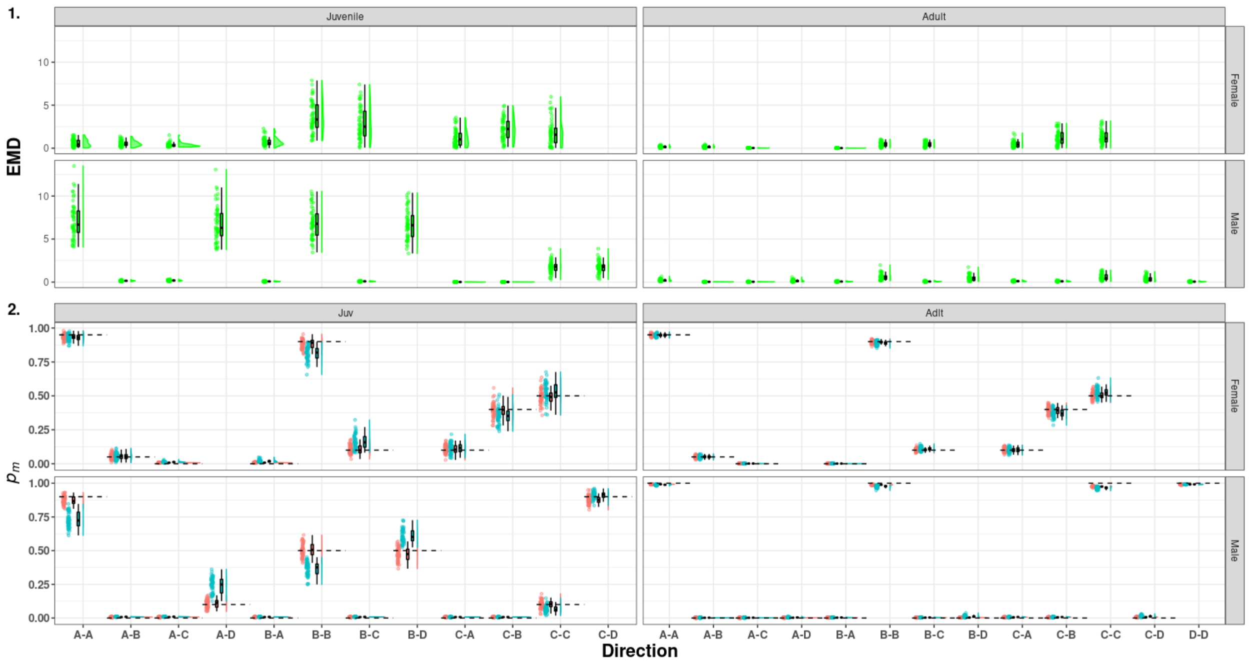

Figure S66: Raincloud plots of the Earth Mover Distance (green) between the medians of the posterior distribution of detection estimated for the model accounting for mark loss and recycling (red) and not accounting for mark loss (blue). On the left capture probability ($p_c$, 1 & 3), on the right resighting probability ($p_r$, 2 & 4) in scenario 1 (long-lived species with high detection). A, B, C, D denotes the name of states and the dashed line correspond to the mean of the true values.

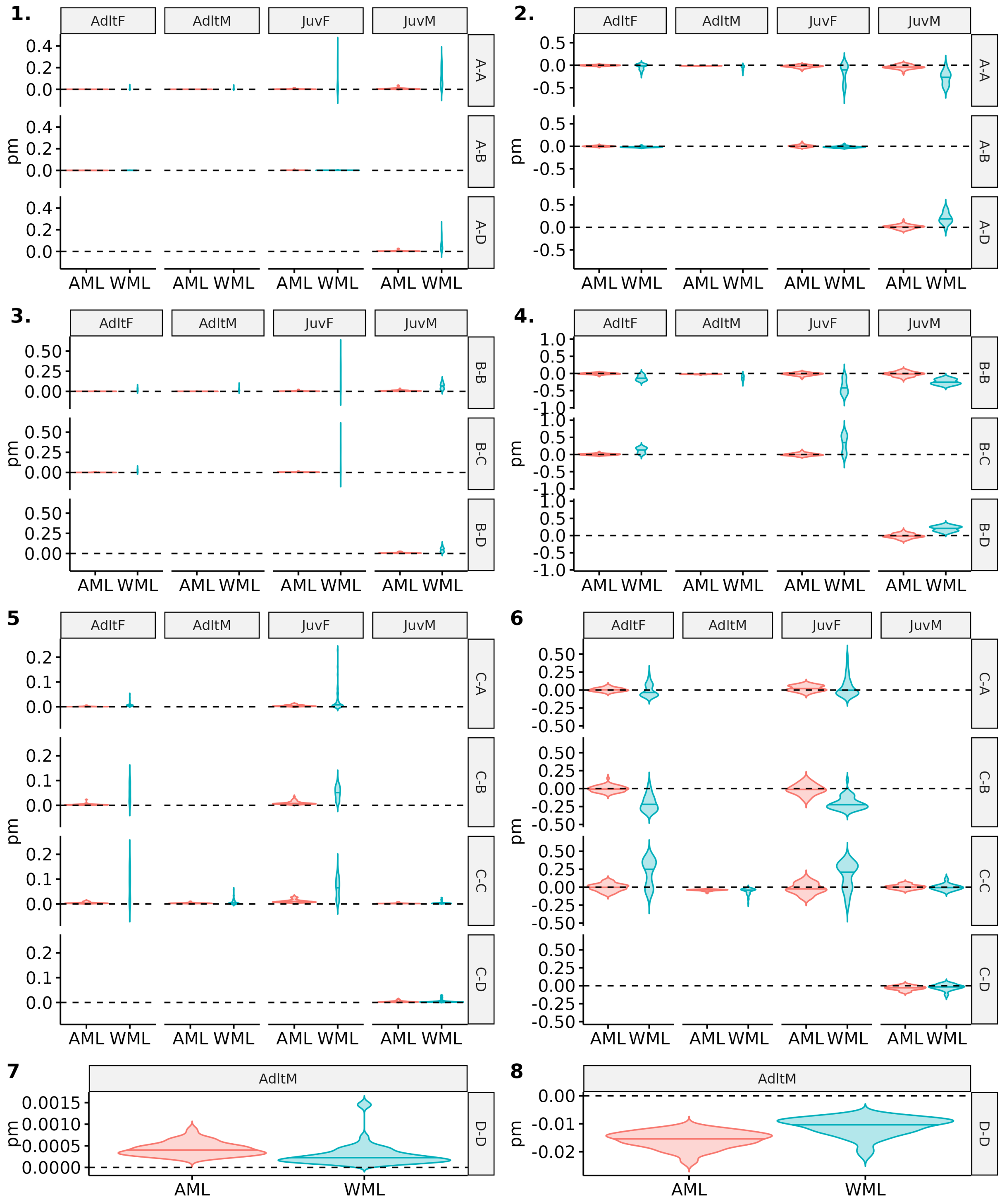

Figure S67: Comparison of precision and bias for estimates of state transition probabilities between model accounting for mark loss (AML) or not (WML). Scenario 2, long-lived and high detection, with a simulated probability of mark loss of 0.4 is shown. Age (Adlt=adult, Juv=juvenile) and sex (F=female, M=male) are specified at the top of each plot and transition "from-to" on the right side, with A, B, C, D the name of the simulated states. Violin plots show the distributions of mean precision (1,3,5,7) and bias (2,4,6,8) over 50 simulations. The median of each distribution is shown with an horizontal line.

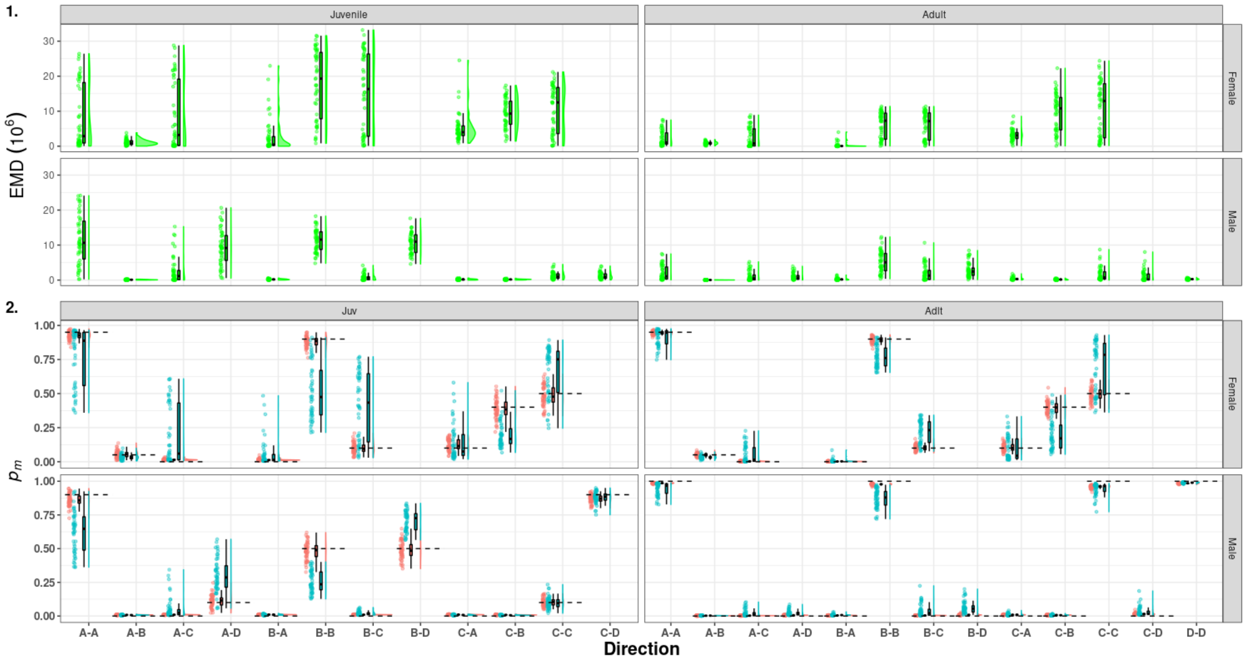

Figure S68: Raincloud plots of the Earth Mover Distance (green) between the medians of the posterior distribution of detection estimated for the model accounting for mark loss and recycling (red) and not accounting for mark loss (blue). On the left capture probability ($p_c$, 1 & 3), on the right resighting probability ($p_r$, 2 & 4) in scenario 2 (short-lived species with high detection). A, B, C, D denotes the name of states and the dashed line correspond to the mean of the true values.

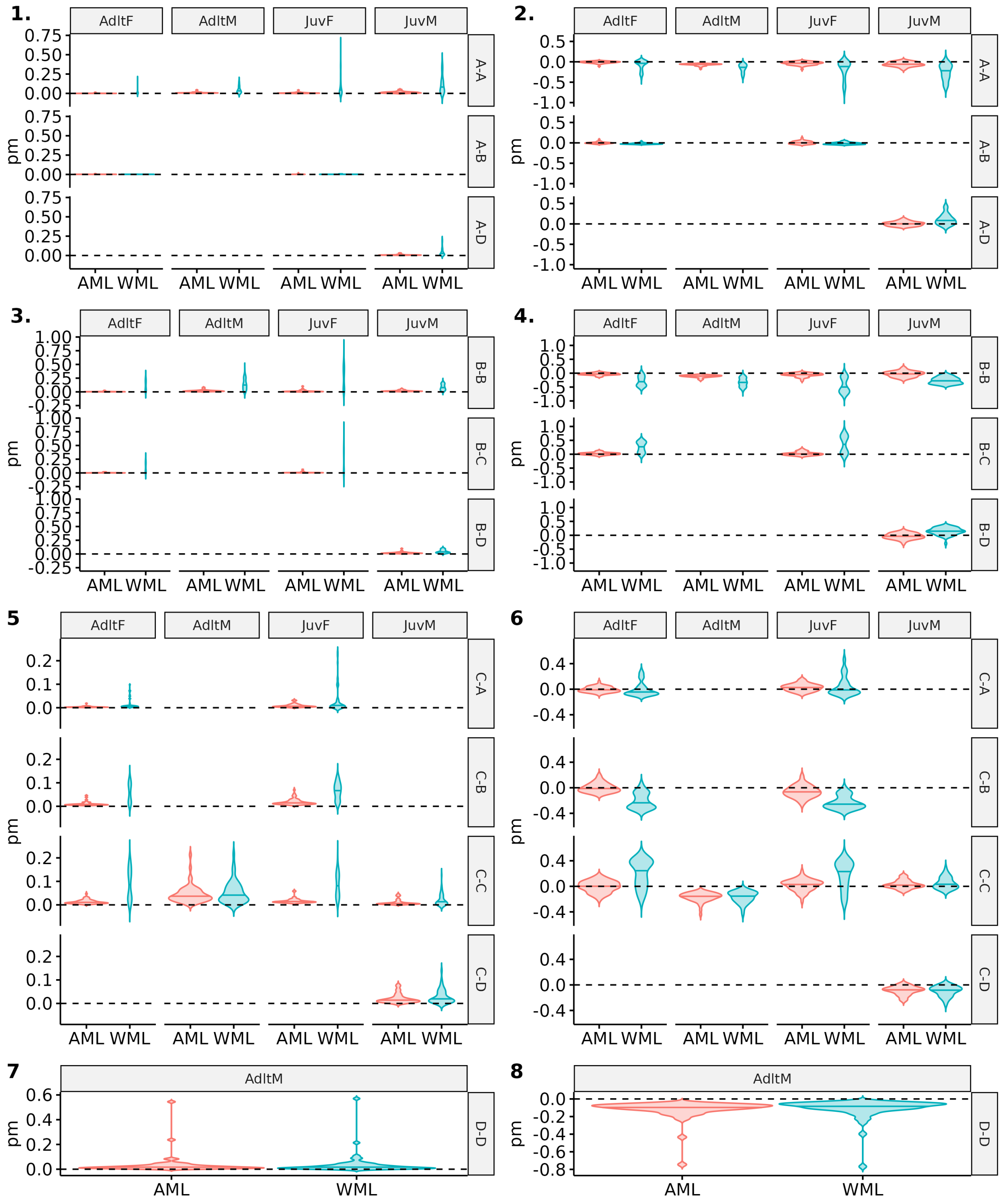

Figure S69: Comparison of precision and bias for estimates of state transition probabilities between model accounting for mark loss (AML) or not (WML). Scenario 3, short-lived and high detection, with a simulated probability of mark loss of 0.4 is shown. Age (Adlt=adult, Juv=juvenile) and sex (F=female, M=male) are specified at the top of each plot and transition "from-to" on the right side, with A, B, C, D the name of the simulated states. Violin plots show the distributions of mean precision (1,3,5,7) and bias (2,4,6,8) over 50 simulations. The median of each distribution is shown with an horizontal line.

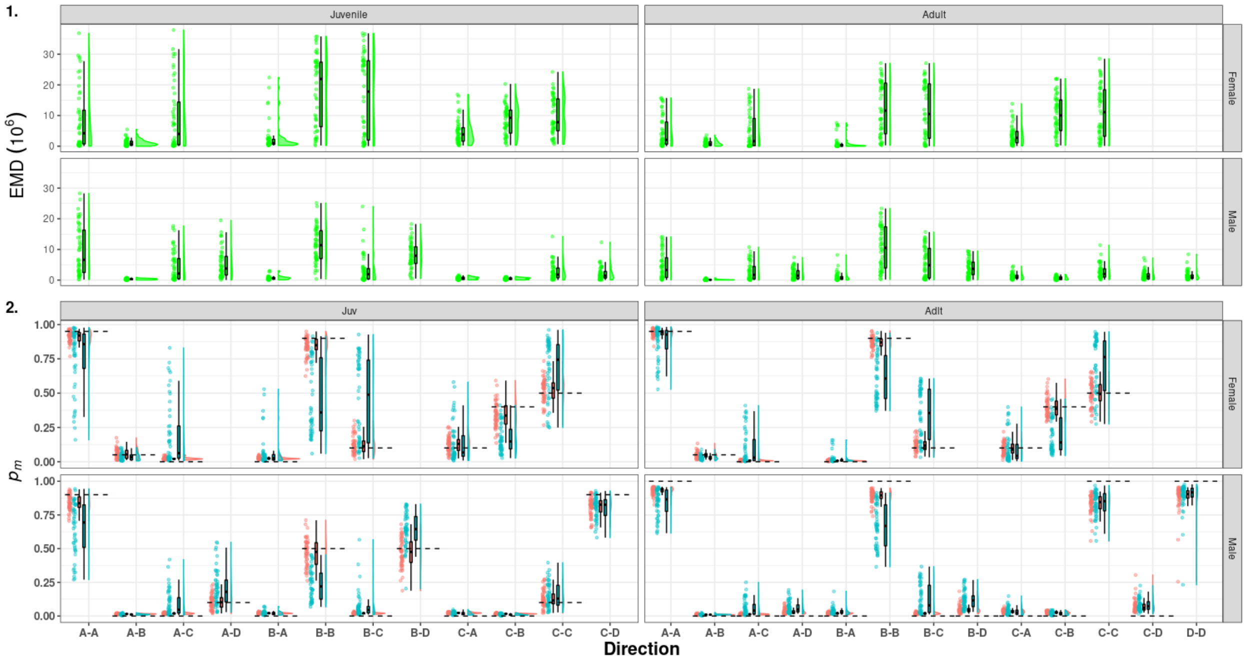

Figure S70: Raincloud plots of the Earth Mover Distance (green) between the medians of the posterior distribution of detection estimated for the model accounting for mark loss and recycling (red) and not accounting for mark loss (blue). On the left capture probability ($p_c$, 1 & 3), on the right resighting probability ($p_r$, 2 & 4) in scenario 3 (short-lived species with low detection). A, B, C, D denotes the name of states and the dashed line correspond to the mean of the true values.

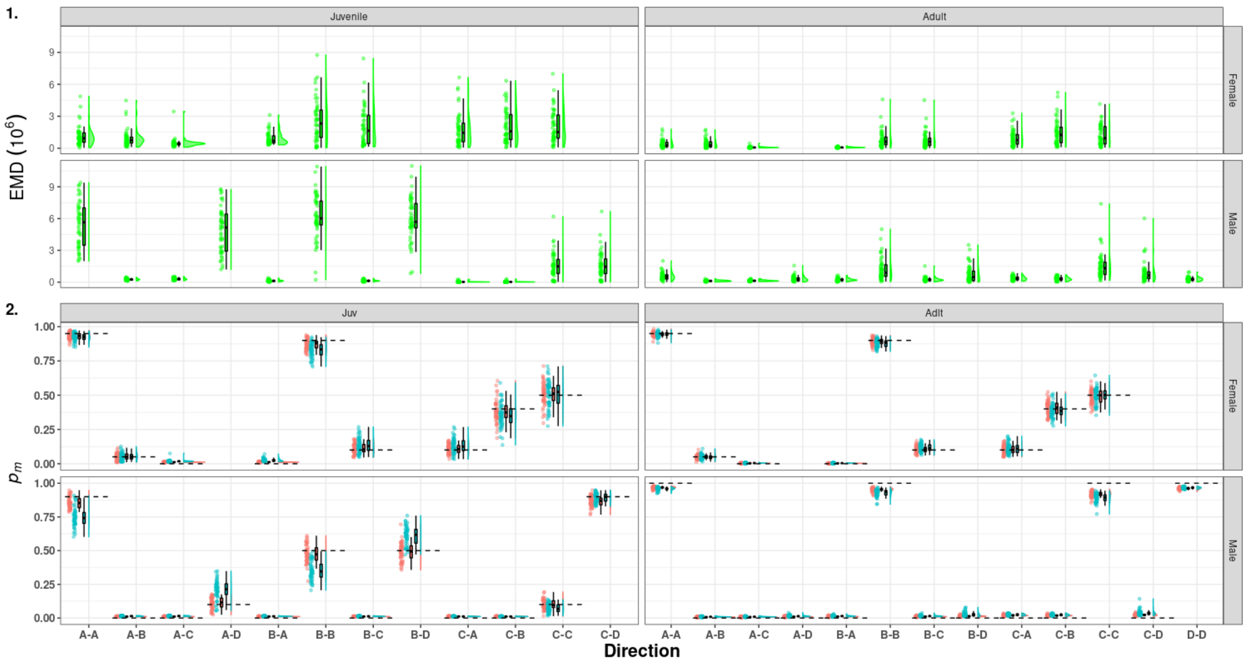

Figure S71: (1) Raincloud plots of the Earth Mover Distance (green) between the medians of the posterior distribution of detection estimated for the model accounting for mark loss and recycling (red) and not accounting for mark loss (blue). On the left capture probability ($p_c$, 1 & 3), on the right resighting probability ($p_r$, 2 & 4) in scenario 4 (long-lived species with low detection). A, B, C, D denotes the name of states and the dashed line correspond to the mean of the true values.

#### 3.6 Mark loss probability estimates

Mark loss rate was calculated with model considering tag loss and recycling (ModelA.jags). Here we displayed the distribution of the median of posterior distribution of the mark loss calculated from 50 simulations for each of the scenarios.

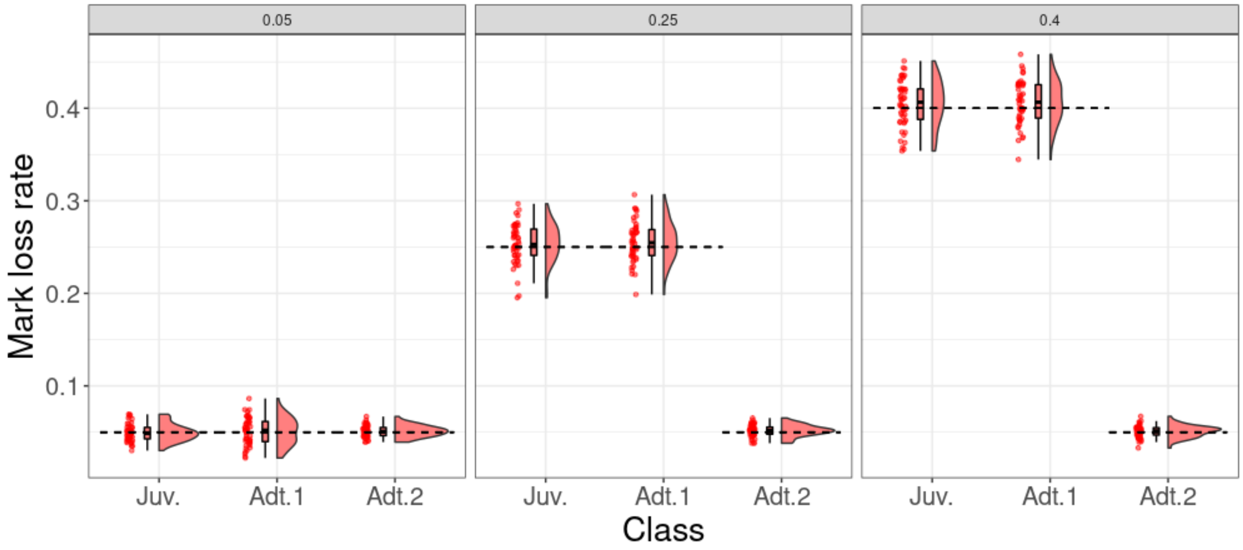

Figure S72: Probability distribution of the estimated mark loss rate for high survival and high detection (scenario 1). "Juv." and "Adt.1" indicate the mark loss rate during the year following marking for juveniles and adults respectively. "Adt.2" indicate mark loss rate for adults since the second year after marking. The true simulated mark loss rate is indicated by the horizontal dashed line. The panel title indicates the simulated mark loss rate during the year following marking, for subsequent years this rate was set to 0.05 for all scenarios.

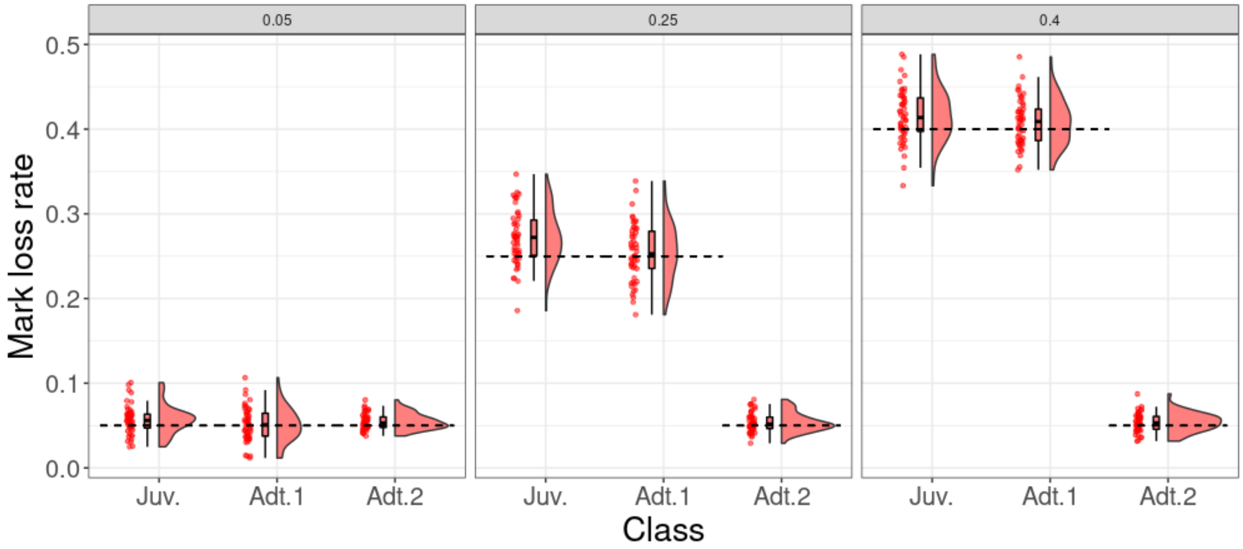

Figure S73: Probability distribution of the estimated mark loss rate for high survival and low detection (scenario 2). "Juv." and "Adt.1" indicate the mark loss rate during the year following marking for juveniles and adults respectively. "Adt.2" indicate mark loss rate for adults since the second year after marking. The true simulated mark loss rate is indicated by the horizontal dashed line. The panel title indicates the simulated mark loss rate during the year following marking, for subsequent years this rate was set to 0.05 for all scenarios.

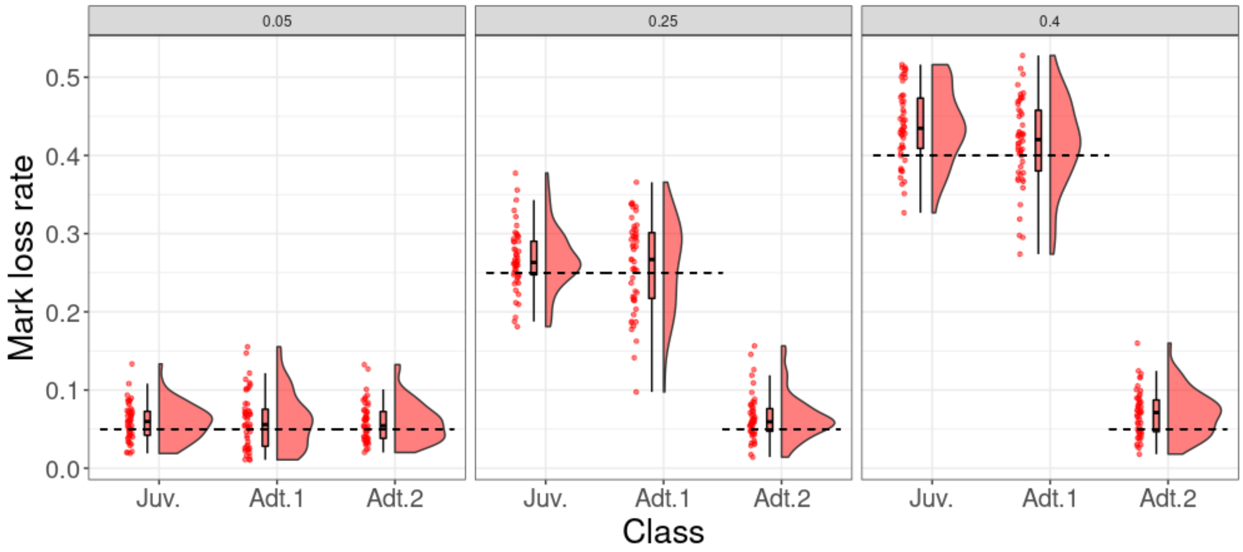

Figure S74: Probability distribution of the estimated mark loss rate for low survival and low detection (scenario 3). "Juv." and "Adt.1" indicate the mark loss rate during the year following marking for juveniles and adults respectively. "Adt.2" indicate mark loss rate for adults since the second year after marking. The true simulated mark loss rate is indicated by the horizontal dashed line. The panel title indicates the simulated mark loss rate during the year following marking, for subsequent years this rate was set to 0.05 for all scenarios.

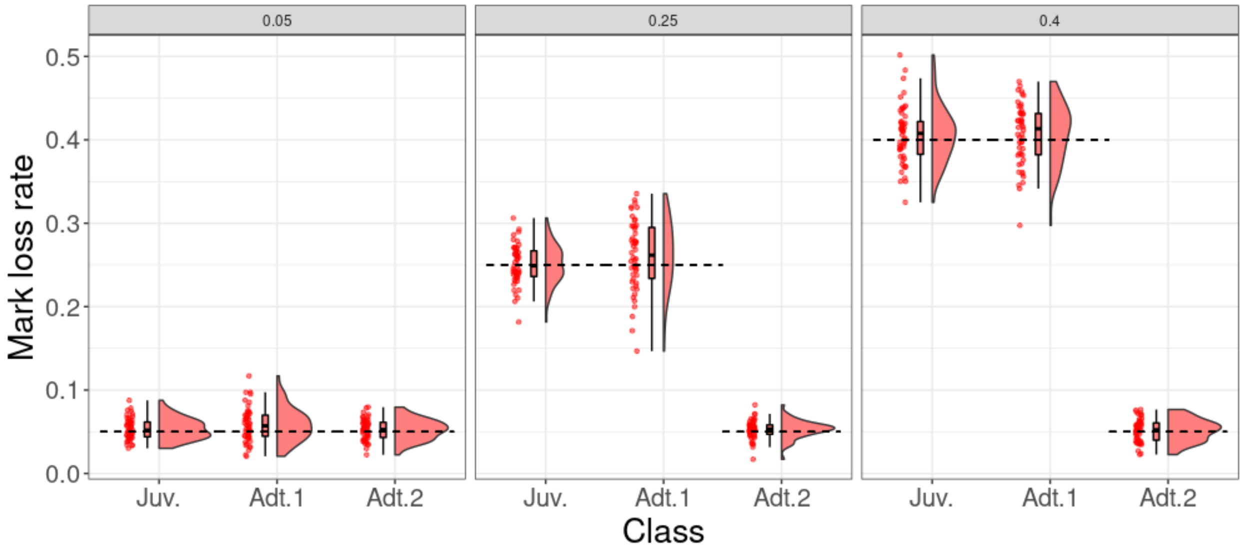

Figure S75: Probability distribution of the estimated mark loss rate for low survival and high detection (scenario 4). "Juv." and "Adt.1" indicate the mark loss rate during the year following marking for juveniles and adults respectively. "Adt.2" indicate mark loss rate for adults since the second year after marking. The true simulated mark loss rate is indicated by the horizontal dashed line. The panel title indicates the simulated mark loss rate during the year following marking, for subsequent years this rate was set to 0.05 for all scenarios.

### 4 ROPE estimation for EMDs

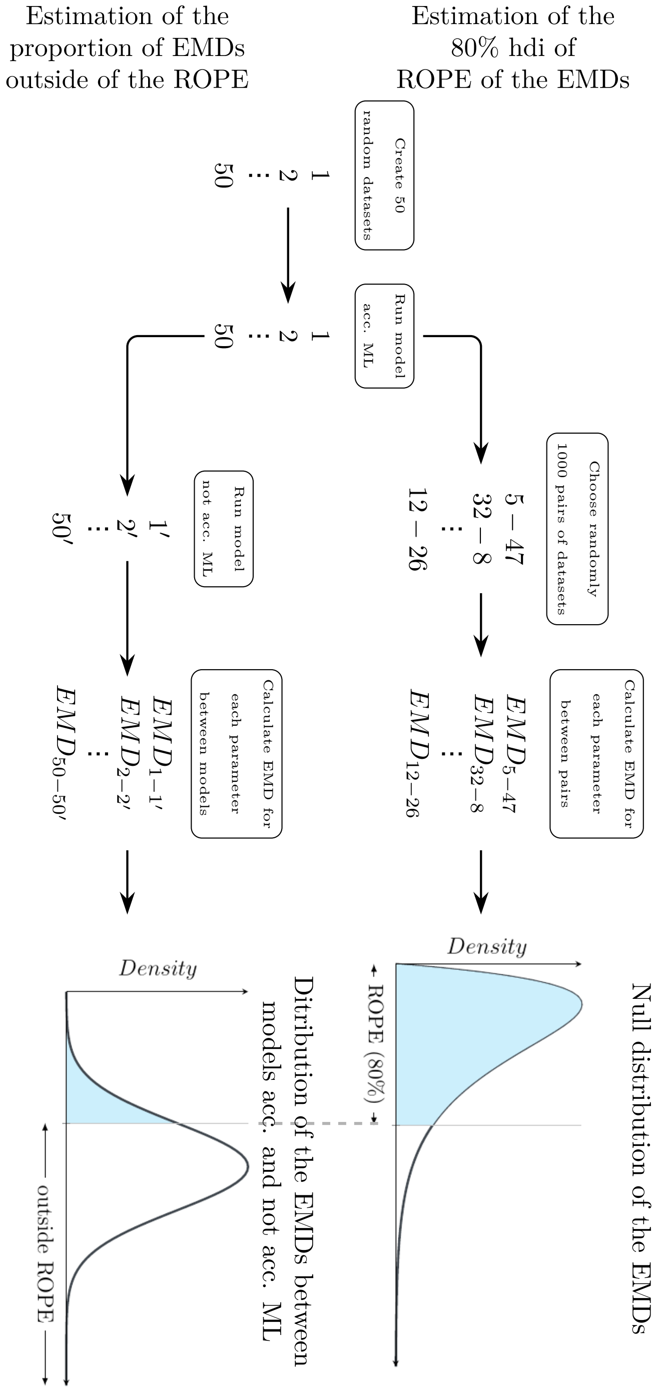

Figure S76: **Flowchart to estimate the proportion of the Earth Mover Distance (EMD)) outside of the Region Of Practical Equivalence (ROPE) between model accounting for mark loss (acc. ML) and not accounting for mark loss (not acc. ML).** The top part of the chart shows the estimation of the ROPE of the EMD. The bottom part of the chart illustrates how the proportion of the EMDs outside the Rope was estimated in the simulation framework. This flowchart shows the procedure which was repeated for each parameter estimated (*cf.* main text for more explanations).

Kéry, Marc, and Michael Schaub. 2012. *Bayesian Population Analysis Using Winbugs: A Hierarchical Perspective.* Waltham, MA, USA: Academic Press Inc.

Conn, Paul B., Kendall, William L. and Samuel, Michael D. 2004. *A General Model for the Analysis of Mark-Resight, Mark-Recapture, and Band-Recovery Data under Tag Loss*, Biometrics, 60(4), 900–909. https://onlinelibrary.wiley.com/doi/abs/10.1111/j.0006-341X.2004.00245.x.

Diefenbach, Duane R. and Alt, Gary L. 1998. *Modeling and Evaluation of Ear Tag Loss in Black Bears*, The Journal of Wildlife Management, 62(4), 1292–1300. https://www.jstor.org/stable/3801993.
