## Supplementary Information 3 for "Mark loss can strongly bias demographic rates in multi-state models: a case study with simulated and empirical datasets"

3 **Tag loss rate in the long term** 14

### 1 Real data: complementary information and model code

#### 1.1 Study design

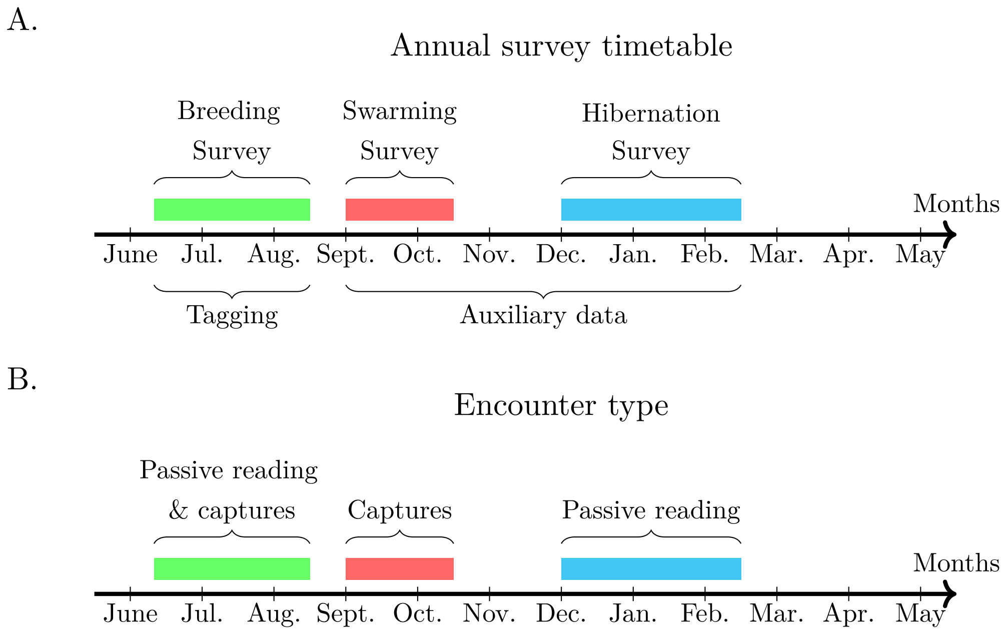

Figure S1: **Diagram of the bat sampling timetable (A) and observation process (B).**

#### 1.2 Tag loss data

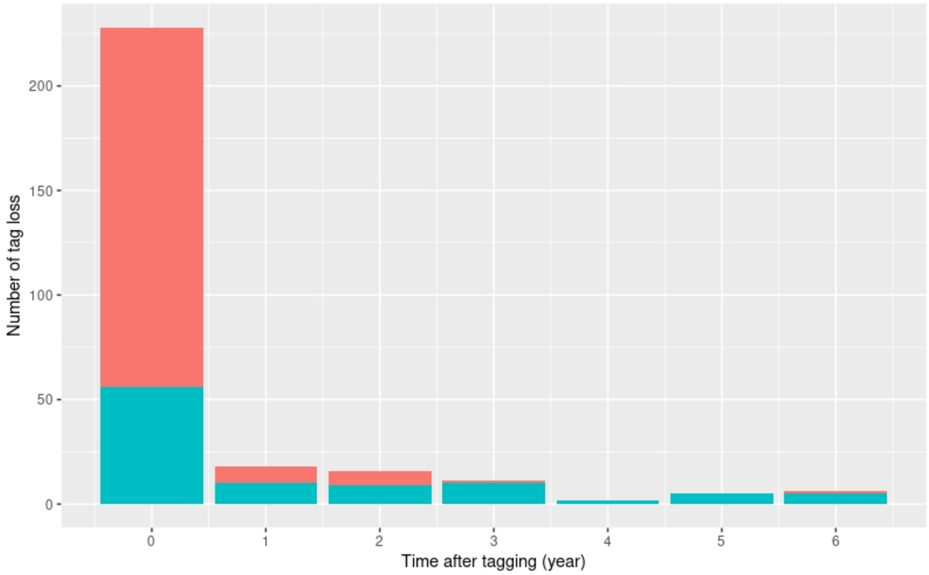

Figure S2: Distribution of tag loss from tagging occasion. Tag loss was assume to occur the day after the last antenna record. The red bars correspond to tag loss identified from PIT-tags found in the breeding sites and the blue from genotypes (recaptured individuals).

#### 1.3 AS model for data analyse

##### 1.3.1 Details on detection process

| Observation class | Probability |
|---|---|
| (1) Capture in colony only | $pc * (1 - m * p.r) - pb + m * p.r * pb$ |
| (2) Read in colony only | $m * p.r * (1 - pc) - pb + pc * pb$ |
| (3) Capture and read in colony | $pc * m * p.r * (1 - pb)$ |
| (4) Observed outside only | $pb * (1 - pc - m * p.r) + pc * m * p.r$ |
| (5) Observed outside and capture in colony | $pb * pc * (1 - m * p.r)$ |
| (6) Observed outside and read in colony | $pb * m * p.r * (1 - pc)$ |
| (7) Observed oumide, capture and read in colony | $pb * pc * m * p.r$ |
| (8) Not observed | $(1 - pc) * (1 - m * p.r) * (1 - pb)$ |

Table S1: Categorical detection probabilities. $pc$ = capture probability, $p.r$ = reading probability, $pb$ = dectected between occasions. $m$ is the tag state, indicating if the tag was retained with value 1 or 0 if it was lost.

##### 1.3.2 Details on state process

For the empirical data, we allowed all possible movements between the 5 colonies studied, but estimated them by sex and age class. We also add another virtual area that illustrate the departure of individuals outside colonies studied thanks to data collected in other sites during swarming and wintering period. Transition probability from this area to colonies is set to 0 for juveniles, as they are only marked in the 5 colonies studied (Fig. S3).

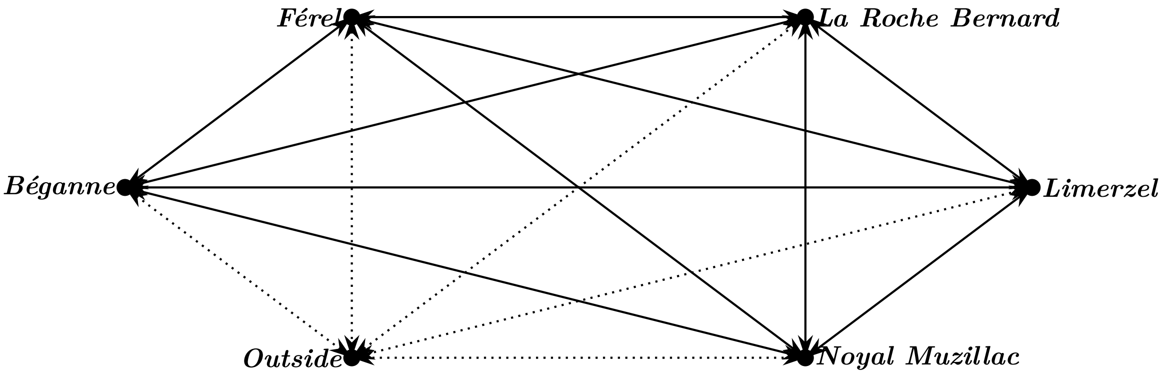

Figure S3: **Colonies and possible movements between them.**

##### 1.3.3 JAGS code for model fitting

```
model{

  for(i in 1:nind){
    for(j in (f[i]+1):l[i]){
      ap[i,j] <- a[i,j] + 1 # indicator variable (1=dead, 2=alive)
      # This specifies the distribution [a|S]
      a[i,j] ~ dbern(sv[i,j-1])
      sv[i,j-1] <- a[i,j-1]*phi[i,j-1]
      # mvts between colonies
      col[i,j] ~ dcat(psi[col[i,j-1],sex[i],age[i,j-1],1:ncol])
      # Detection process
      z[i,j] ~ dcat(pdet[ap[i,j],i,j-1,1:8])
      # indicator variable if (1=marked, 2=not marked)
      tm[i,j-1] <- 2-TM[i,j-1]
      # Tag state: 1 = retained, 0 = lost
      TR[i,j] ~ dbern(pret[age[i,j-1],tm[i,j-1],i,j-1])
    }
  }

  for(i in 1:nind){
    # presence or absence of a tag in the individual
    m[i,f[i]] <- 1 # indicator variable (1 at first tagging)
    for(j in (f[i]+1):l[i]){
      m[i,j] <- TR[i,j] + TM[i,j]  # indicator variable (1=tag present, 0=tag absent)
     }
  }

  for(i in 1:nind){
    for(j in (f[i]+1):l[i]){
      # detection probabilities
      pdet[1,i,j-1,1] <- 0
      pdet[1,i,j-1,2] <- 0
      pdet[1,i,j-1,3] <- 0
      pdet[1,i,j-1,4] <- 0
      pdet[1,i,j-1,5] <- 0
      pdet[1,i,j-1,6] <- 0
```

```
    pdet[1,i,j-1,7] <- 0
    pdet[1,i,j-1,8] <- 1

    pdet[2,i,j-1,1] <- ifelse(col[i,j]==6,0,1)*p.c[col[i,j],j-1]*(1 - m[i,j]*
      p.r[col[i,j],j-1])*(1 - pb) # capture only
    pdet[2,i,j-1,2] <- ifelse(col[i,j]==6,0,1)*m[i,j]*p.r[col[i,j],j-1]*(1 -
      p.c[col[i,j],j-1])*(1 - pb) # antenna only
    pdet[2,i,j-1,3] <- ifelse(col[i,j]==6,0,1)*p.c[col[i,j],j-1]*m[i,j]*
      p.r[col[i,j],j-1]*(1 - pb) # capture and antenna
    pdet[2,i,j-1,4] <- ifelse(col[i,j]==6, pb, (1 - p.c[col[i,j],j-1])*(1 -
      m[i,j]*p.r[col[i,j],j-1])*pb) # observed between season only
    pdet[2,i,j-1,5] <- ifelse(col[i,j]==6,0,1)*p.c[col[i,j],j-1]*(1 - m[i,j]*
      p.r[col[i,j],j-1])*pb # observed between season and captured
    pdet[2,i,j-1,6] <- ifelse(col[i,j]==6,0,1)*(1- p.c[col[i,j],j-1])*m[i,j]*
      p.r[col[i,j],j-1]*pb # observed between season and with antenna
    pdet[2,i,j-1,7] <- ifelse(col[i,j]==6,0,1)*p.c[col[i,j],j-1]*m[i,j]*
      p.r[col[i,j],j-1]*pb # observed between season, captured and read
    pdet[2,i,j-1,8] <- ifelse(col[i,j]==6,1 - pb, (1- p.c[col[i,j],j-1])*(1 -
      m[i,j]*p.r[col[i,j],j-1])*(1 - pb)) # not observed

    # Tag retention probability
    pret[1,1,i,j-1] <- pr[1, glu[i,j-1]]
    pret[1,2,i,j-1] <- pr[2, glu[i,j-1]]
    pret[2,1,i,j-1] <- pr[3, glu[i,j-1]]
    pret[2,2,i,j-1] <- TR[i,j-1]*pr[4, glu[i,j-1]]
  }

  for(j in f[i]:(l[i]-1)){
    # survival probability
    logit(phi[i,j]) <- alpha[col[i,j]] + beta[j]  + delta[age[i,j]] +
      gamma[col[i,j],j,age[i,j]] + eta.s[i]
  }
}

for(c in 1:(ncol-1)){
  for(j in 1:(nocc-1)){
    # mean detection probability in colonies
    pd[c,j] <- p.c[c,j] + p.r[c,j] - p.c[c,j]*p.r[c,j]
  }
}

### Priors and constraints
for(j in 1:(nocc-1)){
  # detcetion and capture probabilities when out of colony is null
  p.c[6,j] <- 0
  pd[6,j] <- 0
  p.r[6,j] <- 0
}

# tag retention probability
for(g in 1:2){# g=1 no surgical adhesive ; g=2 with surgical adhesive
  pr[1,g] ~ dbeta(1,1) # retention for juveniles after tagging
  pr[2,g] <- 0
```

```
  pr[3,g] ~ dbeta(1,1) # retention for adults after tagging
  pr[4,g] ~ dbeta(1,1) # retention for adults if retained after tagging
}

for(c in c(1:3)){
  for(j in 1:(nocc-1)){
  logit(p.c[c,j]) <- alpha.c[c] + beta.c[j] + gamma.c[c,j]
  }
}

# No catch in colony 4 in 2013 & 2014
p.c[4,3] <- 0
p.c[4,4] <- 0
for(j in c(1,2,5:(nocc-1))){
  logit(p.c[4,j]) <- alpha.c[4] + beta.c[j] + gamma.c[4,j]
}

# No catch in colony 5 before 2013
for (j in 1:3){
  p.c[5,j] <- 0
}
for(j in 4:(nocc-1)){
  logit(p.c[5,j]) <- alpha.c[5] + beta.c[j] + gamma.c[5,j]
}

# No reader in colony 1 & 2 before 2012
p.r[1,1] <- 0
p.r[2,1] <- 0
for (c in 1:2){
  for (j in 2:(nocc-1)){
    p.r[c,j] ~ dbeta(1,1)
  }
}

# No reader in colony 3 before 2018
for (j in 1:7){
  p.r[3,j] <- 0
}
p.r[3,8] ~ dbeta(1,1)
p.r[3,9] ~ dbeta(1,1)

# No reader in colony 4
for (j in 1:(nocc-1)){
  p.r[4,j] <- 0
}

# No reader in colony 5 before 2017
for (j in 1:6){
  p.r[5,j] <- 0
}
for (j in 7:(nocc-1)){
  p.r[5,j] ~ dbeta(1,1)
}
```

```
pb ~ dbeta(1,1)

for(c in 1:ncol){
  alpha[c] ~ dnorm(0,0.01)T(-10,10)
}

for(c in 1:(ncol-1)){
  alpha.c[c] ~ dnorm(0,0.01)T(-10,10)
}

beta[1] <- 0
beta.c[1] <- 0
delta[1] <- 0
delta[2] ~ dt(0,0.16,3)

for(j in 2:(nocc-1)){
  beta[j] ~ dt(0,0.16,3)
  beta.c[j] ~ dt(0,0.16,3)
}

for(c in 1:ncol){
  for(j in 1:(nocc-1)){
    for(a in 1:2){
      gamma[c,j,a] ~ dt(0,0.16,3)
    }
  }
}

for(c in 1:(ncol-1)){
  gamma.c[c,1] <- 0
}

for(j in 2:(nocc-1)){
  gamma.c[1,j] <- 0
  for(c in 2:(ncol-1)){
    gamma.c[c,j] ~ dt(0,0.16,3)
  }
}

tau.s <- pow(sigma.s,-2)
sigma.s ~ dunif(0,2)

for(i in 1:nind){
  eta.s[i] ~ dnorm(0, tau.s)
}

# movement between colony
for(c in 1:ncol){
  for(s in 1:2){
    psi[c,s,2,1:6] ~ ddirch(alpha.psi[])
  }
}
```

```
for(c in 1:(ncol-1)){
  for(s in 1:2){
    psi[c,s,1,1:6] ~ ddirch(alpha.psi[])
  }
}

for(c in 1:ncol){
  for(s in 1:2){
    psi[6,s,1,c] <- 0
  }
}

for(c in 1:ncol){
  alpha.psi[c] <- 1
  }
}
```

### 2 Results

The following graphics display the Earth Mover Distance (EMD) between the posterior density distribution of parameters estimated from model accounting for tag loss and recycling and model without. The difference between the medians of the posterior distribution of the parameters from the two models are also displayed.

#### 2.1 Juvenile survival

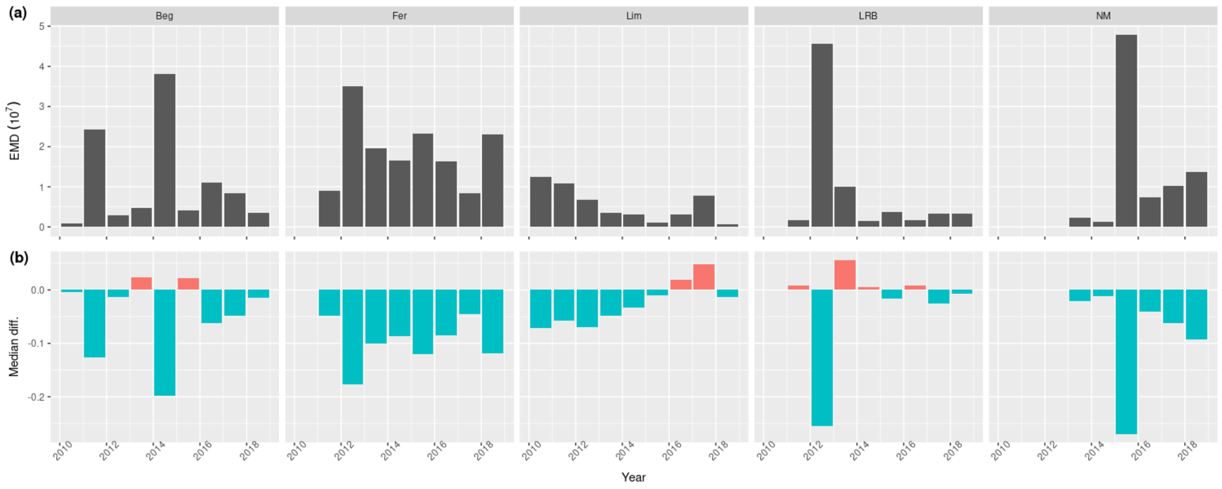

Figure S4: **Assessment of the difference in annual juvenile survival estimates**. EMD (a) and difference between medians (b) of the posterior distributions of survival between model accounting for tag loss and recycling and model without. The blue bars correspond to an underestimate and the red bars to an overestimate of the survival rate by the model ignoring the loss and recycling of the labels relative to the other. Colonies: Beg = Beganne; Fer = Férel; Lim = Limerzel; LRB = La Roche Bernard; NM = Noyal-Muzillac.

##### 2.2 Adult survival

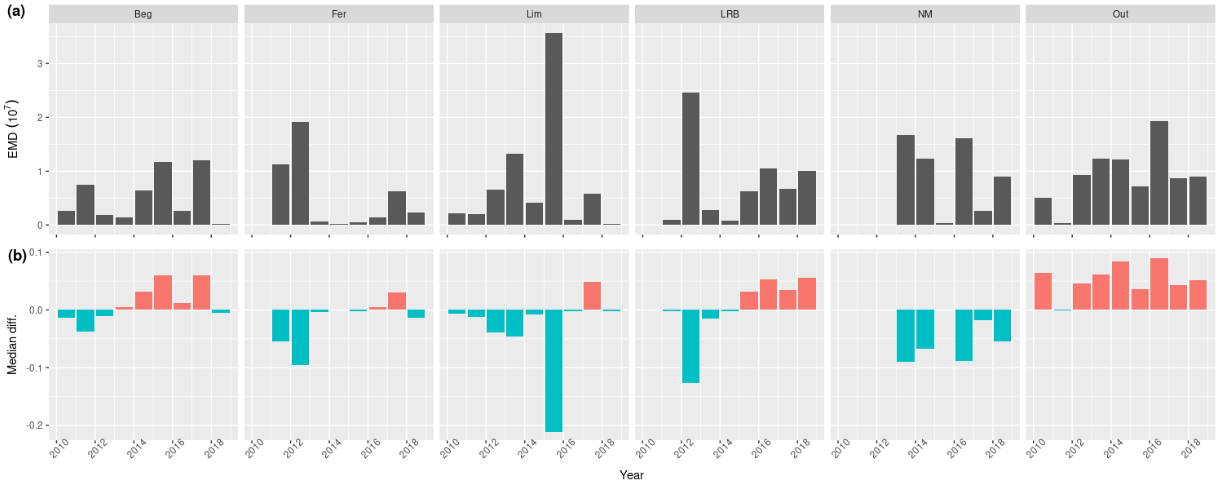

Figure S5: **Assessment of the difference in annual adult survival estimates**. EMD (a) and difference between the medians (b) of the posterior distributions of survival between model accounting for tag loss and recycling and model without. The blue bars correspond to an underestimate and the red bars to an overestimate of the survival rate by the model ignoring the loss and recycling of the labels relative to the other. Colonies: Beg = Beganne; Fer = Férel; Lim = Limerzel; LRB = La Roche Bernard; NM = Noyal-Muzillac; Out = swarming and wintering sites.

##### 2.3 Capture probability

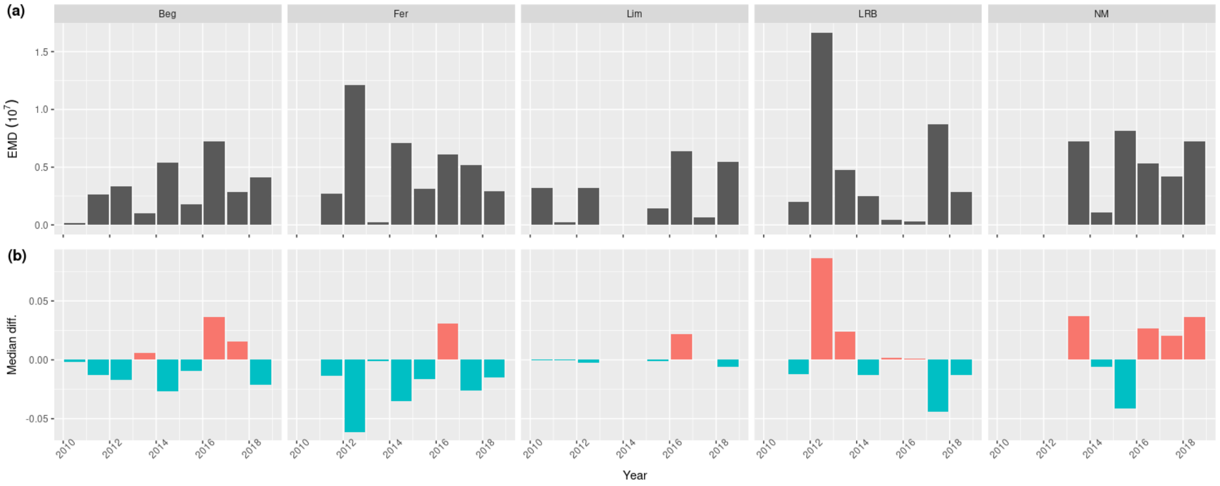

Figure S6: **Assessment of the difference in annual capture probability estimates**. EMD (a) and difference between the medians (b) of the posterior distributions of capture rate between model accounting for tag loss and recycling and model without. The blue bars correspond to an underestimate and the red bars to an overestimate of the survival rate by the model ignoring the loss and recycling of the labels relative to the other. Colonies: Beg = Beganne; Fer = Férel; Lim = Limerzel; LRB = La Roche Bernard; NM = Noyal-Muzillac.

#### 2.4 Resighting probability

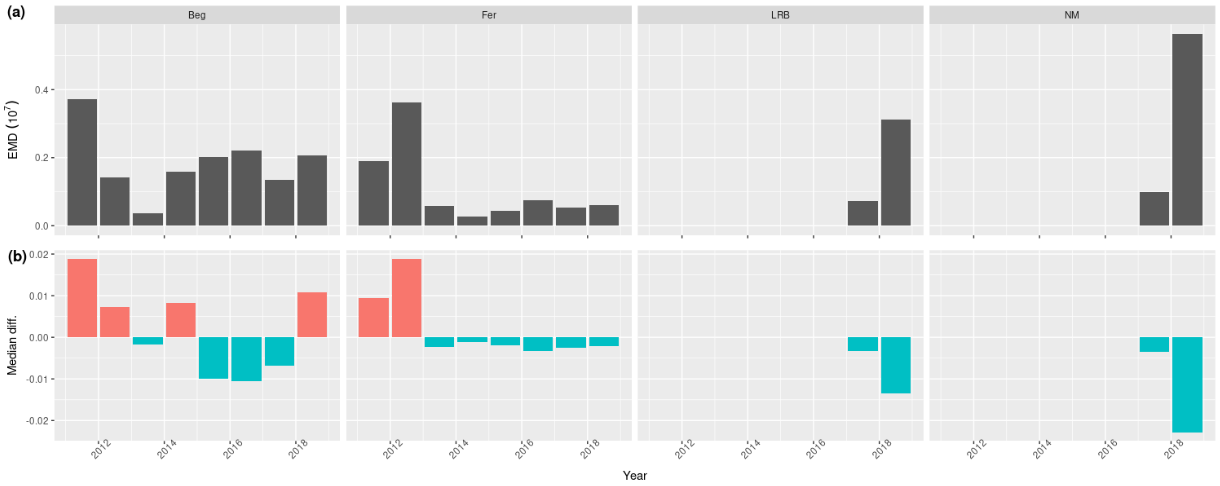

Figure S7: **Resighting probabilities**. EMD (a) and difference between the medians (b) of the posterior densities of resighting rate between model accounting for tag loss and recycling and model without. The blue bars correspond to an underestimate and the red bars to an overestimate of the survival rate by the model ignoring the loss and recycling of the labels relative to the other. Colonies: Beg = Beganne; Fer = Férel; Lim = Limerzel; LRB = La Roche Bernard; NM = Noyal-Muzillac.

#### 2.5 Movement probability

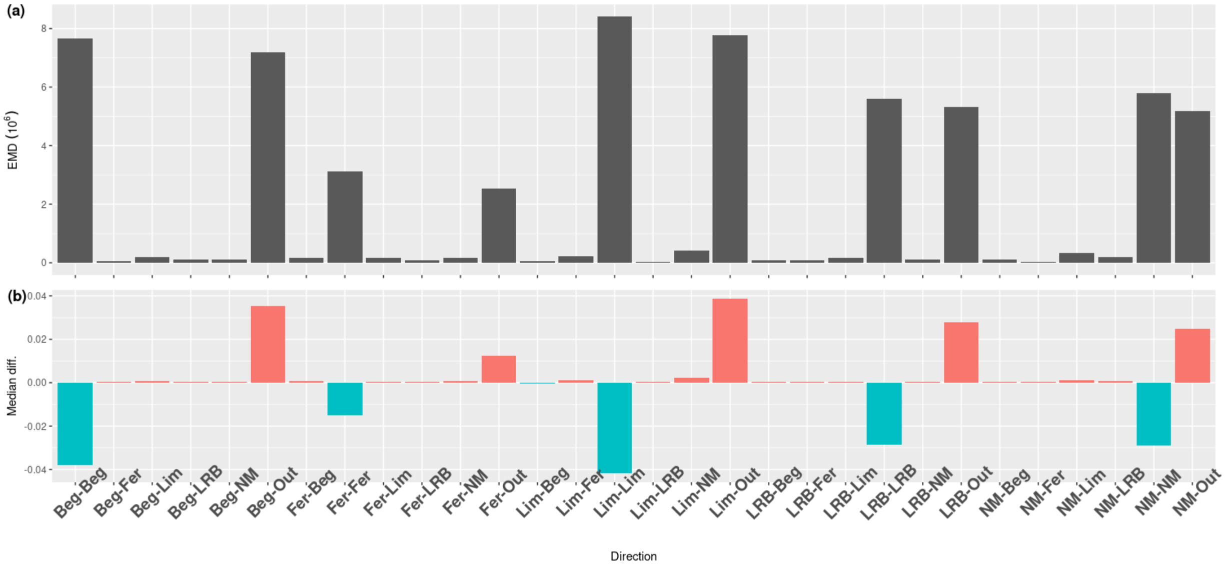

Figure S8: **Assessment of the difference in estimates of movements of juvenile females**. EMD (a) and difference between the medians (b) of the posterior distributions of movement rates between model accounting for tag loss and recycling and model without. The blue bars correspond to an underestimate and the red bars to an overestimate of the survival rate by the model ignoring the loss and recycling of the labels relative to the other. Colonies: Beg = Béganne, Fer = Férel, LRB = La Roche Bernard, Lim = Limerzel, NM = Noyal Muzillac, Out = swarming and wintering sites.

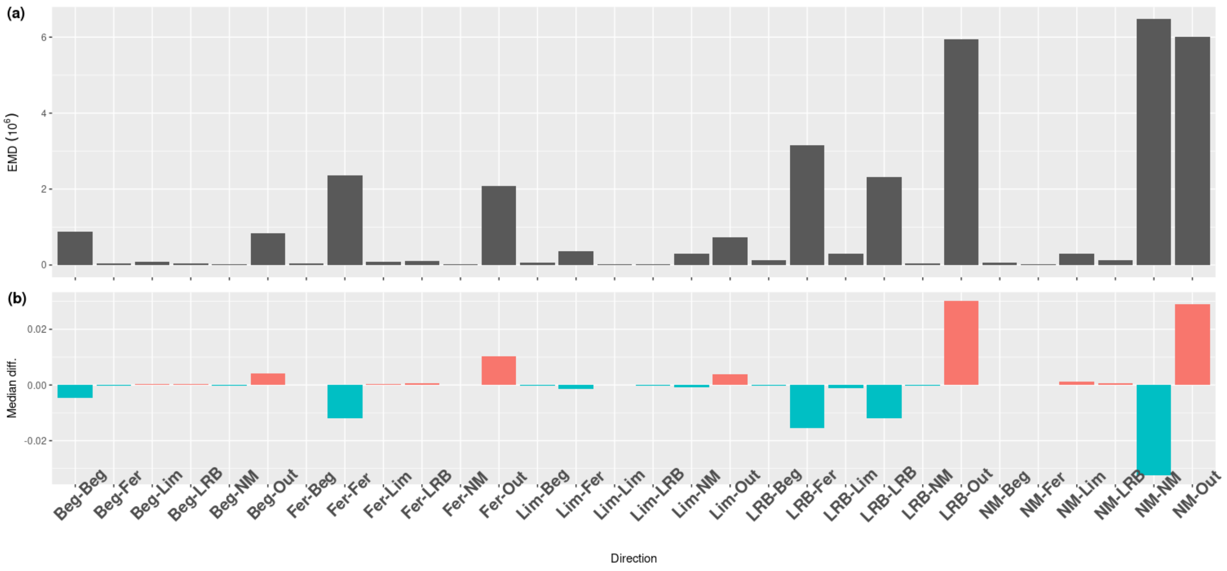

Figure S9: **Assessment of the difference in estimates of movements of juvenile males**.EMD (a) and difference between the medians (b) of the posterior distributions of movement rates between model accounting for tag loss and recycling and model without. The blue bars correspond to an underestimate and the red bars to an overestimate of the survival rate by the model ignoring the loss and recycling of the labels relative to the other. Colonies: Beg= Béganne, Fer=Férel, LRB=La Roche Bernard, Lim=Limerzel, NM=Noyal Muzillac, Out=swarming and wintering sites.

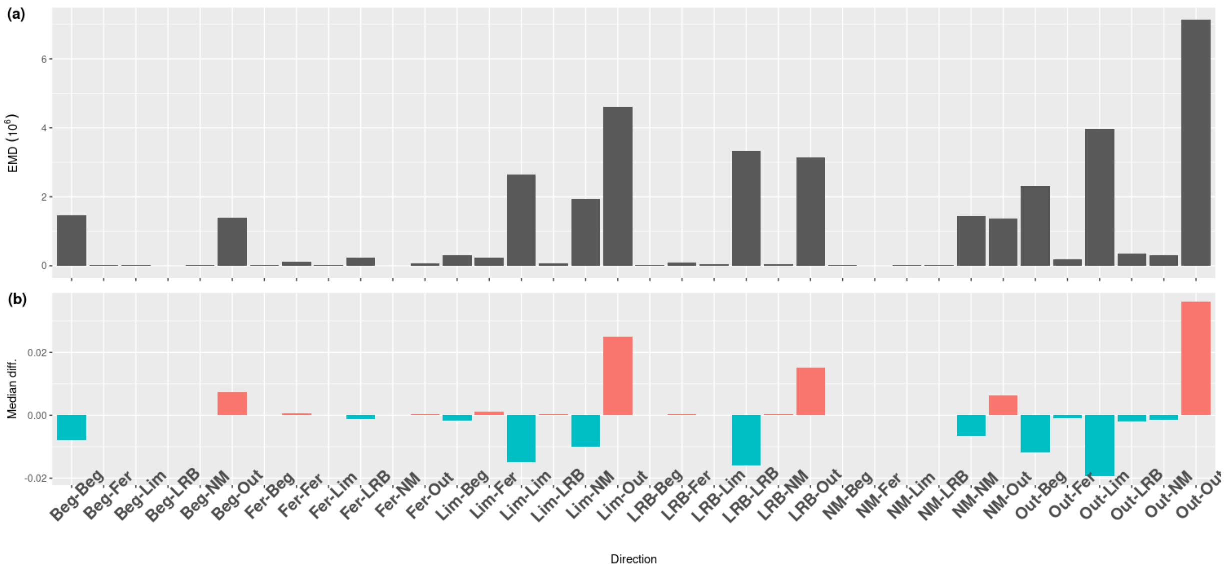

Figure S10: **Assessment of the difference in estimates of movements of adult females**. EMD (a) and difference between the medians (b) of the posterior distributions of movement rates between model accounting for tag loss and recycling and model without. The blue bars correspond to an underestimate and the red bars to an overestimate of the survival rate by the model ignoring the loss and recycling of the labels relative to the other. Colonies: Beg= Béganne, Fer=Férel, LRB=La Roche Bernard, Lim=Limerzel, NM=Noyal Muzillac, Out=swarming and wintering sites.

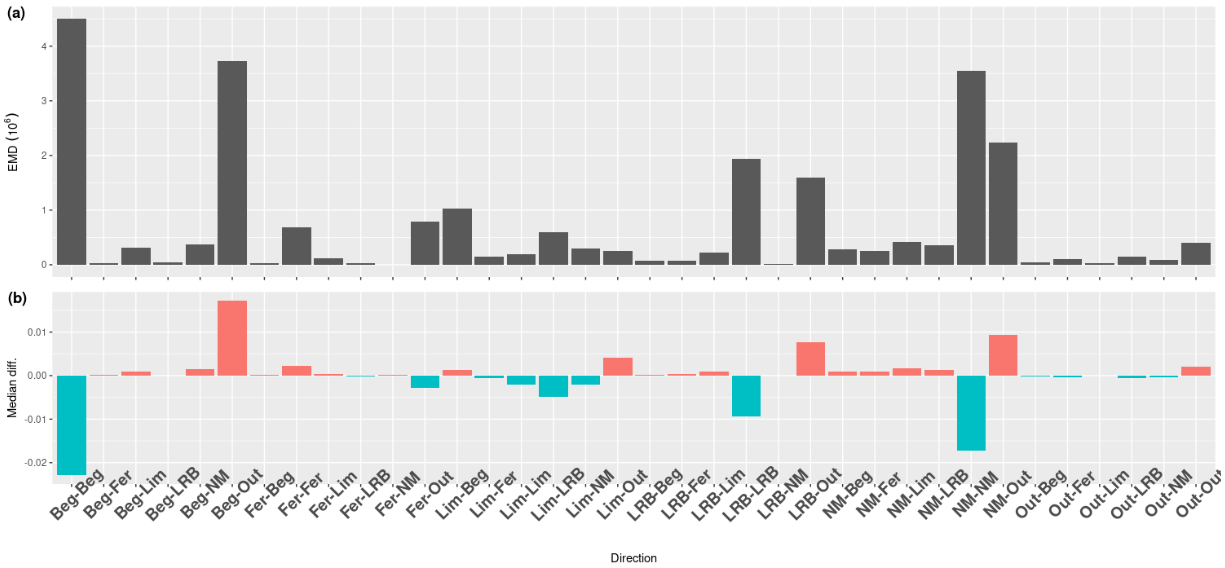

Figure S11: **Assessment of the difference in estimates of movements of adult Males**. EMD (a) and difference between the medians (b) of the posterior distributions of movement rates between model accounting for tag loss and recycling and model without. The blue bars correspond to an underestimate and the red bars to an overestimate of the survival rate by the model ignoring the loss and recycling of the labels relative to the other. Colonies: Beg= Béganne, Fer=Férel, LRB=La Roche Bernard, Lim=Limerzel, NM=Noyal Muzillac, Out=swarming and wintering sites.

#### 2.6 Estimated values for model accounting for tag loss and recycling

| Year | Béganne | Férel | Limerzel | La Roche Bernard | Noyal-Muzillac |
|---|---|---|---|---|---|
| 2010-11 | 0.67 [0.46-0.86] | | 0.71 [0.46-1] | | |
| 2011-12 | 0.6 [0.44-0.77] | 0.93 [0.82-1] | 0.49 [0.18-0.84] | 0.83 [0.33-1] | |
| 2012-13 | 0.14 [0.05-0.25] | 0.42 [0.22-0.62] | 0.28 [0-0.97] | 0.71 [0.45-0.96] | |
| 2013-14 | 0.27 [0.12-0.43] | 0.65 [0.46-0.83] | 0.66 [0.02-1] | 0.53 [0.21-0.88] | 0.7 [0.42-1] |
| 2014-15 | 0.71 [0.52-0.9] | 0.61 [0.45-0.79] | 0.88 [0.08-1] | 0.97 [0.8-1] | 0.91 [0.66-1] |
| 2015-16 | 0.1 [0.02-0.22] | 0.6 [0.45-0.75] | 0.65 [0.36-0.97] | 0.96 [0.82-1] | 0.62 [0.34-1] |
| 2016-17 | 0.76 [0.53-0.96] | 0.82 [0.69-0.93] | 0.75 [0.36-1] | 0.75 [0.42-1] | 0.94 [0.75-1] |
| 2017-18 | 0.88 [0.7-1] | 0.87 [0.75-0.98] | 0.45 [0.13-1] | 0.82 [0.56-1] | 0.82 [0.61-1] |
| 2018-19 | 0.99 [0.93-1] | 0.96 [0.88-1] | 0.76 [0.13-1] | 1 [0.95-1] | 0.86 [0.55-1] |

Table S2: **Annual juvenile survival**. Median and 90% hdi, in bracket, of the posterior density probabilities.

| Year | Béganne | Férel | Limerzel | La Roche Bernard | Noyal-Muzillac | Outside |
|---|---|---|---|---|---|---|
| 2010-11 | 0.78 [0.66-0.89] | | 0.99 [0.91-1] | | | 0.58 [0.02-1] |
| 2011-12 | 0.82 [0.74-0.9] | 0.98 [0.94-1] | 0.5 [0.29-0.71] | 0.97 [0.82-1] | | 0.97 [0.8-1] |
| 2012-13 | 0.76 [0.67-0.84] | 0.74 [0.64-0.83] | 0.42 [0.08-0.84] | 0.88 [0.76-0.99] | | 0.47 [0.21-0.76] |
| 2013-14 | 0.63 [0.5-0.74] | 0.84 [0.76-0.91] | 0.92 [0.45-1] | 0.84 [0.7-0.97] | 0.84 [0.69-0.98] | 0.22 [0.08-0.39] |
| 2014-15 | 0.55 [0.4-0.7] | 0.84 [0.76-0.91] | 0.97 [0.66-1] | 1 [0.97-1] | 0.95 [0.82-1] | 0.77 [0.42-1] |
| 2015-16 | 0.44 [0.28-0.61] | 0.64 [0.53-0.75] | 0.95 [0.76-1] | 0.49 [0.29-0.71] | 0.7 [0.5-0.9] | 0.3 [0.12-0.51] |
| 2016-17 | 0.54 [0.34-0.74] | 0.61 [0.5-0.72] | 0.86 [0.52-1] | 0.56 [0.3-0.81] | 0.91 [0.78-1] | 0.35 [0.13-0.6] |
| 2017-18 | 0.54 [0.33-0.75] | 0.59 [0.46-0.7] | 0.22 [0-0.87] | 0.77 [0.53-1] | 0.88 [0.76-1] | 0.25 [0.09-0.43] |
| 2018-19 | 0.92 [0.77-1] | 0.74 [0.62-0.85] | 0.99 [0.83-1] | 0.72 [0.45-1] | 0.93 [0.81-1] | 0.53 [0.26-1] |

Table S3: **Annual adult survival**. Median and 90% hdi , in bracket, of the posterior density probabilities.

| Year | Béganne | Férel | Limerzel | La Roche Bernard | Noyal-Muzillac |
|---|---|---|---|---|---|
| 2011 | 0.88 [0.81-0.95] | | 1 [0.92-1] | | |
| 2012 | 0.89 [0.84-0.93] | 0.54 [0.43-0.63] | 1 [0.89-1] | 0.74 [0.54-0.93] | |
| 2013 | 0.72 [0.66-0.78] | 0.58 [0.51-0.67] | 0.99 [0.31-1] | 0.85 [0.75-0.94] | |
| 2014 | 0.85 [0.8-0.9] | 0.45 [0.38-0.52] | | 0.74 [0.63-0.85] | 0.59 [0.46-0.73] |
| 2015 | 0.61 [0.53-0.69] | 0.65 [0.58-0.71] | | 0.83 [0.74-0.92] | 0.55 [0.42-0.68] |
| 2016 | 0.72 [0.63-0.8] | 0.57 [0.5-0.63] | 1 [0.93-1] | 0.87 [0.77-0.96] | 0.52 [0.4-0.64] |
| 2017 | 0.44 [0.35-0.54] | 0.36 [0.3-0.41] | 0.46 [0.25-1] | 0.71 [0.57-0.85] | 0.27 [0.18-0.35] |
| 2018 | 0.82 [0.75-0.9] | 0.45 [0.39-0.51] | 1 [0.86-1] | 0.41 [0.28-0.54] | 0.2 [0.13-0.28] |
| 2019 | 0.73 [0.66-0.8] | 0.41 [0.35-0.47] | 0.99 [0.71-1] | 0.47 [0.33-0.61] | 0.2 [0.13-0.27] |

Table S4: **Annual capture probability**. Median and 90% hdi, in bracket, of the posterior density probabilities.

| Year | Beg | Fer | LRB | NM |
|---|---|---|---|---|
| 2012 | 0.97 [0.94-0.99] | 0.95 [0.9-0.99] | | |
| 2013 | 0.98 [0.95-1] | 0.98 [0.95-1] | | |
| 2014 | 0.95 [0.92-0.98] | 0.99 [0.97-1] | | |
| 2015 | 0.94 [0.9-0.98] | 0.99 [0.97-1] | | |
| 2016 | 0.93 [0.87-0.98] | 0.99 [0.97-1] | | |
| 2017 | 0.96 [0.91-0.99] | 0.99 [0.97-1] | | |
| 2018 | 0.95 [0.9-0.99] | 0.98 [0.97-1] | 0.98 [0.93-1] | 0.99 [0.96-1] |
| 2019 | 0.99 [0.96-1] | 0.99 [0.97-1] | 0.96 [0.87-1] | 0.96 [0.88-1] |

Table S5: **Annual resighting probability**. Median and 90% hdi, in bracket, of the posterior density probabilities.

| Movement | Adult female | Juvenile female | Adults male | Juvenile male |
|---|---|---|---|---|
| Beg-Beg | **0.91** [0.89-0.94] | **0.81** [0.73-0.88] | **0.56** [0.42-0.7] | 0.14 [0.09-0.19] |
| Beg-Fer | 0 [0-0.01] | 0 [0-0.02] | 0.02 [0-0.06] | 0.01 [0-0.03] |
| Beg-Lim | 0 [0-0] | 0.01 [0-0.02] | 0.02 [0-0.07] | 0.01 [0-0.02] |
| Beg-LRB | 0 [0-0] | 0.01 [0-0.02] | 0.02 [0-0.06] | 0.01 [0-0.02] |
| Beg-NM | 0 [0-0] | 0.01 [0-0.02] | 0.02 [0-0.08] | 0.01 [0-0.02] |
| Beg-Out | 0.08 [0.05-0.1] | 0.16 [0.09-0.24] | 0.32 [0.17-0.47] | **0.82** [0.76-0.87] |
| Fer-Beg | 0 [0-0.01] | 0.01 [0-0.02] | 0.01 [0-0.02] | 0 [0-0.01] |
| Fer-Fer | **0.99** [0.98-0.99] | **0.92** [0.87-0.96] | **0.7** [0.62-0.78] | 0.26 [0.21-0.31] |
| Fer-Lim | 0 [0-0] | 0 [0-0.01] | 0.01 [0-0.03] | 0 [0-0.01] |
| Fer-LRB | 0 [0-0] | 0 [0-0.01] | 0.01 [0-0.02] | 0.01 [0-0.02] |
| Fer-NM | 0 [0-0] | 0.01 [0-0.02] | 0.01 [0-0.02] | 0 [0-0.01] |
| Fer-Out | 0.01 [0-0.01] | 0.05 [0.01-0.09] | 0.26 [0.18-0.34] | **0.72** [0.66-0.77] |
| Lim-Beg | 0.06 [0.03-0.09] | 0.06 [0.02-0.11] | 0.1 [0-0.31] | 0.01 [0-0.04] |
| Lim-Fer | 0.02 [0.01-0.04] | 0.04 [0.01-0.08] | 0.15 [0-0.39] | 0.03 [0-0.07] |
| Lim-Lim | 0.29 [0.21-0.37] | 0.17 [0.08-0.26] | 0.11 [0-0.32] | 0.01 [0-0.04] |
| Lim-LRB | 0 [0-0.01] | 0.01 [0-0.04] | 0.12 [0-0.35] | 0.01 [0-0.04] |
| Lim-NM | 0.02 [0-0.04] | 0.04 [0-0.08] | 0.13 [0-0.36] | 0.02 [0-0.06] |
| Lim-Out | **0.6** [0.51-0.69] | **0.66** [0.54-0.78] | 0.18 [0-0.46] | **0.9** [0.82-0.96] |
| LRB-Beg | 0 [0-0.01] | 0.01 [0-0.04] | 0.01 [0-0.03] | 0.01 [0-0.03] |
| LRB-Fer | 0.02 [0-0.03] | 0.08 [0.03-0.13] | 0.01 [0-0.03] | 0.04 [0.01-0.08] |
| LRB-Lim | 0 [0-0.01] | 0.01 [0-0.04] | 0.01 [0-0.04] | 0.01 [0-0.03] |
| LRB-LRB | **0.8** [0.74-0.87] | **0.7** [0.58-0.81] | **0.83** [0.74-0.92] | 0.14 [0.07-0.21] |
| LRB-NM | 0 [0-0.01] | 0.01 [0-0.04] | 0.01 [0-0.03] | 0.01 [0-0.04] |
| LRB-Out | 0.16 [0.1-0.23] | 0.16 [0.06-0.27] | 0.11 [0.03-0.2] | **0.77** [0.68-0.86] |
| NM-Beg | 0 [0-0.01] | 0.01 [0-0.04] | 0.01 [0-0.04] | 0.01 [0-0.04] |
| NM-Fer | 0 [0-0.01] | 0.01 [0-0.04] | 0.01 [0-0.04] | 0.01 [0-0.04] |
| NM-Lim | 0 [0-0.01] | 0.01 [0-0.05] | 0.01 [0-0.04] | 0.01 [0-0.04] |
| NM-LRB | 0 [0-0.01] | 0.01 [0-0.04] | 0.01 [0-0.04] | 0.01 [0-0.04] |
| NM-NM | **0.9** [0.85-0.95] | **0.76** [0.64-0.89] | **0.8** [0.68-0.91] | 0.2 [0.11-0.3] |
| NM-Out | 0.08 [0.04-0.13] | 0.16 [0.05-0.29] | 0.14 [0.03-0.24] | **0.73** [0.62-0.84] |
| Out-Beg | 0.07 [0.04-0.1] | 0 [0-0] | 0 [0-0] | 0 [0-0] |
| Out-Fer | 0.01 [0-0.02] | 0 [0-0] | 0 [0-0.01] | 0 [0-0] |
| Out-Lim | 0.14 [0.1-0.19] | 0 [0-0] | 0 [0-0.01] | 0 [0-0] |
| Out-LRB | 0.03 [0.01-0.05] | 0 [0-0] | 0.01 [0-0.01] | 0 [0-0] |
| Out-NM | 0.03 [0.01-0.04] | 0 [0-0] | 0 [0-0.01] | 0 [0-0] |
| Out-Out | **0.71** [0.65-0.77] | 0 [0-0] | **0.98** [0.97-0.99] | 0 [0-0] |

Table S6: **Movement probability**. Median and 90% hdi, in bracket, of the posterior density probabilities. Philopatry rate in bold type.

| P | Surgical glue | Age |
|---|---|---|
| 0.28 [0.23-0.33] | No | Juvenile |
| 0.11 [0.08-0.14] | No | Adult 1st |
| 0.02 [0.01-0.02] | No | Adult 2nd |
| 0.19 [0.16-0.22] | Yes | Juvenile |
| 0.1 [0.06-0.13] | Yes | Adult 1st |
| 0.03 [0.02-0.04] | Yes | Adult 2nd |

Table S7: **Tag loss probability**. Median and 90% hdi, in bracket, of the posterior density probabilities.

### 3 Tag loss rate in the long term

We were not able to collect tags from colonies at the beginning of the study, corresponding to the period when we tagged most of the adults for whom we did not use surgical glue to seal the injection wound. To check how this might have affected our estimates of the probability of long-term tag loss, we re-run our model selecting data only from 2013. The result (Fig. S12) no longer shows differences in long-term tag loss, confirming that the estimate made on the full data set reflects a bias in the tag loss data at the beginning of the study.

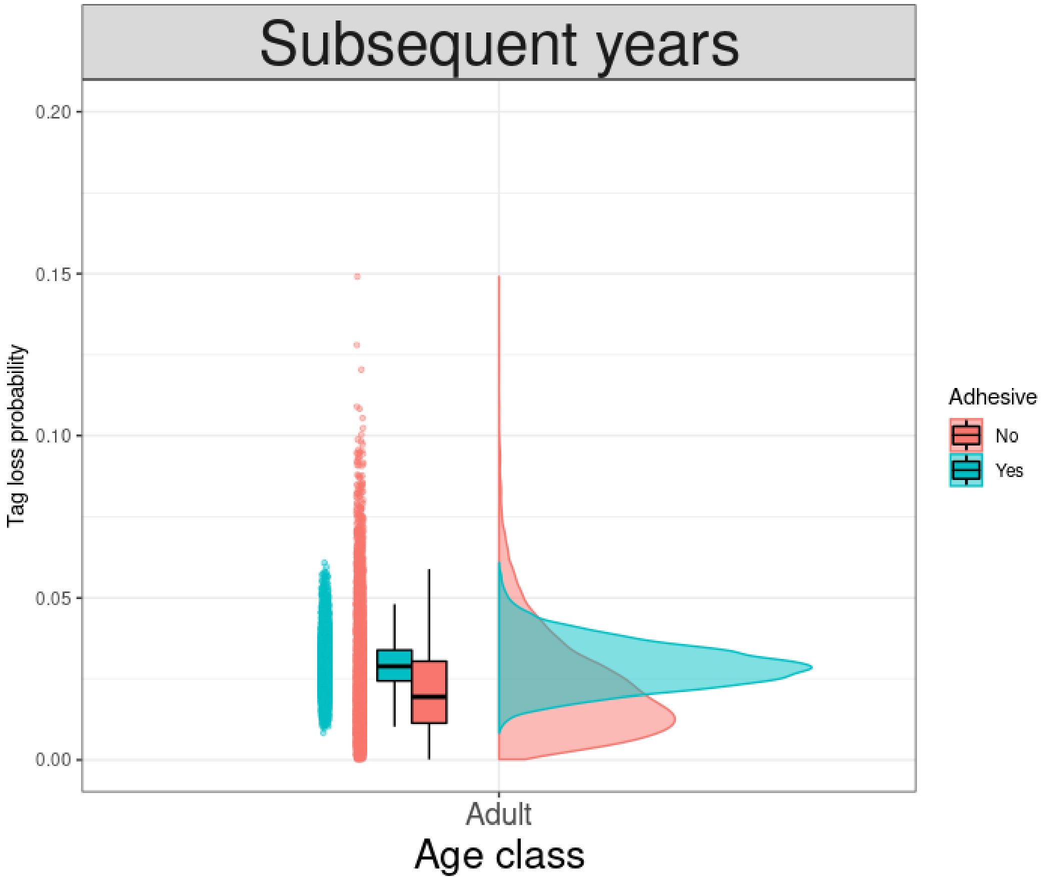

Figure S12: Posterior distribution of the tag loss probabilities according to age class from one year after marking. In blue, distribution if surgical adhesive was used after tag injection and in red, without surgical adhesive.
